# The Effect of Introgression of an R2R3 MYB Transcription Factor on Sulphur Metabolism in *Brassica oleracea*

**DOI:** 10.1101/2020.08.02.232819

**Authors:** Mikhaela Neequaye, Shikha Saha, Martin Trick, Burkhard Steuernagel, Perla Troncoso-Rey, Frans van den Bosch, Pauline Stephenson, Maria H Traka, Lars Østergaard, Richard Mithen

## Abstract

**Background:** A diet rich in cruciferous vegetables is reported to have beneficial health effects, partially mediated by 4-methylsulfinylbutyl glucosinolate, or glucoraphanin, which is predominantly found within broccoli (*Brassica oleracea var italica*). We describe the downstream effects on transcription and metabolism in broccoli following the introgression of a genetic variant of MYB28 into broccoli from a wild *Brassica* relative which has previously been associated with enhancement of glucoraphanin.

**Results:** Whole genome sequencing, RNA expression and metabolite analyses were used to characterise the consequences of the introgression of either one or two copies of a genetic variant of the MYB28 transcription factor into a commercial broccoli genetic background. The introgression of the variant of MYB28 resulted in enhanced expression of genes involved in primary sulphate assimilation, sulphur metabolism and aliphatic glucosinolate biosynthesis, and enhanced accumulation of 4-methylsulphinyl butyl glucosinolate in florets. Other changes in transcription that may be related to non-targeted introgression events are reported. There were no consistent effects upon sulphur metabolites pools, apart from methionine-derived glucosinolates.

**Conclusion:** This study illustrates the downstream effects on transcription and metabolism of the introgression of a genetic variant of MYB28 from a wild species into a commercial broccoli genotype.

## Background

Glucosinolates are sulphur-containing glycosides produced by plants of the order Brassicales. These compounds comprise a common glucosinolate core moiety with a variable side chain derived from amino acids. In *Brassica*, the majority of glucosinolates are derived from methionine or tryptophan [1]. The evolutionary origins of these specialised plant metabolites are believed to be associated with their role in defence against pathogens and herbivores [2]. Following tissue disruption, myrosinase activity results in the degradation of glucosinolates to bioactive compounds that can deter herbivores and prevent pathogen colonisation. Within crop plants, certain of these isothiocyanates are important contributors to the flavour of brassicaceous salad crops such as *Eruca* and *Rorippa*, while these and other isothiocyanates have been associated with health-promoting activity. Foremost amongst these is the isothiocyanate sulforaphane derived from 4-methylsulphinylbutyl glucosinolate, or glucoraphanin. This non-volatile isothiocyanate has been shown in a variety of mammalian model systems to be a potent inducer of the *NRF2* transcription factor that regulates that expression of many phase 2 and anti-oxidation-associated genes [3]. These data are consistent with epidemiological data that has associated diets rich in Brassicaceae vegetables with reduced risk of cardiovascular disease and certain cancers. In order to facilitate research on the potential health promoting effects of sulforaphane, a series of broccoli genotypes that have enhanced levels glucoraphanin have been developed.

The characteristics of glucosinolates depends upon the precursor amino acid from which they are derived. Leucine, isoleucine, valine and methionine form the amino acid precursor of the most prominent and highly expressed group of glucosinolates, the aliphatic glucosinolates. In particular, glucosinolates derived from methionine are the most varied and highly represented, with methionine undergoing the widest variety of modifications [4]. The aliphatic glucosinolate 4-methylsulfinylbutyl glucosinolate (4-MSB), or glucoraphanin, is the most prevalent glucosinolate in broccoli - *Brassica oleracea var italica* [1].

In order to genetically modulate the glucosinolate content of broccoli, a wild Brassica known as *B. villosa* was crossed with a double haploid broccoli breeding line [5]. The F_1_ underwent a series of further backcrosses with a standard broccoli line and inbreeding was undertaken, combined with selection for high glucosinolate content. This produced a series of high-glucoraphanin F_1_ broccoli hybrids which are now commercially available as Beneforte™ broccoli. The Beneforte™ broccoli accumulates higher amounts of total sulphur compared to standard broccoli cultivars as well as increased methionine-derived glucosinolates, such as glucoraphanin, which is up to three times more abundant than in standard broccoli cultivars. A series of linkage maps were produced to assess the level of introgression of the *B. villosa* genome into these F1 broccoli hybrids. In all three of the high-glucoraphanin F_1_ hybrids studied, a region of *B. villosa* genome introgression was shared on C2. Within this shared region of introgression is an R2R3 MYB transcription factor known as *MYB28* or *HAG1 (High Aliphatic Glucosinolate 1)* [6]. The analyses below describe the genomic and transcriptomic composition of the previously uncharacterised, high glucoraphanin accumulating, ‘HG Inbred’ cultivar, alongside comparative metabolomics with a standard broccoli cultivar in addition to the Beneforte™ broccoli cultivar ‘1199’ described in [6].

It is known from studies in the model plant *Arabidopsis thaliana* that *MYB28* specifically regulates the production of methionine-derived glucosinolates such as glucoraphanin [7–9]. In all three independent studies, *myb28* mutants produced decreased levels of long-chain and short-chain aliphatic glucosinolates [7–9]. MYB28 mediates this effect through direct regulation of expression of glucosinolate biosynthesis genes involved in every step of glucosinolate production [7–9] in addition to regulating primary sulphur assimilation genes and partitioning of methionine; a vital precursor for aliphatic glucosinolates [9–11]. MYB28 has also been repeatedly and independently well characterised as a regulator of aliphatic glucosinolate biosynthesis across the *Brassica* genus, including homologues in *B. napus* [12], *B. juncea* [13] and *B. rapa* [14, 15] as well as *B. oleracea* varieties, kohlrabi [16] and Chinese kale [17]. However, the role of *MYB28* in broccoli (*Brassica oleracea var italica*) has yet to be fully characterised.

This study characterises the *MYB28* genotype, gene expression level and downstream phenotypic effect in a series of *MYB28* variant broccoli cultivars with either a single or double introgressed copy of the *MYB28^v^* allele from the wild Brassica, *Brassica villosa*. The following work characterises the expression of this transcription factor in regulating aliphatic glucosinolate production in field experiments of different broccoli cultivars and assesses the downstream effect of this on glucosinolate production, while addressing other major sulphur pools in this important crop plant.

## Results

### The high glucoraphanin broccoli genome contains introgressions from *Brassica villosa*

A high-glucoraphanin commercial broccoli, named Beneforté™ was previously generated from a cross between a standard commercial Broccoli and a high-glucoraphanin Inbred cultivar (HG Inbred) derived from the wild *B. villosa* (Mithen, 2003). To identify genomic regions associated with increased glucoraphanin levels in high-glucoraphanin broccoli, we compared the genome sequences of the HG Inbred with that of the wild *B. villosa* donor. The raw genomic sequence data returned between 122-131 million reads per sample (Table S1). This gave an average depth of coverage of 35.14X. The total length of assemblies varied between 396236782bp and 427100513bp, with number of contigs ranging from 108592 to 195078 (Table S2).

By using k-mer based comparison, general sequence content between the HG Inbred cultivar and the wild *B. villosa* donor, as well as the commercial broccoli background was determined. A map was generated by anchoring to chromosomes using gene models of the published and annotated TO1000 *B. oleracea* genome, a rapid cycling Chinese kale [18] (Fig. 1a). Analysis of total k-mer alignment found 9015484 k-mers shared only with the *B. villosa*, and 210931707 k-mers shared uniquely with the standard commercial broccoli, Ironman. Introgression of the *Brassica villosa* genome into the ‘HG Inbred’ background can be found throughout the genome, particularly on chromosomes 2, 3, 5 and 7.

**Figure 1.**
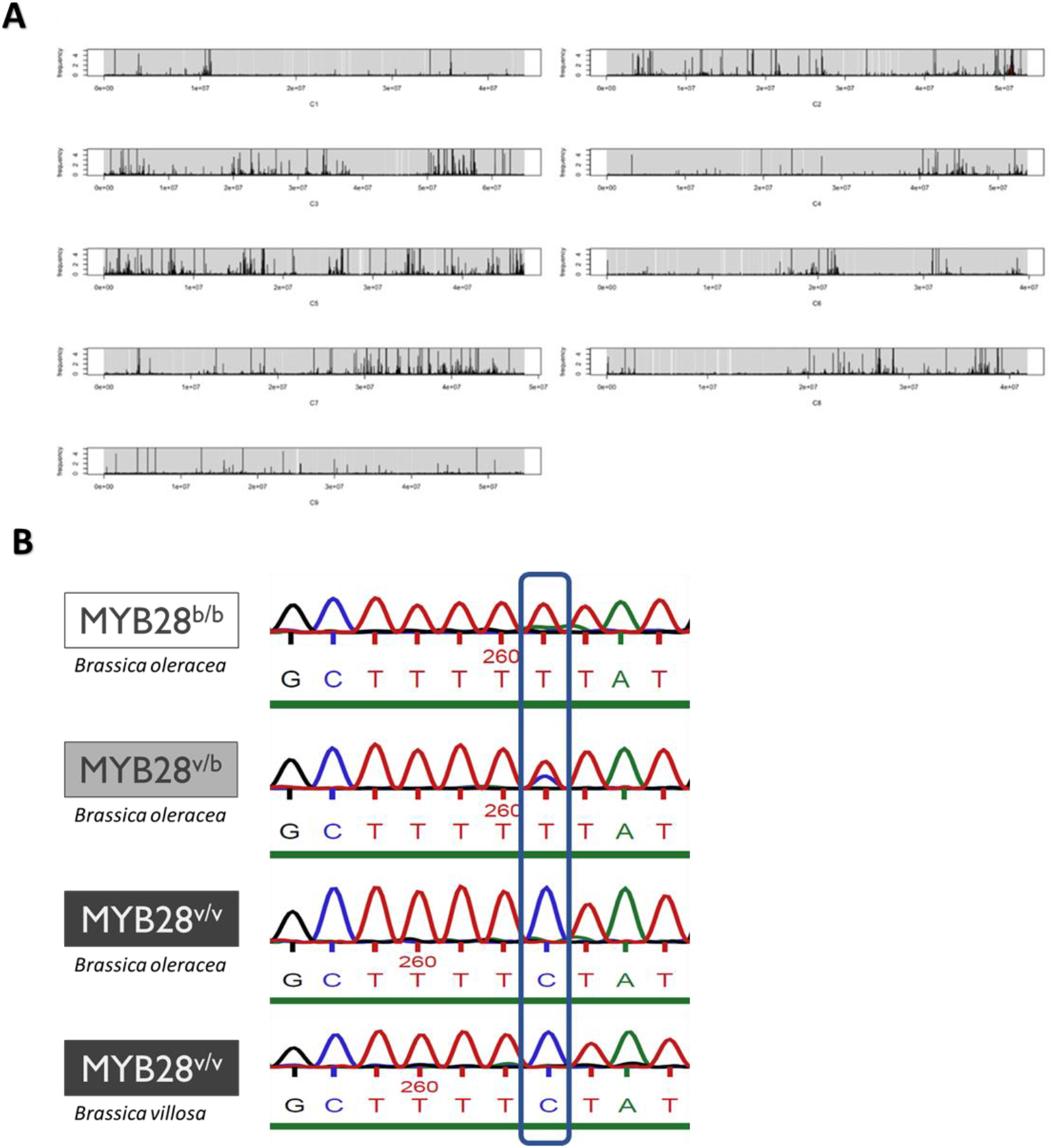
Genetic analysis of high glucoraphanin accumulating broccoli cultivars. **(a)** Introgression of the *B. villosa* genome (black) into the high-glucoraphanin broccoli cultivar ‘HG Inbred’ using k-mer analysis, visualised by chromosome through anchoring to the TO1000 reference sequence. The height of the black lines indicates the number of k-mers in that region which align to *B. villosa* divided by the number of k-mers which align to the commercial broccoli background in that same region. The map was generated by anchoring to chromosomes using gene models of the published and annotated TO1000 *B. oleracea* genome, a rapid cycling Chinese kale. **(b)** MYB28 genotypes of two broccoli cultivars, 1199 *MYB28^v/b^*, and HG Inbred MYB28^v/v^, derived from crossing of standard broccoli *MYB28^b/b^*(top) and *B. villosa MYB28^v/v^* (bottom). The diagnostic SNP visualised is found in the first intron.

Towards the bottom of chromosome C2, a high relative ‘frequency’ of k-mers indicates the introgression of *B. villosa* genomic material, which corresponds to a region containing a gene encoding the R2R3 MYB transcription factor, MYB28 (denoted by a red line). When sequencing specifically at this locus, the standard broccoli (Ironman) plants were confirmed to be homozygous for the standard broccoli allele (MYB28*^*b/b*^*) while the HG Inbred cultivar was found to be homozygous for the *B. villosa* allele (*MYB28^v/v^*) (Fig. 1b). The F1 hybrid ‘1199’ was heterozygous, carrying a single copy of the introgressed *B. villosa* allele (MYB28*^*v/b*^*). A total of 12 SNPs was found in the coding sequences of the HG Inbred cultivar as well as the *B. villosa* when compared to the reference TO1000 sequence (Figure S1). Of these 12 SNPs, 10 were shared between the HG Inbred and *B. villosa*, providing further confirmation that this cultivar contains an introgressed copy of this gene from the *B. villosa* genome. Ironman could not be used in this alignment as no contig was detected with a full-length *MYB28* copy on chromosome, C2 in the Ironman assembly. Instead, we took advantage of a recent genome sequence obtained for the *B. oleracea* species DH1012 [19, 20]. The region upstream of the start of the C2 copy of *MYB28* in *B.* oleracea DH1012 was aligned with corresponding sequence of the HG Inbred cultivar and *B. villosa*. This alignment shows that only the 1kb directly upstream of the start codon is conserved between the HG Inbred, *B. villosa* and the *B. oleracea* 1012 sequence, with 12 shared SNPs between the HG Inbred and the *B. villosa* when compared to the 1012 sequence, in addition to an A/T tract expansion upstream of the start codon. However, *B. villosa* has 7 SNPs exclusive to its promoter sequence (Figure S2). These results show various introgressions of the *B. villosa* genome into the, high-glucoraphanin, HG Inbred, including a region on chromosome C2 in which *MYB28* is located.

### Sulphur metabolism and glucosinolate biosynthesis genes are upregulated in High Glucoraphanin broccoli leaves

A transcriptomic analysis was carried out to study if changes in glucoraphanin levels between varieties is reflected in specific changes in the expression of certain classes of genes involved in glucosinolate biosynthesis. RNA was isolated from vegetative leaf tissue of field-grown HG Inbred and Ironman broccoli cultivars in addition to glasshouse-grown *B. villosa*, however this sample was not used in any statistical analyses. A Multidimensional Scaling (MDS) of FPKMs (Fragments Per Kilobase Million) of the two field-grown broccoli cultivars revealed that PC1 was found to explain 58.7% of variation and distinguishes the three replicates of the high glucoraphanin broccoli (*MYB28^v/v^*) and of the three replicates of the standard broccoli (*MYB28^b/b^*) (Fig. 2a, Table S4). PC2 was found to explain 19.2% of variation between samples but could not distinguish between the two genotypes due to variation in a high glucoraphanin broccoli (*MYB28^v/v^*) sample. It was found that 3286 genes (p threshold of <0.05) were significantly differentially expressed in the HG Inbred broccoli (*MYB28^v/v^*) in comparison to the standard broccoli (*MYB28^b/b^*), which was visualised in an unsupervised heatmap (Fig. 2b). Of the differentially expressed genes, 1668 were downregulated while 1618 were upregulated. In order to confirm the significance of these differentially expressed genes and resolve some of the variation seen within samples, RT-qPCR was used to confirm a selection of genes found to be significantly upregulated in the *MYB28^v/v^* broccoli cultivars (Figure S11).

**Figure 2.**
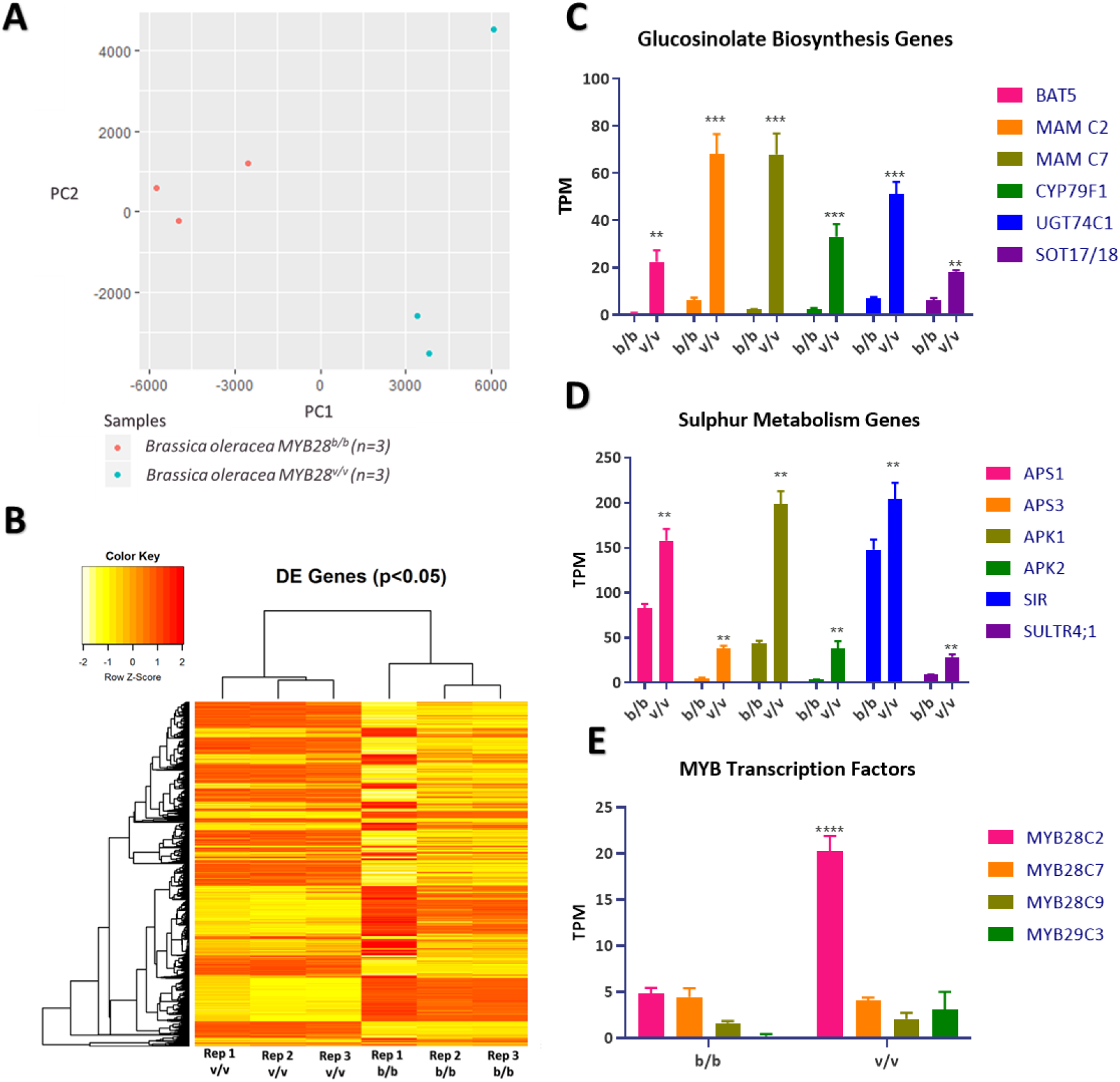
Transcriptome Analysis of MYB28-variant broccoli cultivars. **(a)** MDS of FPKMs of transcriptome profiles of *MYB28^v/v^* broccoli in comparison to the *MYB28^b/b^* broccoli. **(b)** Unsupervised heatmap of differentially expressed genes in *MYB28^v/v^* broccoli in comparison to the *MYB28^b/b^* broccoli visualised in individual replicates. **(c)** Significantly up regulated glucosinolate biosynthesis genes in *MYB28^v/v^* broccoli in comparison to the *MYB28^b/b^* broccoli. Expression levels expressed as TPM (Transcripts Per Kilobase Million). **(d)** Significantly up-regulated primary sulphur metabolism genes in *MYB28^v/v^* broccoli in comparison to the *MYB28^b/b^* broccoli. Expression levels expressed as TPM (Transcripts Per Kilobase Million). **(e)** Expression of *MYB28* and *MYB29* copies in *MYB28^v/v^* broccoli in comparison to the *MYB28^b/b^* broccoli. Values are mean of three independent samples, error bars represent ±SEM. Asterisks indicate significant difference as comparison with standard broccoli (*MYB28^b/b^*) (t-test: ** = p<0.01, *** = p<0.001).

Greater insight into the function of the differentially expressed genes is obtained through investigating which pathways are significantly enriched in the differential transcriptome analysis between *MYB28^v/v^* and *MYB28^b/b^* broccoli cultivars. A gene set enrichment analysis of the significantly differentially expressed genes (p threshold of <0.05 – Table S5) from this analysis was performed using TopGo [21]. This list included glucosinolate biosynthetic processes as well as the generation of aliphatic glucosinolate precursors such as methionine biosynthetic processes. Within the differentially expressed genes were several Gene Ontology (GO) terms relating to sulphur metabolism including regulation of sulphur metabolic process and sulphur assimilation (Table S6). This aligns with genes involved in numerous aspects of glucosinolate biosynthesis and sulphur metabolism.

In accordance with the GO analysis, many genes identified as being differentially expressed (p<0.05) between the *MYB28^v/v^* and *MYB28^b/b^* broccoli cultivars were identified as being directly involved in a range of sulphur metabolism processes, including glucosinolate biosynthesis, amino acid metabolism and sulphur assimilation. This includes well-characterised MYB28 targets involved in the aliphatic glucosinolate biosynthesis pathway (Fig. 2c), as well as genes involved in earlier stages in primary sulphur metabolism (Fig. 2d). Due to the varied positions of these differentially expressed genes across the genome, a subset was selected to be analysed at the single nucleotide level for introgression of the *B. villosa* genome. This included the C2 (Bo2g161100) and C7 (Bo7g098000) copies of *MAM3*, an important gene in methionine elongation, responsible for catalysing the first step in glucosinolate biosynthesis (Figure S6, Figure S7). In the transcriptomic analysis the C2 and C7 copies of *MAM3* were found to have 6-fold and 13-fold increased expression levels in the high-glucoraphanin broccoli (*MYB28^v/v^*) compared to standard broccoli (*MYB28^b/b^*), respectively. Genes involved in the second step of glucosinolate biosynthesis were also interrogated, including *CYP79F1* (Bo5g021810) which was found to have 5-fold increase in expression and *SOT18* (Bo4g120620) which had a 2-fold increase in expression (Figure S8, Figure S9). In addition to this, *APS3* (Bo1g057690), which plays a key role in primary sulphur metabolism and was found to be 4-fold up-regulated was also investigated (Figure S10). It is important to note that due to the historical whole-genome triplication of the *Brassica* genome since separating from *Arabidopsis thaliana* [22], some genes appear more than once as individual copies of important genes.

As described above, the C2 copy of *MYB28* was found to be introgressed in the high-glucoraphanin broccoli cultivars including the *MYB28^b/v^* heterozygous cultivar, 1199. In contrast, two other *MYB28* paralogous genes located on chromosomes C7 and C9 were not (Figure S3, Figure S4). Interestingly, only the C2 copy of *MYB28* is significantly upregulated in the HG Inbred. A low level of transcript expression of the additional paralogues of *MYB28* was detected, however a two-way ANOVA found that these genes as well as a close homologue, *MYB29* on C3 (Figure S5) were not significantly differentially expressed between the two broccoli cultivars nor were they found to contain introgressed segments from the *B. villosa* donor(Fig. 2e). Moreover, MYC transcription factors, also implicated in glucosinolate biosynthesis [23], were undetectable in either of the broccoli cultivars studied. This suggests that introgression of a single copy of *MYB28* was sufficient to cause the increased accumulation of glucoraphanin in these broccoli cultivars. In agreement with this, none of the glucosinolate biosynthesis genes as described above were found to be introgressed from the *B. villosa* genome. These results support the hypothesis that the sulphur metabolism phenotype of the high glucoraphanin broccoli is largely influenced by the introgression specifically of the C2 copy of *MYB28* from *B. villosa*.

In addition to glucosinolate biosynthesis and sulphur metabolism, several additional pathways were differentially regulated between these broccoli cultivars. These pathways included biotic and abiotic responses including ‘response to cytokinin’ and ‘response to bacterium’, which likely relate to the nature of the various interactions these plants had in the field. Additionally, GO terms relating to genetic regulation including ‘negative regulation of gene expression’, ‘RNA splicing’ and ‘chromatin organisation’ were identified (Table S6). The GO responses vary in level of detail, with the results with the lowest p values for this analysis including ‘nucleotide metabolic process’, ‘coenzyme metabolic process’ and ‘ribose phosphate metabolic process’. It is important to view these data alongside the differentially expressed genes to give a better overall perspective of changes between the cultivars. Amongst differentially expressed genes, those with the highest significance were genes such as a ‘HVA22-like protein’, in which *HVA22* is an abscisic acid induced gene, [24] and *PAP24 (PURPLE ACID PHOSPHATASE24)* a stress response gene related to phosphate starvation [25], which aligns with the enriched GO terms associated with stress response. Other genes with highly significant expression changes were *PEX10 (PEROXIN10)* which functions in peroxisomal protein translocation [26]*, OWL1 (ORIENTATION UNDER VERY LOW FLUENCES OF LIGHT 1)* which is involved in light response [27] and *SAR1 (SUPPRESSOR OF AUXIN RESISTANCE1)* which is involved in regulating auxin response [28]. Multiple loci in this HG inbred cultivar are introgressed from the *B. villosa* genome, in addition to *MYB28*. Molecular interactions among these introgressed loci have likely contributed to differential expression amongst numerous regulatory and metabolic pathways.

### Glucoraphanin content is consistently increased in high glucoraphanin broccoli

#### *MYB28* Expression

To determine the reproducibility of the increased expression of *MYB28* and effects on levels of glucosinolates, expression analysis by RT-qPCR was conducted on the vegetative leaf and floret tissue of field-grown broccoli cultivars in two subsequent years. In leaf tissue, the site of glucosinolate biosynthesis, both heterozygous (*MYB28^b/v^*) and homozygous (*MYB28^v/v^*) broccoli cultivars had significantly higher overall levels of *MYB28* expression relative to the standard broccoli cultivar Ironman. This was consistent over two years (Fig. 3a). *MYB28* expression was quantified in the floret tissues to assess conservation of this expression variation in a spatially distinct region of the edible crop parts (Fig. 3b). In year 1 only the *MYB28^v/v^* line was significantly upregulated in the floret, with the *MYB28^b/v^* line showing no significant deviation in expression when compared to the standard broccoli. In the second year, floret tissue showed significantly increased *MYB28* expression in both heterozygous and homozygous *MYB28^v^* broccoli cultivars when compared to standard broccoli.

**Figure 3.**
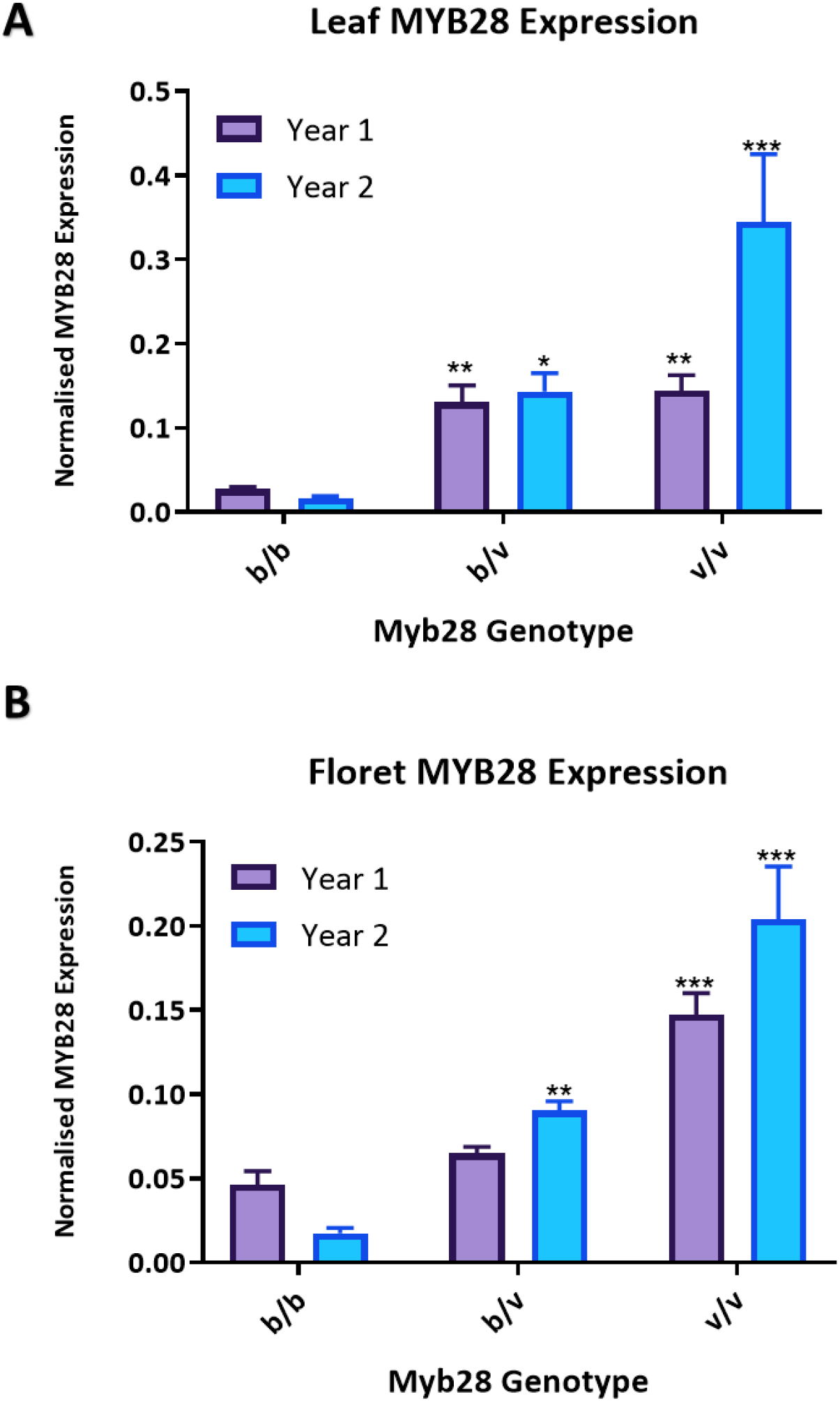
*MYB28* expression in vegetative leaves **(a)** and florets (**b)** of novel broccoli cultivars derived from crossing of standard broccoli (*MYB28^b/b^*) and *B. villosa* (*MYB28^v/v^*) over two trial years. Expression normalised against housekeeping gene Ubiquitin-C. Purple bars = data for Year 1, Blue bars = data for Year 2. Values are mean of four independent samples, error bars represent ±SEM. Asterisks indicate significant difference as comparison with standard broccoli (*MYB28^b/b^*) for the corresponding year, (t-test: * = p<0.05, ** = p<0.01, *** = p<0.001).

#### Glucosinolates

The leaves and florets used in the expression analysis were freeze-dried and profiled for different sulphur metabolites, to determine the effect of increased *MYB28* expression in these tissues. Glucoraphanin shows a significant stepwise increase in accumulation in leaves, which is more pronounced in year 1 when compared to year 2, where only the homozygous *MYB28^v/v^* broccoli cultivar shows a significant increase in glucoraphanin accumulation (Fig. 4a). Glucoraphanin in floret tissue follows the same trend as the leaves, with a significant stepwise increase in glucoraphanin. This trend is consistent over the two years (Fig. 4b). When performing a two-way ANOVA with Sidak's multiple comparisons test for levels of glucoraphanin in leaves compared to florets in the year 1 there is a significant increase in glucoraphanin in florets compared to leaves in all three genotypes studied (Table S8). In year 2 there is a significantly higher accumulation in glucoraphanin in florets when compared to leaves only in the *MYB28^b/v^* and *MYB28^v/v^* broccoli cultivars analysed and not in the commercial *MYB28^b/b^* broccoli (Table S9).

**Figure 4.**
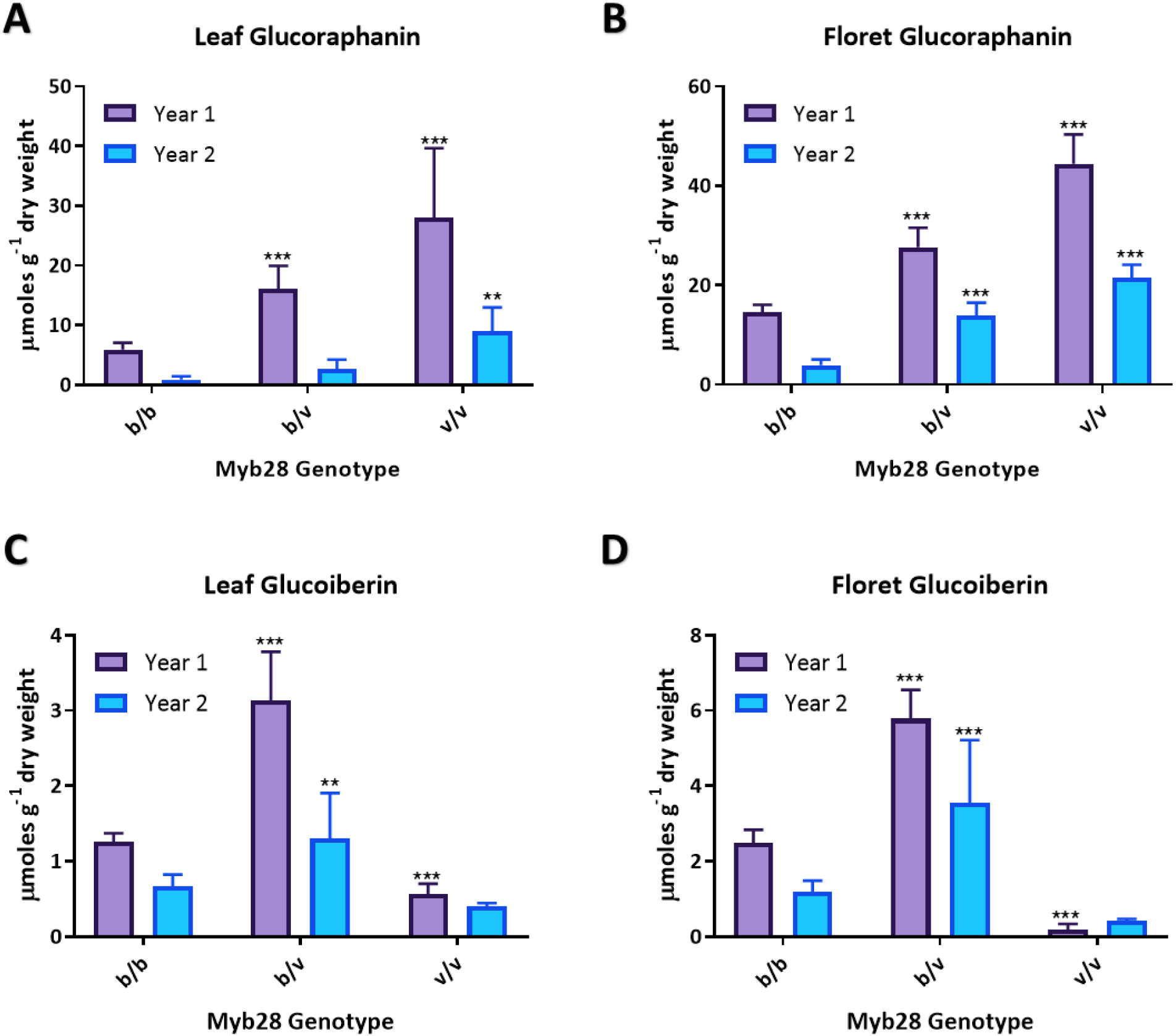
Methionine-derived glucosinolate content of novel broccoli cultivars derived from crossing of standard broccoli (*MYB28^b/b^*) and *B. villosa* (*MYB28^v/^*^v^) over two study years. **(a)** Glucoraphanin (4-MSB) content of vegetative leaves (top left). **(b)** Glucoraphanin (4-MSB) content of florets (top right). **(c)** Glucoiberin (3-MSP) content of vegetative leaves (bottom left). **(d)** Glucoiberin (3-MSP) content of florets (bottom right). Purple bars = data for Year 1, Blue bars = data for Year 2. Values are mean of ten independent samples, error bars represent ±SEM. Asterisks indicate significant difference as comparison with standard broccoli (*MYB28^b/b^*) for the corresponding year, (t-test: ** = p<0.01, *** = p<0.001).

Levels of the aliphatic glucosinolate, glucoiberin (3-MSP) are also significantly increased in the *MYB28^b/v^* broccoli cultivars when compared to the standard broccoli in leaf and floret tissues, consistently over the two trial years. However, the homozygous *MYB28^v/v^* broccoli cultivars show decreased levels of glucoiberin when compared to standard broccoli, in leaves and florets over the two trial years, with only the first year being significant (Fig. 4c and Fig. 4d).

For the parallel glucosinolate biosynthesis pathway, the tryptophan-derived glucosinolates (or indolic glucosinolates), results are more variable between tissues and years of the trial. In year 1 total indolic glucosinolates showed a stepwise decrease in accumulation in leaves, also observed in lower levels of indolyl methyl (I3M) (Fig. 5a) and 1-methoxyindolylmethyl (1MOI3M) (Fig. 5b) with no significant change found in the indolic glucosinolate, 4-methoxyindolylmethyl (4MOI3M) (Fig. 5c). Over the two trial years, levels of individual indolic glucosinolates as well as total glucosinolates were more consistent in floret material (Figs. 5d-f).

**Figure 5.**
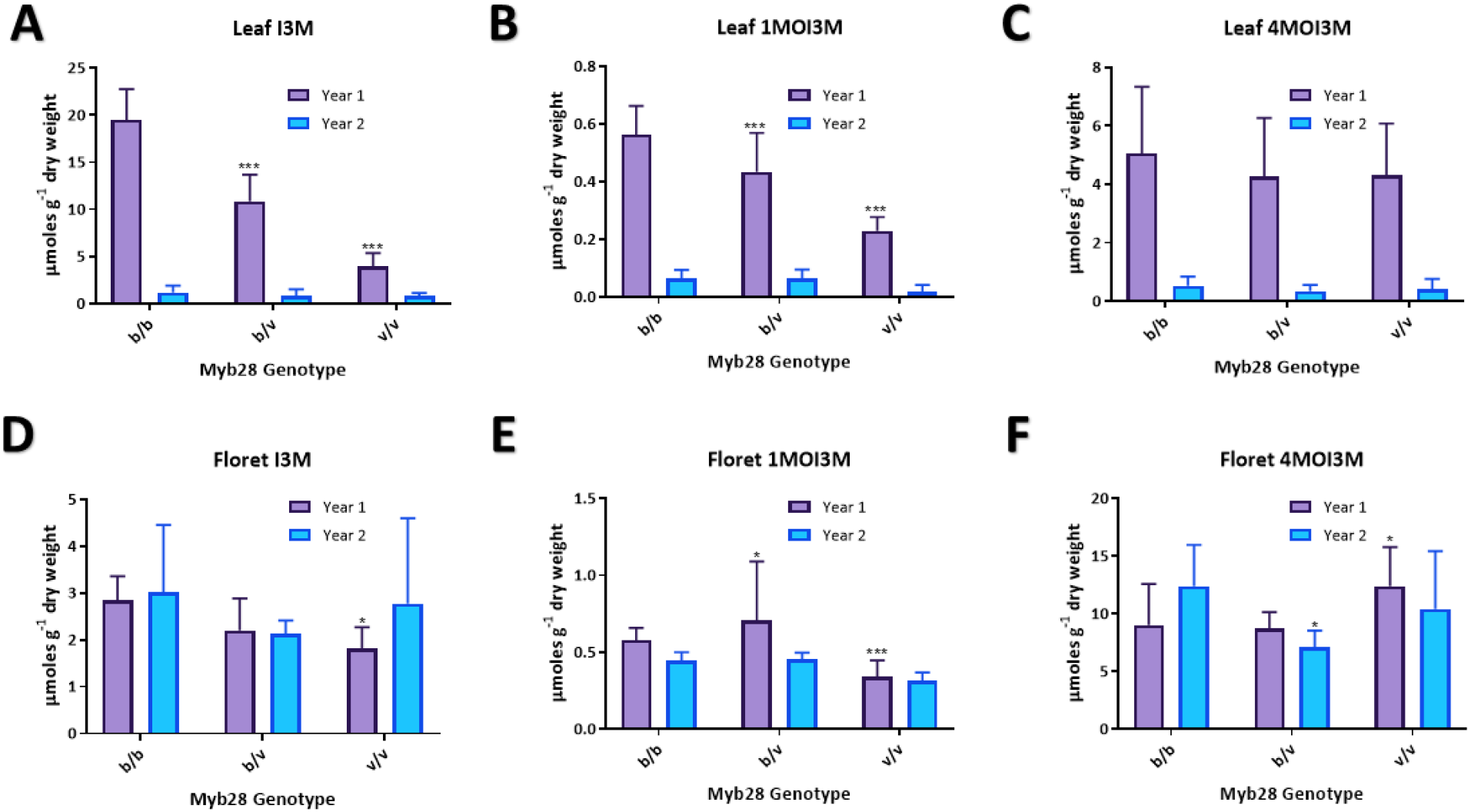
Tryptophan-derived glucosinolate content of novel broccoli cultivars derived from crossing of standard broccoli (*MYB28^b/b^*) and *B. villosa* (*MYB28^v/^*^v^) over two study years. **(a)** Indolyl methyl (I3M) content of vegetative leaves. **(b)**1-methoxyindolyl methyl (1MOI3M) content of vegetative leaves. **(c)**4-methoxyindolyl methyl (4 MOI3M) content of vegetative leaves. **(d)** Indolyl methyl (I3M) content of florets. **(e)**1-methoxyindolyl methyl (1 MOI3M) content of florets. **(f)**4-methoxyindolyl methyl (4 MOI3M) content of florets. Purple bars = data for Year 1, Blue bars = data for Year 2. Values are mean of ten independent samples, error bars represent ±SEM. Asterisks indicate significant difference as comparison with standard broccoli (*MYB28^b/b^*) for the corresponding year, (t-test: * = p<0.05, ** = p<0.01, *** = p<0.001).

#### Primary Sulphur Metabolites

To characterise the effect of the over-expressed *MYB28^v^* allele on the overall sulphur metabolome of the crop, major sulphur pools were quantified. Over both trial years, there was no notable change to sulphate levels in florets, the major sulphur source in broccoli (Fig. 6a). This was consistent in leaves in year 1, however in year 2 a stepwise decrease in sulphate accumulation was found in the heterozygous and homozygous *MYB28^v^* broccoli cultivars (Fig. 6b).

**Figure 6.**
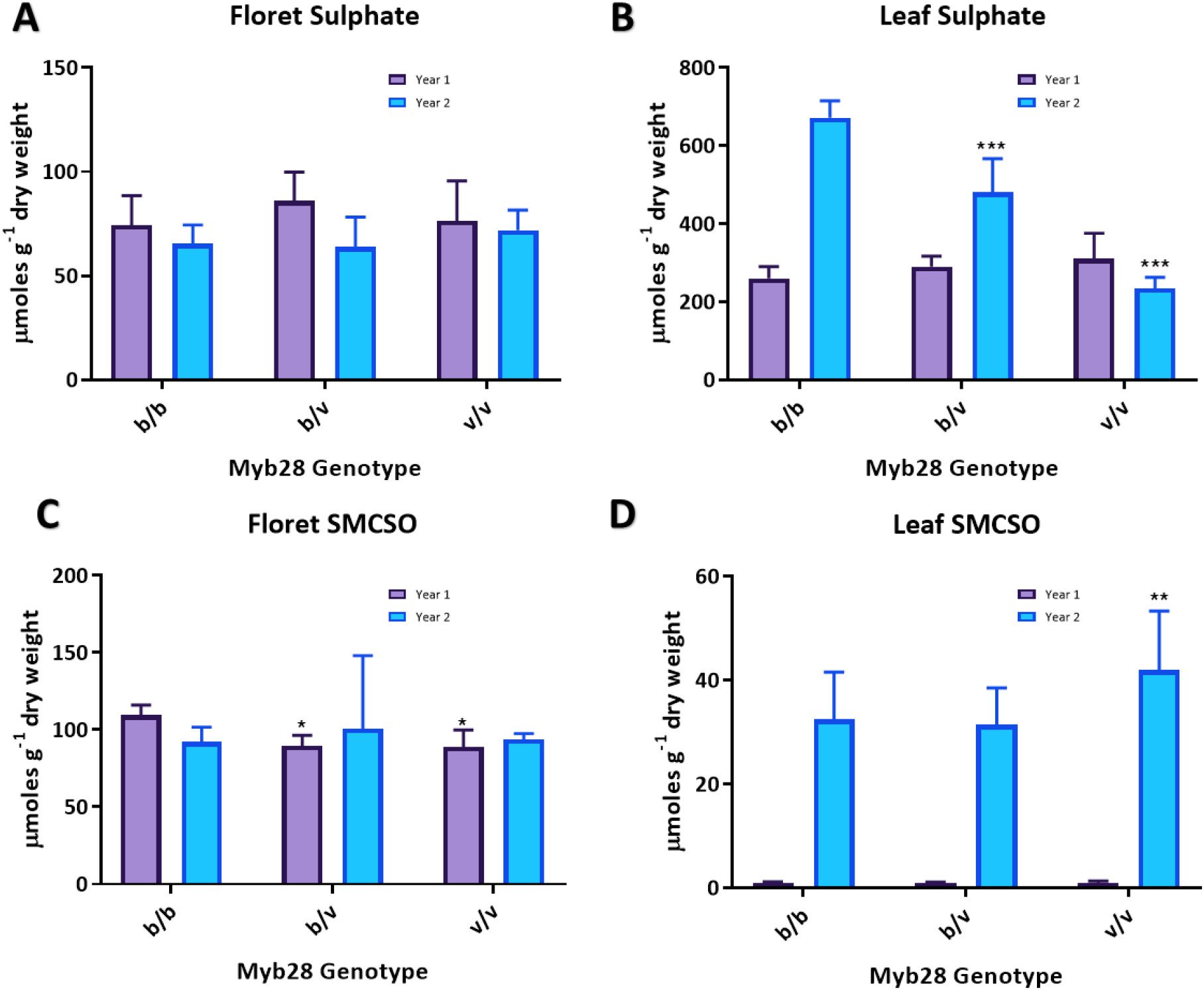
Primary sulphur metabolite content of novel broccoli cultivars derived from crossing of standard broccoli (*MYB28^b/b^*) and *B. villosa* (*MYB28^v/^*^v^) over two trial years. **(a)** Sulphate content of florets. **(b)** Sulphate content of vegetative leaves. **(c)** S-methyl cysteine sulphoxide (SMCSO) content of florets. **(d)** S-methyl cysteine sulphoxide (SMCSO) content of vegetative. Purple bars = data for Year 1, Blue bars = data for Year 2. Values are mean of ten independent samples, error bars represent ±SEM. Asterisks indicate significant difference as comparison with standard broccoli (*MYB28^b/b^*) for the corresponding year, (t-test * = p<0.05, ** = p<0.01, *** = p<0.001).

In year 1 only, the floret tissue showed significantly reduced levels of the sulphur compound s-methyl cysteine sulfoxide (SMCSO) in heterozygous and homozygous *MYB28^v^* broccoli cultivars in comparison to standard broccoli (Fig. 6c). This decrease in SMCSO was also recorded in the published Beneforté^®^ study, confirming previously reported data [6]. Levels of SMCSO were found to be almost 40-fold higher in leaves in year 2 when compared to year 1, with the *MYB28^v/v^* broccoli having higher levels than that of the standard broccoli. In year 1, there was no significant change in accumulation of SMCSO in leaf tissue across the different broccoli cultivars (Fig. 6d).

Total sulphur content of broccoli florets did not significantly differ between the three broccoli cultivars in year 1, however in year 2 the *MYB28^v/v^* broccoli showed a significant increase in total sulphur unlike the heterozygous *MYB28^b/v^* broccoli (Fig. 7a). Levels of the precursor amino acid for glucoraphanin, methionine, did not differ significantly in the florets between any of the genotypes over both trial years. This was similar for the other sulphur-containing amino acid, cysteine (Fig. 7b).

**Figure 7.**
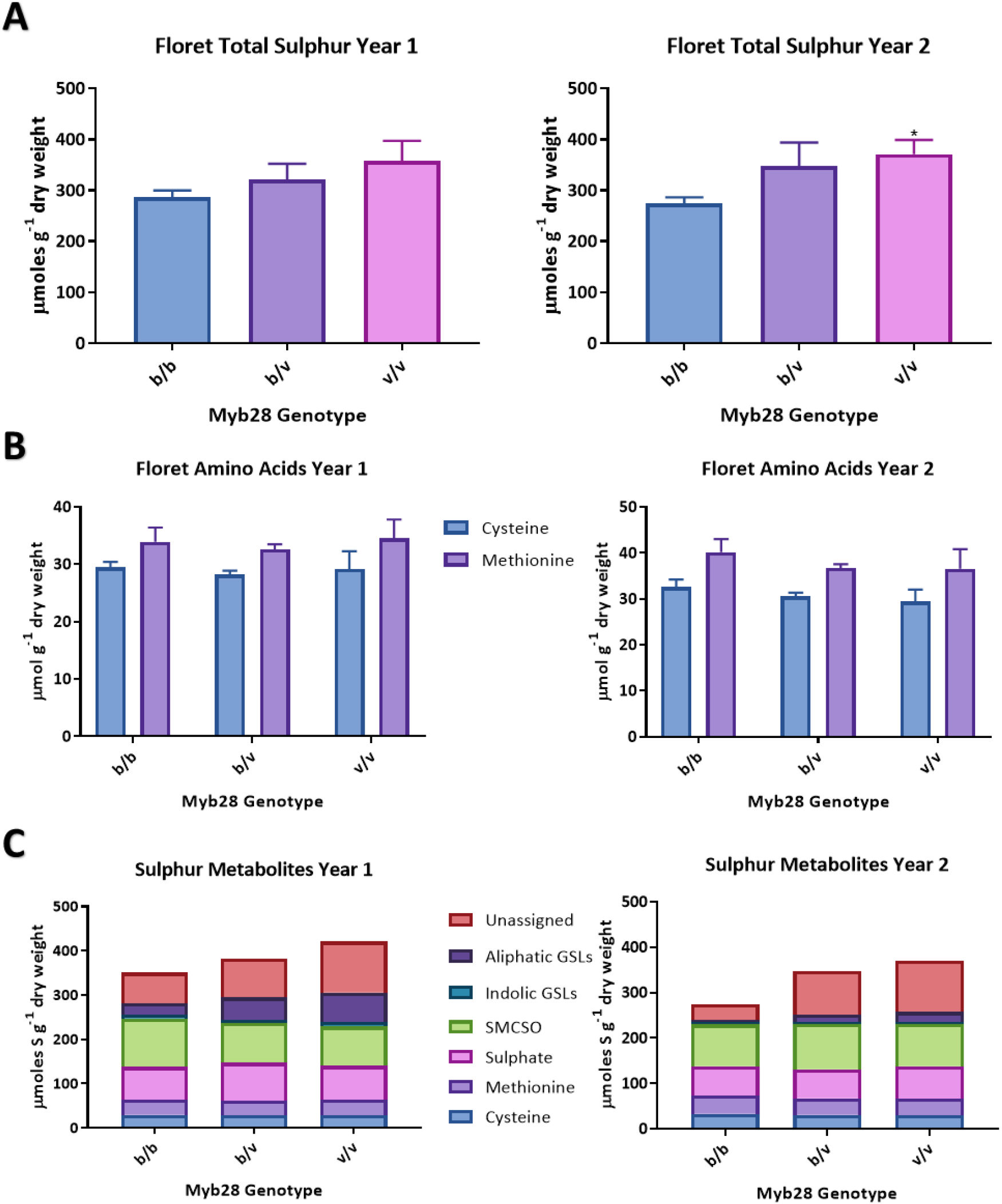
Sulphur metabolite content of florets of novel broccoli cultivars derived from crossing of standard broccoli (*MYB28^b/b^*) and *B. villosa* (*MYB28^v/^*^v^) over two study years. **(a)** Total sulphur content of florets. **(b)** Sulphur-containing amino acid content of florets. **(c)** All sulphur metabolites as a proportion of total sulphur in florets. Values are mean of ten independent samples, error bars represent ±SEM. Asterisks indicate significant difference as comparison with standard broccoli (*MYB28^b/b^*) for the corresponding year, (t-test * = p<0.05).

## Discussion

Whole genome sequencing analysis confirmed the introgression of the C2 copy of *MYB28* from the wild *Brassica*, *B. villosa*, into broccoli cultivar ‘HG Inbred’, without introgression of any of the other copies of *MYB28* or *MYB29*. RNA-seq analysis of the HG Inbred found several genes involved in aliphatic glucosinolate biosynthesis and primary sulphur metabolism to be upregulated in leaves, despite these genes having not been introgressed in the recurrent backcross breeding programme with *B. villosa*. Combining the genomic and transcriptomic analysis of these cultivars with data on the sulphur metabolomes of these crops confirmed that this introgression of the C2 copy of *MYB28* conferred increased accumulation of the aliphatic glucosinolate glucoraphanin, through up-regulation of genes involved in this biosynthesis pathway, without consistent disruption to primary sulphur metabolites, including those required as precursors or indolic glucosinolates.

This analysis provided a genomic perspective of the of introgression of the *B. villosa* genome into a high glucoraphanin broccoli cultivar, ‘HG Inbred’. A previous study also reported the introgression of *B. villosa* into the high glucosinolate F_1_ hybrid 1199 using KASpar markers [6]. This approach lacks the resolution afforded through genome sequencing as used in the current study. In this previous analysis the highest levels of shared *B. villosa* introgression between three hybrid cultivars were found on chromosomes 2, 3 and 8. Further analysis of the shared region of introgression corresponds with the location of our gene of interest, *MYB28* and is this same shared region that is found on chromosome 2 in the analysis described above. The variation observed in introgression levels throughout the genome have undoubtedly influenced the metabolic landscape of this crop, alongside that of *MYB28*. This variation is observed in the range of differentially expressed genes in the HG Inbred and subsequent variation in enriched pathways. The HG Inbred used in the analyses described here is a previously uncharacterised, alternative cultivar generated from the breeding process used to generate the 1199 cultivar from a previous study [6].

Many genes involved in sulphur metabolism were found to be upregulated in the HG Inbred, including genes involved in the three characteristic steps of glucosinolate biosynthesis, with the majority in amino acid side-chain elongation (*BCAT4, BAT5, BASS5, MAM3*) and core structure biosynthesis (*CYP79F1, GSTF11, SUR1, UGT74C1*) [4]. When related to the field experiment data, the increase in expression of these genes correlates with the significant increase in glucoraphanin accumulation in leaf tissue. In addition to assessing the extent of *B. villosa* introgression, the sequence analyses of the respective genomes enabled the interrogation of the origin of the glucosinolate biosynthesis genes, elucidating whether these alleles originate from *B. villosa*, or from the commercial broccoli parent. Genes which had been identified as being up regulated in HG Inbred compared to standard broccoli were interrogated, including those involved in primary sulphur metabolism (*APS3*), methionine elongation (*MAM3* – two copies), glucosinolate core structure biosynthesis (*CYP79F1, UGT74*) and secondary modifications (*SOT18*). These genes were all found to have the highest sequence homology with the commercial broccoli background as opposed to that of *B. villosa*. This suggests that the introgression and subsequent up-regulation of the R2R3 MYB transcription factor, *MYB28*, is able to confer the high accumulation of glucoraphanin found in these cultivars through up-regulation of downstream target genes in the aliphatic glucosinolate biosynthesis pathway, rather than introgression of these respective genes from the wild *Brassica*. This appears to be specific to the C2 copy of *MYB28*, with the C7 and C9 copies exhibiting wild-type sequence in addition to levels of expression.

Increased glucoraphanin is found in both the edible parts, the florets as well as the leaves. These results are consistent with studies in the leaves of *Arabidopsis thaliana* which find that *MYB28* over-expression lines confer increased levels of short-chain aliphatic glucosinolates, including 3-methylsulphinylpropyl (3-MSP) and 5-methylsulphinylpentyl glucosinolate (5-MSP), as well as levels of 4-MSB (4-methylsulphinylbutyl) [7–9]. These results are also supported at the transcript level, with *MYB28* overexpression coinciding with increased expression of genes throughout the aliphatic glucosinolate biosynthesis pathway. This is limited to aliphatic glucosinolate biosynthesis, with genes of the indolic glucosinolate biosynthesis pathway showing no significant change in *MYB28* overexpression and RNAi lines. [7, 9]. In addition, a study found that when *MYB28* was over-expressed in Chinese Kale (*B. oleracea var. alboglabra Bailey*), levels of aliphatic glucosinolates were increased in edible organs, with no change to indolic glucosinolates, which is consistent with the data presented here in *B. oleracea var italica* [17]. Moreover, over-expression lines of *MYB28* conferred increased expression of genes involved in aliphatic glucosinolate that are known MYB28 targets from analysis in *Arabidopsis thaliana*, including *MAM1, MAM3, CYP79F1, CYP79F2, CYP83A1, SOT17* and *SOT18*.

In 2014 the whole genome sequence of *B. oleracea* [18] was published with the use of the TO1000 rapid cycling kale as a reference genome. A subsequent sequencing collaboration project recently described the pan-genome of *B. oleracea* [29]. This study generated *de novo* assemblies for each of the domesticated varieties of *B. oleracea* and anchored them to the published reference TO1000 genome, as demonstrated in the analyses described above. However, their analysis found the pan-genome to have 61,379 gene models as opposed to the 59,225 found in the TO1000 reference sequence, in addition to some contigs generated in the analysis being unable to map to the TO1000 chromosomes. Whilst a perspective of the degree of genomic variation conferred through the breeding programme in the HG Inbred was able to be assessed in this study, it is possible that introgressions from the *B. villosa* were not characterised due to the inability to map to the nine TO1000 chromosomes, particularly as the mapping discussed here is anchored to gene models and will not characterise the non-coding variation found in gene-poor regions. The use of anchoring to the reference TO1000 lines in the generation of the genome introgression maps also introduced a bias in only presenting k-mers on contigs which align to published gene models in a rapid cycling Chinese kale. This analysis was required as the 30X coverage used here could not provide full coverage of the entire genomes of these cultivars in a resolution that could overcome the highly repetitive nature of the *B. oleracea* genome, which has not only a historical whole genome triplication event but a large proportion of transposable elements [30, 31]. Pan-genome analysis also found that genes involved in defence response, including those involved in glucosinolate biosynthesis, tended to have the most variation. This suggests that glucosinolate biosynthesis genes may be unique in sequence and copy number between the *B. villosa* and *Brassica oleracea* cultivars used in this study, and with that of the reference sequence. The possibility of losing genes in this pathway from analysis must be considered, as the final glucosinolate content varies both in volume and glucosinolate profile between these *B. oleracea* cultivars.

In contrast with the previous publication [6], total sulphur did not significantly increase in heterozygous and homozygous *MYB28^villosa^* broccoli cultivars. Other sulphur metabolites are not significantly different in cultivars, including the sulphur containing amino acids cysteine and methionine, methionine being the amino acid precursor for aliphatic glucosinolates such as glucoraphanin. There is a small but significant decrease in S-methyl cysteine sulfoxide (SMCSO) in florets from Year 1, however this was not consistent in the following year. This decrease in SMCSO was also observed in the heterozygous *MYB28^v^* cultivar in previous publication [6]. Amongst genes differentially expressed within leaves during year 1 were *HOMOCYSTEINE S-METHYLTRANSFERASE3* (*HMT3*) which plays an important role in the s-methylmethionine cycle [32] in addition to *METHIONINE SYNTHASE 2* (*MS2*) which catalyses the methylation of the thiol group from homocysteine to form methionine [33]. Taking this into account, alongside observed enrichment for cysteine metabolic processes, it is possible that in spite of unchanged absolute levels of sulphur containing amino acids involved in this process, reallocation between metabolites such as SMCSO and derived compounds such as glucoiberin may be taking place. Interestingly, the field data showed an unexpected reduction in the aliphatic glucosinolate 3-MSP or glucoiberin. In the transcriptome data there is a high fold change in expression level of two copies of ‘*MAM3’*, as well as other side chain elongation genes, which may provide some explanation to the lower levels of glucoiberin. This could be due to the high levels of *MAM3* expression and the central role of *MAM*s in elongation, as glucoiberin is a three-carbon glucosinolate whereas glucoraphanin is a four-carbon glucosinolate, requiring two elongation cycles. It has been suggested that *MAM* variation in the Brassicaceae may be responsible for aliphatic glucosinolate diversity [34]. Moreover, the variation in introgressions of the *B. villosa* genome suggest a wider influence of the breeding programme on the metabolic profiles of these crops, both direct and indirect changes though breeding may be mediated through interactions with regulators such as *MYB28*.

Field experiments were chosen for the analysis of these *B. oleracea* cultivars as this is the best representation of the natural and commercial environment of these crops. However, this type of analysis provides high levels of variation, with changes in conditions in the field from year to year likely having a significant effect on patterns of secondary metabolism, which require concerted efforts of signalling and genetic regulation. The variation observed in gene expression and metabolite levels between the two years are confirmed by two-way ANOVA, with year having a significant effect on metabolite and expression levels. However, in both years, the level of leaf glucoraphanin reflects that of the edible floral organ, with the relatively low leaf glucoraphanin levels of Year 2 reflected in corresponding low levels in the florets. Moreover, glucoraphanin levels follow a consistent pattern between the two genotypes, whilst patterns of other metabolites analysed appear to have a stronger environmental effect. The sulphur metabolism and glucosinolate profile of the crop is heavily influenced by genetic factors, but also in response to environmental cues. A previous study found sulphate and total glucosinolate content to be influenced by salinity and CO^2^ levels, with increased salinity and CO_2_ levels conferring increased aliphatic glucosinolate accumulation in leaves [35]. The variation seen in particular in levels of tryptophan-derived glucosinolates seen between study years is consistent with previous analysis in which aliphatic glucosinolate levels were less perturbed than indolic glucosinolate contents when changing growing conditions of broccoli, repeated over two years [36]. Nitrogen levels have also been known to affect indolic but not aliphatic glucosinolate levels in broccoli, with sulphur being the most significant indicator of aliphatic glucosinolate accumulation [37].

In Year 2 temperatures were higher than Year 1 in Norwich (www.metoffice.gov.uk). The lower levels of the aliphatic glucosinolates glucoraphanin and glucoiberin observed in this study are consistent with previous analysis which found accumulation of glucoraphanin to be reduced at higher temperatures (18°C) compared to lower temperatures (12°) during long day photoperiods, as would be reflected in the field conditions in which these broccoli plants were grown [38]. It is possible that this variation in abiotic and subsequent biotic interactions, such as emergence of insect pests, had an impact on glucosinolate production as well as the proportions of other sulphur metabolites, including SMCSO and sulphate. This highlights the importance of field interactions in studying metabolite profiles of crop plants, with further work potentially exploring the scoring of herbivory on these plants.

Further analysis would uncover the basis of the higher expression levels of the *MYB28^villosa^* allele. It is likely that motifs within the varied upstream regions of *MYB28* in these cultivars can confer the increased expression levels observed with the full extent of *MYB28* regulators and interactors having yet to be characterised. In addition to genetic interactions, *MYB28* has also been found to be induced by glucose signalling [7], which is released upon breakdown of glucoraphanin to sulforaphane [39]. There are likely many factors which directly and indirectly influence *MYB28* expression levels, which may have been disrupted by the sequence change of the promoter of the *MYB28* in *B. villosa* in contrast with the *MYB28* in commercial broccoli. Moreover, understanding the way in which these interactors, such as those relating to sulphur partitioning like *SLIM1* [40], may give some insight as to why some of the genes proposed to be *MYB28* targets from this research show varying susceptibility to *MYB28* expression changes as well as the variability in the field of primary sulphur metabolites such as SMCSO. The basis of this variation may also give some indication of the evolution of *MYB28* function and variation that has helped to sculpt the landscape of variation found in glucosinolates in plants of this group.

## Conclusions

In summary, the work presented here highlights the importance of fundamental analyses in the role and function of transcription factors regulating important biological functions including secondary metabolism and the accumulation of potentially beneficial compounds for human consumption. This work includes the use of modern technologies in whole genome and transcriptome profiling in the ongoing characterisation of the regulation of these processes in the field in often complex crop genomes.

## Methods

### Plant Material

The ‘wild-type’ broccoli cultivar used as a control throughout this study is the standard commercial broccoli cultivar, ‘Ironman’ which has a *MYB28^b/b^* genotype. The high glucoraphanin accumulating cultivars used in these analyses include ‘1199’ (commercially Beneforte™) which is heterozygous for the introgression of the *B. villosa MYB28^v^* allele (*MYB28^b/v^*) and a homozygous cultivar (*MYB28^v/v^*) referred to here as ‘HG Inbred’. If Ironman is A(b/b) x B(b/b) then 1199 is A(b/b) x B(v/v) and the HG inbred would then be C(v/v) at the MYB28 C2 locus. Details of the breeding programme can be found in [5]. The standard commercial broccoli cultivar Ironman (*MYB28^b/b^*), high glucoraphanin cultivars 1199 (*MYB28^b/v^*) and ‘HG Inbred’ (*MYB28^v/v^*) were grown under normal agronomic conditions in an experimental field plot in Norwich in 2016 and repeated in 2017 (Figure S12). Ten plants of each variety were grown in a randomized design. All broccoli heads and pooled vegetative leaf tissue were harvested from individual plants, at a stage equivalent to commercial maturity, freeze dried and ground to a fine powder for metabolite analysis, with some fresh tissue retained for genetic analysis.

### Whole Genome Sequencing Analysis

#### Sequencing

DNA was isolated from an individual leaves of a single individual of Ironman, HG Inbred and *B. villosa* using a phenol-chloroform extraction protocol [41]. Genomic DNA libraries were prepped and sequenced using Illumina HiSeq paired-end sequencing with 30X coverage by Novogene© (https://en.novogene.com) producing an average of 125 million total reads per sample with 35X average depth coverage (Table S1).

#### Assembly

Raw data from each line was trimmed using Trimmomatic version 0.33 [42] with parameters LEADING:20 TRAILING:20 SLIDINGWINDOW:10:20 MINLEN:50. Reads were assembled into contigs using clc assembly cell (https://www.qiagenbioinformatics.com/products/clc-assembly-cell) with default parameters. Quality of assemblies was checked using QUAST[43].

#### Introgression analysis

k-mers were counted in raw-data of *B. villosa* (donor) and Ironman (background) using jellyfish (http://www.cbcb.umd.edu/software/jellyfish) version 2.1.4. The parameter ‘-C’ was used for jellyfish to handle a k-mer along with its reverse complement as one item. Jellyfish is described in [44]. Sequence data from the draft assembly of the introgression line as well as k-mer counts on donor and background were used to determine the ratio of k-mers shared between donor and introgression line versus the ratio of k-mer shared between background and introgression line. A custom java program calculated the ratio of k-mers between the ‘reference’ high glucoraphanin assembly and the ‘wildtype’ (Ironman) and ‘donor’ (*B. villosa*) per contig. For each contig that was able to be anchored to the TO1000 assembly (PubMed ID 24916971), this ratio was visualised. The java program was written by Burkhard Steuernagel at the John Innes Centre.

For each contig of a CLC assembly, a set of k-mers (of length 31bp) was recorded. Then, in a second step, it was determined how many of those k-mers were present in the raw data of the ‘donor’ *B. villosa* and how many were present in the raw data of the commercial broccoli background, Ironman. The resulting plot displays the ratio of k-mer alignment between these lines in which shared genomic regions with the commercial broccoli parent are shown in grey while k-mer alignment with the *B. villosa* appear in black (Fig.1a).

#### Anchoring & Visualisation

Contigs from the CLC draft assemblies were anchored using gene models of the reference genome of *B. oleracea* TO1000 (Assembly BOL, INSDC Assembly GCA_000695525.1 version 97.1 [18]. Coding sequences (CDS) of each gene model were aligned to an assembly using NCBI blastn [45] with default parameters. For each gene model the contig with the highest alignment score was selected and the genomic position of the gene model was assigned to the contig. Plots were visualised and generated using RStudio v.1.0.143 [46].

#### MYB28 Genotyping

DNA from the extraction was used to amplify the fragment containing a region of the *MYB28* C2 locus, as published in [6]. PCR reactions were carried out on a GSTORM Thermocycler and prepared to a final volume of 20μl with 150ng of genomic DNA per reaction, using the New England Biolabs Taq Polymerase with Standard Taq Buffer. The PCR program is as follows; Heated Lid - 110°C, 95°C - 30 sec, Start Cycle - 35 times, 95°C - 20 sec, 58°C - 30 sec, 68°C - 1 min, End Cycle, 68°C - 5 min. Primers used were those published in [6]. PCR products were resolved using (1-2%) agarose gel electrophoresis. The subsequent PCR products were sent to Eurofins Genomics^®^ for sequencing using the Ready2Load sequencing service. A total of 17μl of PCR product (5ng/μl) was sent for sequencing, including 2μl of (10μM) PCR primer.

#### Gene Expression Analysis

Transcriptome analysis was undertaken on vegetative leaves from field-grown individual plants of *B. oleracea var italica* cultivars ‘Ironman’ and ‘HG Inbred’ from 2016 in addition to a glasshouse-grown *B. villosa* leaf of similar developmental age, as this is the site of glucosinolate biosynthesis. All leaf material was harvested while the plants were in a vegetative stage prior to emergence of inflorescences. RNA extraction was performed using E.Z.N.A.^®^ Plant RNA Kit provided by Omega Bio-tek Inc.

A total of 7 individual plant RNA samples (3 Ironman, 3 ‘HG Inbred’, and 1 B. villosa) were used to generate TruSeq non-directional RNA libraries at Earlham Institute (UK). Libraries were sequenced in one pool of 7 (7-plex) and run in on 2 lanes of the Illumina HiSeq2500 with a 125bp paired end read metric that generated an average of 160-170 million reads per sample. Sequencing was performed by the Earlham Institute. Individual plant sample RNA was used to generate 7 TruSeq non-directional RNA libraries. These libraries were sequenced through pooling in one pool of 7 (7-plex) and run in on 2 lanes of the Illumina HiSeq2500 with a 125bp paired end read metric with an average of 160-170 million reads per sample. This data is available on NCBI under Project ID PRJNA623495.

Data analysis of RNA-seq raw data was conducted following the protocol for the “new Tuxedo” suite for short reads with default parameters [47]. Differential expression analysis was performed using the Ballgown package in RStudio [46] [48]. A classical enrichment analysis was conducted on the list of differentially expressed (p<0.05) genes generated using the Ballgown software included in the latest version of the ‘Tuxedo Suite’ for RNA-seq data analysis (Pertea et al., 2016), which provided the significantly differentially expressed genes in the HG Inbred *MYB28^v/v^* broccoli compared to standard broccoli cultivar Ironman *MYB28^b/b^* (p<0.05). This data set was processed using TopGo analysis to gain a list of enriched gene ontologies in the dataset [21]. The gene set analysis statistically compared the representation of GO (Gene Ontology) terms,which were first assigned to the *B. oleracea* gene models using the *Arabidopsis thaliana* org.At.tair.db library (version 3.2.3) [49], in this gene set to that of the ‘expected’ value, to determine those to be considered ‘over-represented’ using the BP parameter with a node size of 10, then confirming the statistical significance of this using Fisher’s exact test. This was performed in RStudio v.1.0.143 [46]. A Multidimensional Scaling (MDS) of FPKMs (Fragments Per Kilobase Million) displaying Euclidean distances between these samples were analysed as a ‘principal component’ analysis to address the comparison of clustering between replicates of the different genotypes. This was performed using gene expression data, following removal of low abundance transcripts.

Gene expression analysis by RT-qPCR was performed using RNA extractions which generated cDNA using the REVERSE TRANSCRIPTASE M-MLV Kit Supplied by Life Technologies Ltd. Reactions included 5μL RNA, 1μl oligo dT, 1μL dNTPs and 5μL distilled H2O and were run at 65°C for 5 minutes. This was followed by addition of 4μl of 5X Buffer and 2μl 0.1M DTT (Dithiothreitol) before being kept at 37°C for 2 minutes. Finally, 1μ M-MLV reverse transcriptase was added to the mixture before being incubated at 37°C for 50 minutes followed by 70°C for 15 minutes. Concentrations of 150 ng/µl of cDNA were used for RT-qPCR reactions. Gene expression was quantified using the QuantiNova SYBR Green PCR Kit from Qiagen. PCRs were carried out in a Bio-Rad CFX96 machine (C1000 Touch). The PCR cycling conditions were 95 °C for 15 min, 40 cycles of 95 °C for 15 s, and 60 °C for 60 s. Primer sequences used can be found in Table S7.

#### Metabolite Analysis

Sulphate analysis was performed, modified from [50]. SMCSO analysis was performed, as modified from [51]. Glucosinolate analysis was performed according to [52]. Freeze-dried inflorescences resembling commercial edible broccoli floret samples were sent to the Eurofins Food and Feed Testing Laboratories for quantification of total sulphur, cysteine and methionine by HP-LC analysis.

#### Statistical Analyses

All statistical analyses were carried out on GraphPad Prism (version 8.2.0). Comparative analyses included t-tests of TPM in the RNA-seq analysis when comparing expression in the standard broccoli with the ‘HG Inbred’. Phenotypic analyses of metabolites included a one-way ANOVA comparing metabolite content of each of the broccoli cultivars for a single year, in addition to Tukey’s multiple comparison tests in which significance identifies a p value of less than 0.05 when comparing each cultivar to the standard commercial *MYB28^b/b^* broccoli cultivar for the corresponding year.

**Table 1.** Species and MYB28 genotype of Brassica line used in these analyses

| Line | Species | MYB28 Genotype |
| --- | --- | --- |
| Ironman | *B. oleracea* | *MYB28<sup>b/b</sup>* |
| 1199 | *B. oleracea* | *MYB28<sup>b/v</sup>* |
| HG Inbred | *B. oleracea* | *MYB28<sup>v/v</sup>* |
| *B. villosa* | *B. villosa* | *MYB28<sup>v/v</sup>* |

## Abbreviations

3-MSP: 3-methylsulfinylpentyl
4-MSB: 4-methylsulfinylbutyl
ANOVA: Analysis of Variance
GO: Gene Ontology
HAG1: High Aliphatic Glucosinolate 1
HP-LC: High Performance Liquid Chromatography
I3M: Indolyl methyl
1MOI3M: 1-methoxyindolyl methyl
4MOI3M: 4-methoxyindolyl methyl
MDS: Multidimensional Scaling
FPKMs: Fragments Per Kilobase Million
SMCSO: S-methyl cysteine sulfoxide
TPM: Transcripts Per Million

## Acknowledgements

We would like to thank Seminis Vegetable Seeds as the supplier of the 1199 and HG Inbred material. We would like to thank Graham Teakle (Warwick) for providing the *B. villosa* seeds. We would like to thank David Marc Jones for contributing the script for the Gene Ontology analysis.

## Authors’ Contributions

RM, LØ, MHT, and MN designed the experiments, analysed the data and wrote the paper. MN conducted the experiments. SS oversaw and supported in the metabolite analysis. MT and PTR provided scripts for RNA-seq analysis. BS provided the script for the whole genome sequencing assembly and performed the k-mer analysis. FB provided the seed material. PS provided supervisory and technical support.

## Availability of data and materials

Sequence data from RNA-seq described in this article has been released at NCBI (http://www.ncbi.nlm.nih.gov/bioproject/623495)

## Ethics approval and consent to participate

Not applicable

## Consent for publication

Not applicable.

