## Supplementary Material for "The Effect of Introgression of an R2R3 MYB Transcription Factor on Sulphur Metabolism in *Brassica oleracea*"

#### Whole Genome Sequencing

##### Supplementary Table 1

| Sample | Species | Genotype | Total reads | Average depth(X) |
|---|---|---|---|---|
| Ironman | *Brassica oleracea* | *MYB28$^{b/b}$* | 122,472,326 | 33.23 |
| HG Inbred | *Brassica oleracea* | *MYB28$^{v/v}$* | 130,769,980 | 36.14 |
| 1086 | *Brassica oleracea* | *MYB28$^{v/v}$* | 122,086,950 | 34.87 |
| Brassica villosa | *Brassica villosa* | *MYB28$^{v/v}$* | 127,819,586 | 36.31 |

*Table of raw sequencing data for each sample including, sample name, species MYB28 genotype, Total number of effective reads in clean data and the average depth of reads at each site, calculated by the total number of bases in the reads divided by size of the assembled genome*

#### Supplementary Table 2

| Name | Species | Genotype | Total Length (bp) | Number of Contigs | Largest Contig (bp) | N50 (bp) |
|---|---|---|---|---|---|---|
| Ironman | *Brassica oleracea* | *MYB28*$^{b/b}$ | 424718746 | 195078 | 83262 | 3840 |
| HG Inbred | *Brassica oleracea* | *MYB28*$^{v/v}$ | 427100513 | 166969 | 83262 | 5084 |
| 1086 | *Brassica oleracea* | *MYB28*$^{v/v}$ | 399827247 | 108592 | 86730 | 8041 |
| *Brassica villosa* | *Brassica villosa* | *MYB28*$^{v/v}$ | 396236782 | 136386 | 109455 | 6114 |

*Table of assembly data including sample name, species, MYB28 genotype, total assembly length (bp), number of contigs making up the assembly, the length of the longest contig (bp) and the N50 which is when half of the total genome is covered by contigs longer than or equal to N50 (bp).*

#### Supplementary Figure 1

Alignments of the MYB28 C2 Coding Sequence and Subsequent Protein Sequence of the high glucoraphanin broccoli lines HG Inbred and 1086, Brassica villosa and the reference Brassica oleracea TO1000 sequence taken from Ensembl Plants (Bolser, Staines et al. 2016) using the Clustal Omega Multiple Sequence Aligner (Sievers, Wilm et al. 2011).

```
CLUSTAL O(1.2.4) multiple sequence alignment

TO1000          ATGTCAAGAAAGCGATGTTGTGTCGGAGAAGGGCTGAAGAAAGGTGCATGGACCATTGAG   60
Bvillosa        ATGTCAAGAAAGCCATGTTGTGTCGGAGAAGGGCTGAAGAAAGGTGCATGGACCACCGAG   60
HGInbred        ATGTCAAGAAAGCCATGTTGTGTCGGAGAAGGGCTGAAGAAAGGTGCATGGACCACCGAG   60
                ************* *****************************************  ***

TO1000          GAAGACAAGAAGCTCATCTCTTATATTCACGAACATGGAGAAGGTGGCTGGCGCGACATT   120
Bvillosa        GAAGACAAGACGCTCATCTCTTACATTCACGAACATGGAGAAGGTGGCTGGCGCGACATT   120
HGInbred        GAAGACAAGACGCTCATCTCTTACATTCACGAACATGGAGAAGGTGGCTGGCGCGACATT   120
                ********** ************ ************************************

TO1000          CCCCAAAAAGCTGGGTTGAAAAGGTGTGGAAAGAGTTGTAGACTGCGATGGACTAACTAC   180
Bvillosa        CCCCAAAAAGCTGGGTTGAAAAGGTGTGGAAAGAGTTGCAGACTGCGATGGACTAACTAC   180
HGInbred        CCCCAAAAAGCTGGGTTGAAAAGGTGTGGAAAGAGTTGCAGACTGCGATGGACTAACTAC   180
                ************************************** *********************

TO1000          CTAAAACCTGAGATCAAAAGAGGCGAGTTTAGTTCAGAGGAGGAACAGATTATCATCATG   240
Bvillosa        CTAAAACCTGAGATCAAAAGAGGCGAGTTTAGTTCAGAGGAGGAACAGATTATCATCATG   240
HGInbred        CTAAAACCTGAGATCAAAAGAGGCGAGTTTAGTTCAGAGGAGGAACAGATTATCATCATG   240
                ************************************************************

TO1000          CTCCATGCTGCTCGTGGCAACAAGTGGTCAGTCATAGCTAGACATTTACCTAGAAGAACC   300
Bvillosa        CTCCATGCTGCTCGTGGCAACAAGTGGTCGGTCATAGCTAGACATTTACCTAGAAGAACC   300
HGInbred        CTCCATGCTGCTCGTGGCAACAAGTGGTCGGTCATAGCTAGACATTTACCTAGAAGAACC   300
                ***************************** ******************************

TO1000          GACAATGAGATCAAGAATTACTGGAACACGCATCTCAAAAAACGTTTGATCGAAGAGGGT   360
Bvillosa        GACAATGAGATCAAGAATTACTGGAACACGCATCTCAAAAAACGTTTGATCGAAGAGGGT   360
HGInbred        GACAATGAGATCAAGAATTACTGGAACACGCATCTCAAAAAACGTTTGATCGAAGAGGGT   360
                ************************************************************

TO1000          ATTGACCCCGTGACTCACAAGCCACTAGCTTCTAATACTAACCCTACTGTCACCACCACG   420
Bvillosa        ATTGACCCCGTGACTCACAAGCCACTAGCTTCTAATACTAACCCTACTGTCACCACCACG   420
HGInbred        ATTGACCCCGTGACTCACAAGCCACTAGCTTCTAATACTAACCCTACTGTCACCACCACG   420
                ************************************************************

TO1000          CAGCCAGAGAACTTGCATTCCCTAGATGCATCTAGTTCCGACAAGCGATACTCCCGGTCA   480
Bvillosa        CAGCCTGAGAACTTGCATTCCCTAGATGCATCTAGTTCCGACAAGCGATACTCCCGGTCA   480
HGInbred        CAGCCAGAGAACTTGCATTCCCTAGATGCATCTAGTTCCGACAAGCGATACTCCCGGTCA   480
                ***** ******************************************************

TO1000          AGCTCAATGCCTCCCATGTCTCTTCCTCTTTCCTCGGGTTTCAACCCGGTTTCCGAGATC   540
Bvillosa        AGCTCAATGCCTCCCATGTCTCTTCCTCTTTCCTCGGGTTTCAACCCGGTTTCCGAGATC   540
HGInbred        AGCTCAATGCCTCCCATGTCTCTTCCTCTTTCCTCGGGTTTCAACCCGGTTTCCGAGATC   540
                ************************************************************

TO1000          ATCACACCAGTTCAGAGAGGTTCCTTGAGTTGCAAGAAAAGTATTAAGAAATCGAACTCT   600
Bvillosa        ACCACACCAGTTCAGAGAGGTTCCTTGAGTTGCAAGAAAAGTATTAAGAAATCGAGCTCT   600
HGInbred        ACCACACCAGTTCAGAGAGGTTCCTTGAGTTGCAAGAAAAGTATTAAGAAATCGAGCTCT   600
                * ***************************************************** ****

TO1000          ACATCAAGGCTTTTAAACAAAGTTGCTGCTAAGGCCACTACCATCAAAGATATCTTGAGT   660
Bvillosa        ACATCAAGGCTTTTAAACAAAGTTGCTGCTAAGGCCACTACCATCAAAGATATCTTGAGT   660
HGInbred        ACATCAACGCTTTTAAACAAAGTTGCTGCTAAGGCCACTACCATCAAAGATATCTTGAGT   660
                ******* ****************************************************

TO1000          GCTACTACAATATCATATGAAAGCTTTTCCAATGGCTTCCCTGAGCAGACTCGCAATGAA   720
```

```
Bvillosa        GCTACTACAATATCATATGAAAGCTTTTCCAATGGCTTCCCTGAGCAGACGCGCAATGAA   720
HGInbred        GCTACTACAATATCATATGAAAGCTTTTCCAATGGCTTCCCTGAGCAGACGCGCAATGAA   720
                ************************************************** *********

TO1000          GAAGATAGTTCTAATGCTCTCGCTGACTTCGATCCATTTCCCCAATCATTGTTGTACACT   780
Bvillosa        GAAGATAGTTCTAATGCTCTCGCTGACTTCGATCCATTTCCCCAATCATTGTTGTACACT   780
HGInbred        GAAGATAGTTCTAATGCTCTCGCTGACTTCGATCCATTTCCCCAATCATTGTTGTACACT   780
                ************************************************************

TO1000          GAACATGAGGTCCATGCTACTTCTGATCTTGATATGGATCAGGGTTACGATTTCTCATAT   840
Bvillosa        GAACATGAGGTCCATGCTACTTCTGATCTTGATATGGATCAGGGTTACGATTTCTCATAT   840
HGInbred        GAACATGAGGTCCATGCTACTTCTGATCTTGATATGGATCAGGGTTACGATTTCTCATAT   840
                ************************************************************

TO1000          TTTCTCGAGACACTCGGGAGAGATGAGCACAATACGAATGTCGAGTATGGTCATGATATT   900
Bvillosa        TTTCTCGAGACACTCGGGAGAGATGAGCACAATACGAATGTCGAGTATGGTCATGATATT   900
HGInbred        TTTCTCGAGACACTCGGGAGAGATGAGCACAATACGAATGTCGAGTATGGTCATGATATT   900
                ************************************************************

TO1000          CTTCTTATGTCCGATGTGTCACAAGAAGTCTCATCAATTAGCGTTGATGATCAAGAAAAT   960
Bvillosa        CTTCTTATGTCCGATGTGTCACAAGAAGTCTCATCAATTAGCGTTGATGATCAAGACAAT   960
HGInbred        CTTCTTATGTCCGATGTGTCACAAGAAGTCTCATCAATTAGCGTTGATGATCAAGACAAT   960
                ******************************************************** ***

TO1000          ATGATAGGAAACTTCGAGGGCTGGTCAAATTATCTTCTTGACCATGATGGTTTGATAAAT   1020
Bvillosa        ATGATAGGAAACTTCGAGGGCTGGTCAAATTATCTTCTTGACCATGATGGTTTGATAAAT   1020
HGInbred        ATGATAGGAAACTTCGAGGGCTGGTCAAATTATCTTCTTGACCATGATGGTTTGATAAAT   1020
                ************************************************************

TO1000          GACACGGACTATGATTTCCTCGAGAAGCAAATCATATGA      1059
Bvillosa        GACACGGACTATGATTTCCTCGAGAAGCAAATCATATGA      1059
HGInbred        GACACGGACTATGATTTCCTCGAGAAGCAAATCATATGA      1059
                ***************************************
```

#### Translation

CLUSTAL O(1.2.4) multiple sequence alignment

R2 DNA Binding Domain
R3 DNA Binding Domain
Conserved residue

```
TO1000          MSRKRCCVGEGLKKGAWTIEEDKKLISYIHEHGEGGWRDIPQKAGLKRCGKSCRLRWTNY   60
Bvillosa        MSRKPCCVGEGLKKGAWTTEEDKTLISYIHEHGEGGWRDIPQKAGLKRCGKSCRLRWTNY   60
BenExtra        MSRKPCCVGEGLKKGAWTTEEDKTLISYIHEHGEGGWRDIPQKAGLKRCGKSCRLRWTNY   60
1086            MSRKPCCVGEGLKKGAWTTEEDKTLISYIHEHGEGGWRDIPQKAGLKRCGKSCRLRWTNY   60
                **** ************* ****.************************************

TO1000          LKPEIKRGEFSSEEEQIIIMLHAARGNKWSVIARHLPRRTDNEIKNYWNTHLKKRLIEEG   120
Bvillosa        LKPEIKRGEFSSEEEQIIIMLHAARGNKWSVIARHLPRRTDNEIKNYWNTHLKKRLIEEG   120
BenExtra        LKPEIKRGEFSSEEEQIIIMLHAARGNKWSVIARHLPRRTDNEIKNYWNTHLKKRLIEEG   120
1086            LKPEIKRGEFSSEEEQIIIMLHAARGNKWSVIARHLPRRTDNEIKNYWNTHLKKRLIEEG   120
                ************************************************************

TO1000          IDPVTHKPLASNTNPTVTTTQPENLHSLDASSSDKRYSRSSSMPPMSLPLSSGFNPVSEI   180
Bvillosa        IDPVTHKPLASNTNPTVTTTQPENLHSLDASSSDKRYSRSSSMPPMSLPLSSGFNPVSEI   180
BenExtra        IDPVTHKPLASNTNPTVTTTQPENLHSLDASSSDKRYSRSSSMPPMSLPLSSGFNPVSEI   180
1086            IDPVTHKPLASNTNPTVTTTQPENLHSLDASSSDKRYSRSSSMPPMSLPLSSGFNPVSEI   180
                ************************************************************

TO1000          ITPVQRGSLSCKKSIKKSNSTSRLLNKVAAKATTIKDILSATTISYESFSNGFPEQTRNE   240
Bvillosa        TTPVQRGSLSCKKSIKKSSSTSRLLNKVAAKATTIKDILSATTISYESFSNGFPEQTRNE   240
BenExtra        TTPVQRGSLSCKKSIKKSSSTSTLLNKVAAKATTIKDILSATTISYESFSNGFPEQTRNE   240
1086            TTPVQRGSLSCKKSIKKSSSTSTLLNKVAAKATTIKDILSATTISYESFSNGFPEQTRNE   240
                 *****************.*** *************************************

TO1000          EDSSNALADFDPFPQSLLYTEHEVHATSDLDMDQGYDFSYFLETLGRDEHNTNVEYGHDI   300
```

```
Bvillosa      EDSSNALADFDPFPQSLLYTEHEVHATSDLDMDQGYDFSYFLETLGRDEHNTNVEYGHDI  300
BenExtra      EDSSNALADFDPFPQSLLYTEHEVHATSDLDMDQGYDFSYFLETLGRDEHNTNVEYGHDI  300
1086          EDSSNALADFDPFPQSLLYTEHEVHATSDLDMDQGYDFSYFLETLGRDEHNTNVEYGHDI  300
              ************************************************************

TO1000        LLMSDVSQEVSSISVDDQENMIGNFEGWSNYLLDHDGLINDTDYDFLEKQII  352
Bvillosa      LLMSDVSQEVSSISVDDQDNMIGNFEGWSNYLLDHDGLINDTDYDFLEKQII  352
BenExtra      LLMSDVSQEVSSISVDDQDNMIGNFEGWSNYLLDHDGLINDTDYDFLEKQII  352
1086          LLMSDVSQEVSSISVDDQDNMIGNFEGWSNYLLDHDGLINDTDYDFLEKQII  352
              ******************:*********************************
```

Amino Acid Changes in Order

1. Arginine to Proline - Positively charged (basic amino acids; non-acidic amino acids); Polar; Hydrophilic; pK=12.5 → Non-polar; Hydrophobic
2. Isoleucine to Threonine - Non-polar; Hydrophobic → No charge (non-acidic amino acids); Polar; Hydrophilic
3. Lysine to Threonine - Positively charged (basic amino acids; non-acidic amino acids); Polar; Hydrophilic; pK=10.5 → No charge (non-acidic amino acids); Polar; Hydrophilic
4. Isoleucine to Threonine - Non-polar; Hydrophobic → No charge (non-acidic amino acids); Polar; Hydrophilic
5. Asparagine to Serine - No charge (non-acidic amino acids); Polar; Hydrophilic → No charge (non-acidic amino acids); Polar; Hydrophilic
6. Arginine to Threonine - Positively charged (basic amino acids; non-acidic amino acids); Polar; Hydrophilic; pK=12.5 → No charge (non-acidic amino acids); Polar; Hydrophilic
7. Glutamate to Aspartate - Negatively charged (acidic amino acids); Polar; Hydrophilic; pK=4.2 → Negatively charged (acidic amino acids); Polar; Hydrophilic; pK=3.9

#### Supplementary Figure 2

Alignment of MYB28 C2 Promoter of the high glucoraphanin broccoli lines HG Inbred and 1086, *Brassica villosa*, Ironman and the reference *Brassica oleracea* DH1012 sequence using the Clustal Omega Multiple Sequence Aligner (Sievers, Wilm et al. 2011).

```
Bvillosa      AATGGAAAACACGTTTGCACAAAAAAAAAAAACCTACCGACTATATCATTTTAATATAAA   1234
1012          TATCTTATTTTTTTCATGATATCATTAGAATAACTACCTACTATATCATTTTAATATAAA   1580
HGInbred      CATCTTATTTTTTTCATGATATCATAAGAATAACTACCTACTATATCATTTTAATATAAA   399
               **   *      *    * *  *    ** * ***** *********************

Bvillosa      CGATATATAGCATAGTACTTTTTATTATAGCATGTTTAAGTTAGTTATCTATTTATTAGT   1294
1012          CGATATATAGCATAGTACTTTTTATTATAGCATGTTTAAGTTAATTATCTATTTATTAGT   1640
HGInbred      CGATATATAGCATAGTACTTTTTATTATAGCATGTTTAAGTTAATTATCTATTTATTAGT   459
              ******************************************* ****************

Bvillosa      ACTGATGTACCATAAAATACCTGGTCGTTAATAGAATATATAACTATATTACTCATTTCT   1354
1012          ACTGATGTACCATAAAATACCTGGTAGTTAATTGAATATATAATTATATTACTCATTTCT   1700
HGInbred      ACTGATGTACCATAAAATACCTGGTAGTTAATTGAATATATAATTATATTACTCATTTCT   519
              ************************* ****** ********** ****************

Bvillosa      GGGTCACCATCAACTAGCAAGTAACACTATGAAATTTATTCTTGTTTGCATTTCTTTTTT   1414
1012          GGGTCACCATCAACTAGCATGTAACACTATGAAATTTATTCTTGTTTGCATTTCTTTTTT   1760
HGInbred      GGGTCACCATCAACTAGCAAGTAACACTATGAAATTTATTCTTGTTTGCATTTCTTTTTT   579
              ******************* ****************************************

Bvillosa      CTTTTTTCTTTTTAAATTAAAATCATTAAGCATTAACTTAACCTTAAAACATGATATTTT   1474
1012          CTTTTTCTTTTTTAAATTAAAATCATTAAGCATTAACTCAACCTTAAAACATGATATTTT   1820
HGInbred      CTTTTTCTTTTTTAAATTAAAATCATTAAGCATTAACTCAACCTTAAAACATGATATTTT   639
              ******  ****************************** *********************

Bvillosa      TCGTGATGTTGAGTTAGTAATTTTATAGAAGTAATAACTTAAAATTGCAATTCAATCATA   1534
1012          TCGTGATGTTGAGTTAGTAATTTTATAGAAGTAATAACTTAAAATTGCAATTCATTCATA   1880
HGInbred      TCGTGATGTTGAGTTAGTAATTTTATAGAAGTAATAACTTAAAATTGCAATTCAATCATA   699
              ****************************************************** *****

Bvillosa      ACACATAATAACTCAAATTTAAAATCAAAAATTTAAATTAATACATGAGTGGTCATTTT   1594
1012          ACACATAATAACTCAAATTTAAAATCAAAAATTTAAATTAATACATGAGTGGTCATTTT   1940
HGInbred      ACACATAATAACTCAAATTTAAAATCAAAAATTTAAATTAATACATGAGTGGTCATTTT   759
              ************************************************************

Bvillosa      TGAGTTACAGCCATAAATCAAACTAAAAAACTCTAACTCAGGGCACCAATCAA-GCTTTA   1653
1012          TGAGTTACAGCCATAAATCAAACTAAAAAACTCTAACTCAGGTCACCAATCAAAGCTTTA   2000
HGInbred      TGAGTTACAGCCATAAATCAAACTAAAAAACTCTAACTCAGGGCACCAATCAA-GCTTTA   818
              ****************************************** ********** ******

Bvillosa      AACCGAAATTTAAACATACATTAATACTGCACGTCATTAGTGCTAATCCTTCTTTTTGT   1713
1012          AACCGAAATTTAAACATACATTAATACTGCACGTCATTAGTGCTAATCCTTCTTTTTGT   2060
HGInbred      AACCGAAATTTAAACATACATTAATACTGCACGTCATTAGTGCTAATCCTTCTTTTTGT   878
              ************************************************************

Bvillosa      GCTAGAAATGCTTCAAAATAAATATGTATAGCAAAATGTAGTAATATATACAGCATCAAA   1773
1012          GCTAGAAATGCTTCAAAATAAATATGTATAGCAAAATGTAGTAATATATACAGCATCAAA   2120
HGInbred      GCTAGAAATGCTTCAAAATAAATATGTATAGCAAAATGTAGTAATATATACAGCATCAAA   938
              ************************************************************

Bvillosa      TTGCAAATATTTTTCATGCTAGAGAATAATTGCGAGCTAGTTTAAACAAATTGATCTATT   1833
1012          TTGCAAATATTTTTCATGTTAGAGAATAATTGCGAGCTAGTTTAAACAAATTGATCTATT   2180
HGInbred      TTGCAAATATTTTTCATGTTAGAGAATAATTGCGAGCTAGTTTAAACAAATTGATCTATT   998
              ****************** *****************************************

Bvillosa      TAAATAATTAATAATTCGCGTACCAAAATATCTATCATAATCAACATTGTGAGCATGTAC   1893
1012          TAAATAATTAATAATTCGCGTACCAAAATATCTATCATAATCAACATTGTGAGCATGTAC   2240
HGInbred      TAAATAATTAATAATTCGCGTACCAAAATATCTATCATAATCAACATTGTGAGCATGTAC   1058
              ************************************************************

Bvillosa      GTGTGTATATATAGAAGAGAATAGCTCGTAAGCTCACCACCATCACACAATTCATTGCTC   1953
```

```
1012          GTGTGTATATATAGAAGAGAATAGCTCGTAAGCTCACCACCATCACACAATTCATTGCTC  2300
HGInbred      GTGTGTATATATAGAAGAGAATAGCTCGTAAGCTCACCACCATCACACAATTCATTGCTC  1118
              ************************************************************

Bvillosa      TTCTTCACAAGTTTCTTTAGTCTGATCTTATAGTGTTATAAAAATACATATAATATTTCA  2013
1012          TTCTTCACAAGTTTCTGTAGTCTGATCTTATAGTGTTATAAAAATACATATAATATTTCA  2360
HGInbred      TTCTTCACAAGTTTCTTTAGTCTGATCTTATAGTGTTATAAAAATACATATAATATTTCA  1178
              **************** *******************************************

Bvillosa      AGGATATATAGAAAGCAAAAACTCAAAGTCTACTTGAAAAACATGAAAACACCTTGCAGC  2073
1012          AGGATATATAGAAGGCAAAAACTCAAAGTCTACTTGAAAAACATGAAAACACCTAGCAGC  2420
HGInbred      AGGATATATAGAAAGCAAAAACTCAAAGTCTACTTGAAAAACATGAAAACACCTTGCAGC  1238
              ************* **************************************** *****

Bvillosa      TCTGTGGGTAAGACCCAAGAGCGCTTCTCGATTAGTCTCATATTCAGATGTATCAGAAAT  2133
1012          TCTGTGGGTAAGACCCAAGAGCGCTTCTCGATTAGTCTCATATTCAGATGTATCAGAGTT  2480
HGInbred      TCTGTGGGTAAGACCCAAGAGCGCTTCTCGATTAGTCTCATATTCAGATGTATCAGAAAT  1298
              *********************************************************  *

Bvillosa      CTCATTAACAGATCTACTTCTTTCTTACCTTTTTAGAAAATTTCCTTTCTGATTTTAGTT  2193
1012          CTCATTAACAGATCTATTTCTTTCTTACCTTTTTAGAAAATTTCCTTTCTGATTTTAGTT  2540
HGInbred      CTCATTAACAGATCTACTTCTTTCTTACCTTTTTAGAAAATTTCCTTTCTGATTTTAGTT  1358
              **************** *******************************************

Bvillosa      TCCTCAAGTATATTTTTCTCAATATAGTATTTCCTTTGGTATTGTTTGTACCTTTTACAC  2253
1012          TCCTCAAGTATATTTTTCTCAATATAGTATTTCCTTTGGTATTGTTTGAACCTTTTACAC  2600
HGInbred      TCCTCAAGTATATTTTTCTCAATATAGTATTTCCTTTGGTATTGTTTGTACCTTTTACAC  1418
              ************************************************ ***********

Bvillosa      ATTAGTGTTCATATATATATATATATATATATATATATATATCGGAGAAA----------  2303
1012          ATTAGTGTTCATCTATATATATCGGAGAAA------------------------------  2630
HGInbred      ATTAGTGTTCATCTATATATATATATATATATATATATATATATATATATATCGGAGAAA  1478
              ************ *********      *
```

### Gene Expression Analysis

#### Supplementary Table 3

| Sample Description | Uniquely Aligned Reads (%) | Multi-mapped Reads (%) | Overall Alignment Rate (%) |
|---|---|---|---|
| Standard Broccoli Replicate 1 | 81.46 | 8.26 | 89.72 |
| Standard Broccoli Replicate 2 | 82.00 | 7.94 | 89.94 |
| Standard Broccoli Replicate 3 | 82.76 | 7.58 | 90.34 |
| High Glucoraphanin Broccoli Replicate 1 | 84.60 | 6.56 | 91.16 |
| High Glucoraphanin Broccoli Replicate 2 | 83.92 | 6.84 | 90.76 |
| High Glucoraphanin Broccoli Replicate 3 | 82.61 | 7.59 | 90.20 |
| Brassica villosa | 73.02 | 9.31 | 82.33 |

*Read mapping statistics for each sample*

#### Supplementary Table 4

Principal component analysis results of PCA using scaling factor F.

| | PC1 | PC2 | PC3 | PC4 | PC5 | PC6 |
|---|---|---|---|---|---|---|
| Standard deviation | 5032.125 | 2879.3326 | 2238.4195 | 1.852e+03 | 1.040e+03 | 1.137e-11 |
| Proportion of Variance | 0.587 | 0.1922 | 0.1162 | 7.955e-02 | 2.508e-02 | 0.000e+00 |
| Cumulative Proportion | 0.587 | 0.7792 | 0.8954 | 9.749e-01 | 1.000e+00 | 1.000e+00 |

#### Supplementary Table 5

| | geneNames | feature | id | fc | pval | qval | Gene ID |
|---|---|---|---|---|---|---|---|
| 4 | Putative endonuclease or glycosyl hydrolase | gene | MSTRG.16209 | 12.69441 | 9.11E-06 | 0.043188 | Bo5g004880.1 |
| 3 | Polynucleotide adenylyltransferase family protein | gene | MSTRG.18490 | 20.24055 | 8.65E-06 | 0.043188 | Bo5g108500.1 |
| 1 | . | gene | MSTRG.25360 | 18.1049 | 3.32E-06 | 0.043188 | Bo7g104030.1 |
| 5 | . | gene | MSTRG.27713 | 5.255825 | 1.01E-05 | 0.043188 | Bo8g070300.1 |
| 2 | . | gene | MSTRG.24122 | 10.12422 | 8.47E-06 | 0.043188 | |
| 6 | . | gene | MSTRG.11629 | 0.056974 | 1.48E-05 | 0.046317 | Bo3g164100.1 |
| 8 | . | gene | MSTRG.16238 | 1.137413 | 1.77E-05 | 0.046317 | Bo5g005250.1 |
| 9 | . | gene | MSTRG.32725 | 0.592952 | 1.94E-05 | 0.046317 | Bo9g152110.1 |
| 7 | . | gene | MSTRG.21015 | 3.906741 | 1.71E-05 | 0.046317 | |
| 16 | . | gene | MSTRG.122 | 0.39172 | 4.52E-05 | 0.047313 | Bo1g004770.1 |
| 26 | ATPAP24 | gene | MSTRG.1279 | 1.589799 | 6.27E-05 | 0.047313 | Bo1g039400.1 |
| 19 | GTP-binding nuclear protein | gene | MSTRG.4306 | 1.898386 | 5.25E-05 | 0.047313 | Bo2g041150.1 |
| 10 | . | gene | MSTRG.6499 | 0.509372 | 2.36E-05 | 0.047313 | Bo2g159490.1 |
| 14 | Copine (Calcium-dependent phospholipid-binding protein) family | gene | MSTRG.6662 | 14.26628 | 4.07E-05 | 0.047313 | Bo2g165960.1 |
| 28 | . | gene | MSTRG.13378 | 0.375067 | 6.39E-05 | 0.047313 | Bo4g061570.1 |
| 11 | . | gene | MSTRG.13757 | 0.590079 | 2.75E-05 | 0.047313 | Bo4g097040.1 |
| 22 | Basic-leucine zipper (BZIP) transcription factor family protein | gene | MSTRG.14787 | 23.31591 | 5.94E-05 | 0.047313 | Bo4g145800.1 |
| 24 | . | gene | MSTRG.16406 | 0.822018 | 6.19E-05 | 0.047313 | Bo5g008660.1 |
| 12 | HVA22-like protein | gene | MSTRG.17168 | 0.446671 | 3.99E-05 | 0.047313 | Bo5g029080.1 |
| 30 | . | gene | MSTRG.20910 | 5.196881 | 6.76E-05 | 0.047313 | Bo6g068460.1 |
| 27 | . | gene | MSTRG.25361 | 8.013577 | 6.29E-05 | 0.047313 | Bo7g104040.1 |
| 25 | . | gene | MSTRG.26462 | 0.542941 | 6.21E-05 | 0.047313 | Bo8g009520.1 |
| 13 | . | gene | MSTRG.27251 | 0.658006 | 4.05E-05 | 0.047313 | Bo8g057950.1 |
| 21 | PEX10 | gene | MSTRG.28762 | 1.701808 | 5.74E-05 | 0.047313 | Bo8g098750.1 |
| 17 | . | gene | MSTRG.30093 | 0.113528 | 4.74E-05 | 0.047313 | Bo9g010300.1 |

| | | | | | | | |
|---|---|---|---|---|---|---|---|
| 15 | . | gene | MSTRG.30826 | 0.716223 | 4.51E-05 | 0.047313 | Bo9g033940.1 |
| 20 | Glutamyl-tRNA reductase | gene | MSTRG.31041 | 0.127258 | 5.32E-05 | 0.047313 | Bo9g050950.1 |
| 18 | . | gene | MSTRG.34623 | 1.492929 | 4.93E-05 | 0.047313 | |
| 23 | . | gene | MSTRG.23803 | 0.171393 | 6.15E-05 | 0.047313 | |
| 29 | . | gene | MSTRG.34721 | 2.85156 | 6.51E-05 | 0.047313 | |
| 31 | . | gene | MSTRG.34442 | 0.666704 | 6.84E-05 | 0.047313 | |
| 32 | . | gene | MSTRG.12880 | 1.013395 | 7.37E-05 | 0.049426 | |
| 33 | . | gene | MSTRG.17689 | 0.233559 | 7.63E-05 | 0.049576 | Bo5g051930.1 |
| 34 | . | gene | MSTRG.1535 | 0.552402 | 8.31E-05 | 0.052345 | Bo1g051270.1 |
| 35 | . | gene | MSTRG.27771 | 3.535931 | 8.92E-05 | 0.052345 | Bo8g071060.1 |
| 36 | . | gene | MSTRG.11058 | 4.969509 | 9.15E-05 | 0.052345 | |
| 37 | . | gene | MSTRG.2356 | 9.81743 | 9.19E-05 | 0.052345 | |
| 38 | . | gene | MSTRG.35558 | 3.181981 | 9.27E-05 | 0.052345 | |
| 39 | . | gene | MSTRG.12842 | 0.560824 | 9.53E-05 | 0.052387 | |
| 40 | . | gene | MSTRG.2424 | 12.70685 | 0.000105 | 0.056139 | Bo1g108300.1 |
| 41 | . | gene | MSTRG.6965 | 0.391848 | 0.000114 | 0.059466 | Bo3g004520.1 |
| 42 | . | gene | MSTRG.1756 | 0.923722 | 0.000122 | 0.062408 | Bo1g058980.1 |
| 45 | . | gene | MSTRG.20882 | 2.562307 | 0.000145 | 0.068149 | Bo6g067990.1 |
| 46 | . | gene | MSTRG.31954 | 1.252355 | 0.000146 | 0.068149 | Bo9g104940.1 |
| 43 | . | gene | MSTRG.16806 | 41.23197 | 0.000139 | 0.068149 | |
| 44 | . | gene | MSTRG.30961 | 0.936967 | 0.000141 | 0.068149 | |
| 47 | . | gene | MSTRG.6996 | 0.410603 | 0.000155 | 0.070696 | |
| 48 | . | gene | MSTRG.14974 | 0.23611 | 0.000169 | 0.073966 | Bo4g154820.1 |
| 49 | . | gene | MSTRG.27395 | 3.057989 | 0.000171 | 0.073966 | Bo8g060110.1 |
| 50 | . | gene | MSTRG.11437 | 1.666328 | 0.000172 | 0.073966 | |
| 52 | . | gene | MSTRG.15327 | 1.771677 | 0.000184 | 0.07592 | Bo4g177870.1 |
| 51 | . | gene | MSTRG.13700 | 0.391954 | 0.000184 | 0.07592 | |
| 53 | . | gene | MSTRG.15280 | 0.195836 | 0.000199 | 0.08042 | Bo4g176140.1 |
| 55 | . | gene | MSTRG.6604 | 4.793871 | 0.000211 | 0.082292 | Bo2g164050.1 |

| | | | | | | | |
|---|---|---|---|---|---|---|---|
| 54 | . | gene | MSTRG.7810 | 17.25744 | 0.000211 | 0.082292 | |
| 56 | . | gene | MSTRG.17324 | 0.571495 | 0.000215 | 0.082292 | |
| 57 | . | gene | MSTRG.17058 | 18.68938 | 0.000224 | 0.084423 | Bo5g025110.1 |
| 58 | . | gene | MSTRG.9913 | 46.52852 | 0.000232 | 0.085944 | Bo3g078820.1 |
| 59 | . | gene | MSTRG.11849 | 1.319412 | 0.000242 | 0.086692 | Bo3g173910.1 |
| 61 | . | gene | MSTRG.15824 | 0.3603 | 0.000247 | 0.086692 | Bo4g191730.1 |
| 60 | . | gene | MSTRG.34006 | 0.483756 | 0.000244 | 0.086692 | |
| 62 | . | gene | MSTRG.22628 | 2.697272 | 0.000251 | 0.086773 | Bo6g121920.1 |
| 63 | . | gene | MSTRG.24427 | 0.294983 | 0.000264 | 0.089822 | |
| 64 | . | gene | MSTRG.16228 | 1.580321 | 0.000272 | 0.091145 | Bo5g005120.1 |
| 65 | . | gene | MSTRG.27599 | 1.141442 | 0.000277 | 0.091519 | Bo8g067280.1 |
| 66 | . | gene | MSTRG.28700 | 3.321822 | 0.000289 | 0.093794 | Bo8g097860.1 |
| 68 | . | gene | MSTRG.975 | 1.166076 | 0.000305 | 0.095182 | Bo1g023840.1 |
| 67 | . | gene | MSTRG.15110 | 0.701226 | 0.000305 | 0.095182 | Bo4g165020.1 |
| 70 | . | gene | MSTRG.16075 | 1.259 | 0.000311 | 0.095182 | Bo5g002760.1 |
| 69 | . | gene | MSTRG.19721 | 1.831299 | 0.00031 | 0.095182 | Bo5g150260.1 |
| 74 | . | gene | MSTRG.9163 | 0.31496 | 0.000329 | 0.095324 | Bo3g055290.1 |
| 71 | . | gene | MSTRG.11574 | 0.226512 | 0.000321 | 0.095324 | Bo3g160920.1 |
| 72 | . | gene | MSTRG.34379 | 2.255425 | 0.000328 | 0.095324 | |
| 73 | . | gene | MSTRG.18426 | 0.855 | 0.000329 | 0.095324 | |
| 75 | . | gene | MSTRG.11582 | 18.55792 | 0.000338 | 0.096681 | Bo3g162170.1 |
| 76 | . | gene | MSTRG.24851 | 1.948459 | 0.000347 | 0.097894 | Bo7g091170.1 |
| 77 | . | gene | MSTRG.13025 | 1.502824 | 0.000354 | 0.098656 | Bo4g040050.1 |
| 80 | . | gene | MSTRG.27315 | 1.369334 | 0.000372 | 0.09986 | Bo8g058870.1 |
| 78 | . | gene | MSTRG.22065 | 1.014666 | 0.000364 | 0.09986 | |
| 79 | . | gene | MSTRG.24884 | 0.443373 | 0.000371 | 0.09986 | |
| 91 | . | gene | MSTRG.1119 | 1.341703 | 0.000448 | 0.101371 | Bo1g032040.1 |
| 103 | . | gene | MSTRG.6135 | 1.547492 | 0.000491 | 0.101371 | Bo2g141720.1 |
| 102 | . | gene | MSTRG.6232 | 0.059099 | 0.000486 | 0.101371 | Bo2g148170.1 |

| | | | | | | | |
|---|---|---|---|---|---|---|---|
| 98 | . | gene | MSTRG.6534 | 1.230759 | 0.000477 | 0.101371 | Bo2g160830.1 |
| 96 | . | gene | MSTRG.6574 | 1.905686 | 0.000476 | 0.101371 | Bo2g161590.1 |
| 100 | . | gene | MSTRG.7440 | 13.56829 | 0.00048 | 0.101371 | Bo3g013550.1 |
| 82 | . | gene | MSTRG.8080 | 1.262938 | 0.000403 | 0.101371 | Bo3g026790.1 |
| 99 | . | gene | MSTRG.9636 | 0.648398 | 0.000479 | 0.101371 | Bo3g066380.1 |
| 92 | . | gene | MSTRG.10077 | 2.234666 | 0.000457 | 0.101371 | Bo3g084170.1 |
| 93 | . | gene | MSTRG.10142 | 0.838409 | 0.000459 | 0.101371 | Bo3g087020.1 |
| 83 | . | gene | MSTRG.14702 | 5.863401 | 0.000408 | 0.101371 | Bo4g142280.1 |
| 81 | . | gene | MSTRG.14933 | 6.289984 | 0.0004 | 0.101371 | Bo4g154050.1 |
| 84 | . | gene | MSTRG.15840 | 4.965684 | 0.00041 | 0.101371 | Bo4g192990.1 |
| 87 | . | gene | MSTRG.16306 | 7.3394 | 0.000426 | 0.101371 | Bo5g007360.1 |
| 97 | . | gene | MSTRG.19725 | 13.69625 | 0.000476 | 0.101371 | Bo5g150300.1 |
| 95 | . | gene | MSTRG.24737 | 1.961586 | 0.000473 | 0.101371 | Bo7g088600.1 |
| 86 | . | gene | MSTRG.29972 | 1.532663 | 0.000424 | 0.101371 | Bo9g008180.1 |
| 85 | . | gene | MSTRG.31090 | 3.100781 | 0.00041 | 0.101371 | Bo9g055660.1 |
| 88 | . | gene | MSTRG.20091 | 1.277484 | 0.000434 | 0.101371 | |
| 89 | . | gene | MSTRG.2560 | 6.226671 | 0.000438 | 0.101371 | |
| 90 | . | gene | MSTRG.20799 | 11.29974 | 0.000441 | 0.101371 | |
| 94 | . | gene | MSTRG.34372 | 0.098245 | 0.000462 | 0.101371 | |
| 101 | . | gene | MSTRG.14356 | 0.488342 | 0.000481 | 0.101371 | |
| 104 | . | gene | MSTRG.34432 | 0.476348 | 0.000492 | 0.101371 | |
| 105 | . | gene | MSTRG.23031 | 2.265188 | 0.000501 | 0.102259 | Bo7g014200.1 |
| 106 | . | gene | MSTRG.30929 | 0.671152 | 0.000506 | 0.102322 | Bo9g042120.1 |
| 109 | . | gene | MSTRG.2040 | 1.352792 | 0.000521 | 0.102559 | Bo1g086220.1 |
| 107 | . | gene | MSTRG.7300 | 7.412906 | 0.000521 | 0.102559 | Bo3g010560.1 |
| 108 | . | gene | MSTRG.27168 | 0.517802 | 0.000521 | 0.102559 | Bo8g051420.1 |
| 111 | . | gene | MSTRG.7466 | 0.528864 | 0.000544 | 0.105015 | Bo3g013900.1 |
| 110 | . | gene | MSTRG.27894 | 0.259669 | 0.000543 | 0.105015 | Bo8g077160.1 |
| 112 | TUB6 | gene | MSTRG.33231 | 1.609084 | 0.000551 | 0.105566 | Bo9g169510.1 |

| | | | | | | | |
|---|---|---|---|---|---|---|---|
| 113 | . | gene | MSTRG.21031 | 1.869659 | 0.000557 | 0.105756 | Bo6g074400.1 |
| 117 | . | gene | MSTRG.8696 | 0.834845 | 0.00059 | 0.106418 | Bo3g039740.1 |
| 118 | . | gene | MSTRG.9448 | 2.018967 | 0.000594 | 0.106418 | Bo3g062750.1 |
| 123 | . | gene | MSTRG.13089 | 2.220166 | 0.000615 | 0.106418 | Bo4g043950.1 |
| 116 | OWL1 | gene | MSTRG.15493 | 1.531256 | 0.000588 | 0.106418 | Bo4g184560.1 |
| 122 | . | gene | MSTRG.15723 | 2.275917 | 0.000614 | 0.106418 | Bo4g190190.1 |
| 124 | . | gene | MSTRG.20732 | 0.183286 | 0.000615 | 0.106418 | Bo6g064780.1 |
| 114 | . | gene | MSTRG.27370 | 0.428689 | 0.000575 | 0.106418 | Bo8g059680.1 |
| 120 | . | gene | MSTRG.33411 | 0.82669 | 0.0006 | 0.106418 | Bo9g174050.1 |
| 115 | . | gene | MSTRG.35655 | 2.472734 | 0.000581 | 0.106418 | |
| 119 | . | gene | MSTRG.35528 | 0.641364 | 0.000595 | 0.106418 | |
| 121 | . | gene | MSTRG.34447 | 0.803551 | 0.000602 | 0.106418 | |
| 125 | . | gene | MSTRG.24561 | 66.19068 | 0.000621 | 0.10655 | Bo7g082200.1 |
| 126 | . | gene | MSTRG.14660 | 0.283331 | 0.000626 | 0.106588 | |
| 127 | . | gene | MSTRG.33666 | 0.826223 | 0.000638 | 0.107489 | Bo9g177660.1 |
| 128 | . | gene | MSTRG.15027 | 14.57431 | 0.000642 | 0.107489 | |
| 129 | . | gene | MSTRG.7105 | 2.406138 | 0.000659 | 0.108214 | Bo3g007310.1 |
| 136 | . | gene | MSTRG.14854 | 3.111281 | 0.000688 | 0.108214 | Bo4g150920.1 |
| 139 | PAK | gene | MSTRG.16708 | 0.284722 | 0.000701 | 0.108214 | Bo5g016880.1 |
| 138 | . | gene | MSTRG.18721 | 1.691797 | 0.0007 | 0.108214 | Bo5g118820.1 |
| 130 | NAKR3 | gene | MSTRG.20843 | 0.425196 | 0.00066 | 0.108214 | Bo6g067540.1 |
| 135 | . | gene | MSTRG.21077 | 0.347047 | 0.000688 | 0.108214 | Bo6g075980.1 |
| 133 | . | gene | MSTRG.22627 | 0.468402 | 0.000683 | 0.108214 | Bo6g121910.1 |
| 132 | . | gene | MSTRG.25383 | 1.201209 | 0.000681 | 0.108214 | Bo7g104280.1 |
| 131 | . | gene | MSTRG.27660 | 1.832162 | 0.000663 | 0.108214 | Bo8g068300.1 |
| 134 | . | gene | MSTRG.11071 | 0.411994 | 0.000687 | 0.108214 | |
| 137 | . | gene | MSTRG.23716 | 0.517808 | 0.000697 | 0.108214 | |
| 140 | . | gene | MSTRG.11451 | 0.213219 | 0.00072 | 0.110307 | Bo3g153740.1 |
| 141 | . | gene | MSTRG.25561 | 1.590739 | 0.000734 | 0.111014 | Bo7g107450.1 |

| | | | | | | | |
|---|---|---|---|---|---|---|---|
| 142 | . | gene | MSTRG.26929 | 0.655475 | 0.000735 | 0.111014 | Bo8g038070.1 |
| 144 | . | gene | MSTRG.18111 | 1.489264 | 0.000745 | 0.111015 | Bo5g079950.1 |
| 145 | . | gene | MSTRG.19744 | 0.468784 | 0.000751 | 0.111015 | Bo5g151520.1 |
| 143 | . | gene | MSTRG.11556 | 5.063255 | 0.000741 | 0.111015 | |
| 147 | . | gene | MSTRG.13086 | 2.565554 | 0.000778 | 0.11358 | Bo4g043860.1 |
| 146 | . | gene | MSTRG.15798 | 1.863959 | 0.000773 | 0.11358 | Bo4g191310.1 |
| 148 | SAR1 | gene | MSTRG.8095 | 1.865913 | 0.000794 | 0.11437 | Bo3g027140.1 |
| 149 | . | gene | MSTRG.10907 | 0.474216 | 0.000798 | 0.11437 | Bo3g129970.1 |
| 150 | . | gene | MSTRG.32670 | 0.401863 | 0.0008 | 0.11437 | Bo9g150100.1 |
| 161 | . | gene | MSTRG.5916 | 2.403095 | 0.000863 | 0.114995 | Bo2g133750.1 |
| 156 | . | gene | MSTRG.9062 | 1.169635 | 0.000855 | 0.114995 | Bo3g052850.1 |
| 153 | VSR6 | gene | MSTRG.17903 | 1.834455 | 0.000849 | 0.114995 | Bo5g065590.1 |
| 151 | OSB4 | gene | MSTRG.17906 | 1.490812 | 0.000811 | 0.114995 | Bo5g065630.1 |
| 157 | . | gene | MSTRG.18489 | 17.29784 | 0.000856 | 0.114995 | Bo5g108480.1 |
| 154 | . | gene | MSTRG.22229 | 1.241081 | 0.000851 | 0.114995 | Bo6g113350.1 |
| 160 | . | gene | MSTRG.27244 | 1.323072 | 0.000863 | 0.114995 | Bo8g055840.1 |
| 152 | . | gene | MSTRG.31714 | 0.48588 | 0.000842 | 0.114995 | |
| 155 | . | gene | MSTRG.14241 | 0.622923 | 0.000853 | 0.114995 | |
| 158 | . | gene | MSTRG.35154 | 0.333691 | 0.000858 | 0.114995 | |
| 159 | . | gene | MSTRG.32694 | 5.249163 | 0.000859 | 0.114995 | |
| 162 | . | gene | MSTRG.19993 | 6.283264 | 0.000876 | 0.116007 | Bo6g018030.1 |
| 175 | . | gene | MSTRG.664 | 0.727947 | 0.000978 | 0.117539 | Bo1g016170.1 |
| 195 | . | gene | MSTRG.1143 | 10.95137 | 0.00108 | 0.117539 | Bo1g034450.1 |
| 198 | . | gene | MSTRG.1708 | 1.628496 | 0.001108 | 0.117539 | Bo1g057270.1 |
| 185 | . | gene | MSTRG.2828 | 1.252306 | 0.00104 | 0.117539 | Bo1g138450.1 |
| 202 | . | gene | MSTRG.6632 | 0.472414 | 0.001117 | 0.117539 | Bo2g165480.1 |
| 193 | . | gene | MSTRG.6640 | 0.370459 | 0.001072 | 0.117539 | Bo2g165620.1 |
| 199 | . | gene | MSTRG.7166 | 0.6665 | 0.001111 | 0.117539 | Bo3g009190.1 |
| 197 | . | gene | MSTRG.9076 | 0.189881 | 0.001097 | 0.117539 | Bo3g053080.1 |

| | | | | | | | |
|---|---|---|---|---|---|---|---|
| 169 | . | gene | MSTRG.9419 | 0.217305 | 0.000943 | 0.117539 | Bo3g062290.1 |
| 164 | . | gene | MSTRG.9756 | 1.70848 | 0.000905 | 0.117539 | Bo3g070280.1 |
| 179 | . | gene | MSTRG.9860 | 0.387733 | 0.000999 | 0.117539 | Bo3g076030.1 |
| 182 | . | gene | MSTRG.10609 | 2.740065 | 0.001009 | 0.117539 | Bo3g107390.1 |
| 192 | . | gene | MSTRG.11949 | 1.229058 | 0.001064 | 0.117539 | Bo3g178330.1 |
| 183 | . | gene | MSTRG.12514 | 1.231962 | 0.001012 | 0.117539 | Bo4g023970.1 |
| 184 | . | gene | MSTRG.15168 | 0.238047 | 0.001031 | 0.117539 | Bo4g169230.1 |
| 191 | . | gene | MSTRG.16954 | 1.699527 | 0.001061 | 0.117539 | Bo5g021620.1 |
| 178 | . | gene | MSTRG.16975 | 4.358534 | 0.000999 | 0.117539 | Bo5g021940.1 |
| 171 | . | gene | MSTRG.20404 | 1.529571 | 0.000966 | 0.117539 | Bo6g040080.1 |
| 180 | . | gene | MSTRG.23242 | 1.446394 | 0.001002 | 0.117539 | Bo7g029070.1 |
| 201 | . | gene | MSTRG.24721 | 9.113334 | 0.001117 | 0.117539 | Bo7g087330.1 |
| 168 | . | gene | MSTRG.25589 | 0.452988 | 0.000939 | 0.117539 | Bo7g107910.1 |
| 190 | . | gene | MSTRG.26372 | 1.439656 | 0.001057 | 0.117539 | Bo8g004270.1 |
| 173 | . | gene | MSTRG.27751 | 0.202619 | 0.00097 | 0.117539 | Bo8g070810.1 |
| 181 | . | gene | MSTRG.28324 | 2.125698 | 0.001009 | 0.117539 | Bo8g090000.1 |
| 166 | . | gene | MSTRG.28766 | 0.710074 | 0.000918 | 0.117539 | Bo8g098800.1 |
| 177 | . | gene | MSTRG.30209 | 2.233635 | 0.000994 | 0.117539 | Bo9g014060.1 |
| 186 | . | gene | MSTRG.30978 | 2.014606 | 0.00104 | 0.117539 | Bo9g044120.1 |
| 174 | . | gene | MSTRG.31198 | 7.555908 | 0.000971 | 0.117539 | Bo9g059410.1 |
| 189 | . | gene | MSTRG.31214 | 1.797165 | 0.001052 | 0.117539 | Bo9g059640.1 |
| 204 | . | gene | MSTRG.33304 | 0.587132 | 0.001123 | 0.117539 | Bo9g171600.1 |
| 163 | . | gene | MSTRG.10545 | 0.16387 | 0.000895 | 0.117539 | |
| 165 | . | gene | MSTRG.14545 | 2.835223 | 0.000908 | 0.117539 | |
| 167 | . | gene | MSTRG.17035 | 0.626133 | 0.000937 | 0.117539 | |
| 170 | . | gene | MSTRG.35493 | 31.82212 | 0.000965 | 0.117539 | |
| 172 | . | gene | MSTRG.34012 | 3.479962 | 0.000967 | 0.117539 | |
| 176 | . | gene | MSTRG.6501 | 0.693047 | 0.000985 | 0.117539 | |
| 187 | . | gene | MSTRG.34545 | 2.293434 | 0.001047 | 0.117539 | |

| | | | | | | | |
|---|---|---|---|---|---|---|---|
| 188 | . | gene | MSTRG.32263 | 0.119421 | 0.001049 | 0.117539 | |
| 194 | . | gene | MSTRG.24148 | 0.681296 | 0.001072 | 0.117539 | |
| 196 | . | gene | MSTRG.6357 | 0.99221 | 0.001089 | 0.117539 | |
| 200 | . | gene | MSTRG.35232 | 3.015923 | 0.001111 | 0.117539 | |
| 203 | . | gene | MSTRG.35198 | 5.697741 | 0.001123 | 0.117539 | |
| 205 | . | gene | MSTRG.34814 | 0.568888 | 0.001123 | 0.117539 | |
| 206 | . | gene | MSTRG.13109 | 2.287957 | 0.00113 | 0.117684 | Bo4g045340.1 |
| 207 | . | gene | MSTRG.19196 | 0.546822 | 0.001136 | 0.117731 | Bo5g135000.1 |
| 213 | . | gene | MSTRG.714 | 1.249885 | 0.001177 | 0.117803 | Bo1g016890.1 |
| 215 | . | gene | MSTRG.2758 | 0.341769 | 0.001192 | 0.117803 | Bo1g131210.1 |
| 208 | . | gene | MSTRG.6557 | 0.578087 | 0.001149 | 0.117803 | Bo2g161210.1 |
| 217 | . | gene | MSTRG.13306 | 0.346106 | 0.001196 | 0.117803 | Bo4g059540.1 |
| 210 | . | gene | MSTRG.22502 | 0.928218 | 0.00117 | 0.117803 | Bo6g120300.1 |
| 218 | . | gene | MSTRG.23007 | 1.444587 | 0.0012 | 0.117803 | Bo7g012730.1 |
| 209 | . | gene | MSTRG.24736 | 0.478804 | 0.001151 | 0.117803 | Bo7g088570.1 |
| 220 | . | gene | MSTRG.26595 | 0.579287 | 0.001208 | 0.117803 | Bo8g016860.1 |
| 211 | . | gene | MSTRG.34427 | 2.368336 | 0.001171 | 0.117803 | |
| 212 | . | gene | MSTRG.31941 | 0.793132 | 0.001175 | 0.117803 | |
| 214 | . | gene | MSTRG.6669 | 0.418181 | 0.001186 | 0.117803 | |
| 216 | . | gene | MSTRG.36137 | 0.798399 | 0.001196 | 0.117803 | |
| 219 | . | gene | MSTRG.29380 | 1.741816 | 0.001206 | 0.117803 | |
| 221 | . | gene | MSTRG.31443 | 0.962523 | 0.001235 | 0.119851 | Bo9g074420.1 |
| 222 | . | gene | MSTRG.30189 | 1.688743 | 0.001252 | 0.120921 | Bo9g013790.1 |
| 223 | . | gene | MSTRG.16945 | 0.451238 | 0.001275 | 0.122626 | Bo5g021540.1 |
| 224 | . | gene | MSTRG.15813 | 0.788064 | 0.001284 | 0.122945 | |
| 225 | . | gene | MSTRG.19345 | 1.512071 | 0.001291 | 0.123047 | Bo5g138120.1 |
| 226 | . | gene | MSTRG.10102 | 0.444898 | 0.001301 | 0.1235 | Bo3g085530.1 |
| 227 | . | gene | MSTRG.3658 | 0.401099 | 0.001324 | 0.124972 | Bo2g011240.1 |
| 228 | . | gene | MSTRG.11037 | 0.578592 | 0.001331 | 0.124972 | |

| | | | | | | | |
|---|---|---|---|---|---|---|---|
| 229 | . | gene | MSTRG.14954 | 0.366393 | 0.001334 | 0.124972 | |
| 230 | . | gene | MSTRG.7268 | 0.243527 | 0.001344 | 0.125357 | Bo3g010260.1 |
| 231 | . | gene | MSTRG.26992 | 1.251879 | 0.001361 | 0.125857 | Bo8g043260.1 |
| 232 | . | gene | MSTRG.27427 | 0.542531 | 0.001361 | 0.125857 | Bo8g062600.1 |
| 233 | . | gene | MSTRG.13342 | 0.162789 | 0.001368 | 0.125911 | |
| 239 | . | gene | MSTRG.11517 | 0.651262 | 0.00141 | 0.126167 | Bo3g155750.1 |
| 235 | . | gene | MSTRG.12756 | 0.424381 | 0.001391 | 0.126167 | Bo4g029780.1 |
| 242 | . | gene | MSTRG.15218 | 2.462616 | 0.001425 | 0.126167 | Bo4g173200.1 |
| 237 | . | gene | MSTRG.17242 | 0.388263 | 0.001397 | 0.126167 | Bo5g031980.1 |
| 234 | . | gene | MSTRG.24804 | 1.283334 | 0.001382 | 0.126167 | Bo7g089470.1 |
| 238 | . | gene | MSTRG.30864 | 0.127561 | 0.001407 | 0.126167 | Bo9g036860.1 |
| 244 | . | gene | MSTRG.33364 | 70.38799 | 0.001435 | 0.126167 | Bo9g173400.1 |
| 236 | . | gene | MSTRG.31958 | 1.234636 | 0.001396 | 0.126167 | |
| 240 | . | gene | MSTRG.17504 | 1.368711 | 0.001422 | 0.126167 | |
| 241 | . | gene | MSTRG.19615 | 1.077957 | 0.001423 | 0.126167 | |
| 243 | . | gene | MSTRG.34592 | 0.636685 | 0.001433 | 0.126167 | |
| 248 | . | gene | MSTRG.2259 | 0.888734 | 0.001475 | 0.126888 | Bo1g101410.1 |
| 252 | . | gene | MSTRG.7478 | 0.238048 | 0.001491 | 0.126888 | Bo3g014050.1 |
| 249 | . | gene | MSTRG.15722 | 6.14732 | 0.001486 | 0.126888 | Bo4g190180.1 |
| 247 | . | gene | MSTRG.17440 | 1.466786 | 0.001468 | 0.126888 | Bo5g039940.1 |
| 251 | . | gene | MSTRG.20143 | 6.257558 | 0.00149 | 0.126888 | Bo6g026490.1 |
| 246 | . | gene | MSTRG.24770 | 0.650365 | 0.001468 | 0.126888 | Bo7g089060.1 |
| 245 | . | gene | MSTRG.27445 | 0.082734 | 0.001465 | 0.126888 | Bo8g064840.1 |
| 250 | . | gene | MSTRG.2925 | 0.969113 | 0.001489 | 0.126888 | |
| 253 | . | gene | MSTRG.31088 | 1.63123 | 0.0015 | 0.127121 | Bo9g055640.1 |
| 262 | . | gene | MSTRG.3922 | 1.5606 | 0.001554 | 0.127239 | Bo2g018540.1 |
| 260 | . | gene | MSTRG.4357 | 3.396673 | 0.001546 | 0.127239 | Bo2g041820.1 |
| 256 | . | gene | MSTRG.12553 | 0.790066 | 0.001531 | 0.127239 | Bo4g024540.1 |
| 255 | . | gene | MSTRG.24060 | 0.457275 | 0.001524 | 0.127239 | Bo7g064910.1 |

| | | | | | | | |
|---|---|---|---|---|---|---|---|
| 258 | . | gene | MSTRG.30877 | 4.010237 | 0.001544 | 0.127239 | Bo9g037180.1 |
| 261 | . | gene | MSTRG.31217 | 0.357179 | 0.001553 | 0.127239 | Bo9g059740.1 |
| 259 | . | gene | MSTRG.32361 | 1.24313 | 0.001544 | 0.127239 | Bo9g133060.1 |
| 254 | . | gene | MSTRG.13505 | 0.799499 | 0.001516 | 0.127239 | |
| 257 | . | gene | MSTRG.23837 | 0.142495 | 0.001535 | 0.127239 | |
| 264 | . | gene | MSTRG.3436 | 0.810989 | 0.001575 | 0.127958 | Bo2g006750.1 |
| 263 | . | gene | MSTRG.5619 | 2.368024 | 0.00157 | 0.127958 | |
| 271 | . | gene | MSTRG.284 | 0.074991 | 0.001628 | 0.128046 | Bo1g006790.1 |
| 277 | . | gene | MSTRG.17775 | 1.410233 | 0.001661 | 0.128046 | Bo5g058440.1 |
| 268 | . | gene | MSTRG.18112 | 6.532048 | 0.001615 | 0.128046 | Bo5g079960.1 |
| 281 | . | gene | MSTRG.21036 | 15.0646 | 0.001679 | 0.128046 | Bo6g074480.1 |
| 266 | . | gene | MSTRG.21761 | 2.533038 | 0.001605 | 0.128046 | Bo6g097380.1 |
| 274 | . | gene | MSTRG.22857 | 0.564692 | 0.001651 | 0.128046 | Bo7g005380.1 |
| 270 | . | gene | MSTRG.24908 | 163.1213 | 0.001627 | 0.128046 | Bo7g093190.1 |
| 273 | . | gene | MSTRG.25275 | 2.627104 | 0.001645 | 0.128046 | Bo7g100630.1 |
| 278 | . | gene | MSTRG.26611 | 1.63515 | 0.001666 | 0.128046 | Bo8g019180.1 |
| 267 | . | gene | MSTRG.28082 | 1.333724 | 0.00161 | 0.128046 | Bo8g081420.1 |
| 276 | . | gene | MSTRG.30547 | 1.490144 | 0.001659 | 0.128046 | Bo9g024090.1 |
| 265 | . | gene | MSTRG.16584 | 0.474379 | 0.001601 | 0.128046 | |
| 269 | . | gene | MSTRG.4302 | 1.307033 | 0.001622 | 0.128046 | |
| 272 | . | gene | MSTRG.34759 | 5.353863 | 0.001636 | 0.128046 | |
| 275 | . | gene | MSTRG.16470 | 1.704715 | 0.001652 | 0.128046 | |
| 279 | . | gene | MSTRG.34264 | 0.53748 | 0.001673 | 0.128046 | |
| 280 | . | gene | MSTRG.32022 | 1.382152 | 0.001678 | 0.128046 | |
| 282 | . | gene | MSTRG.7756 | 0.537337 | 0.001685 | 0.128046 | |
| 283 | . | gene | MSTRG.28124 | 0.288604 | 0.00169 | 0.128046 | |
| 285 | . | gene | MSTRG.1276 | 1.575545 | 0.001712 | 0.128329 | Bo1g039350.1 |
| 287 | . | gene | MSTRG.18678 | 1.5994 | 0.001717 | 0.128329 | Bo5g117120.1 |
| 284 | . | gene | MSTRG.27100 | 1.390676 | 0.001711 | 0.128329 | Bo8g050310.1 |

| | | | | | | | |
|---|---|---|---|---|---|---|---|
| 286 | . | gene | MSTRG.25285 | 1.803309 | 0.001713 | 0.128329 | |
| 291 | . | gene | MSTRG.7380 | 0.340443 | 0.001747 | 0.128773 | Bo3g012640.1 |
| 288 | . | gene | MSTRG.19061 | 2.052214 | 0.001734 | 0.128773 | Bo5g131810.1 |
| 290 | . | gene | MSTRG.20798 | 0.325436 | 0.001741 | 0.128773 | Bo6g066750.1 |
| 289 | . | gene | MSTRG.24941 | 0.343701 | 0.001736 | 0.128773 | Bo7g093620.1 |
| 296 | Nek7 | gene | MSTRG.2893 | 1.621318 | 0.001802 | 0.129702 | Bo1g139510.1 |
| 298 | . | gene | MSTRG.10495 | 1.30681 | 0.001806 | 0.129702 | Bo3g101680.1 |
| 295 | . | gene | MSTRG.16324 | 0.713512 | 0.001793 | 0.129702 | Bo5g007570.1 |
| 293 | . | gene | MSTRG.27432 | 1.453486 | 0.001782 | 0.129702 | Bo8g063710.1 |
| 292 | . | gene | MSTRG.27722 | 15.23901 | 0.001769 | 0.129702 | Bo8g070380.1 |
| 294 | . | gene | MSTRG.35994 | 0.234756 | 0.001791 | 0.129702 | |
| 297 | . | gene | MSTRG.15221 | 0.990509 | 0.001804 | 0.129702 | |
| 299 | . | gene | MSTRG.10989 | 8.280622 | 0.001808 | 0.129702 | |
| 301 | . | gene | MSTRG.8511 | 0.3878 | 0.001829 | 0.13029 | Bo3g036020.1 |
| 300 | . | gene | MSTRG.16512 | 0.823605 | 0.001827 | 0.13029 | |
| 303 | . | gene | MSTRG.3297 | 1.355623 | 0.001841 | 0.130327 | Bo1g157520.1 |
| 302 | . | gene | MSTRG.35391 | 1.17951 | 0.001838 | 0.130327 | |
| 305 | . | gene | MSTRG.6740 | 0.284081 | 0.001856 | 0.130535 | Bo2g168080.1 |
| 304 | . | gene | MSTRG.19820 | 1.681262 | 0.001851 | 0.130535 | Bo5g156500.1 |
| 311 | . | gene | MSTRG.1492 | 8.014604 | 0.001904 | 0.13112 | Bo1g049430.1 |
| 309 | . | gene | MSTRG.2353 | 1.399904 | 0.0019 | 0.13112 | Bo1g105200.1 |
| 306 | . | gene | MSTRG.15934 | 0.662738 | 0.001871 | 0.13112 | Bo4g195730.1 |
| 310 | . | gene | MSTRG.32668 | 2.022843 | 0.0019 | 0.13112 | Bo9g150080.1 |
| 307 | . | gene | MSTRG.12906 | 0.45137 | 0.001888 | 0.13112 | |
| 308 | . | gene | MSTRG.6247 | 0.512412 | 0.001888 | 0.13112 | |
| 312 | . | gene | MSTRG.5005 | 0.203479 | 0.001907 | 0.13112 | |
| 313 | . | gene | MSTRG.6038 | 0.444713 | 0.001928 | 0.131966 | Bo2g137020.1 |
| 314 | . | gene | MSTRG.31742 | 0.936607 | 0.001932 | 0.131966 | |
| 315 | . | gene | MSTRG.27963 | 0.282268 | 0.001952 | 0.13288 | Bo8g079230.1 |

| | | | | | | | |
|---|---|---|---|---|---|---|---|
| 353 | . | gene | MSTRG.3082 | 0.549616 | 0.002211 | 0.133735 | Bo1g147350.1 |
| 339 | . | gene | MSTRG.3626 | 0.462094 | 0.002128 | 0.133735 | Bo2g010720.1 |
| 337 | . | gene | MSTRG.4119 | 1.658533 | 0.002119 | 0.133735 | Bo2g025570.1 |
| 326 | . | gene | MSTRG.6077 | 0.287759 | 0.00207 | 0.133735 | Bo2g138750.1 |
| 343 | . | gene | MSTRG.6671 | 1.404342 | 0.002152 | 0.133735 | Bo2g166100.1 |
| 335 | . | gene | MSTRG.6757 | 0.255621 | 0.002095 | 0.133735 | Bo3g001090.1 |
| 327 | . | gene | MSTRG.7283 | 0.670118 | 0.002076 | 0.133735 | Bo3g010410.1 |
| 324 | . | gene | MSTRG.7499 | 2.383779 | 0.002044 | 0.133735 | Bo3g015380.1 |
| 348 | . | gene | MSTRG.10908 | 1.443602 | 0.002191 | 0.133735 | Bo3g130010.1 |
| 322 | . | gene | MSTRG.11104 | 0.623237 | 0.00202 | 0.133735 | Bo3g140020.1 |
| 319 | . | gene | MSTRG.15017 | 2.414558 | 0.001998 | 0.133735 | Bo4g158770.1 |
| 357 | . | gene | MSTRG.15244 | 8.567491 | 0.002235 | 0.133735 | Bo4g173560.1 |
| 349 | . | gene | MSTRG.15898 | 8.579741 | 0.002192 | 0.133735 | Bo4g195120.1 |
| 320 | . | gene | MSTRG.17128 | 7.839464 | 0.00201 | 0.133735 | Bo5g027470.1 |
| 359 | . | gene | MSTRG.17322 | 1.664849 | 0.002249 | 0.133735 | Bo5g035020.1 |
| 351 | . | gene | MSTRG.17634 | 0.408416 | 0.002203 | 0.133735 | Bo5g049050.1 |
| 328 | . | gene | MSTRG.17648 | 4.671861 | 0.002083 | 0.133735 | Bo5g049240.1 |
| 329 | . | gene | MSTRG.17818 | 1.356781 | 0.002089 | 0.133735 | Bo5g061140.1 |
| 345 | PKp3 | gene | MSTRG.18220 | 1.37764 | 0.002168 | 0.133735 | Bo5g087140.1 |
| 334 | . | gene | MSTRG.18871 | 1.623178 | 0.002093 | 0.133735 | Bo5g126240.1 |
| 330 | . | gene | MSTRG.20802 | 0.468766 | 0.002089 | 0.133735 | Bo6g066830.1 |
| 325 | . | gene | MSTRG.24523 | 4.720431 | 0.002053 | 0.133735 | Bo7g081440.1 |
| 318 | . | gene | MSTRG.26333 | 0.687709 | 0.001988 | 0.133735 | Bo8g002730.1 |
| 344 | . | gene | MSTRG.27128 | 0.410284 | 0.002156 | 0.133735 | Bo8g050810.1 |
| 354 | . | gene | MSTRG.28575 | 1.745221 | 0.002212 | 0.133735 | Bo8g094670.1 |
| 336 | . | gene | MSTRG.29206 | 12.50076 | 0.002117 | 0.133735 | Bo8g106410.1 |
| 346 | ATARLA1B | gene | MSTRG.29444 | 0.473572 | 0.002169 | 0.133735 | Bo8g110970.1 |
| 317 | . | gene | MSTRG.30100 | 1.574842 | 0.001983 | 0.133735 | Bo9g010410.1 |
| 358 | . | gene | MSTRG.30404 | 0.94082 | 0.002242 | 0.133735 | Bo9g019160.1 |

| | | | | | | | |
|---|---|---|---|---|---|---|---|
| 361 | . | gene | MSTRG.30445 | 0.71511 | 0.002251 | 0.133735 | Bo9g021610.1 |
| 333 | . | gene | MSTRG.30656 | 0.589961 | 0.002092 | 0.133735 | Bo9g028080.1 |
| 332 | . | gene | MSTRG.32482 | 0.785021 | 0.00209 | 0.133735 | Bo9g136950.1 |
| 316 | . | gene | MSTRG.28155 | 4.942731 | 0.001979 | 0.133735 | |
| 321 | . | gene | MSTRG.20295 | 0.838101 | 0.002013 | 0.133735 | |
| 323 | . | gene | MSTRG.35810 | 53.7543 | 0.002036 | 0.133735 | |
| 331 | . | gene | MSTRG.33968 | 4.657625 | 0.002089 | 0.133735 | |
| 338 | . | gene | MSTRG.13902 | 0.562873 | 0.002127 | 0.133735 | |
| 340 | . | gene | MSTRG.2638 | 0.613926 | 0.002129 | 0.133735 | |
| 341 | . | gene | MSTRG.6670 | 0.20224 | 0.002137 | 0.133735 | |
| 342 | . | gene | MSTRG.11060 | 8.812401 | 0.002142 | 0.133735 | |
| 347 | . | gene | MSTRG.34508 | 2.694498 | 0.002185 | 0.133735 | |
| 350 | . | gene | MSTRG.30361 | 0.23843 | 0.002199 | 0.133735 | |
| 352 | . | gene | MSTRG.23237 | 0.442012 | 0.002204 | 0.133735 | |
| 355 | . | gene | MSTRG.14826 | 4.517705 | 0.002225 | 0.133735 | |
| 356 | . | gene | MSTRG.34002 | 1.316116 | 0.002232 | 0.133735 | |
| 360 | . | gene | MSTRG.27221 | 0.485475 | 0.002249 | 0.133735 | |
| 362 | . | gene | MSTRG.11564 | 1.418977 | 0.002281 | 0.134824 | Bo3g158710.1 |
| 363 | . | gene | MSTRG.15839 | 3.116407 | 0.002282 | 0.134824 | |
| 364 | . | gene | MSTRG.28232 | 0.236063 | 0.002291 | 0.134978 | Bo8g083670.1 |
| 365 | . | gene | MSTRG.12840 | 6.357089 | 0.002302 | 0.135278 | Bo4g032370.1 |
| 374 | . | gene | MSTRG.2046 | 1.361094 | 0.002386 | 0.135563 | Bo1g087360.1 |
| 386 | . | gene | MSTRG.2398 | 0.320411 | 0.002446 | 0.135563 | Bo1g105930.1 |
| 369 | . | gene | MSTRG.2413 | 0.505559 | 0.002351 | 0.135563 | Bo1g107200.1 |
| 367 | . | gene | MSTRG.2955 | 1.218344 | 0.002347 | 0.135563 | Bo1g141360.1 |
| 372 | . | gene | MSTRG.3109 | 0.625816 | 0.002376 | 0.135563 | Bo1g147750.1 |
| 387 | . | gene | MSTRG.3452 | 1.275849 | 0.002451 | 0.135563 | Bo2g007060.1 |
| 377 | . | gene | MSTRG.3455 | 0.272093 | 0.002401 | 0.135563 | Bo2g007120.1 |
| 376 | . | gene | MSTRG.6558 | 6.780267 | 0.002398 | 0.135563 | Bo2g161230.1 |

| | | | | | | | |
|---|---|---|---|---|---|---|---|
| 371 | . | gene | MSTRG.7693 | 0.644899 | 0.002367 | 0.135563 | Bo3g018860.1 |
| 375 | . | gene | MSTRG.8072 | 0.60872 | 0.002389 | 0.135563 | Bo3g026700.1 |
| 382 | . | gene | MSTRG.9100 | 1.817837 | 0.00243 | 0.135563 | Bo3g054500.1 |
| 366 | . | gene | MSTRG.23384 | 1.327366 | 0.002329 | 0.135563 | Bo7g037600.1 |
| 390 | . | gene | MSTRG.25904 | 3.213064 | 0.002465 | 0.135563 | Bo7g114140.1 |
| 380 | . | gene | MSTRG.26724 | 0.757219 | 0.002409 | 0.135563 | Bo8g027220.1 |
| 370 | . | gene | MSTRG.27880 | 4.166219 | 0.002352 | 0.135563 | Bo8g076910.1 |
| 383 | . | gene | MSTRG.28368 | 0.991954 | 0.002431 | 0.135563 | Bo8g090560.1 |
| 378 | . | gene | MSTRG.30562 | 2.168742 | 0.002406 | 0.135563 | Bo9g024390.1 |
| 368 | . | gene | MSTRG.9946 | 0.479197 | 0.002351 | 0.135563 | |
| 373 | . | gene | MSTRG.4675 | 0.598374 | 0.002376 | 0.135563 | |
| 379 | . | gene | MSTRG.31467 | 0.228873 | 0.002408 | 0.135563 | |
| 381 | . | gene | MSTRG.2011 | 0.9006 | 0.002425 | 0.135563 | |
| 384 | . | gene | MSTRG.6363 | 1.99501 | 0.002432 | 0.135563 | |
| 385 | . | gene | MSTRG.35477 | 6.294104 | 0.002443 | 0.135563 | |
| 388 | . | gene | MSTRG.23062 | 0.323276 | 0.002455 | 0.135563 | |
| 389 | . | gene | MSTRG.18576 | 0.522985 | 0.002459 | 0.135563 | |
| 391 | . | gene | MSTRG.7032 | 2.549461 | 0.002483 | 0.135872 | Bo3g005380.1 |
| 393 | . | gene | MSTRG.10558 | 0.328824 | 0.002496 | 0.135872 | Bo3g103590.1 |
| 394 | . | gene | MSTRG.14862 | 0.516113 | 0.002496 | 0.135872 | Bo4g151060.1 |
| 392 | . | gene | MSTRG.14114 | 0.430219 | 0.002489 | 0.135872 | |
| 397 | . | gene | MSTRG.23238 | 0.374024 | 0.002525 | 0.136417 | Bo7g029040.1 |
| 396 | . | gene | MSTRG.26067 | 0.21834 | 0.002524 | 0.136417 | Bo7g116470.1 |
| 395 | . | gene | MSTRG.20049 | 0.374379 | 0.002524 | 0.136417 | |
| 398 | . | gene | MSTRG.3669 | 0.37525 | 0.002538 | 0.136761 | |
| 399 | . | gene | MSTRG.34977 | 2.210201 | 0.002559 | 0.137527 | |
| 400 | . | gene | MSTRG.33999 | 1.204138 | 0.002592 | 0.138991 | |
| 405 | . | gene | MSTRG.1264 | 1.353935 | 0.002653 | 0.139356 | Bo1g039190.1 |
| 417 | . | gene | MSTRG.4519 | 1.772055 | 0.002729 | 0.139356 | Bo2g050820.1 |

| | | | | | | | |
|---|---|---|---|---|---|---|---|
| 403 | . | gene | MSTRG.6537 | 1.250079 | 0.002633 | 0.139356 | Bo2g160890.1 |
| 416 | . | gene | MSTRG.6661 | 1.615573 | 0.002727 | 0.139356 | Bo2g165950.1 |
| 404 | . | gene | MSTRG.6956 | 0.782851 | 0.002638 | 0.139356 | Bo3g004280.1 |
| 419 | . | gene | MSTRG.7020 | 2.460239 | 0.00274 | 0.139356 | Bo3g005200.1 |
| 423 | . | gene | MSTRG.9067 | 0.359664 | 0.002749 | 0.139356 | Bo3g052980.1 |
| 406 | . | gene | MSTRG.9136 | 0.653361 | 0.002655 | 0.139356 | Bo3g054920.1 |
| 407 | . | gene | MSTRG.13524 | 1.530132 | 0.002658 | 0.139356 | Bo4g076990.1 |
| 421 | . | gene | MSTRG.16203 | 1.296787 | 0.002744 | 0.139356 | Bo5g004820.1 |
| 420 | . | gene | MSTRG.16217 | 1.410196 | 0.00274 | 0.139356 | Bo5g005000.1 |
| 410 | . | gene | MSTRG.16283 | 0.736925 | 0.002679 | 0.139356 | Bo5g006960.1 |
| 408 | . | gene | MSTRG.25277 | 2.454954 | 0.002667 | 0.139356 | Bo7g100660.1 |
| 402 | . | gene | MSTRG.27720 | 3.333214 | 0.002629 | 0.139356 | Bo8g070360.1 |
| 422 | . | gene | MSTRG.29778 | 1.308616 | 0.002748 | 0.139356 | Bo9g003710.1 |
| 414 | . | gene | MSTRG.30167 | 0.37624 | 0.002716 | 0.139356 | Bo9g012560.1 |
| 411 | . | gene | MSTRG.31934 | 1.806892 | 0.00268 | 0.139356 | Bo9g103630.1 |
| 401 | . | gene | MSTRG.22961 | 0.728706 | 0.002626 | 0.139356 | |
| 409 | . | gene | MSTRG.1376 | 0.411385 | 0.002669 | 0.139356 | |
| 412 | . | gene | MSTRG.3231 | 0.868173 | 0.002684 | 0.139356 | |
| 413 | . | gene | MSTRG.20280 | 1.887552 | 0.002698 | 0.139356 | |
| 415 | . | gene | MSTRG.522 | 0.094425 | 0.002725 | 0.139356 | |
| 418 | . | gene | MSTRG.14870 | 0.224177 | 0.00273 | 0.139356 | |
| 424 | . | gene | MSTRG.13001 | 1.743318 | 0.002766 | 0.139494 | Bo4g039710.1 |
| 425 | . | gene | MSTRG.35090 | 0.881839 | 0.002767 | 0.139494 | |
| 426 | . | gene | MSTRG.34088 | 0.379595 | 0.002771 | 0.139494 | |
| 427 | . | gene | MSTRG.31680 | 1.091691 | 0.002779 | 0.139602 | |
| 429 | . | gene | MSTRG.10361 | 0.294759 | 0.002796 | 0.139742 | Bo3g097930.1 |
| 428 | . | gene | MSTRG.20288 | 1.957862 | 0.002789 | 0.139742 | Bo6g031760.1 |
| 430 | . | gene | MSTRG.31794 | 0.326413 | 0.002802 | 0.139742 | |
| 431 | . | gene | MSTRG.23528 | 4.207297 | 0.002814 | 0.140003 | |

| | | | | | | | |
|---|---|---|---|---|---|---|---|
| 436 | . | gene | MSTRG.25212 | 3.579578 | 0.002852 | 0.140293 | Bo7g098500.1 |
| 435 | . | gene | MSTRG.25394 | 0.742542 | 0.002852 | 0.140293 | Bo7g104410.1 |
| 433 | . | gene | MSTRG.26889 | 0.319366 | 0.002844 | 0.140293 | Bo8g034290.1 |
| 434 | . | gene | MSTRG.32671 | 0.376435 | 0.002844 | 0.140293 | Bo9g150120.1 |
| 432 | . | gene | MSTRG.27867 | 0.25574 | 0.002836 | 0.140293 | |
| 437 | . | gene | MSTRG.34107 | 0.434125 | 0.002865 | 0.140324 | |
| 438 | . | gene | MSTRG.13271 | 0.422271 | 0.002869 | 0.140324 | |
| 439 | . | gene | MSTRG.33164 | 0.185745 | 0.002872 | 0.140324 | |
| 442 | . | gene | MSTRG.15569 | 0.262249 | 0.002895 | 0.14049 | Bo4g186880.1 |
| 441 | . | gene | MSTRG.17471 | 1.727276 | 0.002894 | 0.14049 | Bo5g041440.1 |
| 440 | . | gene | MSTRG.9021 | 0.668166 | 0.002892 | 0.14049 | |
| 443 | . | gene | MSTRG.31086 | 0.679698 | 0.002906 | 0.140492 | Bo9g054630.1 |
| 444 | . | gene | MSTRG.16948 | 0.421184 | 0.002909 | 0.140492 | |
| 445 | . | gene | MSTRG.6009 | 0.572631 | 0.002928 | 0.141128 | |
| 446 | . | gene | MSTRG.2382 | 0.84168 | 0.002936 | 0.141174 | |
| 447 | PSAN | gene | MSTRG.10499 | 0.796305 | 0.002944 | 0.14125 | Bo3g102750.1 |
| 448 | . | gene | MSTRG.24757 | 0.668128 | 0.002966 | 0.142008 | |
| 451 | . | gene | MSTRG.4512 | 0.741301 | 0.002987 | 0.142051 | Bo2g050690.1 |
| 449 | . | gene | MSTRG.14798 | 4.881541 | 0.002976 | 0.142051 | Bo4g145950.1 |
| 450 | . | gene | MSTRG.35467 | 0.419972 | 0.002985 | 0.142051 | |
| 452 | . | gene | MSTRG.34995 | 20.15026 | 0.003004 | 0.142528 | |
| 453 | . | gene | MSTRG.7071 | 1.5086 | 0.003019 | 0.142796 | Bo3g006790.1 |
| 454 | . | gene | MSTRG.29953 | 0.459578 | 0.003023 | 0.142796 | Bo9g007960.1 |
| 455 | . | gene | MSTRG.7027 | 0.69091 | 0.003044 | 0.143344 | Bo3g005300.1 |
| 456 | . | gene | MSTRG.8016 | 6.931457 | 0.003048 | 0.143344 | Bo3g024840.1 |
| 475 | . | gene | MSTRG.432 | 0.318378 | 0.003199 | 0.143739 | Bo1g008770.1 |
| 460 | . | gene | MSTRG.6747 | 2.20238 | 0.003109 | 0.143739 | Bo2g168300.1 |
| 464 | . | gene | MSTRG.7224 | 0.266558 | 0.003135 | 0.143739 | Bo3g009810.1 |
| 480 | . | gene | MSTRG.7811 | 0.047417 | 0.003227 | 0.143739 | Bo3g021390.1 |

| | | | | | | | |
|---|---|---|---|---|---|---|---|
| 473 | . | gene | MSTRG.10060 | 1.826629 | 0.00319 | 0.143739 | Bo3g083960.1 |
| 471 | . | gene | MSTRG.11500 | 0.599467 | 0.003182 | 0.143739 | Bo3g155500.1 |
| 457 | . | gene | MSTRG.15074 | 0.301719 | 0.003072 | 0.143739 | Bo4g162530.1 |
| 483 | . | gene | MSTRG.15825 | 1.518816 | 0.003242 | 0.143739 | Bo4g191750.1 |
| 484 | . | gene | MSTRG.17505 | 4.270009 | 0.003246 | 0.143739 | Bo5g042910.1 |
| 467 | ATSYP24 | gene | MSTRG.17625 | 1.30026 | 0.003159 | 0.143739 | Bo5g048930.1 |
| 474 | . | gene | MSTRG.18538 | 0.428601 | 0.003197 | 0.143739 | Bo5g111210.1 |
| 485 | . | gene | MSTRG.20083 | 1.430388 | 0.003252 | 0.143739 | Bo6g022440.1 |
| 469 | . | gene | MSTRG.23239 | 0.494864 | 0.003177 | 0.143739 | Bo7g029050.1 |
| 479 | . | gene | MSTRG.23598 | 3.676068 | 0.003224 | 0.143739 | Bo7g046900.1 |
| 466 | . | gene | MSTRG.23975 | 1.649709 | 0.003145 | 0.143739 | Bo7g062300.1 |
| 461 | . | gene | MSTRG.26006 | 1.242071 | 0.003115 | 0.143739 | Bo7g115530.1 |
| 459 | . | gene | MSTRG.27381 | 0.269726 | 0.003086 | 0.143739 | Bo8g059890.1 |
| 481 | . | gene | MSTRG.27958 | 1.310346 | 0.003235 | 0.143739 | Bo8g079150.1 |
| 482 | . | gene | MSTRG.28107 | 0.676291 | 0.003241 | 0.143739 | Bo8g081770.1 |
| 486 | . | gene | MSTRG.29282 | 1.545774 | 0.003257 | 0.143739 | Bo8g107730.1 |
| 458 | . | gene | MSTRG.30267 | 0.403246 | 0.003074 | 0.143739 | Bo9g015080.1 |
| 462 | . | gene | MSTRG.20903 | 0.512847 | 0.003119 | 0.143739 | |
| 463 | . | gene | MSTRG.34338 | 0.320179 | 0.003121 | 0.143739 | |
| 465 | . | gene | MSTRG.31838 | 0.343774 | 0.003141 | 0.143739 | |
| 468 | . | gene | MSTRG.35164 | 0.493728 | 0.003162 | 0.143739 | |
| 470 | . | gene | MSTRG.35406 | 10.51896 | 0.003178 | 0.143739 | |
| 472 | . | gene | MSTRG.30681 | 1.411144 | 0.003183 | 0.143739 | |
| 476 | . | gene | MSTRG.20436 | 2.292821 | 0.003202 | 0.143739 | |
| 477 | . | gene | MSTRG.30998 | 0.399072 | 0.003212 | 0.143739 | |
| 478 | . | gene | MSTRG.17336 | 1.594025 | 0.003219 | 0.143739 | |
| 487 | . | gene | MSTRG.20892 | 2.164242 | 0.00328 | 0.144441 | Bo6g068170.1 |
| 488 | . | gene | MSTRG.22429 | 0.62399 | 0.003299 | 0.14499 | |
| 489 | . | gene | MSTRG.13499 | 0.439075 | 0.003325 | 0.145823 | Bo4g075430.1 |

| | | | | | | | |
|---|---|---|---|---|---|---|---|
| 499 | . | gene | MSTRG.4382 | 0.287428 | 0.003406 | 0.146144 | Bo2g043350.1 |
| 491 | . | gene | MSTRG.11381 | 1.34575 | 0.003347 | 0.146144 | Bo3g151300.1 |
| 497 | . | gene | MSTRG.16528 | 1.854578 | 0.003402 | 0.146144 | Bo5g010120.1 |
| 494 | . | gene | MSTRG.17236 | 0.156154 | 0.003381 | 0.146144 | Bo5g030880.1 |
| 500 | . | gene | MSTRG.17637 | 0.255457 | 0.003407 | 0.146144 | Bo5g049090.1 |
| 498 | . | gene | MSTRG.18308 | 7.587442 | 0.003403 | 0.146144 | Bo5g094570.1 |
| 495 | . | gene | MSTRG.18429 | 0.512216 | 0.003384 | 0.146144 | Bo5g102350.1 |
| 490 | . | gene | MSTRG.28361 | 0.673761 | 0.003347 | 0.146144 | Bo8g090470.1 |
| 493 | . | gene | MSTRG.28976 | 0.470556 | 0.003364 | 0.146144 | Bo8g102000.1 |
| 492 | . | gene | MSTRG.32732 | 0.276035 | 0.003357 | 0.146144 | Bo9g152200.1 |
| 496 | . | gene | MSTRG.18294 | 0.48752 | 0.003386 | 0.146144 | |
| 501 | . | gene | MSTRG.30726 | 0.483737 | 0.003417 | 0.146279 | Bo9g029230.1 |
| 502 | . | gene | MSTRG.11346 | 1.84661 | 0.003426 | 0.146287 | Bo3g149730.1 |
| 503 | . | gene | MSTRG.20712 | 0.089658 | 0.003431 | 0.146287 | Bo6g064380.1 |
| 504 | . | gene | MSTRG.3731 | 0.313828 | 0.003439 | 0.146353 | |
| 505 | . | gene | MSTRG.8455 | 1.900719 | 0.003451 | 0.146361 | Bo3g035120.1 |
| 507 | . | gene | MSTRG.17091 | 0.53673 | 0.00346 | 0.146361 | Bo5g025790.1 |
| 506 | . | gene | MSTRG.10073 | 2.221174 | 0.003454 | 0.146361 | |
| 508 | . | gene | MSTRG.6642 | 2.371538 | 0.003476 | 0.146759 | Bo2g165700.1 |
| 509 | . | gene | MSTRG.9280 | 2.514211 | 0.003483 | 0.146759 | Bo3g058990.1 |
| 510 | . | gene | MSTRG.11801 | 0.468553 | 0.00351 | 0.147613 | Bo3g171180.1 |
| 511 | . | gene | MSTRG.15737 | 0.444985 | 0.003521 | 0.147791 | Bo4g190420.1 |
| 512 | . | gene | MSTRG.2282 | 0.522324 | 0.003528 | 0.147794 | |
| 513 | . | gene | MSTRG.7416 | 0.727766 | 0.003567 | 0.148306 | Bo3g013260.1 |
| 514 | . | gene | MSTRG.12081 | 1.569955 | 0.003574 | 0.148306 | Bo3g184520.1 |
| 515 | . | gene | MSTRG.25249 | 0.17015 | 0.003574 | 0.148306 | Bo7g099130.1 |
| 516 | . | gene | MSTRG.7338 | 2.906866 | 0.003574 | 0.148306 | |
| 517 | . | gene | MSTRG.11466 | 0.302206 | 0.003575 | 0.148306 | |
| 518 | . | gene | MSTRG.15858 | 5.4208 | 0.003596 | 0.148887 | Bo4g194290.1 |

| | | | | | | | |
|---|---|---|---|---|---|---|---|
| 519 | . | gene | MSTRG.23102 | 0.240098 | 0.003605 | 0.148962 | Bo7g018610.1 |
| 520 | APS3 | gene | MSTRG.1740 | 4.142095 | 0.003646 | 0.150172 | Bo1g057690.1 |
| 521 | . | gene | MSTRG.30626 | 0.56373 | 0.003648 | 0.150172 | |
| 522 | . | gene | MSTRG.7957 | 1.634263 | 0.003706 | 0.151996 | Bo3g023930.1 |
| 523 | . | gene | MSTRG.8304 | 0.813257 | 0.003707 | 0.151996 | |
| 524 | . | gene | MSTRG.19338 | 1.565598 | 0.003718 | 0.152156 | Bo5g138000.1 |
| 525 | . | gene | MSTRG.33926 | 0.086483 | 0.003729 | 0.152355 | |
| 526 | . | gene | MSTRG.20917 | 0.295197 | 0.00374 | 0.152479 | Bo6g068520.1 |
| 529 | . | gene | MSTRG.5271 | 0.304848 | 0.003774 | 0.152843 | Bo2g095520.1 |
| 528 | . | gene | MSTRG.17778 | 2.392074 | 0.003772 | 0.152843 | Bo5g058510.1 |
| 527 | . | gene | MSTRG.27690 | 2.505318 | 0.003757 | 0.152843 | Bo8g069880.1 |
| 530 | . | gene | MSTRG.14433 | 0.368924 | 0.003777 | 0.152843 | |
| 531 | . | gene | MSTRG.27838 | 0.317222 | 0.003792 | 0.153158 | |
| 537 | . | gene | MSTRG.6958 | 1.808605 | 0.003842 | 0.153441 | Bo3g004300.1 |
| 534 | . | gene | MSTRG.7206 | 0.476833 | 0.003828 | 0.153441 | Bo3g009640.1 |
| 535 | . | gene | MSTRG.15872 | 0.737865 | 0.003841 | 0.153441 | Bo4g194540.1 |
| 536 | . | gene | MSTRG.23801 | 0.75832 | 0.003842 | 0.153441 | Bo7g055880.1 |
| 532 | . | gene | MSTRG.31963 | 0.787341 | 0.003807 | 0.153441 | Bo9g106020.1 |
| 533 | . | gene | MSTRG.1793 | 0.290894 | 0.003824 | 0.153441 | |
| 538 | . | gene | MSTRG.8773 | 2.259802 | 0.003864 | 0.154032 | Bo3g042010.1 |
| 540 | . | gene | MSTRG.19675 | 1.219033 | 0.003882 | 0.154196 | Bo5g148410.1 |
| 539 | . | gene | MSTRG.36106 | 0.394517 | 0.003881 | 0.154196 | |
| 541 | . | gene | MSTRG.36069 | 3.043909 | 0.003892 | 0.154289 | |
| 542 | . | gene | MSTRG.17113 | 0.528892 | 0.003905 | 0.154489 | Bo5g027200.1 |
| 543 | . | gene | MSTRG.20749 | 0.470973 | 0.003911 | 0.154489 | |
| 544 | . | gene | MSTRG.31139 | 1.342419 | 0.003919 | 0.154504 | Bo9g058360.1 |
| 548 | . | gene | MSTRG.12271 | 1.328219 | 0.003971 | 0.155297 | Bo4g009280.1 |
| 545 | . | gene | MSTRG.34045 | 9.408057 | 0.003952 | 0.155297 | |
| 546 | . | gene | MSTRG.17106 | 0.468882 | 0.003957 | 0.155297 | |

| | | | | | | | |
|---|---|---|---|---|---|---|---|
| 547 | . | gene | MSTRG.6551 | 0.296215 | 0.003964 | 0.155297 | |
| 549 | . | gene | MSTRG.4254 | 0.556428 | 0.003975 | 0.155297 | |
| 550 | . | gene | MSTRG.10264 | 0.320187 | 0.004014 | 0.156537 | |
| 556 | . | gene | MSTRG.1036 | 5.892514 | 0.004076 | 0.156972 | Bo1g025830.1 |
| 552 | . | gene | MSTRG.6224 | 0.455194 | 0.004051 | 0.156972 | Bo2g148040.1 |
| 557 | . | gene | MSTRG.6665 | 2.342992 | 0.004077 | 0.156972 | Bo2g166010.1 |
| 553 | . | gene | MSTRG.8154 | 0.293747 | 0.004054 | 0.156972 | Bo3g027930.1 |
| 551 | . | gene | MSTRG.8908 | 0.13826 | 0.004042 | 0.156972 | Bo3g045130.1 |
| 555 | . | gene | MSTRG.14536 | 0.407386 | 0.004065 | 0.156972 | Bo4g137290.1 |
| 554 | . | gene | MSTRG.16919 | 0.723838 | 0.004063 | 0.156972 | |
| 558 | . | gene | MSTRG.6639 | 0.130121 | 0.004104 | 0.157565 | Bo2g165610.1 |
| 560 | . | gene | MSTRG.11962 | 1.674383 | 0.004126 | 0.157565 | Bo3g179550.1 |
| 564 | . | gene | MSTRG.12910 | 0.259435 | 0.004148 | 0.157565 | Bo4g038480.1 |
| 565 | HSP98.7 | gene | MSTRG.28806 | 1.941678 | 0.004151 | 0.157565 | Bo8g099360.1 |
| 563 | . | gene | MSTRG.31637 | 0.409959 | 0.004146 | 0.157565 | Bo9g086970.1 |
| 559 | . | gene | MSTRG.19408 | 0.830428 | 0.004125 | 0.157565 | |
| 561 | . | gene | MSTRG.23230 | 0.789667 | 0.004135 | 0.157565 | |
| 562 | . | gene | MSTRG.17010 | 0.547605 | 0.004142 | 0.157565 | |
| 566 | . | gene | MSTRG.17761 | 0.496283 | 0.004176 | 0.158225 | |
| 568 | . | gene | MSTRG.10539 | 5.188544 | 0.0042 | 0.158605 | Bo3g103270.1 |
| 567 | . | gene | MSTRG.33584 | 0.134209 | 0.004199 | 0.158605 | Bo9g176490.1 |
| 569 | . | gene | MSTRG.2947 | 0.201737 | 0.004208 | 0.158625 | Bo1g141240.1 |
| 570 | . | gene | MSTRG.7829 | 0.649483 | 0.004236 | 0.158631 | Bo3g021690.1 |
| 572 | . | gene | MSTRG.25321 | 1.63314 | 0.004238 | 0.158631 | Bo7g101400.1 |
| 571 | . | gene | MSTRG.15136 | 0.277259 | 0.004236 | 0.158631 | |
| 573 | . | gene | MSTRG.12666 | 1.525653 | 0.004238 | 0.158631 | |
| 606 | . | gene | MSTRG.762 | 1.451049 | 0.004498 | 0.158785 | Bo1g017600.1 |
| 585 | . | gene | MSTRG.1673 | 0.653821 | 0.00434 | 0.158785 | Bo1g056650.1 |
| 592 | . | gene | MSTRG.1862 | 0.714258 | 0.004407 | 0.158785 | Bo1g070980.1 |

| | | | | | | | |
|---|---|---|---|---|---|---|---|
| 578 | . | gene | MSTRG.2899 | 0.82138 | 0.0043 | 0.158785 | Bo1g139580.1 |
| 588 | . | gene | MSTRG.4321 | 26.79924 | 0.004364 | 0.158785 | Bo2g041340.1 |
| 598 | . | gene | MSTRG.7072 | 1.915838 | 0.004448 | 0.158785 | Bo3g006800.1 |
| 599 | . | gene | MSTRG.11969 | 1.522519 | 0.00445 | 0.158785 | Bo3g179630.1 |
| 582 | . | gene | MSTRG.12080 | 0.140469 | 0.004323 | 0.158785 | Bo3g184510.1 |
| 595 | . | gene | MSTRG.14871 | 0.340072 | 0.004434 | 0.158785 | Bo4g151150.1 |
| 601 | . | gene | MSTRG.15308 | 5.801129 | 0.004475 | 0.158785 | Bo4g177560.1 |
| 584 | . | gene | MSTRG.17697 | 1.353128 | 0.004337 | 0.158785 | Bo5g052010.1 |
| 576 | . | gene | MSTRG.18072 | 0.122611 | 0.004288 | 0.158785 | Bo5g076490.1 |
| 589 | . | gene | MSTRG.18939 | 1.93003 | 0.004371 | 0.158785 | Bo5g127230.1 |
| 579 | . | gene | MSTRG.19216 | 0.722027 | 0.004304 | 0.158785 | Bo5g136280.1 |
| 574 | . | gene | MSTRG.20516 | 0.613128 | 0.004275 | 0.158785 | Bo6g049430.1 |
| 577 | . | gene | MSTRG.21862 | 2.320969 | 0.004292 | 0.158785 | Bo6g101120.1 |
| 590 | . | gene | MSTRG.24516 | 0.287751 | 0.004399 | 0.158785 | Bo7g081370.1 |
| 605 | . | gene | MSTRG.26840 | 1.188626 | 0.004497 | 0.158785 | Bo8g032290.1 |
| 593 | . | gene | MSTRG.27695 | 0.491368 | 0.004418 | 0.158785 | Bo8g069940.1 |
| 581 | . | gene | MSTRG.28164 | 1.747266 | 0.004307 | 0.158785 | Bo8g082730.1 |
| 600 | . | gene | MSTRG.29636 | 1.671698 | 0.004458 | 0.158785 | Bo8g115880.1 |
| 603 | . | gene | MSTRG.29758 | 1.742929 | 0.004492 | 0.158785 | Bo9g002430.1 |
| 580 | . | gene | MSTRG.30143 | 0.30968 | 0.004305 | 0.158785 | Bo9g011190.1 |
| 597 | . | gene | MSTRG.31657 | 0.145847 | 0.00444 | 0.158785 | Bo9g088210.1 |
| 591 | . | gene | MSTRG.32076 | 0.152535 | 0.0044 | 0.158785 | Bo9g112650.1 |
| 575 | . | gene | MSTRG.30801 | 0.339664 | 0.004276 | 0.158785 | |
| 583 | . | gene | MSTRG.4146 | 0.57783 | 0.004326 | 0.158785 | |
| 586 | . | gene | MSTRG.34203 | 0.447828 | 0.004346 | 0.158785 | |
| 587 | . | gene | MSTRG.13480 | 0.56672 | 0.004361 | 0.158785 | |
| 594 | . | gene | MSTRG.27542 | 0.311326 | 0.004422 | 0.158785 | |
| 596 | . | gene | MSTRG.36150 | 0.771183 | 0.004438 | 0.158785 | |
| 602 | . | gene | MSTRG.14877 | 4.115233 | 0.00449 | 0.158785 | |

| | | | | | | | |
|---|---|---|---|---|---|---|---|
| 604 | . | gene | MSTRG.11448 | 0.568854 | 0.004494 | 0.158785 | |
| 607 | . | gene | MSTRG.20890 | 1.348314 | 0.004504 | 0.158785 | |
| 608 | . | gene | MSTRG.19984 | 0.227433 | 0.004505 | 0.158785 | |
| 609 | . | gene | MSTRG.22324 | 10.55283 | 0.004515 | 0.158785 | |
| 610 | . | gene | MSTRG.13465 | 0.813343 | 0.004516 | 0.158785 | |
| 613 | . | gene | MSTRG.29327 | 2.040332 | 0.00454 | 0.158844 | Bo8g108600.1 |
| 611 | . | gene | MSTRG.31018 | 0.752332 | 0.00453 | 0.158844 | Bo9g047660.1 |
| 612 | . | gene | MSTRG.32579 | 2.281781 | 0.004537 | 0.158844 | |
| 617 | . | gene | MSTRG.6442 | 24.57875 | 0.004618 | 0.159466 | Bo2g158420.1 |
| 619 | . | gene | MSTRG.8395 | 0.462514 | 0.004627 | 0.159466 | Bo3g034270.1 |
| 616 | . | gene | MSTRG.9610 | 0.299691 | 0.004602 | 0.159466 | Bo3g065900.1 |
| 623 | . | gene | MSTRG.18644 | 0.277444 | 0.004636 | 0.159466 | Bo5g115590.1 |
| 620 | . | gene | MSTRG.24329 | 0.693111 | 0.004627 | 0.159466 | Bo7g077420.1 |
| 614 | . | gene | MSTRG.34642 | 1.409732 | 0.00457 | 0.159466 | |
| 615 | . | gene | MSTRG.26943 | 0.48973 | 0.004586 | 0.159466 | |
| 618 | . | gene | MSTRG.20506 | 0.523085 | 0.004621 | 0.159466 | |
| 621 | . | gene | MSTRG.30918 | 0.335217 | 0.00463 | 0.159466 | |
| 622 | . | gene | MSTRG.14260 | 0.216273 | 0.00463 | 0.159466 | |
| 624 | . | gene | MSTRG.5266 | 0.782547 | 0.004642 | 0.159466 | |
| 625 | . | gene | MSTRG.31996 | 0.57511 | 0.004653 | 0.159466 | |
| 626 | . | gene | MSTRG.32078 | 0.489699 | 0.004655 | 0.159466 | |
| 629 | . | gene | MSTRG.2122 | 1.224262 | 0.004677 | 0.159477 | Bo1g091740.1 |
| 627 | . | gene | MSTRG.32706 | 1.581058 | 0.004668 | 0.159477 | Bo9g151840.1 |
| 628 | . | gene | MSTRG.26820 | 0.621054 | 0.004674 | 0.159477 | |
| 630 | . | gene | MSTRG.18438 | 0.565809 | 0.004686 | 0.159531 | Bo5g103450.1 |
| 633 | . | gene | MSTRG.15486 | 7.278356 | 0.004719 | 0.159696 | Bo4g184480.1 |
| 631 | PTAC2 | gene | MSTRG.22404 | 0.1993 | 0.004715 | 0.159696 | Bo6g119020.1 |
| 632 | . | gene | MSTRG.29287 | 0.808465 | 0.004717 | 0.159696 | Bo8g107870.1 |
| 634 | . | gene | MSTRG.29316 | 0.197907 | 0.004721 | 0.159696 | Bo8g108360.1 |

| | | | | | | | |
|---|---|---|---|---|---|---|---|
| 636 | . | gene | MSTRG.21172 | 1.291463 | 0.004764 | 0.160663 | Bo6g077340.1 |
| 635 | . | gene | MSTRG.33980 | 0.407345 | 0.004761 | 0.160663 | |
| 649 | . | gene | MSTRG.3304 | 0.394644 | 0.004897 | 0.160775 | Bo1g157620.1 |
| 653 | . | gene | MSTRG.6741 | 1.556956 | 0.004923 | 0.160775 | Bo2g168130.1 |
| 640 | . | gene | MSTRG.7178 | 0.618838 | 0.004834 | 0.160775 | Bo3g009320.1 |
| 646 | . | gene | MSTRG.10280 | 0.29826 | 0.004883 | 0.160775 | Bo3g093700.1 |
| 647 | . | gene | MSTRG.15721 | 0.389447 | 0.00489 | 0.160775 | Bo4g190200.1 |
| 660 | . | gene | MSTRG.15761 | 0.445527 | 0.004954 | 0.160775 | Bo4g190720.1 |
| 651 | . | gene | MSTRG.17141 | 1.962328 | 0.004908 | 0.160775 | Bo5g027630.1 |
| 658 | . | gene | MSTRG.17700 | 2.116687 | 0.004946 | 0.160775 | Bo5g052080.1 |
| 659 | . | gene | MSTRG.18193 | 0.298055 | 0.004951 | 0.160775 | Bo5g086670.1 |
| 656 | . | gene | MSTRG.20618 | 0.679102 | 0.004944 | 0.160775 | Bo6g056140.1 |
| 648 | . | gene | MSTRG.21010 | 2.051753 | 0.004894 | 0.160775 | Bo6g073060.1 |
| 642 | . | gene | MSTRG.21830 | 2.559475 | 0.00486 | 0.160775 | Bo6g099600.1 |
| 645 | . | gene | MSTRG.23943 | 0.634797 | 0.004878 | 0.160775 | Bo7g061690.1 |
| 638 | . | gene | MSTRG.24298 | 0.400331 | 0.004799 | 0.160775 | Bo7g075860.1 |
| 657 | . | gene | MSTRG.28850 | 0.270407 | 0.004945 | 0.160775 | Bo8g100100.1 |
| 650 | . | gene | MSTRG.29635 | 1.421183 | 0.004905 | 0.160775 | Bo8g115830.1 |
| 641 | . | gene | MSTRG.30921 | 2.657923 | 0.004857 | 0.160775 | Bo9g041000.1 |
| 637 | . | gene | MSTRG.33194 | 1.768971 | 0.004784 | 0.160775 | Bo9g168890.1 |
| 639 | . | gene | MSTRG.19303 | 0.270115 | 0.004812 | 0.160775 | |
| 643 | . | gene | MSTRG.23256 | 0.180268 | 0.004865 | 0.160775 | |
| 644 | . | gene | MSTRG.31814 | 0.779937 | 0.004875 | 0.160775 | |
| 652 | . | gene | MSTRG.31809 | 0.820663 | 0.004909 | 0.160775 | |
| 654 | . | gene | MSTRG.31652 | 0.289459 | 0.004924 | 0.160775 | |
| 655 | . | gene | MSTRG.34713 | 3.634553 | 0.00493 | 0.160775 | |
| 661 | . | gene | MSTRG.35579 | 0.123022 | 0.004965 | 0.160775 | |
| 662 | . | gene | MSTRG.10118 | 0.88545 | 0.004968 | 0.160775 | |
| 663 | . | gene | MSTRG.34992 | 2.047122 | 0.00497 | 0.160775 | |

| | | | | | | | |
|---|---|---|---|---|---|---|---|
| 664 | . | gene | MSTRG.8228 | 0.466012 | 0.004978 | 0.160792 | Bo3g029710.1 |
| 665 | . | gene | MSTRG.25393 | 2.048601 | 0.004986 | 0.160792 | |
| 666 | . | gene | MSTRG.14582 | 0.902503 | 0.005006 | 0.161103 | |
| 667 | . | gene | MSTRG.8632 | 0.732537 | 0.00501 | 0.161103 | |
| 668 | . | gene | MSTRG.2221 | 0.301534 | 0.005025 | 0.161204 | Bo1g098730.1 |
| 670 | . | gene | MSTRG.18647 | 0.582085 | 0.005039 | 0.161204 | Bo5g115620.1 |
| 669 | . | gene | MSTRG.21007 | 1.755707 | 0.00503 | 0.161204 | Bo6g072990.1 |
| 671 | . | gene | MSTRG.30760 | 0.456775 | 0.005044 | 0.161204 | Bo9g030780.1 |
| 673 | . | gene | MSTRG.6229 | 0.196612 | 0.005064 | 0.16138 | Bo2g149410.1 |
| 672 | . | gene | MSTRG.24167 | 0.363817 | 0.005057 | 0.16138 | Bo7g067640.1 |
| 674 | . | gene | MSTRG.3234 | 2.220619 | 0.005073 | 0.161386 | Bo1g153700.1 |
| 676 | . | gene | MSTRG.6760 | 2.155715 | 0.00509 | 0.161386 | Bo3g001130.1 |
| 677 | . | gene | MSTRG.14882 | 0.504998 | 0.005094 | 0.161386 | Bo4g151340.1 |
| 675 | . | gene | MSTRG.18108 | 0.945312 | 0.00508 | 0.161386 | |
| 678 | . | gene | MSTRG.23529 | 1.557178 | 0.005109 | 0.161621 | Bo7g044850.1 |
| 679 | . | gene | MSTRG.30571 | 1.690536 | 0.005125 | 0.161891 | Bo9g024490.1 |
| 680 | . | gene | MSTRG.20437 | 21.58562 | 0.005136 | 0.161994 | |
| 681 | . | gene | MSTRG.17964 | 1.684853 | 0.00516 | 0.162494 | Bo5g070770.1 |
| 683 | . | gene | MSTRG.8111 | 2.068476 | 0.005177 | 0.162521 | Bo3g027380.1 |
| 685 | . | gene | MSTRG.31499 | 0.571153 | 0.005191 | 0.162521 | Bo9g076420.1 |
| 682 | . | gene | MSTRG.20294 | 0.978893 | 0.005174 | 0.162521 | |
| 684 | . | gene | MSTRG.34159 | 2.218407 | 0.005184 | 0.162521 | |
| 687 | . | gene | MSTRG.2674 | 1.583607 | 0.005222 | 0.162791 | Bo1g123640.1 |
| 688 | . | gene | MSTRG.27357 | 0.457497 | 0.005222 | 0.162791 | Bo8g059470.1 |
| 686 | . | gene | MSTRG.14842 | 0.724719 | 0.005213 | 0.162791 | |
| 689 | . | gene | MSTRG.15592 | 0.952297 | 0.005256 | 0.163587 | Bo4g187300.1 |
| 690 | . | gene | MSTRG.31752 | 1.317492 | 0.005263 | 0.163587 | Bo9g093850.1 |
| 698 | . | gene | MSTRG.13715 | 0.361867 | 0.005327 | 0.163674 | Bo4g094190.1 |
| 693 | . | gene | MSTRG.18117 | 0.43206 | 0.00531 | 0.163674 | Bo5g080050.1 |

| | | | | | | | |
|---|---|---|---|---|---|---|---|
| 697 | . | gene | MSTRG.18298 | 2.47008 | 0.005326 | 0.163674 | Bo5g093400.1 |
| 691 | . | gene | MSTRG.21867 | 2.020858 | 0.005291 | 0.163674 | Bo6g101230.1 |
| 694 | . | gene | MSTRG.26508 | 0.332642 | 0.005317 | 0.163674 | Bo8g011470.1 |
| 692 | . | gene | MSTRG.34263 | 0.276363 | 0.005305 | 0.163674 | |
| 695 | . | gene | MSTRG.19979 | 0.079946 | 0.005322 | 0.163674 | |
| 696 | . | gene | MSTRG.18074 | 2.993207 | 0.005325 | 0.163674 | |
| 712 | . | gene | MSTRG.1181 | 0.940083 | 0.005447 | 0.163831 | Bo1g037050.1 |
| 713 | . | gene | MSTRG.8921 | 2.295837 | 0.005453 | 0.163831 | Bo3g045420.1 |
| 705 | . | gene | MSTRG.9582 | 0.399668 | 0.005408 | 0.163831 | Bo3g065510.1 |
| 700 | . | gene | MSTRG.9599 | 1.739169 | 0.005366 | 0.163831 | Bo3g065780.1 |
| 707 | . | gene | MSTRG.9825 | 19.21473 | 0.005431 | 0.163831 | Bo3g073430.1 |
| 704 | . | gene | MSTRG.16185 | 0.687916 | 0.005396 | 0.163831 | Bo5g004560.1 |
| 702 | . | gene | MSTRG.16588 | 0.607213 | 0.005378 | 0.163831 | Bo5g012000.1 |
| 710 | . | gene | MSTRG.17090 | 0.37473 | 0.005445 | 0.163831 | Bo5g025690.1 |
| 708 | . | gene | MSTRG.18427 | 0.484904 | 0.005439 | 0.163831 | Bo5g102290.1 |
| 706 | AAR3 | gene | MSTRG.24706 | 1.499638 | 0.005419 | 0.163831 | Bo7g087140.1 |
| 711 | . | gene | MSTRG.24869 | 1.293755 | 0.005446 | 0.163831 | Bo7g091480.1 |
| 699 | . | gene | MSTRG.35151 | 2.236511 | 0.005355 | 0.163831 | |
| 701 | . | gene | MSTRG.35221 | 4.507362 | 0.005367 | 0.163831 | |
| 703 | . | gene | MSTRG.15435 | 1.463002 | 0.005382 | 0.163831 | |
| 709 | . | gene | MSTRG.34504 | 1.600916 | 0.005439 | 0.163831 | |
| 714 | . | gene | MSTRG.34619 | 0.250639 | 0.005454 | 0.163831 | |
| 715 | . | gene | MSTRG.8769 | 1.661043 | 0.005463 | 0.163878 | Bo3g041960.1 |
| 716 | . | gene | MSTRG.2520 | 1.705597 | 0.00549 | 0.164198 | Bo1g115980.1 |
| 718 | . | gene | MSTRG.13957 | 0.472505 | 0.005502 | 0.164198 | Bo4g108140.1 |
| 717 | . | gene | MSTRG.14364 | 1.814679 | 0.005502 | 0.164198 | |
| 719 | . | gene | MSTRG.14749 | 0.531504 | 0.005505 | 0.164198 | |
| 722 | . | gene | MSTRG.3894 | 0.6954 | 0.005542 | 0.164619 | Bo2g018100.1 |
| 720 | . | gene | MSTRG.22304 | 1.210027 | 0.005527 | 0.164619 | Bo6g116530.1 |

| | | | | | | | |
|---|---|---|---|---|---|---|---|
| 721 | . | gene | MSTRG.29889 | 2.20919 | 0.005538 | 0.164619 | Bo9g005990.1 |
| 723 | . | gene | MSTRG.20394 | 0.668666 | 0.005551 | 0.164667 | Bo6g039740.1 |
| 727 | . | gene | MSTRG.1323 | 0.455948 | 0.005606 | 0.164824 | Bo1g040120.1 |
| 724 | . | gene | MSTRG.1380 | 1.727672 | 0.005573 | 0.164824 | Bo1g041900.1 |
| 736 | . | gene | MSTRG.3030 | 0.690815 | 0.005667 | 0.164824 | Bo1g144470.1 |
| 730 | . | gene | MSTRG.5728 | 0.505339 | 0.005632 | 0.164824 | Bo2g124550.1 |
| 737 | . | gene | MSTRG.16410 | 1.233565 | 0.005668 | 0.164824 | Bo5g008700.1 |
| 738 | . | gene | MSTRG.19382 | 0.260913 | 0.005672 | 0.164824 | Bo5g139620.1 |
| 729 | . | gene | MSTRG.20751 | 0.147953 | 0.005624 | 0.164824 | Bo6g065180.1 |
| 732 | . | gene | MSTRG.26858 | 1.541491 | 0.005644 | 0.164824 | Bo8g032490.1 |
| 726 | . | gene | MSTRG.28296 | 0.642968 | 0.005602 | 0.164824 | Bo8g089560.1 |
| 725 | . | gene | MSTRG.32804 | 0.549868 | 0.005602 | 0.164824 | Bo9g154350.1 |
| 734 | . | gene | MSTRG.33661 | 0.392301 | 0.005648 | 0.164824 | Bo9g177620.1 |
| 728 | . | gene | MSTRG.32497 | 0.468438 | 0.005611 | 0.164824 | |
| 731 | . | gene | MSTRG.21731 | 0.418929 | 0.005642 | 0.164824 | |
| 733 | . | gene | MSTRG.9075 | 1.453337 | 0.005645 | 0.164824 | |
| 735 | . | gene | MSTRG.11440 | 0.78282 | 0.005652 | 0.164824 | |
| 740 | . | gene | MSTRG.3433 | 2.116222 | 0.005702 | 0.165043 | Bo2g006700.1 |
| 739 | . | gene | MSTRG.27301 | 6.227849 | 0.005691 | 0.165043 | |
| 741 | . | gene | MSTRG.14928 | 0.712022 | 0.005702 | 0.165043 | |
| 742 | . | gene | MSTRG.12147 | 0.078921 | 0.005741 | 0.165903 | Bo3g185500.1 |
| 743 | . | gene | MSTRG.11837 | 0.457213 | 0.005747 | 0.165903 | |
| 744 | . | gene | MSTRG.14952 | 3.049863 | 0.005768 | 0.166266 | |
| 745 | . | gene | MSTRG.9488 | 0.399122 | 0.005785 | 0.16631 | Bo3g064320.1 |
| 746 | . | gene | MSTRG.30739 | 1.739888 | 0.005785 | 0.16631 | Bo9g029370.1 |
| 747 | . | gene | MSTRG.14964 | 2.216452 | 0.005802 | 0.166578 | Bo4g154610.1 |
| 748 | . | gene | MSTRG.10423 | 2.670936 | 0.005815 | 0.166727 | Bo3g100650.1 |
| 749 | . | gene | MSTRG.2530 | 0.697457 | 0.005844 | 0.167295 | Bo1g116190.1 |
| 750 | . | gene | MSTRG.33935 | 1.180326 | 0.00585 | 0.167295 | |

| | | | | | | | |
|---|---|---|---|---|---|---|---|
| 751 | . | gene | MSTRG.1832 | 0.199708 | 0.005869 | 0.167608 | Bo1g067400.1 |
| 752 | . | gene | MSTRG.25304 | 0.710409 | 0.005896 | 0.168159 | Bo7g101100.1 |
| 753 | TRH1 | gene | MSTRG.1178 | 4.319686 | 0.005919 | 0.168437 | Bo1g036910.1 |
| 754 | . | gene | MSTRG.34914 | 0.394869 | 0.005922 | 0.168437 | |
| 755 | . | gene | MSTRG.26606 | 0.125707 | 0.005932 | 0.168498 | Bo8g018100.1 |
| 760 | . | gene | MSTRG.9466 | 1.51501 | 0.005984 | 0.168743 | Bo3g064010.1 |
| 757 | ATSC35 | gene | MSTRG.10484 | 1.227836 | 0.005976 | 0.168743 | Bo3g101530.1 |
| 758 | . | gene | MSTRG.11841 | 0.186698 | 0.005979 | 0.168743 | Bo3g173750.1 |
| 756 | . | gene | MSTRG.15706 | 0.615957 | 0.005964 | 0.168743 | Bo4g190030.1 |
| 764 | . | gene | MSTRG.16524 | 4.633621 | 0.006017 | 0.168743 | Bo5g010070.1 |
| 761 | . | gene | MSTRG.22609 | 1.416672 | 0.006001 | 0.168743 | Bo6g121670.1 |
| 759 | . | gene | MSTRG.35149 | 0.124532 | 0.005982 | 0.168743 | |
| 762 | . | gene | MSTRG.9816 | 0.276231 | 0.006001 | 0.168743 | |
| 763 | . | gene | gene:Bo4g030120 | 0.949115 | 0.00601 | 0.168743 | |
| 765 | . | gene | MSTRG.24054 | 0.504125 | 0.006022 | 0.168743 | |
| 766 | . | gene | MSTRG.15985 | 0.652128 | 0.006027 | 0.168743 | |
| 771 | . | gene | MSTRG.120 | 0.334549 | 0.006098 | 0.168869 | Bo1g003720.1 |
| 794 | . | gene | MSTRG.2942 | 0.363303 | 0.00627 | 0.168869 | Bo1g141220.1 |
| 769 | . | gene | MSTRG.8280 | 1.432974 | 0.006075 | 0.168869 | Bo3g031550.1 |
| 793 | . | gene | MSTRG.8669 | 1.515633 | 0.006256 | 0.168869 | Bo3g039270.1 |
| 783 | FBR12 | gene | MSTRG.11045 | 1.533232 | 0.006209 | 0.168869 | Bo3g137090.1 |
| 777 | . | gene | MSTRG.13073 | 0.296311 | 0.006161 | 0.168869 | Bo4g043690.1 |
| 785 | . | gene | MSTRG.14079 | 0.201945 | 0.006223 | 0.168869 | Bo4g115540.1 |
| 791 | . | gene | MSTRG.15032 | 0.553891 | 0.006254 | 0.168869 | Bo4g159930.1 |
| 772 | . | gene | MSTRG.17097 | 0.836539 | 0.006099 | 0.168869 | Bo5g025910.1 |
| 788 | . | gene | MSTRG.19689 | 0.614073 | 0.006238 | 0.168869 | Bo5g148710.1 |
| 796 | . | gene | MSTRG.20748 | 4.060321 | 0.006273 | 0.168869 | Bo6g065090.1 |
| 797 | . | gene | MSTRG.20924 | 1.247732 | 0.006275 | 0.168869 | Bo6g069590.1 |
| 789 | . | gene | MSTRG.20987 | 8.51954 | 0.00625 | 0.168869 | Bo6g072660.1 |

| | | | | | | | |
|---|---|---|---|---|---|---|---|
| 786 | . | gene | MSTRG.22160 | 3.133607 | 0.006232 | 0.168869 | Bo6g112360.1 |
| 770 | . | gene | MSTRG.22505 | 2.632637 | 0.006092 | 0.168869 | Bo6g120340.1 |
| 776 | . | gene | MSTRG.25177 | 12.99973 | 0.00616 | 0.168869 | Bo7g098000.1 |
| 778 | . | gene | MSTRG.25263 | 0.323282 | 0.006174 | 0.168869 | Bo7g099470.1 |
| 779 | . | gene | MSTRG.27862 | 1.348436 | 0.006188 | 0.168869 | Bo8g076700.1 |
| 787 | . | gene | MSTRG.28089 | 0.803301 | 0.006233 | 0.168869 | Bo8g081500.1 |
| 784 | . | gene | MSTRG.30499 | 0.732014 | 0.006214 | 0.168869 | Bo9g022270.1 |
| 782 | . | gene | MSTRG.30872 | 0.275256 | 0.006196 | 0.168869 | Bo9g036990.1 |
| 774 | . | gene | MSTRG.32056 | 0.596747 | 0.006117 | 0.168869 | Bo9g111300.1 |
| 792 | . | gene | MSTRG.32655 | 0.628516 | 0.006255 | 0.168869 | Bo9g149910.1 |
| 790 | . | gene | MSTRG.33241 | 1.649667 | 0.006251 | 0.168869 | Bo9g169630.1 |
| 767 | . | gene | MSTRG.13131 | 0.746908 | 0.006067 | 0.168869 | |
| 768 | . | gene | MSTRG.22529 | 16.84475 | 0.006071 | 0.168869 | |
| 773 | . | gene | MSTRG.35689 | 0.52243 | 0.006114 | 0.168869 | |
| 775 | . | gene | MSTRG.17599 | 0.122658 | 0.006125 | 0.168869 | |
| 780 | . | gene | MSTRG.34695 | 3.273872 | 0.006191 | 0.168869 | |
| 781 | . | gene | MSTRG.35187 | 43.52291 | 0.006195 | 0.168869 | |
| 795 | . | gene | MSTRG.36127 | 0.61927 | 0.006271 | 0.168869 | |
| 801 | . | gene | MSTRG.16637 | 0.134489 | 0.006309 | 0.168938 | Bo5g014720.1 |
| 798 | . | gene | MSTRG.20027 | 4.733313 | 0.006302 | 0.168938 | |
| 799 | . | gene | MSTRG.10855 | 0.595042 | 0.006305 | 0.168938 | |
| 800 | . | gene | MSTRG.33892 | 0.101595 | 0.006306 | 0.168938 | |
| 802 | . | gene | MSTRG.17144 | 0.129066 | 0.006322 | 0.169063 | Bo5g027670.1 |
| 804 | . | gene | MSTRG.3804 | 1.535354 | 0.006344 | 0.169108 | Bo2g013780.1 |
| 803 | . | gene | MSTRG.5216 | 0.446499 | 0.006335 | 0.169108 | Bo2g093390.1 |
| 805 | . | gene | MSTRG.15426 | 0.616443 | 0.006347 | 0.169108 | Bo4g182480.1 |
| 807 | . | gene | MSTRG.11447 | 0.086486 | 0.006369 | 0.169265 | Bo3g153720.1 |
| 806 | . | gene | MSTRG.25626 | 0.767746 | 0.006369 | 0.169265 | Bo7g108630.1 |
| 808 | . | gene | MSTRG.35679 | 0.266839 | 0.006392 | 0.169558 | |

| | | | | | | | |
|---|---|---|---|---|---|---|---|
| 809 | . | gene | MSTRG.34853 | 1.723203 | 0.006396 | 0.169558 | |
| 810 | ALA1 | gene | MSTRG.6887 | 0.623357 | 0.006412 | 0.169637 | Bo3g003060.1 |
| 811 | . | gene | MSTRG.19143 | 1.825375 | 0.006415 | 0.169637 | Bo5g133140.1 |
| 814 | . | gene | MSTRG.4034 | 0.39177 | 0.006483 | 0.170218 | Bo2g024310.1 |
| 817 | . | gene | MSTRG.6345 | 0.343548 | 0.00649 | 0.170218 | Bo2g154020.1 |
| 819 | . | gene | MSTRG.8814 | 1.354386 | 0.0065 | 0.170218 | Bo3g042560.1 |
| 812 | . | gene | MSTRG.9318 | 2.042126 | 0.006456 | 0.170218 | Bo3g060510.1 |
| 816 | . | gene | MSTRG.13513 | 2.095789 | 0.006486 | 0.170218 | Bo4g076740.1 |
| 815 | . | gene | MSTRG.29486 | 1.415835 | 0.006483 | 0.170218 | Bo8g112600.1 |
| 813 | . | gene | MSTRG.33396 | 0.596153 | 0.006471 | 0.170218 | |
| 818 | . | gene | MSTRG.35135 | 0.60777 | 0.0065 | 0.170218 | |
| 821 | . | gene | MSTRG.18574 | 14.93419 | 0.006531 | 0.170422 | Bo5g113720.1 |
| 822 | . | gene | MSTRG.21803 | 10.94853 | 0.006532 | 0.170422 | Bo6g099140.1 |
| 820 | . | gene | MSTRG.34824 | 0.310938 | 0.006524 | 0.170422 | |
| 834 | . | gene | MSTRG.8218 | 0.894441 | 0.006665 | 0.171204 | Bo3g028560.1 |
| 824 | . | gene | MSTRG.15912 | 0.24858 | 0.006603 | 0.171204 | Bo4g195360.1 |
| 831 | FTSH1 | gene | MSTRG.21941 | 0.791259 | 0.006646 | 0.171204 | Bo6g105140.1 |
| 826 | . | gene | MSTRG.30763 | 0.103853 | 0.006618 | 0.171204 | Bo9g030880.1 |
| 835 | . | gene | MSTRG.31419 | 0.114129 | 0.006666 | 0.171204 | Bo9g072980.1 |
| 829 | . | gene | MSTRG.33180 | 1.461129 | 0.006626 | 0.171204 | Bo9g166760.1 |
| 823 | . | gene | MSTRG.35491 | 0.205629 | 0.006602 | 0.171204 | |
| 825 | . | gene | MSTRG.35337 | 0.145609 | 0.006613 | 0.171204 | |
| 827 | . | gene | MSTRG.13736 | 0.493512 | 0.00662 | 0.171204 | |
| 828 | . | gene | MSTRG.31486 | 0.407245 | 0.006624 | 0.171204 | |
| 830 | . | gene | MSTRG.34362 | 0.286908 | 0.00664 | 0.171204 | |
| 832 | . | gene | MSTRG.36016 | 0.510719 | 0.006649 | 0.171204 | |
| 833 | . | gene | MSTRG.14586 | 0.274381 | 0.006655 | 0.171204 | |
| 838 | . | gene | MSTRG.27846 | 1.589459 | 0.006717 | 0.171743 | Bo8g076430.1 |
| 836 | . | gene | MSTRG.29139 | 1.689694 | 0.006695 | 0.171743 | Bo8g105530.1 |

| | | | | | | | |
|---|---|---|---|---|---|---|---|
| 837 | . | gene | MSTRG.29957 | 0.442842 | 0.006716 | 0.171743 | Bo9g008010.1 |
| 839 | . | gene | MSTRG.34622 | 2.385272 | 0.006719 | 0.171743 | |
| 840 | . | gene | MSTRG.27832 | 2.278818 | 0.00674 | 0.172075 | |
| 842 | . | gene | MSTRG.6698 | 0.824738 | 0.006769 | 0.172419 | Bo2g166450.1 |
| 841 | . | gene | MSTRG.35623 | 0.35416 | 0.006768 | 0.172419 | |
| 843 | . | gene | MSTRG.24153 | 0.492046 | 0.00681 | 0.173244 | |
| 853 | . | gene | MSTRG.666 | 0.202719 | 0.006925 | 0.173289 | Bo1g016200.1 |
| 866 | . | gene | MSTRG.1173 | 0.152317 | 0.007004 | 0.173289 | Bo1g036870.1 |
| 856 | . | gene | MSTRG.3772 | 1.872222 | 0.006952 | 0.173289 | Bo2g013180.1 |
| 848 | . | gene | MSTRG.11571 | 2.612245 | 0.006881 | 0.173289 | Bo3g159900.1 |
| 867 | CSP41A | gene | MSTRG.13770 | 0.602876 | 0.007005 | 0.173289 | Bo4g098190.1 |
| 855 | . | gene | MSTRG.14072 | 0.273083 | 0.006943 | 0.173289 | Bo4g115460.1 |
| 850 | . | gene | MSTRG.14357 | 6.316531 | 0.006901 | 0.173289 | Bo4g129600.1 |
| 847 | . | gene | MSTRG.15335 | 1.535972 | 0.006853 | 0.173289 | Bo4g178100.1 |
| 861 | . | gene | MSTRG.15646 | 1.633337 | 0.006968 | 0.173289 | Bo4g188070.1 |
| 862 | . | gene | MSTRG.19881 | 0.413385 | 0.006971 | 0.173289 | Bo6g007000.1 |
| 864 | . | gene | MSTRG.21045 | 2.10489 | 0.006993 | 0.173289 | Bo6g074620.1 |
| 863 | . | gene | MSTRG.22345 | 0.615504 | 0.006978 | 0.173289 | Bo6g118150.1 |
| 852 | PFK4 | gene | MSTRG.25235 | 0.362606 | 0.006923 | 0.173289 | Bo7g098840.1 |
| 854 | . | gene | MSTRG.25431 | 6.257176 | 0.006928 | 0.173289 | Bo7g105090.1 |
| 858 | . | gene | MSTRG.25667 | 0.476453 | 0.006955 | 0.173289 | Bo7g109290.1 |
| 844 | . | gene | MSTRG.28969 | 1.294248 | 0.006825 | 0.173289 | Bo8g101880.1 |
| 857 | . | gene | MSTRG.29348 | 0.918819 | 0.006952 | 0.173289 | Bo8g108930.1 |
| 860 | . | gene | MSTRG.30068 | 1.333774 | 0.006967 | 0.173289 | Bo9g010060.1 |
| 865 | . | gene | MSTRG.30597 | 0.523277 | 0.006996 | 0.173289 | Bo9g026090.1 |
| 859 | . | gene | MSTRG.31630 | 0.138209 | 0.006957 | 0.173289 | Bo9g085850.1 |
| 845 | . | gene | MSTRG.15724 | 1.887151 | 0.006845 | 0.173289 | |
| 846 | . | gene | MSTRG.35133 | 0.22061 | 0.00685 | 0.173289 | |
| 849 | . | gene | MSTRG.34131 | 2.124889 | 0.006885 | 0.173289 | |

| | | | | | | | |
|---|---|---|---|---|---|---|---|
| 851 | . | gene | MSTRG.10029 | 0.633505 | 0.006917 | 0.173289 | |
| 869 | . | gene | MSTRG.6844 | 0.458936 | 0.007024 | 0.173324 | Bo3g002340.1 |
| 870 | . | gene | MSTRG.10996 | 0.739274 | 0.007031 | 0.173324 | Bo3g134560.1 |
| 868 | . | gene | MSTRG.21415 | 4.277479 | 0.007023 | 0.173324 | Bo6g083280.1 |
| 871 | . | gene | MSTRG.22945 | 0.780393 | 0.007052 | 0.17365 | Bo7g009650.1 |
| 872 | . | gene | MSTRG.30858 | 0.479166 | 0.007079 | 0.174098 | Bo9g036760.1 |
| 880 | . | gene | MSTRG.2373 | 0.653543 | 0.007154 | 0.174288 | Bo1g105520.1 |
| 877 | ERD2B | gene | MSTRG.10052 | 1.225152 | 0.00714 | 0.174288 | Bo3g083750.1 |
| 874 | . | gene | MSTRG.11416 | 3.001256 | 0.007104 | 0.174288 | Bo3g153120.1 |
| 876 | . | gene | MSTRG.12562 | 2.820982 | 0.007138 | 0.174288 | Bo4g024680.1 |
| 878 | . | gene | MSTRG.17095 | 2.489655 | 0.007142 | 0.174288 | Bo5g025860.1 |
| 873 | . | gene | MSTRG.19688 | 1.665315 | 0.007101 | 0.174288 | Bo5g148630.1 |
| 881 | . | gene | MSTRG.33337 | 0.187432 | 0.007159 | 0.174288 | Bo9g172040.1 |
| 875 | . | gene | MSTRG.34386 | 1.339656 | 0.007134 | 0.174288 | |
| 879 | . | gene | MSTRG.35031 | 0.487186 | 0.007145 | 0.174288 | |
| 882 | . | gene | MSTRG.10689 | 1.632813 | 0.007168 | 0.174294 | Bo3g109690.1 |
| 883 | . | gene | MSTRG.5119 | 11.70541 | 0.007184 | 0.174409 | Bo2g089970.1 |
| 884 | . | gene | MSTRG.25316 | 0.218538 | 0.007189 | 0.174409 | Bo7g101340.1 |
| 886 | . | gene | MSTRG.16912 | 1.396319 | 0.007205 | 0.174413 | Bo5g021150.1 |
| 885 | . | gene | MSTRG.16435 | 5.303561 | 0.007203 | 0.174413 | |
| 889 | . | gene | MSTRG.13059 | 0.693785 | 0.007232 | 0.174456 | Bo4g043500.1 |
| 893 | ENOC | gene | MSTRG.13365 | 1.516185 | 0.007264 | 0.174456 | Bo4g061400.1 |
| 892 | . | gene | MSTRG.18394 | 0.434816 | 0.007261 | 0.174456 | Bo5g098830.1 |
| 891 | . | gene | MSTRG.20539 | 1.409434 | 0.007259 | 0.174456 | Bo6g050710.1 |
| 888 | . | gene | MSTRG.27526 | 0.312834 | 0.007224 | 0.174456 | Bo8g066190.1 |
| 887 | . | gene | MSTRG.29447 | 2.784454 | 0.007217 | 0.174456 | Bo8g111020.1 |
| 890 | . | gene | MSTRG.23447 | 0.990763 | 0.007251 | 0.174456 | |
| 894 | . | gene | MSTRG.25832 | 0.750254 | 0.007305 | 0.175248 | Bo7g111820.1 |
| 895 | . | gene | MSTRG.23652 | 0.181624 | 0.007321 | 0.175411 | Bo7g049780.1 |

| | | | | | | | |
|---|---|---|---|---|---|---|---|
| 896 | . | gene | MSTRG.1884 | 0.351657 | 0.007328 | 0.175411 | |
| 898 | . | gene | MSTRG.6842 | 0.49751 | 0.007351 | 0.175557 | Bo3g002320.1 |
| 897 | . | gene | MSTRG.15115 | 0.434683 | 0.007348 | 0.175557 | |
| 899 | . | gene | MSTRG.19072 | 0.130404 | 0.007367 | 0.17574 | |
| 900 | . | gene | MSTRG.7160 | 0.589778 | 0.007399 | 0.176317 | |
| 901 | . | gene | MSTRG.4232 | 4.533015 | 0.007422 | 0.176371 | Bo2g032480.1 |
| 903 | . | gene | MSTRG.6398 | 1.684071 | 0.007432 | 0.176371 | Bo2g155680.1 |
| 902 | . | gene | MSTRG.27292 | 1.399037 | 0.00743 | 0.176371 | Bo8g058490.1 |
| 904 | . | gene | MSTRG.34678 | 0.768204 | 0.007434 | 0.176371 | |
| 906 | . | gene | MSTRG.1261 | 1.739225 | 0.007482 | 0.177039 | Bo1g039160.1 |
| 905 | . | gene | MSTRG.14866 | 0.126595 | 0.007474 | 0.177039 | |
| 907 | . | gene | MSTRG.19039 | 0.914089 | 0.007489 | 0.177039 | |
| 908 | . | gene | MSTRG.31890 | 4.81715 | 0.007495 | 0.177039 | |
| 909 | . | gene | MSTRG.7989 | 0.237429 | 0.007508 | 0.177122 | Bo3g024400.1 |
| 913 | . | gene | MSTRG.12222 | 2.700338 | 0.00755 | 0.177122 | Bo4g004630.1 |
| 910 | . | gene | MSTRG.14581 | 0.287202 | 0.007521 | 0.177122 | Bo4g137930.1 |
| 915 | . | gene | MSTRG.15968 | 0.374982 | 0.007559 | 0.177122 | Bo4g197170.1 |
| 911 | . | gene | MSTRG.24760 | 1.676654 | 0.007529 | 0.177122 | Bo7g088910.1 |
| 912 | . | gene | MSTRG.2946 | 0.127789 | 0.007542 | 0.177122 | |
| 914 | . | gene | MSTRG.34793 | 0.675622 | 0.007559 | 0.177122 | |
| 916 | . | gene | MSTRG.24409 | 0.822026 | 0.007565 | 0.177122 | |
| 917 | . | gene | MSTRG.27249 | 0.25879 | 0.007577 | 0.177213 | Bo8g056920.1 |
| 918 | CRK2 | gene | MSTRG.4848 | 1.880498 | 0.007586 | 0.177235 | Bo2g068260.1 |
| 919 | . | gene | MSTRG.11338 | 0.370388 | 0.007612 | 0.177297 | Bo3g149580.1 |
| 920 | . | gene | MSTRG.15121 | 0.591149 | 0.007618 | 0.177297 | Bo4g165260.1 |
| 921 | . | gene | MSTRG.27803 | 2.424851 | 0.007621 | 0.177297 | Bo8g071620.1 |
| 922 | . | gene | MSTRG.35424 | 0.106218 | 0.007622 | 0.177297 | |
| 923 | . | gene | MSTRG.23262 | 1.154948 | 0.007631 | 0.177327 | Bo7g030520.1 |
| 924 | . | gene | MSTRG.7368 | 0.61684 | 0.00765 | 0.17756 | Bo3g012380.1 |

| | | | | | | | |
|---|---|---|---|---|---|---|---|
| 926 | . | gene | MSTRG.27353 | 2.483773 | 0.007686 | 0.178007 | Bo8g059440.1 |
| 925 | . | gene | MSTRG.34738 | 1.731771 | 0.007682 | 0.178007 | |
| 928 | . | gene | MSTRG.376 | 1.646462 | 0.007713 | 0.178258 | Bo1g008020.1 |
| 927 | . | gene | MSTRG.2826 | 1.38196 | 0.007707 | 0.178258 | Bo1g138430.1 |
| 930 | . | gene | MSTRG.11199 | 0.099588 | 0.007736 | 0.178403 | Bo3g143670.1 |
| 929 | . | gene | MSTRG.26494 | 0.790829 | 0.007734 | 0.178403 | |
| 933 | . | gene | MSTRG.3735 | 1.269671 | 0.007762 | 0.178428 | Bo2g012570.1 |
| 931 | . | gene | MSTRG.9542 | 0.142909 | 0.007758 | 0.178428 | Bo3g064940.1 |
| 932 | . | gene | MSTRG.19362 | 1.398905 | 0.007762 | 0.178428 | Bo5g139370.1 |
| 934 | . | gene | MSTRG.13753 | 0.264086 | 0.007782 | 0.178521 | |
| 935 | . | gene | MSTRG.33412 | 1.80496 | 0.007783 | 0.178521 | |
| 937 | . | gene | MSTRG.16930 | 1.492645 | 0.007839 | 0.179093 | Bo5g021400.1 |
| 940 | . | gene | MSTRG.20745 | 2.57861 | 0.007851 | 0.179093 | Bo6g065040.1 |
| 941 | . | gene | MSTRG.25281 | 0.258838 | 0.007858 | 0.179093 | Bo7g100740.1 |
| 936 | . | gene | MSTRG.25512 | 2.707299 | 0.007818 | 0.179093 | Bo7g106490.1 |
| 938 | . | gene | MSTRG.30519 | 4.321298 | 0.007842 | 0.179093 | Bo9g022580.1 |
| 939 | . | gene | MSTRG.34001 | 0.515576 | 0.007849 | 0.179093 | |
| 942 | . | gene | MSTRG.9424 | 0.880432 | 0.007902 | 0.179779 | Bo3g062360.1 |
| 943 | . | gene | MSTRG.19629 | 0.342226 | 0.007905 | 0.179779 | Bo5g146780.1 |
| 946 | . | gene | MSTRG.13597 | 0.62032 | 0.007935 | 0.179901 | Bo4g081170.1 |
| 944 | . | gene | MSTRG.30876 | 1.66339 | 0.007933 | 0.179901 | Bo9g037160.1 |
| 945 | . | gene | MSTRG.17169 | 0.781195 | 0.007933 | 0.179901 | |
| 947 | . | gene | MSTRG.19360 | 3.812698 | 0.007952 | 0.180086 | Bo5g139310.1 |
| 948 | . | gene | MSTRG.31885 | 0.155002 | 0.007996 | 0.180887 | |
| 949 | . | gene | MSTRG.3539 | 0.82866 | 0.008004 | 0.180889 | Bo2g009460.1 |
| 950 | . | gene | MSTRG.11559 | 0.452953 | 0.008034 | 0.181098 | Bo3g158590.1 |
| 952 | . | gene | MSTRG.15202 | 3.507092 | 0.008039 | 0.181098 | Bo4g171890.1 |
| 951 | . | gene | MSTRG.19508 | 0.213279 | 0.008036 | 0.181098 | |
| 954 | . | gene | MSTRG.2348 | 0.124863 | 0.008075 | 0.181259 | Bo1g105130.1 |

| | | | | | | | |
|---|---|---|---|---|---|---|---|
| 955 | BUB3.2 | gene | MSTRG.11941 | 1.845262 | 0.008092 | 0.181259 | Bo3g178240.1 |
| 958 | . | gene | MSTRG.18829 | 3.447409 | 0.0081 | 0.181259 | Bo5g123530.1 |
| 953 | . | gene | MSTRG.24626 | 1.500892 | 0.008073 | 0.181259 | Bo7g084090.1 |
| 959 | . | gene | MSTRG.30389 | 0.24115 | 0.008105 | 0.181259 | Bo9g018860.1 |
| 956 | . | gene | MSTRG.27284 | 2.472778 | 0.008097 | 0.181259 | |
| 957 | . | gene | MSTRG.9402 | 0.648205 | 0.008099 | 0.181259 | |
| 961 | . | gene | MSTRG.14913 | 0.077664 | 0.00816 | 0.181987 | Bo4g151820.1 |
| 963 | . | gene | MSTRG.15077 | 0.30552 | 0.008171 | 0.181987 | Bo4g163640.1 |
| 962 | . | gene | MSTRG.17661 | 0.321618 | 0.008166 | 0.181987 | Bo5g049430.1 |
| 960 | . | gene | MSTRG.19407 | 4.896061 | 0.008158 | 0.181987 | Bo5g139880.1 |
| 964 | . | gene | MSTRG.27689 | 2.098884 | 0.008188 | 0.182167 | Bo8g069870.1 |
| 966 | . | gene | MSTRG.10297 | 0.375075 | 0.008231 | 0.182565 | Bo3g094030.1 |
| 965 | . | gene | MSTRG.12231 | 2.457401 | 0.008219 | 0.182565 | Bo4g005780.1 |
| 968 | . | gene | MSTRG.13186 | 9.970184 | 0.00824 | 0.182565 | Bo4g049480.1 |
| 967 | . | gene | MSTRG.26566 | 1.465195 | 0.008234 | 0.182565 | Bo8g016260.1 |
| 974 | . | gene | MSTRG.8364 | 0.708551 | 0.008312 | 0.182937 | Bo3g032780.1 |
| 976 | . | gene | MSTRG.25891 | 1.379812 | 0.008325 | 0.182937 | Bo7g113860.1 |
| 972 | . | gene | MSTRG.27305 | 1.750604 | 0.008302 | 0.182937 | Bo8g058750.1 |
| 970 | . | gene | MSTRG.27546 | 1.43345 | 0.008287 | 0.182937 | Bo8g066490.1 |
| 969 | . | gene | MSTRG.22926 | 0.662742 | 0.008282 | 0.182937 | |
| 971 | . | gene | MSTRG.2374 | 0.139002 | 0.008299 | 0.182937 | |
| 973 | . | gene | MSTRG.13094 | 1.327954 | 0.008309 | 0.182937 | |
| 975 | . | gene | MSTRG.6122 | 0.553661 | 0.008323 | 0.182937 | |
| 977 | . | gene | MSTRG.15080 | 1.87877 | 0.008343 | 0.182999 | Bo4g163710.1 |
| 978 | . | gene | MSTRG.29324 | 4.162912 | 0.008345 | 0.182999 | Bo8g108500.1 |
| 980 | . | gene | MSTRG.21071 | 2.034856 | 0.008365 | 0.183075 | Bo6g074920.1 |
| 979 | . | gene | MSTRG.14509 | 0.212546 | 0.008364 | 0.183075 | |
| 982 | . | gene | MSTRG.1867 | 1.447339 | 0.008411 | 0.183407 | Bo1g071050.1 |
| 988 | . | gene | MSTRG.7070 | 1.343321 | 0.008456 | 0.183407 | Bo3g006780.1 |

| | | | | | | | |
|---|---|---|---|---|---|---|---|
| 983 | . | gene | MSTRG.9870 | 3.159396 | 0.008432 | 0.183407 | Bo3g076020.1 |
| 986 | . | gene | MSTRG.11316 | 0.416064 | 0.008445 | 0.183407 | Bo3g149360.1 |
| 984 | . | gene | MSTRG.13557 | 1.44179 | 0.008436 | 0.183407 | Bo4g078540.1 |
| 985 | . | gene | MSTRG.23705 | 0.057781 | 0.008438 | 0.183407 | Bo7g050630.1 |
| 987 | . | gene | MSTRG.26696 | 0.741282 | 0.008452 | 0.183407 | Bo8g025650.1 |
| 989 | . | gene | MSTRG.31981 | 0.26976 | 0.008459 | 0.183407 | Bo9g108370.1 |
| 981 | . | gene | MSTRG.32637 | 0.629393 | 0.008409 | 0.183407 | |
| 990 | . | gene | MSTRG.1828 | 0.281215 | 0.008466 | 0.183407 | |
| 992 | . | gene | MSTRG.327 | 2.50271 | 0.008494 | 0.183636 | Bo1g007370.1 |
| 991 | . | gene | MSTRG.9689 | 0.091236 | 0.008486 | 0.183636 | |
| 995 | . | gene | MSTRG.9317 | 49.08921 | 0.008552 | 0.184247 | Bo3g060500.1 |
| 993 | . | gene | MSTRG.22653 | 2.03408 | 0.008545 | 0.184247 | Bo6g122380.1 |
| 994 | . | gene | MSTRG.28512 | 1.383441 | 0.008547 | 0.184247 | Bo8g092640.1 |
| 996 | . | gene | MSTRG.7355 | 0.197026 | 0.008556 | 0.184247 | |
| 997 | . | gene | MSTRG.6575 | 0.413784 | 0.008578 | 0.184517 | Bo2g161610.1 |
| 998 | . | gene | MSTRG.27156 | 1.211094 | 0.008593 | 0.184654 | |
| 999 | . | gene | MSTRG.3178 | 1.432908 | 0.008654 | 0.185642 | Bo1g151900.1 |
| 1000 | . | gene | MSTRG.21926 | 3.378343 | 0.00866 | 0.185642 | Bo6g104030.1 |
| 1001 | . | gene | MSTRG.33426 | 1.192833 | 0.008665 | 0.185642 | Bo9g174220.1 |
| 1003 | . | gene | MSTRG.18063 | 0.369353 | 0.008688 | 0.185775 | Bo5g076310.1 |
| 1002 | . | gene | MSTRG.20949 | 1.344661 | 0.008683 | 0.185775 | Bo6g069970.1 |
| 1004 | . | gene | MSTRG.16025 | 1.394577 | 0.0087 | 0.185845 | Bo5g001040.1 |
| 1006 | . | gene | MSTRG.473 | 1.54211 | 0.008736 | 0.185925 | Bo1g010390.1 |
| 1025 | . | gene | MSTRG.1003 | 1.257268 | 0.008897 | 0.185925 | Bo1g024270.1 |
| 1027 | . | gene | MSTRG.3103 | 0.632816 | 0.008908 | 0.185925 | Bo1g147670.1 |
| 1010 | . | gene | MSTRG.3784 | 1.361938 | 0.008781 | 0.185925 | Bo2g013400.1 |
| 1023 | . | gene | MSTRG.9534 | 0.550021 | 0.008889 | 0.185925 | Bo3g064890.1 |
| 1011 | . | gene | MSTRG.10517 | 1.239582 | 0.008783 | 0.185925 | Bo3g102980.1 |
| 1008 | . | gene | MSTRG.11143 | 0.703934 | 0.008768 | 0.185925 | Bo3g141680.1 |

| | | | | | | | |
|---|---|---|---|---|---|---|---|
| 1016 | . | gene | MSTRG.11487 | 0.386558 | 0.008853 | 0.185925 | Bo3g154220.1 |
| 1020 | . | gene | MSTRG.12281 | 0.702455 | 0.008865 | 0.185925 | Bo4g010430.1 |
| 1018 | . | gene | MSTRG.12402 | 0.296357 | 0.00886 | 0.185925 | Bo4g018460.1 |
| 1007 | . | gene | MSTRG.18094 | 0.143776 | 0.008763 | 0.185925 | Bo5g078790.1 |
| 1005 | . | gene | MSTRG.18974 | 0.537219 | 0.008725 | 0.185925 | Bo5g127640.1 |
| 1028 | . | gene | MSTRG.19298 | 10.32378 | 0.008912 | 0.185925 | Bo5g137410.1 |
| 1015 | . | gene | MSTRG.19561 | 1.212442 | 0.008815 | 0.185925 | Bo5g144830.1 |
| 1012 | . | gene | MSTRG.25863 | 0.36709 | 0.008795 | 0.185925 | Bo7g113340.1 |
| 1022 | . | gene | MSTRG.28105 | 1.363116 | 0.008889 | 0.185925 | Bo8g081740.1 |
| 1021 | . | gene | MSTRG.29651 | 0.808422 | 0.008881 | 0.185925 | Bo8g117130.1 |
| 1019 | . | gene | MSTRG.33284 | 1.208082 | 0.008863 | 0.185925 | Bo9g171400.1 |
| 1009 | . | gene | MSTRG.34118 | 0.397323 | 0.008774 | 0.185925 | |
| 1013 | . | gene | MSTRG.29387 | 1.56762 | 0.008812 | 0.185925 | |
| 1014 | . | gene | MSTRG.35711 | 4.579513 | 0.008814 | 0.185925 | |
| 1017 | . | gene | MSTRG.9716 | 0.466735 | 0.00886 | 0.185925 | |
| 1024 | . | gene | MSTRG.32530 | 0.173072 | 0.008891 | 0.185925 | |
| 1026 | . | gene | MSTRG.31158 | 0.388774 | 0.008908 | 0.185925 | |
| 1029 | . | gene | MSTRG.6362 | 2.828387 | 0.008933 | 0.186193 | Bo2g155190.1 |
| 1030 | . | gene | MSTRG.29783 | 0.931783 | 0.008961 | 0.186479 | Bo9g003750.1 |
| 1031 | . | gene | MSTRG.33449 | 0.370901 | 0.008964 | 0.186479 | Bo9g174460.1 |
| 1034 | . | gene | MSTRG.10438 | 1.216409 | 0.008993 | 0.186523 | Bo3g100850.1 |
| 1032 | . | gene | MSTRG.13558 | 1.394954 | 0.008976 | 0.186523 | Bo4g078560.1 |
| 1033 | . | gene | MSTRG.19753 | 1.629957 | 0.008986 | 0.186523 | Bo5g152600.1 |
| 1037 | CKA1 | gene | MSTRG.6735 | 0.352941 | 0.009066 | 0.186795 | Bo2g167980.1 |
| 1038 | . | gene | MSTRG.15089 | 1.720982 | 0.009066 | 0.186795 | Bo4g164790.1 |
| 1040 | . | gene | MSTRG.15196 | 0.39725 | 0.009072 | 0.186795 | Bo4g170750.1 |
| 1035 | . | gene | MSTRG.21727 | 4.054948 | 0.009036 | 0.186795 | Bo6g095850.1 |
| 1042 | . | gene | MSTRG.26216 | 10.56046 | 0.009091 | 0.186795 | Bo7g118850.1 |
| 1043 | . | gene | MSTRG.27696 | 0.683654 | 0.009092 | 0.186795 | Bo8g069950.1 |

| | | | | | | | |
|---|---|---|---|---|---|---|---|
| 10 39 | . | gene | MSTRG.2846 2 | 1.1889 21 | 0.0090 67 | 0.1867 95 | Bo8g0918 20.1 |
| 10 44 | SIR | gene | MSTRG.3368 0 | 2.0154 81 | 0.0090 93 | 0.1867 95 | Bo9g1788 90.1 |
| 10 36 | . | gene | MSTRG.2291 5 | 0.7909 21 | 0.0090 44 | 0.1867 95 | |
| 10 41 | . | gene | MSTRG.3471 7 | 0.2819 53 | 0.0090 75 | 0.1867 95 | |
| 10 45 | . | gene | MSTRG.3296 7 | 1.1809 38 | 0.0091 02 | 0.1867 95 | Bo9g1618 00.1 |
| 10 46 | . | gene | MSTRG.2770 2 | 7.5733 61 | 0.0091 25 | 0.1871 03 | Bo8g0700 50.1 |
| 10 62 | . | gene | MSTRG.1360 5 | 0.6495 97 | 0.0092 69 | 0.1871 26 | Bo4g0823 50.1 |
| 10 49 | . | gene | MSTRG.1765 5 | 0.1767 34 | 0.0091 8 | 0.1871 26 | Bo5g0493 00.1 |
| 10 55 | . | gene | MSTRG.2094 7 | 3.6804 63 | 0.0092 24 | 0.1871 26 | Bo6g0699 20.1 |
| 10 59 | . | gene | MSTRG.2226 0 | 2.0747 19 | 0.0092 5 | 0.1871 26 | Bo6g1158 10.1 |
| 10 58 | . | gene | MSTRG.2232 0 | 0.3583 2 | 0.0092 38 | 0.1871 26 | Bo6g1168 10.1 |
| 10 53 | . | gene | MSTRG.2296 3 | 1.8887 12 | 0.0092 19 | 0.1871 26 | Bo7g0099 90.1 |
| 10 61 | . | gene | MSTRG.2688 6 | 0.3495 62 | 0.0092 68 | 0.1871 26 | Bo8g0341 50.1 |
| 10 50 | . | gene | MSTRG.2941 9 | 0.4880 55 | 0.0091 94 | 0.1871 26 | Bo8g1106 70.1 |
| 10 47 | . | gene | MSTRG.3467 4 | 0.2735 17 | 0.0091 47 | 0.1871 26 | |
| 10 48 | . | gene | MSTRG.3034 6 | 0.6112 79 | 0.0091 47 | 0.1871 26 | |
| 10 51 | . | gene | MSTRG.1425 | 0.3234 53 | 0.0092 02 | 0.1871 26 | |
| 10 52 | . | gene | MSTRG.2655 9 | 0.2010 46 | 0.0092 04 | 0.1871 26 | |
| 10 54 | . | gene | MSTRG.3484 3 | 8.1094 53 | 0.0092 23 | 0.1871 26 | |
| 10 56 | . | gene | MSTRG.6621 | 0.8758 36 | 0.0092 32 | 0.1871 26 | |
| 10 57 | . | gene | MSTRG.4221 | 0.3558 92 | 0.0092 36 | 0.1871 26 | |
| 10 60 | . | gene | MSTRG.6173 | 1.0693 4 | 0.0092 51 | 0.1871 26 | |
| 10 63 | . | gene | MSTRG.3136 4 | 0.1877 91 | 0.0092 75 | 0.1871 26 | |
| 10 64 | . | gene | MSTRG.4243 | 0.3374 77 | 0.0093 06 | 0.1875 35 | Bo2g0327 90.1 |
| 10 65 | . | gene | MSTRG.1692 1 | 4.1230 23 | 0.0093 13 | 0.1875 35 | Bo5g0212 80.1 |
| 10 66 | . | gene | MSTRG.3054 9 | 0.4340 04 | 0.0093 39 | 0.1878 9 | Bo9g0241 40.1 |
| 10 75 | . | gene | MSTRG.443 | 0.7123 6 | 0.0094 23 | 0.1879 94 | Bo1g0100 00.1 |
| 10 70 | . | gene | MSTRG.9185 | 0.1683 1 | 0.0093 89 | 0.1879 94 | Bo3g0556 50.1 |
| 10 71 | . | gene | MSTRG.1024 8 | 5.9071 12 | 0.0094 | 0.1879 94 | Bo3g0920 80.1 |

| | | | | | | | |
|---|---|---|---|---|---|---|---|
| 10 72 | . | gene | MSTRG.1682 2 | 0.5923 3 | 0.0094 12 | 0.1879 94 | Bo5g0196 70.1 |
| 10 68 | . | gene | MSTRG.1724 3 | 0.4167 27 | 0.0093 76 | 0.1879 94 | Bo5g0320 00.1 |
| 10 67 | . | gene | MSTRG.1749 7 | 5.7997 05 | 0.0093 71 | 0.1879 94 | Bo5g0427 90.1 |
| 10 74 | . | gene | MSTRG.2049 2 | 2.5266 69 | 0.0094 18 | 0.1879 94 | Bo6g0479 00.1 |
| 10 69 | . | gene | MSTRG.9623 | 0.3113 01 | 0.0093 83 | 0.1879 94 | |
| 10 73 | . | gene | MSTRG.6860 | 0.6653 54 | 0.0094 12 | 0.1879 94 | |
| 10 76 | . | gene | MSTRG.7011 | 1.6252 19 | 0.0094 55 | 0.1884 54 | Bo3g0050 80.1 |
| 10 77 | . | gene | MSTRG.2173 3 | 0.7088 07 | 0.0094 68 | 0.1885 41 | Bo6g0959 70.1 |
| 10 78 | . | gene | MSTRG.4291 | 0.5267 12 | 0.0095 07 | 0.1891 36 | |
| 10 87 | . | gene | MSTRG.907 | 0.7443 75 | 0.0096 37 | 0.1896 71 | Bo1g0217 00.1 |
| 10 89 | . | gene | MSTRG.6282 | 1.7326 87 | 0.0096 38 | 0.1896 71 | Bo2g1498 70.1 |
| 10 82 | . | gene | MSTRG.1663 8 | 0.2417 39 | 0.0095 87 | 0.1896 71 | Bo5g0147 50.1 |
| 10 86 | . | gene | MSTRG.1924 5 | 2.1919 73 | 0.0096 22 | 0.1896 71 | Bo5g1366 50.1 |
| 10 80 | . | gene | MSTRG.2878 6 | 0.1669 58 | 0.0095 66 | 0.1896 71 | Bo8g0991 10.1 |
| 10 79 | . | gene | MSTRG.1690 7 | 2.6438 97 | 0.0095 55 | 0.1896 71 | |
| 10 81 | . | gene | MSTRG.1806 7 | 0.4219 46 | 0.0095 81 | 0.1896 71 | |
| 10 83 | . | gene | MSTRG.3439 5 | 0.2249 72 | 0.0095 92 | 0.1896 71 | |
| 10 84 | . | gene | MSTRG.3554 4 | 0.4868 43 | 0.0095 93 | 0.1896 71 | |
| 10 85 | . | gene | MSTRG.1370 9 | 0.4089 29 | 0.0095 99 | 0.1896 71 | |
| 10 88 | . | gene | MSTRG.2322 2 | 0.3148 57 | 0.0096 38 | 0.1896 71 | |
| 10 90 | . | gene | MSTRG.3410 6 | 0.7337 93 | 0.0096 4 | 0.1896 71 | |
| 10 92 | . | gene | MSTRG.7742 | 0.7118 46 | 0.0096 84 | 0.1897 15 | Bo3g0196 10.1 |
| 10 91 | . | gene | MSTRG.1576 0 | 0.1018 99 | 0.0096 72 | 0.1897 15 | Bo4g1907 10.1 |
| 10 98 | . | gene | MSTRG.1588 2 | 1.5569 46 | 0.0097 28 | 0.1897 15 | Bo4g1947 90.1 |
| 10 96 | . | gene | MSTRG.1938 0 | 0.2696 36 | 0.0097 01 | 0.1897 15 | Bo5g1396 00.1 |
| 10 99 | . | gene | MSTRG.2084 4 | 1.7117 42 | 0.0097 29 | 0.1897 15 | Bo6g0675 50.1 |
| 10 94 | . | gene | MSTRG.2359 0 | 1.3032 75 | 0.0096 96 | 0.1897 15 | Bo7g0457 50.1 |
| 11 00 | . | gene | MSTRG.2458 8 | 0.3767 49 | 0.0097 3 | 0.1897 15 | Bo7g0825 70.1 |
| 10 93 | . | gene | MSTRG.3401 5 | 0.4913 73 | 0.0096 9 | 0.1897 15 | |

| | | | | | | | |
|---|---|---|---|---|---|---|---|
| 1095 | . | gene | MSTRG.24204 | 0.964888 | 0.009701 | 0.189715 | |
| 1097 | . | gene | MSTRG.23600 | 8.646093 | 0.00972 | 0.189715 | |
| 1104 | . | gene | MSTRG.276 | 0.603711 | 0.009809 | 0.18979 | Bo1g006670.1 |
| 1113 | . | gene | MSTRG.2790 | 1.470437 | 0.009863 | 0.18979 | Bo1g134680.1 |
| 1112 | . | gene | MSTRG.7003 | 1.215423 | 0.009854 | 0.18979 | Bo3g004980.1 |
| 1115 | . | gene | MSTRG.13055 | 0.28043 | 0.009874 | 0.18979 | Bo4g043430.1 |
| 1114 | . | gene | MSTRG.15009 | 0.376287 | 0.00987 | 0.18979 | Bo4g158620.1 |
| 1108 | . | gene | MSTRG.22654 | 1.285916 | 0.009825 | 0.18979 | Bo6g122390.1 |
| 1110 | . | gene | MSTRG.25422 | 1.75882 | 0.009838 | 0.18979 | Bo7g104990.1 |
| 1102 | . | gene | MSTRG.25992 | 4.07759 | 0.009759 | 0.18979 | Bo7g115290.1 |
| 1111 | . | gene | MSTRG.27125 | 2.823995 | 0.00984 | 0.18979 | Bo8g050630.1 |
| 1106 | . | gene | MSTRG.29843 | 1.474817 | 0.009817 | 0.18979 | Bo9g004470.1 |
| 1103 | . | gene | MSTRG.30968 | 0.534696 | 0.009776 | 0.18979 | Bo9g043930.1 |
| 1116 | . | gene | MSTRG.31495 | 4.25367 | 0.009882 | 0.18979 | Bo9g076350.1 |
| 1101 | . | gene | MSTRG.4488 | 0.570122 | 0.009753 | 0.18979 | |
| 1105 | . | gene | MSTRG.2202 | 0.359485 | 0.009815 | 0.18979 | |
| 1107 | . | gene | MSTRG.31912 | 0.705474 | 0.009823 | 0.18979 | |
| 1109 | . | gene | MSTRG.35656 | 0.298398 | 0.009827 | 0.18979 | |
| 1117 | . | gene | MSTRG.11936 | 0.208005 | 0.009885 | 0.18979 | |
| 1118 | . | gene | MSTRG.20885 | 2.627035 | 0.0099 | 0.189914 | Bo6g068020.1 |
| 1119 | . | gene | MSTRG.23666 | 1.323258 | 0.009933 | 0.190243 | Bo7g049960.1 |
| 1120 | . | gene | MSTRG.29727 | 1.619284 | 0.009935 | 0.190243 | Bo9g001010.1 |
| 1121 | . | gene | MSTRG.28059 | 2.087146 | 0.009984 | 0.191013 | Bo8g081060.1 |
| 1122 | . | gene | MSTRG.10027 | 0.106704 | 0.01 | 0.191154 | |
| 1124 | . | gene | MSTRG.5238 | 2.094105 | 0.010019 | 0.191166 | Bo2g094720.1 |
| 1123 | . | gene | MSTRG.23013 | 0.282492 | 0.010018 | 0.191166 | |
| 1131 | . | gene | MSTRG.3585 | 1.251681 | 0.010096 | 0.191182 | Bo2g010060.1 |
| 1133 | . | gene | MSTRG.5438 | 28.76272 | 0.010104 | 0.191182 | Bo2g102190.1 |
| 1132 | . | gene | MSTRG.13831 | 0.313607 | 0.010101 | 0.191182 | Bo4g101150.1 |

| | | | | | | | |
|---|---|---|---|---|---|---|---|
| 11 29 | . | gene | MSTRG.1726 6 | 1.1609 3 | 0.0100 71 | 0.1911 82 | Bo5g0342 30.1 |
| 11 25 | . | gene | MSTRG.1885 9 | 0.1270 26 | 0.0100 35 | 0.1911 82 | Bo5g1260 80.1 |
| 11 34 | . | gene | MSTRG.2652 1 | 1.7777 72 | 0.0101 09 | 0.1911 82 | Bo8g0127 10.1 |
| 11 26 | . | gene | MSTRG.2986 3 | 1.0212 49 | 0.0100 39 | 0.1911 82 | Bo9g0057 40.1 |
| 11 27 | . | gene | MSTRG.1432 5 | 0.1590 43 | 0.0100 58 | 0.1911 82 | |
| 11 28 | . | gene | MSTRG.2634 0 | 0.3152 91 | 0.0100 62 | 0.1911 82 | |
| 11 30 | . | gene | MSTRG.3332 6 | 0.1782 07 | 0.0100 86 | 0.1911 82 | |
| 11 35 | . | gene | MSTRG.880 | 1.7585 77 | 0.0101 2 | 0.1912 23 | Bo1g0203 60.1 |
| 11 37 | . | gene | MSTRG.3066 0 | 1.3973 77 | 0.0101 46 | 0.1913 91 | Bo9g0281 20.1 |
| 11 36 | . | gene | MSTRG.3613 4 | 1.5831 55 | 0.0101 45 | 0.1913 91 | |
| 11 40 | . | gene | MSTRG.259 | 1.3131 23 | 0.0101 75 | 0.1914 32 | Bo1g0064 90.1 |
| 11 38 | . | gene | MSTRG.1608 2 | 0.8165 52 | 0.0101 72 | 0.1914 32 | Bo5g0028 30.1 |
| 11 39 | . | gene | MSTRG.2104 4 | 1.9514 24 | 0.0101 73 | 0.1914 32 | Bo6g0746 00.1 |
| 11 41 | . | gene | MSTRG.2095 5 | 0.0824 23 | 0.0101 91 | 0.1915 06 | Bo6g0710 70.1 |
| 11 42 | . | gene | MSTRG.2779 1 | 2.6689 73 | 0.0101 97 | 0.1915 06 | Bo8g0714 70.1 |
| 11 50 | . | gene | MSTRG.8052 | 1.2673 64 | 0.0102 97 | 0.1917 18 | Bo3g0264 00.1 |
| 11 51 | . | gene | MSTRG.9086 | 1.8342 17 | 0.0102 98 | 0.1917 18 | Bo3g0533 40.1 |
| 11 43 | . | gene | MSTRG.1371 4 | 11.937 97 | 0.0102 27 | 0.1917 18 | Bo4g0941 20.1 |
| 11 48 | . | gene | MSTRG.2588 6 | 2.1502 56 | 0.0102 88 | 0.1917 18 | Bo7g1137 80.1 |
| 11 52 | . | gene | MSTRG.2942 8 | 1.2503 42 | 0.0102 98 | 0.1917 18 | Bo8g1107 80.1 |
| 11 44 | . | gene | MSTRG.3188 7 | 1.5254 71 | 0.0102 29 | 0.1917 18 | Bo9g1010 80.1 |
| 11 46 | . | gene | MSTRG.3206 8 | 0.6457 59 | 0.0102 64 | 0.1917 18 | Bo9g1114 50.1 |
| 11 49 | . | gene | MSTRG.3289 3 | 1.6776 74 | 0.0102 95 | 0.1917 18 | Bo9g1596 30.1 |
| 11 45 | . | gene | MSTRG.3132 3 | 0.1724 76 | 0.0102 38 | 0.1917 18 | |
| 11 47 | . | gene | MSTRG.1325 3 | 1.8007 48 | 0.0102 64 | 0.1917 18 | |
| 11 55 | . | gene | MSTRG.557 | 0.4831 97 | 0.0103 29 | 0.1917 23 | Bo1g0116 70.1 |
| 11 56 | . | gene | MSTRG.9824 | 1.5701 65 | 0.0103 34 | 0.1917 23 | Bo3g0733 80.1 |
| 11 54 | . | gene | MSTRG.1474 7 | 0.2185 38 | 0.0103 21 | 0.1917 23 | Bo4g1441 20.1 |
| 11 53 | . | gene | MSTRG.1412 | 0.8281 59 | 0.0103 18 | 0.1917 23 | |

| | | | | | | | |
|---|---|---|---|---|---|---|---|
| 11 58 | . | gene | MSTRG.1864 6 | 0.6303 01 | 0.0103 58 | 0.1918 44 | Bo5g1156 10.1 |
| 11 57 | . | gene | MSTRG.3137 8 | 0.6664 8 | 0.0103 5 | 0.1918 44 | |
| 11 59 | . | gene | MSTRG.1141 3 | 0.6030 69 | 0.0103 71 | 0.1918 54 | Bo3g1530 60.1 |
| 11 60 | . | gene | MSTRG.1292 6 | 0.6379 11 | 0.0103 77 | 0.1918 54 | Bo4g0386 70.1 |
| 11 61 | . | gene | MSTRG.9239 | 2.0605 92 | 0.0103 87 | 0.1918 73 | Bo3g0574 70.1 |
| 11 62 | . | gene | MSTRG.1489 3 | 0.1396 66 | 0.0103 99 | 0.1919 26 | |
| 11 63 | . | gene | MSTRG.1217 6 | 0.8297 16 | 0.0104 09 | 0.1919 49 | Bo3g1858 50.1 |
| 11 65 | . | gene | MSTRG.1932 6 | 0.7164 7 | 0.0104 51 | 0.1923 97 | Bo5g1377 60.1 |
| 11 64 | . | gene | MSTRG.3268 3 | 1.7445 3 | 0.0104 43 | 0.1923 97 | Bo9g1503 70.1 |
| 11 66 | . | gene | MSTRG.3533 8 | 3.1133 07 | 0.0104 82 | 0.1928 07 | |
| 11 67 | . | gene | MSTRG.6199 | 0.4691 59 | 0.0105 05 | 0.1930 6 | Bo2g1478 20.1 |
| 11 68 | . | gene | MSTRG.1347 0 | 2.1764 32 | 0.0105 27 | 0.1931 41 | Bo4g0731 30.1 |
| 11 69 | . | gene | MSTRG.2566 1 | 0.7956 69 | 0.0105 27 | 0.1931 41 | Bo7g1092 10.1 |
| 11 70 | . | gene | MSTRG.2660 2 | 0.1631 16 | 0.0105 4 | 0.1932 11 | |
| 11 71 | . | gene | MSTRG.9576 | 2.1182 01 | 0.0105 63 | 0.1933 38 | Bo3g0654 10.1 |
| 11 72 | . | gene | MSTRG.1867 4 | 1.2630 36 | 0.0105 65 | 0.1933 38 | Bo5g1170 40.1 |
| 11 76 | . | gene | MSTRG.4183 | 1.8700 35 | 0.0106 58 | 0.1941 44 | Bo2g0287 00.1 |
| 11 73 | . | gene | MSTRG.8108 | 2.9795 98 | 0.0106 23 | 0.1941 44 | Bo3g0273 00.1 |
| 11 74 | . | gene | MSTRG.2595 9 | 0.5806 74 | 0.0106 4 | 0.1941 44 | Bo7g1148 30.1 |
| 11 79 | . | gene | MSTRG.2641 4 | 0.2810 75 | 0.0106 73 | 0.1941 44 | Bo8g0059 40.1 |
| 11 75 | . | gene | MSTRG.1565 8 | 0.5015 23 | 0.0106 48 | 0.1941 44 | |
| 11 77 | . | gene | MSTRG.1489 6 | 0.6260 35 | 0.0106 65 | 0.1941 44 | |
| 11 78 | . | gene | MSTRG.1146 9 | 0.4294 87 | 0.0106 72 | 0.1941 44 | |
| 11 80 | . | gene | MSTRG.3558 | 1.2620 27 | 0.0106 96 | 0.1944 01 | Bo2g0098 10.1 |
| 11 81 | . | gene | MSTRG.2102 0 | 0.2035 29 | 0.0107 08 | 0.1944 01 | Bo6g0732 10.1 |
| 11 82 | . | gene | MSTRG.3154 4 | 0.2182 9 | 0.0107 14 | 0.1944 01 | Bo9g0782 20.1 |
| 11 83 | . | gene | MSTRG.1931 0 | 0.4826 05 | 0.0107 31 | 0.1945 41 | Bo5g1376 00.1 |
| 11 84 | . | gene | MSTRG.3442 2 | 2.7432 64 | 0.0107 58 | 0.1948 64 | |
| 11 85 | . | gene | MSTRG.2802 0 | 2.1437 11 | 0.0107 68 | 0.1948 84 | Bo8g0806 00.1 |

| | | | | | | | |
|---|---|---|---|---|---|---|---|
| 1186 | . | gene | MSTRG.24223 | 4.348582 | 0.010779 | 0.194927 | Bo7g072720.1 |
| 1191 | . | gene | MSTRG.6994 | 0.574133 | 0.010833 | 0.195076 | Bo3g004910.1 |
| 1189 | . | gene | MSTRG.8987 | 0.349328 | 0.010819 | 0.195076 | Bo3g048700.1 |
| 1188 | . | gene | MSTRG.9577 | 1.333003 | 0.010817 | 0.195076 | Bo3g065440.1 |
| 1190 | . | gene | MSTRG.18982 | 0.394489 | 0.010825 | 0.195076 | Bo5g127740.1 |
| 1187 | . | gene | MSTRG.32865 | 1.349862 | 0.010798 | 0.195076 | Bo9g157340.1 |
| 1192 | . | gene | MSTRG.23398 | 0.512468 | 0.010843 | 0.195098 | Bo7g038780.1 |
| 1194 | . | gene | MSTRG.15259 | 1.611086 | 0.010862 | 0.1951 | Bo4g173750.1 |
| 1193 | . | gene | MSTRG.36033 | 0.200412 | 0.010859 | 0.1951 | |
| 1195 | . | gene | MSTRG.4579 | 2.542883 | 0.010872 | 0.195115 | Bo2g053880.1 |
| 1198 | . | gene | MSTRG.3077 | 1.371523 | 0.010908 | 0.195283 | Bo1g147230.1 |
| 1196 | . | gene | MSTRG.34016 | 1.226425 | 0.010898 | 0.195283 | |
| 1197 | . | gene | MSTRG.17623 | 0.370814 | 0.010903 | 0.195283 | |
| 1199 | . | gene | MSTRG.1065 | 1.285767 | 0.010923 | 0.195378 | Bo1g027230.1 |
| 1200 | . | gene | MSTRG.9139 | 1.527865 | 0.010934 | 0.195419 | Bo3g054950.1 |
| 1203 | . | gene | MSTRG.6624 | 0.579751 | 0.010976 | 0.195464 | Bo2g165350.1 |
| 1205 | . | gene | MSTRG.14962 | 0.260272 | 0.010982 | 0.195464 | Bo4g154580.1 |
| 1202 | . | gene | MSTRG.26485 | 0.449798 | 0.010969 | 0.195464 | Bo8g010890.1 |
| 1201 | . | gene | MSTRG.17647 | 5.091365 | 0.010946 | 0.195464 | |
| 1204 | . | gene | MSTRG.6462 | 0.077577 | 0.01098 | 0.195464 | |
| 1206 | . | gene | MSTRG.25453 | 1.830935 | 0.010994 | 0.195515 | Bo7g105400.1 |
| 1208 | . | gene | MSTRG.8120 | 0.303303 | 0.01102 | 0.195648 | Bo3g027560.1 |
| 1207 | . | gene | MSTRG.31330 | 0.862656 | 0.011018 | 0.195648 | Bo9g068090.1 |
| 1210 | . | gene | MSTRG.30058 | 0.645588 | 0.01104 | 0.195673 | Bo9g009910.1 |
| 1209 | . | gene | MSTRG.35693 | 0.375415 | 0.011032 | 0.195673 | |
| 1211 | . | gene | MSTRG.24659 | 1.513159 | 0.011072 | 0.195928 | Bo7g084570.1 |
| 1213 | . | gene | MSTRG.29231 | 2.364127 | 0.011081 | 0.195928 | Bo8g106810.1 |
| 1212 | . | gene | MSTRG.14691 | 0.572303 | 0.011073 | 0.195928 | |
| 1214 | . | gene | MSTRG.2440 | 4.155608 | 0.011132 | 0.19667 | Bo1g108520.1 |

| | | | | | | | |
|---|---|---|---|---|---|---|---|
| 12 15 | . | gene | MSTRG.3164 4 | 1.5215 93 | 0.0111 44 | 0.1967 14 | |
| 12 16 | . | gene | MSTRG.9177 | 0.9623 77 | 0.0112 13 | 0.1975 6 | Bo3g0555 30.1 |
| 12 17 | . | gene | MSTRG.1150 4 | 2.5320 42 | 0.0112 14 | 0.1975 6 | Bo3g1555 90.1 |
| 12 18 | . | gene | MSTRG.2326 0 | 0.1243 98 | 0.0112 2 | 0.1975 6 | Bo7g0304 90.1 |
| 12 20 | . | gene | MSTRG.7120 | 0.6622 88 | 0.0112 57 | 0.1978 52 | Bo3g0075 80.1 |
| 12 19 | . | gene | MSTRG.2575 2 | 1.4953 21 | 0.0112 48 | 0.1978 52 | Bo7g1105 70.1 |
| 12 21 | . | gene | MSTRG.2778 5 | 0.2065 1 | 0.0112 64 | 0.1978 52 | Bo8g0714 10.1 |
| 12 28 | . | gene | MSTRG.4562 | 1.5529 78 | 0.0113 57 | 0.1979 03 | Bo2g0526 40.1 |
| 12 27 | . | gene | MSTRG.5722 | 1.4412 75 | 0.0113 49 | 0.1979 03 | Bo2g1243 30.1 |
| 12 33 | . | gene | MSTRG.6677 | 0.8536 1 | 0.0113 82 | 0.1979 03 | Bo2g1661 80.1 |
| 12 24 | . | gene | MSTRG.9544 | 0.1146 88 | 0.0113 1 | 0.1979 03 | Bo3g0649 80.1 |
| 12 22 | . | gene | MSTRG.1493 6 | 3.2055 1 | 0.0112 83 | 0.1979 03 | Bo4g1540 90.1 |
| 12 31 | . | gene | MSTRG.1516 9 | 4.4674 99 | 0.0113 71 | 0.1979 03 | Bo4g1692 40.1 |
| 12 32 | . | gene | MSTRG.1545 0 | 0.4821 37 | 0.0113 72 | 0.1979 03 | Bo4g1828 40.1 |
| 12 29 | . | gene | MSTRG.2533 7 | 0.4397 17 | 0.0113 63 | 0.1979 03 | Bo7g1037 00.1 |
| 12 23 | . | gene | MSTRG.3543 3 | 4.1424 49 | 0.0113 09 | 0.1979 03 | |
| 12 25 | . | gene | MSTRG.3448 4 | 1.7267 8 | 0.0113 4 | 0.1979 03 | |
| 12 26 | . | gene | MSTRG.1881 | 0.2915 05 | 0.0113 48 | 0.1979 03 | |
| 12 30 | . | gene | MSTRG.3309 5 | 0.4920 46 | 0.0113 66 | 0.1979 03 | |
| 12 34 | . | gene | MSTRG.1501 9 | 0.9293 75 | 0.0113 87 | 0.1979 03 | |
| 12 35 | . | gene | MSTRG.9608 | 1.5446 57 | 0.0114 37 | 0.1980 64 | Bo3g0658 80.1 |
| 12 40 | . | gene | MSTRG.1986 5 | 1.1942 78 | 0.0114 6 | 0.1980 64 | Bo6g0057 10.1 |
| 12 38 | . | gene | MSTRG.2267 2 | 1.3117 6 | 0.0114 49 | 0.1980 64 | Bo6g1236 60.1 |
| 12 39 | . | gene | MSTRG.2362 6 | 0.0594 29 | 0.0114 5 | 0.1980 64 | Bo7g0483 10.1 |
| 12 36 | . | gene | MSTRG.2451 2 | 0.5300 74 | 0.0114 39 | 0.1980 64 | |
| 12 37 | . | gene | MSTRG.2012 | 0.0994 61 | 0.0114 41 | 0.1980 64 | |
| 12 41 | . | gene | MSTRG.6074 | 1.4902 23 | 0.0114 61 | 0.1980 64 | |
| 12 42 | . | gene | MSTRG.1950 5 | 0.7193 97 | 0.0114 7 | 0.1980 67 | |
| 12 43 | . | gene | MSTRG.3608 4 | 0.3788 78 | 0.0115 25 | 0.1988 6 | |

| | | | | | | | |
|---|---|---|---|---|---|---|---|
| 1248 | . | gene | MSTRG.17979 | 1.411 | 0.011587 | 0.199128 | Bo5g071170.1 |
| 1245 | . | gene | MSTRG.18492 | 1.83712 | 0.01157 | 0.199128 | Bo5g108540.1 |
| 1244 | . | gene | MSTRG.19824 | 1.94573 | 0.01156 | 0.199128 | Bo6g001000.1 |
| 1246 | . | gene | MSTRG.28037 | 1.352193 | 0.011581 | 0.199128 | Bo8g080790.1 |
| 1247 | . | gene | MSTRG.18484 | 0.145483 | 0.011583 | 0.199128 | |
| 1249 | . | gene | MSTRG.11278 | 2.611199 | 0.011607 | 0.199314 | Bo3g147870.1 |
| 1250 | . | gene | MSTRG.19714 | 1.64338 | 0.011629 | 0.199446 | Bo5g150110.1 |
| 1251 | . | gene | MSTRG.30104 | 0.407167 | 0.01164 | 0.199446 | Bo9g010450.1 |
| 1252 | . | gene | MSTRG.7144 | 1.942247 | 0.011645 | 0.199446 | |
| 1253 | . | gene | MSTRG.2287 | 0.519143 | 0.011652 | 0.199446 | |
| 1254 | . | gene | MSTRG.22954 | 0.652799 | 0.011681 | 0.199782 | Bo7g009770.1 |
| 1255 | . | gene | MSTRG.27694 | 0.648694 | 0.01173 | 0.200463 | |
| 1257 | . | gene | MSTRG.16460 | 4.204137 | 0.011752 | 0.200519 | Bo5g009310.1 |
| 1256 | . | gene | MSTRG.27835 | 0.559801 | 0.01175 | 0.200519 | Bo8g076230.1 |
| 1258 | . | gene | MSTRG.10990 | 0.407687 | 0.011776 | 0.20077 | Bo3g134480.1 |
| 1259 | . | gene | MSTRG.28062 | 0.099308 | 0.011792 | 0.200882 | Bo8g081130.1 |
| 1260 | . | gene | MSTRG.27407 | 1.704668 | 0.011813 | 0.201076 | Bo8g061320.1 |
| 1261 | . | gene | MSTRG.116 | 0.564572 | 0.011856 | 0.201549 | Bo1g003680.1 |
| 1269 | . | gene | MSTRG.2972 | 0.662207 | 0.011934 | 0.201549 | Bo1g143630.1 |
| 1270 | . | gene | MSTRG.3681 | 0.590419 | 0.011945 | 0.201549 | Bo2g011530.1 |
| 1262 | . | gene | MSTRG.4522 | 0.761709 | 0.011873 | 0.201549 | Bo2g050870.1 |
| 1266 | . | gene | MSTRG.7773 | 0.534173 | 0.011925 | 0.201549 | Bo3g021020.1 |
| 1267 | . | gene | MSTRG.10037 | 1.658804 | 0.011926 | 0.201549 | Bo3g083490.1 |
| 1264 | . | gene | MSTRG.11016 | 0.730892 | 0.01191 | 0.201549 | Bo3g135790.1 |
| 1273 | . | gene | MSTRG.21292 | 0.495374 | 0.011975 | 0.201549 | Bo6g080340.1 |
| 1265 | . | gene | MSTRG.26907 | 0.203782 | 0.011915 | 0.201549 | Bo8g036630.1 |
| 1271 | . | gene | MSTRG.26941 | 0.114489 | 0.011965 | 0.201549 | Bo8g038240.1 |
| 1263 | . | gene | MSTRG.29072 | 0.129932 | 0.011877 | 0.201549 | Bo8g104400.1 |
| 1275 | . | gene | MSTRG.32393 | 0.668647 | 0.011982 | 0.201549 | Bo9g133590.1 |

| | | | | | | | |
|---|---|---|---|---|---|---|---|
| 1274 | . | gene | MSTRG.33740 | 1.442484 | 0.011979 | 0.201549 | Bo9g181760.1 |
| 1268 | . | gene | MSTRG.34390 | 6.711374 | 0.011933 | 0.201549 | |
| 1272 | . | gene | MSTRG.19991 | 0.485239 | 0.011965 | 0.201549 | |
| 1280 | . | gene | MSTRG.5332 | 1.659289 | 0.012058 | 0.201949 | Bo2g097390.1 |
| 1281 | . | gene | MSTRG.15128 | 2.91133 | 0.01207 | 0.201949 | Bo4g165340.1 |
| 1282 | . | gene | MSTRG.25960 | 1.283989 | 0.012072 | 0.201949 | Bo7g114840.1 |
| 1277 | . | gene | MSTRG.26115 | 1.537562 | 0.01204 | 0.201949 | Bo7g117330.1 |
| 1276 | . | gene | MSTRG.4279 | 0.402677 | 0.012036 | 0.201949 | |
| 1278 | . | gene | MSTRG.4988 | 0.441212 | 0.012053 | 0.201949 | |
| 1279 | . | gene | MSTRG.30593 | 0.610284 | 0.012056 | 0.201949 | |
| 1283 | . | gene | MSTRG.26383 | 1.490877 | 0.012089 | 0.202085 | Bo8g004510.1 |
| 1288 | . | gene | MSTRG.4536 | 0.117201 | 0.012182 | 0.202192 | Bo2g051130.1 |
| 1299 | . | gene | MSTRG.6548 | 6.117539 | 0.012267 | 0.202192 | Bo2g161100.1 |
| 1304 | . | gene | MSTRG.7871 | 1.449889 | 0.012294 | 0.202192 | Bo3g022680.1 |
| 1300 | . | gene | MSTRG.8006 | 0.324489 | 0.012277 | 0.202192 | Bo3g024700.1 |
| 1286 | . | gene | MSTRG.9873 | 1.176787 | 0.012166 | 0.202192 | Bo3g076050.1 |
| 1293 | . | gene | MSTRG.10688 | 1.392839 | 0.012214 | 0.202192 | Bo3g109680.1 |
| 1292 | . | gene | MSTRG.12689 | 0.587812 | 0.012206 | 0.202192 | Bo4g026550.1 |
| 1290 | . | gene | MSTRG.17054 | 1.335641 | 0.012197 | 0.202192 | Bo5g025060.1 |
| 1289 | . | gene | MSTRG.23954 | 0.817158 | 0.012185 | 0.202192 | Bo7g061890.1 |
| 1285 | . | gene | MSTRG.26573 | 0.584116 | 0.012126 | 0.202192 | Bo8g016350.1 |
| 1291 | . | gene | MSTRG.27511 | 1.443519 | 0.012199 | 0.202192 | Bo8g065930.1 |
| 1303 | . | gene | MSTRG.29052 | 0.065039 | 0.012292 | 0.202192 | Bo8g104150.1 |
| 1301 | . | gene | MSTRG.31218 | 0.228705 | 0.012281 | 0.202192 | Bo9g059760.1 |
| 1284 | . | gene | MSTRG.32628 | 0.673764 | 0.012121 | 0.202192 | Bo9g148580.1 |
| 1296 | . | gene | MSTRG.33649 | 1.369707 | 0.012235 | 0.202192 | Bo9g177500.1 |
| 1287 | . | gene | MSTRG.35025 | 7.522182 | 0.012167 | 0.202192 | |
| 1294 | . | gene | MSTRG.35909 | 0.63361 | 0.012217 | 0.202192 | |
| 1295 | . | gene | MSTRG.35147 | 0.617385 | 0.012226 | 0.202192 | |

| | | | | | | | |
|---|---|---|---|---|---|---|---|
| 1297 | . | gene | MSTRG.34120 | 0.308854 | 0.012261 | 0.202192 | |
| 1298 | . | gene | MSTRG.33916 | 0.148204 | 0.012266 | 0.202192 | |
| 1302 | . | gene | MSTRG.3113 | 0.323299 | 0.012289 | 0.202192 | |
| 1307 | . | gene | MSTRG.5947 | 0.782195 | 0.012334 | 0.20239 | Bo2g134250.1 |
| 1306 | . | gene | MSTRG.14883 | 0.202544 | 0.012332 | 0.20239 | Bo4g151360.1 |
| 1305 | . | gene | MSTRG.19292 | 0.274969 | 0.012324 | 0.20239 | Bo5g137340.1 |
| 1308 | . | gene | MSTRG.7101 | 2.605948 | 0.012371 | 0.202845 | Bo3g007270.1 |
| 1312 | iqd17 | gene | MSTRG.9083 | 0.75713 | 0.01244 | 0.20325 | Bo3g053300.1 |
| 1313 | . | gene | MSTRG.9659 | 0.777022 | 0.012443 | 0.20325 | Bo3g066680.1 |
| 1311 | . | gene | MSTRG.16749 | 6.771205 | 0.012426 | 0.20325 | Bo5g017520.1 |
| 1309 | UCH2 | gene | MSTRG.21821 | 1.932328 | 0.012406 | 0.20325 | Bo6g099420.1 |
| 1310 | . | gene | MSTRG.33565 | 0.54265 | 0.01242 | 0.20325 | Bo9g176180.1 |
| 1314 | . | gene | MSTRG.22145 | 0.75171 | 0.012478 | 0.203658 | Bo6g112190.1 |
| 1316 | . | gene | MSTRG.2780 | 1.454617 | 0.012515 | 0.203907 | Bo1g134570.1 |
| 1315 | . | gene | MSTRG.9667 | 16.38303 | 0.012508 | 0.203907 | Bo3g066670.1 |
| 1317 | . | gene | MSTRG.15894 | 1.203629 | 0.012521 | 0.203907 | Bo4g195050.1 |
| 1318 | . | gene | MSTRG.31777 | 2.126619 | 0.012531 | 0.203912 | Bo9g094130.1 |
| 1320 | . | gene | MSTRG.3425 | 0.533645 | 0.012589 | 0.204406 | Bo2g006550.1 |
| 1324 | . | gene | MSTRG.13193 | 1.317847 | 0.012619 | 0.204406 | Bo4g049610.1 |
| 1319 | . | gene | MSTRG.24135 | 0.870517 | 0.01258 | 0.204406 | Bo7g067120.1 |
| 1321 | . | gene | MSTRG.17690 | 14.36077 | 0.012599 | 0.204406 | |
| 1322 | . | gene | MSTRG.17638 | 0.38341 | 0.012612 | 0.204406 | |
| 1323 | . | gene | MSTRG.30725 | 1.404482 | 0.012615 | 0.204406 | |
| 1325 | . | gene | MSTRG.8560 | 0.615497 | 0.012636 | 0.204536 | |
| 1326 | . | gene | MSTRG.7580 | 1.430325 | 0.012659 | 0.204751 | Bo3g017410.1 |
| 1327 | . | gene | MSTRG.33479 | 0.99724 | 0.01267 | 0.204778 | |
| 1330 | . | gene | MSTRG.11464 | 1.367571 | 0.012737 | 0.205195 | Bo3g153880.1 |
| 1334 | . | gene | MSTRG.22686 | 2.177616 | 0.012763 | 0.205195 | Bo6g123910.1 |
| 1332 | . | gene | MSTRG.22809 | 1.747386 | 0.012756 | 0.205195 | Bo7g003680.1 |

| | | | | | | | |
|---|---|---|---|---|---|---|---|
| 13<br>28 | . | gene | MSTRG.2427<br>0 | 0.7704<br>89 | 0.0127<br>21 | 0.2051<br>95 | Bo7g0754<br>40.1 |
| 13<br>33 | . | gene | MSTRG.2586<br>1 | 0.5018<br>72 | 0.0127<br>59 | 0.2051<br>95 | Bo7g1133<br>10.1 |
| 13<br>31 | . | gene | MSTRG.3095<br>2 | 0.7438<br>79 | 0.0127<br>55 | 0.2051<br>95 | Bo9g0435<br>70.1 |
| 13<br>29 | . | gene | MSTRG.3435<br>0 | 0.2098<br>5 | 0.0127<br>26 | 0.2051<br>95 | |
| 13<br>36 | . | gene | MSTRG.2092<br>3 | 0.6381<br>45 | 0.0128<br>24 | 0.2058<br>64 | Bo6g0695<br>80.1 |
| 13<br>35 | . | gene | MSTRG.1474<br>8 | 0.0862<br>79 | 0.0128<br>18 | 0.2058<br>64 | |
| 13<br>37 | . | gene | MSTRG.5832 | 1.3866<br>26 | 0.0128<br>57 | 0.2062<br>37 | Bo2g1303<br>50.1 |
| 13<br>38 | . | gene | MSTRG.2516<br>7 | 0.7147<br>07 | 0.0128<br>83 | 0.2065<br>03 | Bo7g0979<br>10.1 |
| 13<br>39 | ATMES12 | gene | MSTRG.3163<br>9 | 0.3213<br>8 | 0.0129<br>24 | 0.2070<br>12 | Bo9g0870<br>20.1 |
| 13<br>41 | . | gene | MSTRG.1618<br>6 | 1.4302<br>08 | 0.0129<br>6 | 0.2072<br>81 | Bo5g0045<br>70.1 |
| 13<br>40 | . | gene | MSTRG.2216<br>1 | 1.4930<br>73 | 0.0129<br>52 | 0.2072<br>81 | Bo6g1123<br>70.1 |
| 13<br>48 | . | gene | MSTRG.1091 | 0.1085<br>16 | 0.0130<br>8 | 0.2081<br>12 | Bo1g0306<br>50.1 |
| 13<br>44 | . | gene | MSTRG.4313 | 0.8076<br>61 | 0.0130<br>65 | 0.2081<br>12 | Bo2g0412<br>20.1 |
| 13<br>45 | . | gene | MSTRG.7632 | 0.0732<br>19 | 0.0130<br>72 | 0.2081<br>12 | Bo3g0180<br>00.1 |
| 13<br>46 | . | gene | MSTRG.2710<br>3 | 0.6404<br>74 | 0.0130<br>73 | 0.2081<br>12 | Bo8g0503<br>40.1 |
| 13<br>42 | . | gene | MSTRG.3522<br>5 | 0.2112<br>3 | 0.0130<br>32 | 0.2081<br>12 | |
| 13<br>43 | . | gene | MSTRG.3451<br>1 | 0.1997<br>14 | 0.0130<br>52 | 0.2081<br>12 | |
| 13<br>47 | . | gene | MSTRG.2307<br>3 | 1.4333<br>26 | 0.0130<br>75 | 0.2081<br>12 | |
| 13<br>49 | . | gene | MSTRG.1512<br>3 | 0.1380<br>55 | 0.0130<br>94 | 0.2081<br>78 | Bo4g1652<br>50.1 |
| 13<br>51 | . | gene | MSTRG.3188<br>9 | 1.2990<br>69 | 0.0131<br>25 | 0.2083<br>41 | Bo9g1011<br>20.1 |
| 13<br>50 | . | gene | MSTRG.2469<br>1 | 1.8502<br>03 | 0.0131<br>15 | 0.2083<br>41 | |
| 13<br>52 | . | gene | MSTRG.8237 | 0.5764<br>83 | 0.0131<br>34 | 0.2083<br>41 | |
| 13<br>53 | . | gene | MSTRG.9504 | 0.5993<br>33 | 0.0131<br>67 | 0.2087<br>17 | Bo3g0645<br>20.1 |
| 13<br>54 | . | gene | MSTRG.3457 | 2.4803<br>98 | 0.0132 | 0.2090<br>5 | Bo2g0071<br>30.1 |
| 13<br>55 | . | gene | MSTRG.3363<br>6 | 0.3491<br>87 | 0.0132<br>08 | 0.2090<br>5 | Bo9g1772<br>50.1 |
| 13<br>57 | . | gene | MSTRG.2186<br>6 | 0.6475<br>97 | 0.0132<br>36 | 0.2091<br>84 | Bo6g1012<br>20.1 |
| 13<br>56 | . | gene | MSTRG.2146 | 0.0985<br>82 | 0.0132<br>26 | 0.2091<br>84 | |
| 13<br>58 | HO2 | gene | MSTRG.1765<br>7 | 0.4446<br>38 | 0.0132<br>51 | 0.2092<br>78 | Bo5g0493<br>20.1 |
| 13<br>59 | . | gene | MSTRG.6711 | 1.4410<br>9 | 0.0132<br>99 | 0.2098<br>71 | |

| | | | | | | | |
|---|---|---|---|---|---|---|---|
| 13<br>60 | . | gene | MSTRG.1407 | 1.0949<br>31 | 0.0133<br>35 | 0.2101<br>21 | Bo1g0452<br>20.1 |
| 13<br>62 | . | gene | MSTRG.7112 | 1.4575<br>62 | 0.0133<br>51 | 0.2101<br>21 | Bo3g0074<br>40.1 |
| 13<br>64 | . | gene | MSTRG.1140<br>0 | 0.4330<br>99 | 0.0133<br>78 | 0.2101<br>21 | Bo3g1527<br>20.1 |
| 13<br>65 | . | gene | MSTRG.1519<br>4 | 0.4160<br>36 | 0.0133<br>8 | 0.2101<br>21 | Bo4g1707<br>20.1 |
| 13<br>63 | . | gene | MSTRG.3101<br>1 | 3.0496<br>48 | 0.0133<br>58 | 0.2101<br>21 | Bo9g0465<br>90.1 |
| 13<br>61 | . | gene | MSTRG.1317<br>2 | 0.0775<br>39 | 0.0133<br>48 | 0.2101<br>21 | |
| 13<br>66 | . | gene | MSTRG.1818<br>3 | 0.3439<br>66 | 0.0133<br>83 | 0.2101<br>21 | |
| 13<br>67 | . | gene | MSTRG.3176<br>5 | 47.911 | 0.0133<br>99 | 0.2102<br>1 | Bo9g0940<br>20.1 |
| 13<br>69 | . | gene | MSTRG.3771 | 0.2439<br>27 | 0.0134<br>29 | 0.2103<br>84 | Bo2g0131<br>70.1 |
| 13<br>68 | . | gene | MSTRG.3465<br>9 | 0.7765<br>65 | 0.0134<br>25 | 0.2103<br>84 | |
| 13<br>72 | . | gene | MSTRG.7662 | 1.4398<br>78 | 0.0134<br>66 | 0.2104<br>67 | Bo3g0184<br>30.1 |
| 13<br>75 | . | gene | MSTRG.1195<br>7 | 1.9937<br>08 | 0.0134<br>93 | 0.2104<br>67 | Bo3g1784<br>20.1 |
| 13<br>74 | . | gene | MSTRG.2270<br>7 | 0.7482<br>47 | 0.0134<br>85 | 0.2104<br>67 | Bo6g1241<br>40.1 |
| 13<br>70 | . | gene | MSTRG.2901<br>1 | 16.924<br>36 | 0.0134<br>48 | 0.2104<br>67 | Bo8g1024<br>20.1 |
| 13<br>71 | . | gene | MSTRG.1626<br>4 | 1.3749<br>19 | 0.0134<br>56 | 0.2104<br>67 | |
| 13<br>73 | . | gene | MSTRG.2824 | 1.4389<br>15 | 0.0134<br>84 | 0.2104<br>67 | |
| 13<br>77 | ATMKK2 | gene | MSTRG.642 | 1.4204<br>03 | 0.0135<br>48 | 0.2106<br>04 | Bo1g0149<br>40.1 |
| 13<br>91 | . | gene | MSTRG.4390 | 0.0882<br>78 | 0.0136<br>84 | 0.2106<br>04 | Bo2g0445<br>10.1 |
| 13<br>90 | . | gene | MSTRG.5142 | 0.1153<br>51 | 0.0136<br>8 | 0.2106<br>04 | Bo2g0912<br>60.1 |
| 13<br>93 | . | gene | MSTRG.8123 | 1.9618<br>09 | 0.0136<br>92 | 0.2106<br>04 | Bo3g0276<br>30.1 |
| 13<br>82 | . | gene | MSTRG.8305 | 1.3725<br>63 | 0.0135<br>95 | 0.2106<br>04 | Bo3g0319<br>50.1 |
| 13<br>89 | . | gene | MSTRG.1059<br>7 | 0.2570<br>26 | 0.0136<br>73 | 0.2106<br>04 | Bo3g1072<br>00.1 |
| 13<br>96 | . | gene | MSTRG.1139<br>2 | 0.6963<br>31 | 0.0137<br>3 | 0.2106<br>04 | Bo3g1526<br>20.1 |
| 13<br>85 | . | gene | MSTRG.1766<br>0 | 0.1289<br>15 | 0.0136<br>19 | 0.2106<br>04 | Bo5g0494<br>20.1 |
| 13<br>99 | . | gene | MSTRG.2252<br>3 | 0.5205<br>41 | 0.0137<br>38 | 0.2106<br>04 | Bo6g1206<br>10.1 |
| 13<br>84 | . | gene | MSTRG.2436<br>8 | 0.8452<br>17 | 0.0136<br>1 | 0.2106<br>04 | Bo7g0778<br>70.1 |
| 13<br>88 | . | gene | MSTRG.2783<br>7 | 1.1655<br>2 | 0.0136<br>35 | 0.2106<br>04 | Bo8g0762<br>70.1 |
| 13<br>81 | . | gene | MSTRG.2786<br>4 | 1.8455<br>27 | 0.0135<br>9 | 0.2106<br>04 | Bo8g0767<br>30.1 |
| 13<br>83 | . | gene | MSTRG.2940<br>8 | 1.4419<br>52 | 0.0136<br>06 | 0.2106<br>04 | Bo8g1095<br>70.1 |

| | | | | | | | |
|---|---|---|---|---|---|---|---|
| 1380 | . | gene | MSTRG.30325 | 3.23565 | 0.013582 | 0.210604 | Bo9g017940.1 |
| 1397 | . | gene | MSTRG.31091 | 0.79454 | 0.013731 | 0.210604 | Bo9g055670.1 |
| 1395 | . | gene | MSTRG.31350 | 0.576182 | 0.013726 | 0.210604 | Bo9g069500.1 |
| 1379 | . | gene | MSTRG.31918 | 0.241353 | 0.013572 | 0.210604 | Bo9g102520.1 |
| 1378 | . | gene | MSTRG.32027 | 0.187526 | 0.013549 | 0.210604 | Bo9g110930.1 |
| 1387 | . | gene | MSTRG.32272 | 0.321613 | 0.013633 | 0.210604 | Bo9g123700.1 |
| 1376 | . | gene | MSTRG.21482 | 0.296986 | 0.013541 | 0.210604 | |
| 1386 | . | gene | MSTRG.31374 | 0.74448 | 0.013633 | 0.210604 | |
| 1392 | . | gene | MSTRG.32485 | 2.0268 | 0.013692 | 0.210604 | |
| 1394 | . | gene | MSTRG.36055 | 0.138649 | 0.013716 | 0.210604 | |
| 1398 | . | gene | MSTRG.14872 | 0.101946 | 0.013737 | 0.210604 | |
| 1400 | . | gene | MSTRG.2166 | 2.089641 | 0.013772 | 0.210895 | Bo1g094620.1 |
| 1401 | . | gene | MSTRG.9927 | 1.267112 | 0.013776 | 0.210895 | Bo3g080070.1 |
| 1402 | . | gene | MSTRG.20276 | 1.163511 | 0.013788 | 0.21092 | Bo6g031600.1 |
| 1414 | . | gene | MSTRG.5214 | 2.052342 | 0.013913 | 0.211019 | Bo2g093360.1 |
| 1406 | . | gene | MSTRG.13488 | 0.629338 | 0.013858 | 0.211019 | Bo4g074310.1 |
| 1405 | . | gene | MSTRG.13923 | 1.596521 | 0.013848 | 0.211019 | Bo4g106490.1 |
| 1403 | . | gene | MSTRG.15925 | 0.647729 | 0.013816 | 0.211019 | Bo4g195670.1 |
| 1416 | . | gene | MSTRG.18099 | 2.785493 | 0.013932 | 0.211019 | Bo5g079840.1 |
| 1413 | . | gene | MSTRG.18673 | 1.983648 | 0.013906 | 0.211019 | Bo5g117030.1 |
| 1409 | . | gene | MSTRG.22478 | 1.625836 | 0.013872 | 0.211019 | Bo6g120010.1 |
| 1407 | . | gene | MSTRG.23499 | 1.502288 | 0.013864 | 0.211019 | Bo7g043440.1 |
| 1404 | . | gene | MSTRG.26417 | 1.534734 | 0.013833 | 0.211019 | Bo8g005990.1 |
| 1408 | . | gene | MSTRG.27968 | 5.301653 | 0.013867 | 0.211019 | Bo8g079310.1 |
| 1412 | . | gene | MSTRG.31194 | 3.045071 | 0.013902 | 0.211019 | Bo9g059370.1 |
| 1410 | . | gene | MSTRG.19773 | 0.636105 | 0.01389 | 0.211019 | |
| 1411 | . | gene | MSTRG.12491 | 1.171828 | 0.013901 | 0.211019 | |
| 1415 | . | gene | MSTRG.13270 | 1.319261 | 0.013931 | 0.211019 | |
| 1421 | . | gene | MSTRG.707 | 0.384603 | 0.013985 | 0.21104 | Bo1g016810.1 |

| | | | | | | | |
|---|---|---|---|---|---|---|---|
| 14<br>18 | . | gene | MSTRG.3460 | 1.5602<br>78 | 0.0139<br>7 | 0.2110<br>4 | Bo2g0071<br>60.1 |
| 14<br>17 | . | gene | MSTRG.1025<br>0 | 4.2598<br>98 | 0.0139<br>54 | 0.2110<br>4 | Bo3g0922<br>10.1 |
| 14<br>19 | . | gene | MSTRG.1639<br>9 | 1.2251<br>24 | 0.0139<br>76 | 0.2110<br>4 | Bo5g0086<br>00.1 |
| 14<br>23 | . | gene | MSTRG.1916<br>5 | 1.9588<br>21 | 0.0140<br>02 | 0.2110<br>4 | Bo5g1334<br>80.1 |
| 14<br>22 | . | gene | MSTRG.2429<br>2 | 0.4035<br>31 | 0.0139<br>99 | 0.2110<br>4 | Bo7g0758<br>10.1 |
| 14<br>20 | . | gene | MSTRG.1491<br>2 | 0.0655<br>07 | 0.0139<br>78 | 0.2110<br>4 | |
| 14<br>24 | . | gene | MSTRG.1105<br>4 | 8.3303<br>82 | 0.0140<br>32 | 0.2113<br>44 | Bo3g1373<br>40.1 |
| 14<br>26 | . | gene | MSTRG.3048<br>9 | 27.575<br>57 | 0.0140<br>66 | 0.2115<br>47 | Bo9g0221<br>50.1 |
| 14<br>25 | . | gene | MSTRG.5149 | 0.7481<br>89 | 0.0140<br>58 | 0.2115<br>47 | |
| 14<br>27 | . | gene | MSTRG.2278<br>0 | 3.6751<br>14 | 0.0140<br>77 | 0.2115<br>66 | Bo7g0033<br>70.1 |
| 14<br>28 | . | gene | MSTRG.3364<br>3 | 0.3695<br>46 | 0.0141<br>09 | 0.2118<br>97 | Bo9g1773<br>70.1 |
| 14<br>30 | . | gene | MSTRG.1307 | 0.3167<br>21 | 0.0142<br>18 | 0.2130<br>07 | Bo1g0397<br>70.1 |
| 14<br>31 | . | gene | MSTRG.1699<br>5 | 1.7772<br>61 | 0.0142<br>21 | 0.2130<br>07 | Bo5g0221<br>70.1 |
| 14<br>29 | . | gene | MSTRG.2929<br>9 | 1.3729<br>47 | 0.0142<br>12 | 0.2130<br>07 | Bo8g1080<br>60.1 |
| 14<br>32 | . | gene | MSTRG.2466<br>3 | 0.2811<br>18 | 0.0142<br>22 | 0.2130<br>07 | |
| 14<br>34 | ZCF61 | gene | MSTRG.1377<br>5 | 0.2702<br>1 | 0.0142<br>5 | 0.2131<br>25 | Bo4g0982<br>50.1 |
| 14<br>33 | . | gene | MSTRG.1577<br>3 | 0.7380<br>92 | 0.0142<br>41 | 0.2131<br>25 | Bo4g1908<br>90.1 |
| 14<br>35 | . | gene | MSTRG.2523<br>7 | 4.0884<br>49 | 0.0142<br>6 | 0.2131<br>25 | |
| 14<br>36 | . | gene | MSTRG.2250<br>9 | 7.9563<br>2 | 0.0142<br>75 | 0.2132<br>03 | Bo6g1203<br>80.1 |
| 14<br>37 | . | gene | MSTRG.2267<br>3 | 1.2313<br>3 | 0.0142<br>88 | 0.2132<br>46 | Bo6g1237<br>00.1 |
| 14<br>38 | . | gene | MSTRG.2166<br>1 | 159.47<br>33 | 0.0143<br>51 | 0.2140<br>41 | Bo6g0943<br>80.1 |
| 14<br>39 | . | gene | MSTRG.4178 | 0.7022<br>32 | 0.0143<br>84 | 0.2143<br>74 | Bo2g0286<br>50.1 |
| 14<br>40 | . | gene | MSTRG.2682<br>9 | 2.1752<br>51 | 0.0144 | 0.2144<br>74 | Bo8g0320<br>50.1 |
| 14<br>41 | . | gene | MSTRG.2120<br>2 | 1.3188<br>39 | 0.0144<br>18 | 0.2145<br>86 | |
| 14<br>42 | . | gene | MSTRG.2633<br>2 | 1.4887<br>37 | 0.0144<br>49 | 0.2149<br>06 | Bo8g0027<br>20.1 |
| 14<br>43 | . | gene | MSTRG.2927 | 1.4191<br>73 | 0.0144<br>63 | 0.2149<br>6 | Bo1g1410<br>00.1 |
| 14<br>44 | . | gene | MSTRG.4236 | 0.6972<br>37 | 0.0144<br>75 | 0.2149<br>83 | Bo2g0326<br>40.1 |
| 14<br>45 | . | gene | MSTRG.3566<br>4 | 1.5426<br>58 | 0.0145<br>03 | 0.2151<br>02 | |
| 14<br>46 | . | gene | MSTRG.3524<br>0 | 21.956<br>96 | 0.0145<br>11 | 0.2151<br>02 | |

| | | | | | | | |
|---|---|---|---|---|---|---|---|
| 14 47 | . | gene | MSTRG.1758 9 | 0.9430 92 | 0.0145 19 | 0.2151 02 | |
| 14 48 | . | gene | MSTRG.2962 2 | 0.7420 34 | 0.0145 23 | 0.2151 02 | |
| 14 49 | . | gene | MSTRG.2258 9 | 3.5918 42 | 0.0145 62 | 0.2155 38 | Bo6g1213 80.1 |
| 14 51 | . | gene | MSTRG.9963 | 0.4119 44 | 0.0145 9 | 0.2156 48 | Bo3g0816 00.1 |
| 14 50 | . | gene | MSTRG.1890 9 | 0.6422 34 | 0.0145 87 | 0.2156 48 | Bo5g1267 80.1 |
| 14 52 | . | gene | MSTRG.1013 6 | 0.6966 18 | 0.0146 11 | 0.2158 18 | Bo3g0869 70.1 |
| 14 61 | . | gene | MSTRG.9872 | 2.0175 02 | 0.0147 11 | 0.2159 49 | Bo3g0760 40.1 |
| 14 59 | . | gene | MSTRG.1245 7 | 0.4336 04 | 0.0146 96 | 0.2159 49 | Bo4g0211 30.1 |
| 14 55 | . | gene | MSTRG.1584 1 | 3.8437 2 | 0.0146 61 | 0.2159 49 | Bo4g1930 00.1 |
| 14 54 | . | gene | MSTRG.2004 6 | 0.5953 84 | 0.0146 58 | 0.2159 49 | Bo6g0209 60.1 |
| 14 56 | . | gene | MSTRG.2100 9 | 3.2122 16 | 0.0146 7 | 0.2159 49 | Bo6g0730 30.1 |
| 14 57 | . | gene | MSTRG.2980 8 | 0.5641 04 | 0.0146 91 | 0.2159 49 | Bo9g0040 20.1 |
| 14 53 | . | gene | MSTRG.2242 8 | 0.6256 34 | 0.0146 33 | 0.2159 49 | |
| 14 58 | . | gene | MSTRG.2775 | 0.6471 4 | 0.0146 94 | 0.2159 49 | |
| 14 60 | . | gene | MSTRG.3490 4 | 2.0207 | 0.0147 05 | 0.2159 49 | |
| 14 85 | . | gene | MSTRG.7259 | 0.5915 87 | 0.0149 96 | 0.2165 82 | Bo3g0101 70.1 |
| 14 82 | . | gene | MSTRG.9497 | 0.0830 68 | 0.0149 68 | 0.2165 82 | Bo3g0644 20.1 |
| 14 64 | . | gene | MSTRG.1121 0 | 2.1300 25 | 0.0148 2 | 0.2165 82 | Bo3g1438 30.1 |
| 14 83 | . | gene | MSTRG.1239 2 | 1.3063 56 | 0.0149 77 | 0.2165 82 | Bo4g0173 80.1 |
| 14 68 | . | gene | MSTRG.1254 5 | 1.5602 87 | 0.0148 65 | 0.2165 82 | Bo4g0244 40.1 |
| 14 73 | . | gene | MSTRG.1749 1 | 3.2192 02 | 0.0149 13 | 0.2165 82 | Bo5g0427 10.1 |
| 14 65 | . | gene | MSTRG.1755 8 | 0.6440 84 | 0.0148 42 | 0.2165 82 | Bo5g0438 60.1 |
| 14 80 | . | gene | MSTRG.2307 9 | 4.0507 97 | 0.0149 53 | 0.2165 82 | Bo7g0162 00.1 |
| 14 66 | . | gene | MSTRG.2419 9 | 0.2906 59 | 0.0148 46 | 0.2165 82 | Bo7g0712 70.1 |
| 14 81 | . | gene | MSTRG.2562 2 | 1.8348 45 | 0.0149 63 | 0.2165 82 | Bo7g1085 80.1 |
| 14 67 | . | gene | MSTRG.2669 3 | 1.4961 11 | 0.0148 59 | 0.2165 82 | Bo8g0256 20.1 |
| 14 74 | . | gene | MSTRG.2818 8 | 2.3926 87 | 0.0149 15 | 0.2165 82 | Bo8g0830 70.1 |
| 14 71 | . | gene | MSTRG.2825 2 | 1.7296 88 | 0.0148 9 | 0.2165 82 | Bo8g0869 40.1 |
| 14 62 | . | gene | MSTRG.3065 7 | 0.2099 86 | 0.0147 8 | 0.2165 82 | Bo9g0280 90.1 |

| | | | | | | | |
|---|---|---|---|---|---|---|---|
| 14 75 | . | gene | MSTRG.3075 5 | 1.4351 75 | 0.0149 25 | 0.2165 82 | Bo9g0306 80.1 |
| 14 79 | EMB1706 | gene | MSTRG.3167 2 | 1.1887 14 | 0.0149 42 | 0.2165 82 | Bo9g0896 40.1 |
| 14 84 | . | gene | MSTRG.3238 5 | 1.7639 59 | 0.0149 88 | 0.2165 82 | Bo9g1335 10.1 |
| 14 78 | . | gene | MSTRG.3370 2 | 0.5917 17 | 0.0149 39 | 0.2165 82 | Bo9g1812 70.1 |
| 14 63 | . | gene | MSTRG.2070 9 | 0.4147 52 | 0.0148 01 | 0.2165 82 | |
| 14 69 | . | gene | MSTRG.3389 8 | 0.5515 99 | 0.0148 7 | 0.2165 82 | |
| 14 70 | . | gene | MSTRG.1625 4 | 1.3692 | 0.0148 74 | 0.2165 82 | |
| 14 72 | . | gene | MSTRG.2149 | 1.2405 6 | 0.0149 04 | 0.2165 82 | |
| 14 76 | . | gene | MSTRG.2052 5 | 0.5741 8 | 0.0149 28 | 0.2165 82 | |
| 14 77 | . | gene | MSTRG.1739 5 | 0.8645 48 | 0.0149 38 | 0.2165 82 | |
| 14 86 | . | gene | MSTRG.3148 4 | 0.4496 37 | 0.0150 28 | 0.2168 31 | |
| 14 87 | . | gene | MSTRG.3371 2 | 0.7702 24 | 0.0150 34 | 0.2168 31 | |
| 14 88 | . | gene | MSTRG.1813 1 | 0.0951 47 | 0.0150 59 | 0.2170 55 | |
| 14 89 | . | gene | MSTRG.2683 | 1.5576 57 | 0.0150 85 | 0.2172 79 | Bo1g1249 80.1 |
| 14 90 | . | gene | MSTRG.1954 3 | 1.4098 61 | 0.0151 02 | 0.2173 77 | Bo5g1435 30.1 |
| 14 91 | . | gene | MSTRG.3820 | 1.9117 23 | 0.0151 28 | 0.2176 02 | Bo2g0140 90.1 |
| 14 93 | . | gene | MSTRG.1514 9 | 0.2618 05 | 0.0151 56 | 0.2177 18 | Bo4g1679 10.1 |
| 14 92 | . | gene | MSTRG.2823 6 | 1.3902 72 | 0.0151 48 | 0.2177 18 | Bo8g0837 40.1 |
| 14 95 | . | gene | MSTRG.2112 9 | 1.2951 89 | 0.0152 06 | 0.2181 39 | Bo6g0767 60.1 |
| 14 94 | . | gene | MSTRG.5226 | 0.6429 16 | 0.0152 | 0.2181 39 | |
| 14 96 | . | gene | MSTRG.7232 | 1.4347 1 | 0.0152 2 | 0.2182 01 | |
| 14 97 | . | gene | MSTRG.1997 7 | 1.1378 | 0.0152 48 | 0.2184 59 | Bo6g0157 80.1 |
| 15 00 | . | gene | MSTRG.9704 | 1.3111 84 | 0.0153 01 | 0.2186 49 | Bo3g0685 30.1 |
| 14 99 | . | gene | MSTRG.1126 9 | 4.8011 72 | 0.0152 9 | 0.2186 49 | Bo3g1476 70.1 |
| 15 03 | . | gene | MSTRG.1394 5 | 2.1656 26 | 0.0153 27 | 0.2186 49 | Bo4g1079 40.1 |
| 15 02 | . | gene | MSTRG.2234 7 | 1.4342 55 | 0.0153 19 | 0.2186 49 | Bo6g1181 90.1 |
| 15 01 | . | gene | MSTRG.3077 2 | 1.5295 36 | 0.0153 12 | 0.2186 49 | Bo9g0310 00.1 |
| 14 98 | . | gene | MSTRG.1370 6 | 0.3304 72 | 0.0152 84 | 0.2186 49 | |
| 15 04 | . | gene | MSTRG.2238 4 | 0.5839 89 | 0.0153 33 | 0.2186 49 | |

| | | | | | | | |
|---|---|---|---|---|---|---|---|
| 1505 | . | gene | MSTRG.35516 | 2.51952 | 0.015365 | 0.218956 | |
| 1506 | . | gene | MSTRG.21933 | 0.779696 | 0.01539 | 0.219035 | Bo6g104080.1 |
| 1507 | . | gene | MSTRG.26382 | 1.255682 | 0.015399 | 0.219035 | Bo8g004450.1 |
| 1508 | . | gene | MSTRG.10852 | 0.492772 | 0.015401 | 0.219035 | |
| 1510 | . | gene | MSTRG.7030 | 1.452867 | 0.015434 | 0.219177 | Bo3g005340.1 |
| 1509 | . | gene | MSTRG.25877 | 0.853795 | 0.015423 | 0.219177 | Bo7g113530.1 |
| 1511 | emb1129 | gene | MSTRG.26517 | 1.349768 | 0.015442 | 0.219177 | Bo8g011670.1 |
| 1512 | . | gene | MSTRG.17000 | 1.177827 | 0.015467 | 0.21939 | Bo5g022220.1 |
| 1513 | . | gene | MSTRG.2077 | 0.582067 | 0.015498 | 0.219591 | Bo1g087940.1 |
| 1514 | . | gene | MSTRG.15952 | 0.970399 | 0.015505 | 0.219591 | Bo4g195950.1 |
| 1515 | . | gene | MSTRG.26972 | 0.338611 | 0.015512 | 0.219591 | |
| 1516 | . | gene | MSTRG.7846 | 1.37768 | 0.015536 | 0.219795 | Bo3g022050.1 |
| 1517 | . | gene | MSTRG.9393 | 1.689818 | 0.015559 | 0.219963 | Bo3g061850.1 |
| 1518 | . | gene | MSTRG.11436 | 0.172613 | 0.015586 | 0.220207 | Bo3g153530.1 |
| 1519 | . | gene | MSTRG.28944 | 2.124982 | 0.015602 | 0.220284 | Bo8g101550.1 |
| 1520 | . | gene | MSTRG.31427 | 1.366582 | 0.015621 | 0.220409 | Bo9g073120.1 |
| 1526 | . | gene | MSTRG.1766 | 1.419017 | 0.015691 | 0.220455 | Bo1g059170.1 |
| 1525 | . | gene | MSTRG.7135 | 0.675116 | 0.015689 | 0.220455 | Bo3g008770.1 |
| 1527 | . | gene | MSTRG.13632 | 1.65931 | 0.015696 | 0.220455 | Bo4g085100.1 |
| 1522 | . | gene | MSTRG.21243 | 2.040214 | 0.015659 | 0.220455 | Bo6g079410.1 |
| 1521 | . | gene | MSTRG.34017 | 0.40137 | 0.015657 | 0.220455 | |
| 1523 | . | gene | MSTRG.15284 | 0.413538 | 0.015676 | 0.220455 | |
| 1524 | . | gene | MSTRG.22381 | 0.661524 | 0.015681 | 0.220455 | |
| 1528 | . | gene | MSTRG.18361 | 1.456307 | 0.015707 | 0.220459 | Bo5g098230.1 |
| 1534 | . | gene | MSTRG.1408 | 0.438645 | 0.015788 | 0.22051 | Bo1g045240.1 |
| 1537 | . | gene | MSTRG.11880 | 0.597824 | 0.015815 | 0.22051 | Bo3g175330.1 |
| 1536 | . | gene | MSTRG.16646 | 1.350587 | 0.015809 | 0.22051 | Bo5g014800.1 |
| 1530 | . | gene | MSTRG.18070 | 5.604772 | 0.015731 | 0.22051 | Bo5g076460.1 |
| 1542 | . | gene | MSTRG.19102 | 1.37348 | 0.015854 | 0.22051 | Bo5g132520.1 |

| | | | | | | | |
|---|---|---|---|---|---|---|---|
| 1535 | . | gene | MSTRG.21096 | 0.661992 | 0.015805 | 0.22051 | Bo6g076230.1 |
| 1540 | . | gene | MSTRG.22360 | 0.246999 | 0.015846 | 0.22051 | Bo6g118450.1 |
| 1531 | . | gene | MSTRG.26585 | 1.619382 | 0.015758 | 0.22051 | Bo8g016660.1 |
| 1529 | . | gene | MSTRG.26847 | 0.084704 | 0.015723 | 0.22051 | Bo8g032430.1 |
| 1539 | . | gene | MSTRG.27780 | 2.049498 | 0.015842 | 0.22051 | Bo8g071310.1 |
| 1532 | . | gene | MSTRG.2367 | 0.363787 | 0.015764 | 0.22051 | |
| 1533 | . | gene | MSTRG.26363 | 0.201914 | 0.01577 | 0.22051 | |
| 1538 | . | gene | MSTRG.19450 | 1.511749 | 0.015825 | 0.22051 | |
| 1541 | . | gene | MSTRG.34342 | 1.736768 | 0.015846 | 0.22051 | |
| 1543 | . | gene | MSTRG.2060 | 0.748425 | 0.015915 | 0.221125 | Bo1g087660.1 |
| 1544 | . | gene | MSTRG.3544 | 1.211509 | 0.015927 | 0.221125 | Bo2g009580.1 |
| 1545 | ATERDJ3A | gene | MSTRG.19381 | 0.220221 | 0.015929 | 0.221125 | Bo5g139610.1 |
| 1548 | . | gene | MSTRG.6636 | 0.064455 | 0.016 | 0.221566 | Bo2g165560.1 |
| 1547 | . | gene | MSTRG.17516 | 1.825124 | 0.015986 | 0.221566 | Bo5g043090.1 |
| 1546 | . | gene | MSTRG.17231 | 0.234234 | 0.015984 | 0.221566 | |
| 1549 | . | gene | MSTRG.35223 | 0.763881 | 0.016003 | 0.221566 | |
| 1550 | . | gene | MSTRG.32684 | 0.714127 | 0.016013 | 0.221575 | |
| 1551 | . | gene | MSTRG.2676 | 0.300546 | 0.01603 | 0.221588 | Bo1g123670.1 |
| 1553 | . | gene | MSTRG.8107 | 1.908701 | 0.016045 | 0.221588 | Bo3g027270.1 |
| 1552 | . | gene | MSTRG.23406 | 0.068486 | 0.016045 | 0.221588 | Bo7g038880.1 |
| 1554 | . | gene | MSTRG.25033 | 1.163863 | 0.016073 | 0.221824 | Bo7g095870.1 |
| 1555 | . | gene | MSTRG.21289 | 2.766463 | 0.016134 | 0.222531 | Bo6g080300.1 |
| 1556 | . | gene | MSTRG.17958 | 3.661155 | 0.016147 | 0.22254 | Bo5g070670.1 |
| 1557 | . | gene | MSTRG.25772 | 2.63516 | 0.016156 | 0.22254 | Bo7g110950.1 |
| 1559 | . | gene | MSTRG.5219 | 0.7718 | 0.016179 | 0.222568 | Bo2g093430.1 |
| 1558 | . | gene | MSTRG.12063 | 1.542344 | 0.016176 | 0.222568 | Bo3g184270.1 |
| 1563 | . | gene | MSTRG.17153 | 0.365528 | 0.016223 | 0.222601 | Bo5g027750.1 |
| 1561 | . | gene | MSTRG.23025 | 1.435678 | 0.016215 | 0.222601 | Bo7g014080.1 |
| 1562 | . | gene | MSTRG.29909 | 15.78126 | 0.016218 | 0.222601 | Bo9g006220.1 |

| | | | | | | | |
|---|---|---|---|---|---|---|---|
| 1560 | . | gene | MSTRG.29556 | 2.10611 | 0.016201 | 0.222601 | |
| 1564 | . | gene | MSTRG.16372 | 9.637451 | 0.016261 | 0.22298 | Bo5g008190.1 |
| 1567 | . | gene | MSTRG.7543 | 0.054609 | 0.016295 | 0.223022 | Bo3g016930.1 |
| 1566 | . | gene | MSTRG.18961 | 2.057817 | 0.016289 | 0.223022 | Bo5g127510.1 |
| 1565 | . | gene | MSTRG.29190 | 1.237205 | 0.016278 | 0.223022 | Bo8g106220.1 |
| 1568 | . | gene | MSTRG.34651 | 3.076887 | 0.016315 | 0.223153 | |
| 1569 | . | gene | MSTRG.17603 | 0.206238 | 0.016332 | 0.223247 | Bo5g048540.1 |
| 1570 | . | gene | MSTRG.6666 | 0.36593 | 0.016367 | 0.223588 | |
| 1571 | . | gene | MSTRG.22461 | 0.696228 | 0.016394 | 0.223691 | Bo6g119810.1 |
| 1572 | . | gene | MSTRG.25794 | 0.58587 | 0.016396 | 0.223691 | |
| 1574 | . | gene | MSTRG.18942 | 0.684101 | 0.01644 | 0.223982 | Bo5g127260.1 |
| 1575 | . | gene | MSTRG.30552 | 0.652986 | 0.016449 | 0.223982 | Bo9g024160.1 |
| 1573 | . | gene | MSTRG.10016 | 0.235103 | 0.016428 | 0.223982 | |
| 1578 | . | gene | MSTRG.1434 | 0.761554 | 0.016504 | 0.224191 | Bo1g046610.1 |
| 1580 | . | gene | MSTRG.15886 | 0.677507 | 0.016517 | 0.224191 | Bo4g194860.1 |
| 1577 | . | gene | MSTRG.15974 | 3.405757 | 0.016495 | 0.224191 | Bo4g197240.1 |
| 1581 | . | gene | MSTRG.33101 | 2.931548 | 0.016527 | 0.224191 | Bo9g165660.1 |
| 1576 | . | gene | MSTRG.35012 | 2.388727 | 0.016482 | 0.224191 | |
| 1579 | . | gene | MSTRG.32156 | 0.71419 | 0.01651 | 0.224191 | |
| 1588 | . | gene | MSTRG.1642 | 0.970409 | 0.016641 | 0.224604 | Bo1g055090.1 |
| 1583 | . | gene | MSTRG.9763 | 0.559157 | 0.016622 | 0.224604 | Bo3g071430.1 |
| 1589 | . | gene | MSTRG.11624 | 0.975682 | 0.016644 | 0.224604 | Bo3g163990.1 |
| 1587 | . | gene | MSTRG.11802 | 0.08458 | 0.016635 | 0.224604 | Bo3g171190.1 |
| 1585 | EMB1705 | gene | MSTRG.13242 | 1.280687 | 0.016626 | 0.224604 | Bo4g052420.1 |
| 1584 | . | gene | MSTRG.16633 | 0.543321 | 0.016624 | 0.224604 | Bo5g014690.1 |
| 1582 | . | gene | MSTRG.23953 | 2.726195 | 0.01658 | 0.224604 | |
| 1586 | . | gene | MSTRG.26891 | 0.267933 | 0.016627 | 0.224604 | |
| 1590 | . | gene | MSTRG.11653 | 0.907679 | 0.016651 | 0.224604 | |
| 1606 | . | gene | MSTRG.397 | 0.485967 | 0.01687 | 0.224629 | Bo1g008300.1 |

| | | | | | | | |
|---|---|---|---|---|---|---|---|
| 15 92 | . | gene | MSTRG.544 | 1.2885 95 | 0.0166 93 | 0.2246 29 | Bo1g0114 10.1 |
| 15 99 | . | gene | MSTRG.850 | 1.1517 61 | 0.0167 8 | 0.2246 29 | Bo1g0199 60.1 |
| 15 96 | . | gene | MSTRG.1420 | 15.835 75 | 0.0167 47 | 0.2246 29 | Bo1g0464 50.1 |
| 16 47 | . | gene | MSTRG.1589 | 0.3826 5 | 0.0176 05 | 0.2246 29 | Bo1g0532 40.1 |
| 17 11 | . | gene | MSTRG.2315 | 0.2114 99 | 0.0181 28 | 0.2246 29 | Bo1g1035 20.1 |
| 16 75 | . | gene | MSTRG.2981 | 1.8140 93 | 0.0179 73 | 0.2246 29 | Bo1g1438 10.1 |
| 17 18 | . | gene | MSTRG.3056 | 0.5094 42 | 0.0181 28 | 0.2246 29 | Bo1g1449 90.1 |
| 16 60 | . | gene | MSTRG.3546 | 1.4123 17 | 0.0177 49 | 0.2246 29 | Bo2g0096 00.1 |
| 16 59 | . | gene | MSTRG.3686 | 0.6346 68 | 0.0177 39 | 0.2246 29 | Bo2g0116 10.1 |
| 16 22 | . | gene | MSTRG.3703 | 1.3924 54 | 0.0171 35 | 0.2246 29 | Bo2g0119 90.1 |
| 17 24 | . | gene | MSTRG.3717 | 0.2432 08 | 0.0181 28 | 0.2246 29 | Bo2g0122 70.1 |
| 17 25 | . | gene | MSTRG.4694 | 0.9974 61 | 0.0181 28 | 0.2246 29 | Bo2g0576 40.1 |
| 16 35 | . | gene | MSTRG.5248 | 1.5153 46 | 0.0173 42 | 0.2246 29 | Bo2g0948 80.1 |
| 16 25 | . | gene | MSTRG.5990 | 3.0495 77 | 0.0171 85 | 0.2246 29 | Bo2g1361 80.1 |
| 16 90 | . | gene | MSTRG.6139 | 0.9934 34 | 0.0181 28 | 0.2246 29 | Bo2g1427 60.1 |
| 17 26 | . | gene | MSTRG.6433 | 0.9949 06 | 0.0181 28 | 0.2246 29 | Bo2g1562 30.1 |
| 16 62 | . | gene | MSTRG.6972 | 1.8462 12 | 0.0177 64 | 0.2246 29 | Bo3g0046 00.1 |
| 15 98 | . | gene | MSTRG.7208 | 0.6160 64 | 0.0167 76 | 0.2246 29 | Bo3g0096 70.1 |
| 16 30 | ADF4 | gene | MSTRG.7603 | 1.5322 37 | 0.0172 62 | 0.2246 29 | Bo3g0176 70.1 |
| 16 91 | . | gene | MSTRG.8164 | 0.2079 96 | 0.0181 28 | 0.2246 29 | Bo3g0280 20.1 |
| 15 93 | . | gene | MSTRG.8351 | 0.8223 43 | 0.0166 97 | 0.2246 29 | Bo3g0326 10.1 |
| 15 97 | . | gene | MSTRG.8529 | 1.2382 | 0.0167 52 | 0.2246 29 | Bo3g0362 10.1 |
| 16 43 | . | gene | MSTRG.8611 | 0.5465 15 | 0.0175 65 | 0.2246 29 | Bo3g0374 30.1 |
| 17 28 | . | gene | MSTRG.8789 | 0.4385 37 | 0.0181 28 | 0.2246 29 | Bo3g0422 10.1 |
| 17 29 | . | gene | MSTRG.9117 | 0.9967 73 | 0.0181 28 | 0.2246 29 | Bo3g0546 90.1 |
| 16 27 | . | gene | MSTRG.9467 | 1.4915 26 | 0.0172 31 | 0.2246 29 | Bo3g0640 20.1 |
| 16 02 | . | gene | MSTRG.9842 | 1.2630 22 | 0.0168 18 | 0.2246 29 | Bo3g0746 40.1 |
| 16 41 | . | gene | MSTRG.1013 3 | 1.2959 05 | 0.0174 82 | 0.2246 29 | Bo3g0868 90.1 |
| 16 92 | . | gene | MSTRG.1022 1 | 0.9890 57 | 0.0181 28 | 0.2246 29 | Bo3g0914 30.1 |

| | | | | | | | |
|---|---|---|---|---|---|---|---|
| 1693 | . | gene | MSTRG.10919 | 0.984467 | 0.018128 | 0.224629 | Bo3g130190.1 |
| 1617 | . | gene | MSTRG.11502 | 0.540984 | 0.017047 | 0.224629 | Bo3g155550.1 |
| 1679 | . | gene | MSTRG.11785 | 1.992051 | 0.018049 | 0.224629 | Bo3g170960.1 |
| 1637 | . | gene | MSTRG.11830 | 0.141668 | 0.017385 | 0.224629 | Bo3g173650.1 |
| 1607 | . | gene | MSTRG.11844 | 1.250698 | 0.016893 | 0.224629 | Bo3g173810.1 |
| 1642 | . | gene | MSTRG.11873 | 0.138513 | 0.017511 | 0.224629 | Bo3g175260.1 |
| 1694 | . | gene | MSTRG.11924 | 0.26405 | 0.018128 | 0.224629 | Bo3g176080.1 |
| 1594 | . | gene | MSTRG.12145 | 0.709477 | 0.016717 | 0.224629 | Bo3g185470.1 |
| 1644 | . | gene | MSTRG.12297 | 0.468512 | 0.017579 | 0.224629 | Bo4g012670.1 |
| 1674 | . | gene | MSTRG.12405 | 0.934457 | 0.017954 | 0.224629 | Bo4g018500.1 |
| 1696 | . | gene | MSTRG.12651 | 0.991988 | 0.018128 | 0.224629 | Bo4g025870.1 |
| 1697 | . | gene | MSTRG.12701 | 0.961203 | 0.018128 | 0.224629 | Bo4g026740.1 |
| 1698 | . | gene | MSTRG.12803 | 0.996581 | 0.018128 | 0.224629 | Bo4g030680.1 |
| 1700 | . | gene | MSTRG.13096 | 0.98626 | 0.018128 | 0.224629 | Bo4g044100.1 |
| 1608 | . | gene | MSTRG.13181 | 0.159836 | 0.016897 | 0.224629 | Bo4g048420.1 |
| 1683 | . | gene | MSTRG.13218 | 0.946622 | 0.018128 | 0.224629 | Bo4g052130.1 |
| 1604 | . | gene | MSTRG.13258 | 4.906723 | 0.016827 | 0.224629 | Bo4g052640.1 |
| 1681 | . | gene | MSTRG.14171 | 2.296086 | 0.018113 | 0.224629 | Bo4g119930.1 |
| 1731 | . | gene | MSTRG.14216 | 1.825376 | 0.01813 | 0.224629 | Bo4g120620.1 |
| 1600 | . | gene | MSTRG.15332 | 9.556897 | 0.016793 | 0.224629 | Bo4g177980.1 |
| 1702 | . | gene | MSTRG.15414 | 0.705381 | 0.018128 | 0.224629 | Bo4g182290.1 |
| 1671 | . | gene | MSTRG.16243 | 1.689218 | 0.017938 | 0.224629 | Bo5g005300.1 |
| 1673 | . | gene | MSTRG.16249 | 0.283676 | 0.017946 | 0.224629 | Bo5g005370.1 |
| 1614 | . | gene | MSTRG.16280 | 0.096378 | 0.01697 | 0.224629 | Bo5g006920.1 |
| 1664 | . | gene | MSTRG.16338 | 4.337957 | 0.017787 | 0.224629 | Bo5g007710.1 |
| 1638 | . | gene | MSTRG.16416 | 0.811032 | 0.017391 | 0.224629 | Bo5g008830.1 |
| 1676 | . | gene | MSTRG.16938 | 1.978904 | 0.017988 | 0.224629 | Bo5g021470.1 |
| 1704 | . | gene | MSTRG.17039 | 0.924893 | 0.018128 | 0.224629 | Bo5g023780.1 |
| 1611 | . | gene | MSTRG.17173 | 2.160348 | 0.016918 | 0.224629 | Bo5g029120.1 |

| | | | | | | | |
|---|---|---|---|---|---|---|---|
| 1656 | . | gene | MSTRG.17613 | 0.40374 | 0.017727 | 0.224629 | Bo5g048700.1 |
| 1619 | . | gene | MSTRG.18603 | 1.396075 | 0.017096 | 0.224629 | Bo5g114080.1 |
| 1632 | . | gene | MSTRG.18618 | 3.401157 | 0.017307 | 0.224629 | Bo5g115220.1 |
| 1616 | . | gene | MSTRG.18662 | 0.508181 | 0.016996 | 0.224629 | Bo5g116870.1 |
| 1636 | . | gene | MSTRG.19734 | 0.793466 | 0.017347 | 0.224629 | Bo5g151410.1 |
| 1707 | . | gene | MSTRG.19930 | 0.989572 | 0.018128 | 0.224629 | Bo6g010940.1 |
| 1634 | . | gene | MSTRG.20026 | 1.384803 | 0.017342 | 0.224629 | Bo6g018660.1 |
| 1708 | . | gene | MSTRG.20205 | 0.996051 | 0.018128 | 0.224629 | Bo6g029490.1 |
| 1685 | . | gene | MSTRG.20240 | 0.447049 | 0.018128 | 0.224629 | Bo6g031110.1 |
| 1640 | . | gene | MSTRG.20920 | 0.443196 | 0.017443 | 0.224629 | Bo6g068550.1 |
| 1613 | . | gene | MSTRG.21682 | 12.54903 | 0.016968 | 0.224629 | Bo6g094680.1 |
| 1653 | . | gene | MSTRG.21863 | 0.671687 | 0.017677 | 0.224629 | Bo6g101160.1 |
| 1655 | . | gene | MSTRG.22133 | 0.399146 | 0.017698 | 0.224629 | Bo6g111010.1 |
| 1665 | ATY1 | gene | MSTRG.22541 | 1.290667 | 0.017815 | 0.224629 | Bo6g120810.1 |
| 1610 | . | gene | MSTRG.22751 | 0.579156 | 0.016916 | 0.224629 | Bo6g125750.1 |
| 1658 | . | gene | MSTRG.22978 | 0.363514 | 0.017736 | 0.224629 | Bo7g011310.1 |
| 1651 | . | gene | MSTRG.23838 | 1.703219 | 0.01767 | 0.224629 | Bo7g057750.1 |
| 1624 | . | gene | MSTRG.24470 | 1.484483 | 0.017173 | 0.224629 | Bo7g080500.1 |
| 1713 | . | gene | MSTRG.24630 | 0.125493 | 0.018128 | 0.224629 | Bo7g084180.1 |
| 1649 | . | gene | MSTRG.25168 | 1.367467 | 0.017654 | 0.224629 | Bo7g097930.1 |
| 1626 | . | gene | MSTRG.25370 | 0.339885 | 0.017208 | 0.224629 | Bo7g104140.1 |
| 1670 | . | gene | MSTRG.25487 | 0.494276 | 0.017936 | 0.224629 | Bo7g106010.1 |
| 1714 | . | gene | MSTRG.25875 | 0.994079 | 0.018128 | 0.224629 | Bo7g113500.1 |
| 1612 | . | gene | MSTRG.26233 | 0.503632 | 0.016925 | 0.224629 | Bo7g119080.1 |
| 1680 | . | gene | MSTRG.26261 | 0.659965 | 0.018099 | 0.224629 | Bo7g119480.1 |
| 1678 | . | gene | MSTRG.26488 | 1.102215 | 0.018047 | 0.224629 | Bo8g010990.1 |
| 1668 | . | gene | MSTRG.27406 | 1.512754 | 0.017923 | 0.224629 | Bo8g061350.1 |
| 1716 | . | gene | MSTRG.27777 | 0.960992 | 0.018128 | 0.224629 | Bo8g071230.1 |
| 1623 | . | gene | MSTRG.28220 | 0.667719 | 0.017159 | 0.224629 | Bo8g083510.1 |

| | | | | | | | |
|---|---|---|---|---|---|---|---|
| 1717 | OTP84 | gene | MSTRG.28463 | 0.030688 | 0.018128 | 0.224629 | Bo8g091830.1 |
| 1633 | . | gene | MSTRG.29802 | 0.670394 | 0.01733 | 0.224629 | Bo9g003970.1 |
| 1628 | . | gene | MSTRG.30401 | 1.401581 | 0.017256 | 0.224629 | Bo9g019110.1 |
| 1666 | . | gene | MSTRG.31197 | 1.570485 | 0.01786 | 0.224629 | Bo9g059400.1 |
| 1719 | . | gene | MSTRG.31512 | 0.219201 | 0.018128 | 0.224629 | Bo9g076570.1 |
| 1657 | . | gene | MSTRG.31522 | 0.761782 | 0.017732 | 0.224629 | Bo9g077760.1 |
| 1645 | . | gene | MSTRG.32165 | 1.655274 | 0.017581 | 0.224629 | Bo9g120200.1 |
| 1595 | . | gene | MSTRG.33289 | 2.166638 | 0.016733 | 0.224629 | Bo9g171450.1 |
| 1591 | . | gene | MSTRG.5841 | 0.085563 | 0.016678 | 0.224629 | |
| 1601 | . | gene | MSTRG.5999 | 0.124772 | 0.016798 | 0.224629 | |
| 1603 | . | gene | MSTRG.34086 | 0.530424 | 0.016819 | 0.224629 | |
| 1605 | . | gene | MSTRG.32254 | 0.477891 | 0.016869 | 0.224629 | |
| 1609 | . | gene | MSTRG.15871 | 1.499312 | 0.016901 | 0.224629 | |
| 1615 | . | gene | MSTRG.9578 | 1.251377 | 0.016991 | 0.224629 | |
| 1618 | . | gene | MSTRG.21934 | 0.373534 | 0.017077 | 0.224629 | |
| 1620 | . | gene | MSTRG.34788 | 13.03412 | 0.017105 | 0.224629 | |
| 1621 | . | gene | MSTRG.21861 | 0.921565 | 0.017116 | 0.224629 | |
| 1629 | . | gene | MSTRG.8414 | 0.59857 | 0.017258 | 0.224629 | |
| 1631 | . | gene | MSTRG.33766 | 2.573405 | 0.017282 | 0.224629 | |
| 1639 | . | gene | MSTRG.10246 | 2.662065 | 0.017441 | 0.224629 | |
| 1646 | . | gene | MSTRG.34176 | 0.831351 | 0.017591 | 0.224629 | |
| 1648 | . | gene | MSTRG.35379 | 1.212722 | 0.017621 | 0.224629 | |
| 1650 | . | gene | MSTRG.35749 | 0.266192 | 0.017667 | 0.224629 | |
| 1652 | . | gene | MSTRG.2625 | 0.199626 | 0.017671 | 0.224629 | |
| 1654 | . | gene | MSTRG.15487 | 0.160945 | 0.017696 | 0.224629 | |
| 1661 | . | gene | MSTRG.13849 | 0.764244 | 0.017755 | 0.224629 | |
| 1663 | . | gene | MSTRG.35938 | 0.514808 | 0.017782 | 0.224629 | |
| 1667 | . | gene | MSTRG.33987 | 0.066555 | 0.017882 | 0.224629 | |
| 1669 | . | gene | MSTRG.26687 | 0.113328 | 0.01793 | 0.224629 | |

| | | | | | | | |
|---|---|---|---|---|---|---|---|
| 1672 | . | gene | MSTRG.35874 | 2.643378 | 0.017942 | 0.224629 | |
| 1677 | . | gene | MSTRG.11435 | 0.095787 | 0.018017 | 0.224629 | |
| 1682 | . | gene | gene:Bo9g173840 | 0.997944 | 0.018128 | 0.224629 | |
| 1684 | . | gene | MSTRG.1332 | 0.251442 | 0.018128 | 0.224629 | |
| 1686 | . | gene | MSTRG.20644 | 0.965107 | 0.018128 | 0.224629 | |
| 1687 | . | gene | MSTRG.23961 | 0.990194 | 0.018128 | 0.224629 | |
| 1688 | . | gene | MSTRG.28815 | 0.964908 | 0.018128 | 0.224629 | |
| 1689 | . | gene | MSTRG.4939 | 0.98248 | 0.018128 | 0.224629 | |
| 1695 | . | gene | MSTRG.12544 | 0.994777 | 0.018128 | 0.224629 | |
| 1699 | . | gene | MSTRG.12879 | 0.96884 | 0.018128 | 0.224629 | |
| 1701 | . | gene | MSTRG.13222 | 0.992369 | 0.018128 | 0.224629 | |
| 1703 | . | gene | MSTRG.16753 | 0.989219 | 0.018128 | 0.224629 | |
| 1705 | . | gene | MSTRG.18586 | 0.973257 | 0.018128 | 0.224629 | |
| 1706 | . | gene | MSTRG.19090 | 0.93055 | 0.018128 | 0.224629 | |
| 1709 | . | gene | MSTRG.20518 | 0.989005 | 0.018128 | 0.224629 | |
| 1710 | . | gene | MSTRG.22085 | 0.986288 | 0.018128 | 0.224629 | |
| 1712 | . | gene | MSTRG.24136 | 0.979391 | 0.018128 | 0.224629 | |
| 1715 | . | gene | MSTRG.26906 | 0.983735 | 0.018128 | 0.224629 | |
| 1720 | . | gene | MSTRG.32799 | 0.993845 | 0.018128 | 0.224629 | |
| 1721 | . | gene | MSTRG.34563 | 0.967895 | 0.018128 | 0.224629 | |
| 1722 | . | gene | MSTRG.35812 | 0.95956 | 0.018128 | 0.224629 | |
| 1723 | . | gene | MSTRG.36123 | 0.991894 | 0.018128 | 0.224629 | |
| 1727 | . | gene | MSTRG.6511 | 0.311649 | 0.018128 | 0.224629 | |
| 1730 | . | gene | MSTRG.9510 | 0.974683 | 0.018128 | 0.224629 | |
| 1735 | . | gene | MSTRG.546 | 1.426022 | 0.018226 | 0.224632 | Bo1g011450.1 |
| 1746 | . | gene | MSTRG.2408 | 0.139988 | 0.01836 | 0.224632 | Bo1g107090.1 |
| 1758 | . | gene | MSTRG.4253 | 0.755696 | 0.018419 | 0.224632 | Bo2g033970.1 |
| 1748 | . | gene | MSTRG.6561 | 3.40627 | 0.018366 | 0.224632 | Bo2g161260.1 |
| 1741 | . | gene | MSTRG.6873 | 0.592431 | 0.018311 | 0.224632 | Bo3g002900.1 |

| | | | | | | | |
|---|---|---|---|---|---|---|---|
| 1739 | . | gene | MSTRG.7086 | 1.754569 | 0.0183 | 0.224632 | Bo3g007040.1 |
| 1756 | . | gene | MSTRG.8824 | 0.650408 | 0.018403 | 0.224632 | Bo3g042730.1 |
| 1751 | . | gene | MSTRG.9977 | 3.322122 | 0.018389 | 0.224632 | Bo3g081910.1 |
| 1738 | . | gene | MSTRG.10259 | 2.079349 | 0.018289 | 0.224632 | Bo3g092370.1 |
| 1740 | . | gene | MSTRG.10260 | 0.389839 | 0.018305 | 0.224632 | Bo3g092380.1 |
| 1733 | . | gene | MSTRG.11686 | 1.577366 | 0.018186 | 0.224632 | Bo3g167190.1 |
| 1732 | . | gene | MSTRG.12085 | 0.16532 | 0.018149 | 0.224632 | Bo3g184600.1 |
| 1737 | . | gene | MSTRG.17299 | 1.570026 | 0.018276 | 0.224632 | Bo5g034700.1 |
| 1750 | . | gene | MSTRG.17765 | 0.141075 | 0.018375 | 0.224632 | Bo5g057230.1 |
| 1743 | . | gene | MSTRG.21032 | 2.110698 | 0.018331 | 0.224632 | Bo6g074410.1 |
| 1753 | . | gene | MSTRG.21342 | 1.702375 | 0.018393 | 0.224632 | Bo6g081130.1 |
| 1759 | . | gene | MSTRG.24071 | 1.398685 | 0.018423 | 0.224632 | Bo7g065110.1 |
| 1742 | . | gene | MSTRG.27735 | 0.734758 | 0.018328 | 0.224632 | Bo8g070650.1 |
| 1747 | . | gene | MSTRG.29660 | 2.239925 | 0.018361 | 0.224632 | Bo8g117250.1 |
| 1736 | . | gene | MSTRG.32749 | 1.688084 | 0.018258 | 0.224632 | Bo9g152450.1 |
| 1734 | . | gene | MSTRG.27620 | 1.704569 | 0.01819 | 0.224632 | |
| 1744 | . | gene | MSTRG.6407 | 0.208988 | 0.018342 | 0.224632 | |
| 1745 | . | gene | MSTRG.23459 | 0.33185 | 0.018355 | 0.224632 | |
| 1749 | . | gene | MSTRG.36078 | 5.101846 | 0.018372 | 0.224632 | |
| 1752 | . | gene | MSTRG.24114 | 0.241642 | 0.01839 | 0.224632 | |
| 1754 | . | gene | MSTRG.31797 | 1.65365 | 0.018396 | 0.224632 | |
| 1755 | . | gene | MSTRG.11900 | 0.689378 | 0.018396 | 0.224632 | |
| 1757 | . | gene | MSTRG.35254 | 1.726081 | 0.018416 | 0.224632 | |
| 1760 | . | gene | MSTRG.29836 | 0.827749 | 0.018439 | 0.224694 | Bo9g004350.1 |
| 1761 | . | gene | MSTRG.13754 | 1.356519 | 0.018456 | 0.224774 | Bo4g097010.1 |
| 1763 | . | gene | MSTRG.11607 | 2.324254 | 0.0185 | 0.224861 | Bo3g162760.1 |
| 1762 | . | gene | MSTRG.15173 | 0.318796 | 0.018481 | 0.224861 | Bo4g169310.1 |
| 1765 | . | gene | MSTRG.26395 | 1.141549 | 0.018505 | 0.224861 | Bo8g004710.1 |
| 1764 | . | gene | MSTRG.32322 | 1.779075 | 0.018504 | 0.224861 | |

| | | | | | | | |
|---|---|---|---|---|---|---|---|
| 17 66 | . | gene | MSTRG.3564 4 | 3.4836 94 | 0.0185 2 | 0.2249 17 | |
| 17 68 | . | gene | MSTRG.5647 | 0.7722 6 | 0.0185 58 | 0.2250 58 | Bo2g1181 30.1 |
| 17 67 | . | gene | MSTRG.6792 | 1.0551 3 | 0.0185 57 | 0.2250 58 | Bo3g0016 60.1 |
| 17 69 | . | gene | MSTRG.1592 9 | 3.6577 57 | 0.0185 63 | 0.2250 58 | Bo4g1956 50.1 |
| 17 70 | . | gene | MSTRG.2690 | 1.2731 12 | 0.0185 99 | 0.2252 05 | Bo1g1248 50.1 |
| 17 71 | . | gene | MSTRG.5827 | 0.4879 03 | 0.0186 01 | 0.2252 05 | Bo2g1302 90.1 |
| 17 72 | . | gene | MSTRG.1798 7 | 1.7500 71 | 0.0186 07 | 0.2252 05 | |
| 17 73 | . | gene | MSTRG.1812 9 | 0.0594 42 | 0.0186 29 | 0.2252 14 | Bo5g0813 50.1 |
| 17 75 | . | gene | MSTRG.1975 7 | 1.4987 17 | 0.0186 39 | 0.2252 14 | Bo5g1526 90.1 |
| 17 74 | . | gene | MSTRG.1076 4 | 0.9936 89 | 0.0186 37 | 0.2252 14 | |
| 17 76 | . | gene | MSTRG.6759 | 0.3863 5 | 0.0186 52 | 0.2252 37 | Bo3g0011 20.1 |
| 17 78 | . | gene | MSTRG.3492 | 0.5409 52 | 0.0186 94 | 0.2254 17 | Bo2g0087 70.1 |
| 17 77 | . | gene | MSTRG.2915 5 | 0.5396 63 | 0.0186 91 | 0.2254 17 | Bo8g1056 90.1 |
| 17 79 | . | gene | MSTRG.2865 9 | 2.0637 87 | 0.0187 06 | 0.2254 17 | |
| 17 80 | . | gene | MSTRG.3052 5 | 0.1019 99 | 0.0187 09 | 0.2254 17 | |
| 17 83 | . | gene | MSTRG.273 | 1.5471 55 | 0.0187 65 | 0.2255 26 | Bo1g0066 40.1 |
| 17 87 | ATCEN2 | gene | MSTRG.2015 | 1.5114 97 | 0.0187 95 | 0.2255 26 | Bo1g0847 00.1 |
| 17 81 | . | gene | MSTRG.7549 | 1.4021 59 | 0.0187 51 | 0.2255 26 | Bo3g0170 50.1 |
| 17 86 | . | gene | MSTRG.1029 6 | 0.0913 91 | 0.0187 94 | 0.2255 26 | Bo3g0939 40.1 |
| 17 88 | . | gene | MSTRG.1182 9 | 2.7719 84 | 0.0188 1 | 0.2255 26 | Bo3g1736 40.1 |
| 17 84 | . | gene | MSTRG.2681 6 | 0.6231 58 | 0.0187 7 | 0.2255 26 | Bo8g0308 00.1 |
| 17 82 | . | gene | MSTRG.1790 2 | 1.6840 14 | 0.0187 51 | 0.2255 26 | |
| 17 85 | . | gene | MSTRG.2655 6 | 0.3307 89 | 0.0187 93 | 0.2255 26 | |
| 17 89 | . | gene | MSTRG.2098 9 | 2.7098 32 | 0.0188 12 | 0.2255 26 | |
| 17 91 | . | gene | MSTRG.1160 1 | 0.7046 88 | 0.0188 47 | 0.2256 92 | Bo3g1625 70.1 |
| 17 90 | . | gene | MSTRG.3521 9 | 19.228 66 | 0.0188 46 | 0.2256 92 | |
| 17 92 | . | gene | MSTRG.2438 2 | 0.3635 65 | 0.0188 7 | 0.2258 36 | Bo7g0780 30.1 |
| 17 93 | . | gene | MSTRG.6811 | 0.5377 6 | 0.0188 82 | 0.2258 58 | Bo3g0018 60.1 |
| 17 94 | . | gene | MSTRG.2114 0 | 1.9730 07 | 0.0189 02 | 0.2259 13 | Bo6g0770 20.1 |

| | | | | | | | |
|---|---|---|---|---|---|---|---|
| 1795 | . | gene | MSTRG.29674 | 1.041756 | 0.018908 | 0.225913 | Bo8g117580.1 |
| 1796 | . | gene | MSTRG.14669 | 4.162931 | 0.018923 | 0.225973 | Bo4g140770.1 |
| 1797 | . | gene | MSTRG.26557 | 0.19325 | 0.018969 | 0.226391 | |
| 1798 | . | gene | MSTRG.32947 | 3.024918 | 0.018983 | 0.22643 | |
| 1799 | . | gene | MSTRG.19741 | 0.674501 | 0.018993 | 0.226432 | Bo5g151500.1 |
| 1800 | . | gene | MSTRG.7611 | 0.456057 | 0.019006 | 0.226462 | |
| 1806 | . | gene | MSTRG.3264 | 0.624111 | 0.019088 | 0.226509 | Bo1g155120.1 |
| 1804 | . | gene | MSTRG.8729 | 0.779411 | 0.019071 | 0.226509 | Bo3g040290.1 |
| 1801 | . | gene | MSTRG.16581 | 1.62748 | 0.019039 | 0.226509 | Bo5g010910.1 |
| 1808 | . | gene | MSTRG.22048 | 0.543812 | 0.019095 | 0.226509 | Bo6g108740.1 |
| 1805 | . | gene | MSTRG.25618 | 1.411633 | 0.019082 | 0.226509 | Bo7g108500.1 |
| 1802 | . | gene | MSTRG.30847 | 0.576992 | 0.019044 | 0.226509 | Bo9g035470.1 |
| 1807 | SNX2a | gene | MSTRG.32433 | 0.954866 | 0.019092 | 0.226509 | Bo9g135080.1 |
| 1803 | . | gene | MSTRG.33407 | 3.39798 | 0.019054 | 0.226509 | Bo9g173950.1 |
| 1809 | . | gene | MSTRG.21910 | 0.781062 | 0.019108 | 0.226537 | Bo6g103900.1 |
| 1812 | . | gene | MSTRG.9900 | 2.46914 | 0.019149 | 0.226646 | Bo3g077420.1 |
| 1811 | . | gene | MSTRG.14544 | 0.110027 | 0.019147 | 0.226646 | Bo4g137360.1 |
| 1810 | . | gene | MSTRG.14487 | 0.436495 | 0.019133 | 0.226646 | |
| 1820 | . | gene | MSTRG.4425 | 0.405751 | 0.019266 | 0.22669 | Bo2g046160.1 |
| 1818 | . | gene | MSTRG.8410 | 1.344705 | 0.019238 | 0.22669 | Bo3g034480.1 |
| 1815 | . | gene | MSTRG.12306 | 0.543045 | 0.019204 | 0.22669 | Bo4g012790.1 |
| 1822 | . | gene | MSTRG.13112 | 0.586821 | 0.019267 | 0.22669 | Bo4g045380.1 |
| 1813 | . | gene | MSTRG.14908 | 0.321993 | 0.019193 | 0.22669 | Bo4g151750.1 |
| 1823 | . | gene | MSTRG.15517 | 0.429589 | 0.01927 | 0.22669 | Bo4g186020.1 |
| 1817 | . | gene | MSTRG.17865 | 1.53662 | 0.019217 | 0.22669 | Bo5g062980.1 |
| 1824 | . | gene | MSTRG.26308 | 0.39706 | 0.019279 | 0.22669 | Bo8g001260.1 |
| 1819 | . | gene | MSTRG.26805 | 1.256854 | 0.019264 | 0.22669 | Bo8g030620.1 |
| 1816 | . | gene | MSTRG.29182 | 0.608607 | 0.019209 | 0.22669 | Bo8g106090.1 |
| 1821 | CPD | gene | MSTRG.33617 | 0.532475 | 0.019266 | 0.22669 | Bo9g177000.1 |

| | | | | | | | |
|---|---|---|---|---|---|---|---|
| 18<br>14 | . | gene | MSTRG.2615<br>5 | 0.4065<br>22 | 0.0192<br>01 | 0.2266<br>9 | |
| 18<br>25 | . | gene | MSTRG.2556 | 1.3017<br>78 | 0.0192<br>96 | 0.2267<br>63 | Bo1g1177<br>00.1 |
| 18<br>26 | . | gene | MSTRG.1762 | 0.5483<br>52 | 0.0193<br>22 | 0.2269<br>44 | Bo1g0590<br>60.1 |
| 18<br>27 | . | gene | MSTRG.1738<br>5 | 1.2849<br>07 | 0.0193<br>77 | 0.2273<br>96 | Bo5g0390<br>60.1 |
| 18<br>28 | . | gene | MSTRG.1776<br>2 | 0.6419<br>78 | 0.0193<br>82 | 0.2273<br>96 | |
| 18<br>30 | . | gene | MSTRG.2920<br>1 | 1.9737<br>68 | 0.0194<br>16 | 0.2275<br>54 | Bo8g1063<br>30.1 |
| 18<br>29 | . | gene | MSTRG.3478<br>7 | 12.024<br>29 | 0.0194<br>06 | 0.2275<br>54 | |
| 18<br>31 | . | gene | MSTRG.2516 | 1.6647<br>82 | 0.0194<br>5 | 0.2278<br>2 | |
| 18<br>32 | . | gene | MSTRG.1357<br>1 | 0.9907<br>52 | 0.0194<br>81 | 0.2280<br>6 | Bo4g0787<br>00.1 |
| 18<br>34 | . | gene | MSTRG.1757<br>5 | 0.8431<br>37 | 0.0195<br>13 | 0.2280<br>85 | Bo5g0452<br>20.1 |
| 18<br>33 | . | gene | MSTRG.1959<br>4 | 1.6490<br>38 | 0.0194<br>97 | 0.2280<br>85 | Bo5g1453<br>90.1 |
| 18<br>35 | . | gene | MSTRG.9736 | 0.3947<br>38 | 0.0195<br>15 | 0.2280<br>85 | |
| 18<br>38 | . | gene | MSTRG.7397 | 1.9338<br>27 | 0.0195<br>58 | 0.2281<br>07 | Bo3g0129<br>70.1 |
| 18<br>39 | . | gene | MSTRG.1097<br>1 | 0.5001<br>24 | 0.0195<br>59 | 0.2281<br>07 | Bo3g1342<br>20.1 |
| 18<br>37 | . | gene | MSTRG.1587<br>0 | 1.8089<br>94 | 0.0195<br>4 | 0.2281<br>07 | Bo4g1944<br>70.1 |
| 18<br>36 | . | gene | MSTRG.2637<br>5 | 1.4240<br>77 | 0.0195<br>29 | 0.2281<br>07 | Bo8g0043<br>50.1 |
| 18<br>40 | . | gene | MSTRG.1952<br>3 | 1.4292<br>89 | 0.0195<br>91 | 0.2283<br>56 | Bo5g1433<br>00.1 |
| 18<br>41 | . | gene | MSTRG.3247 | 1.3520<br>16 | 0.0196<br>28 | 0.2285<br>86 | Bo1g1549<br>10.1 |
| 18<br>42 | . | gene | MSTRG.9524 | 0.1785<br>49 | 0.0196<br>32 | 0.2285<br>86 | Bo3g0647<br>80.1 |
| 18<br>43 | . | gene | MSTRG.2786<br>8 | 0.3962<br>75 | 0.0196<br>8 | 0.2290<br>14 | |
| 18<br>44 | . | gene | MSTRG.6023 | 2.3182<br>05 | 0.0197<br>12 | 0.2292<br>7 | Bo2g1366<br>60.1 |
| 18<br>46 | . | gene | MSTRG.1079 | 0.1199<br>43 | 0.0197<br>44 | 0.2293<br>89 | Bo1g0284<br>40.1 |
| 18<br>45 | . | gene | MSTRG.3129<br>3 | 0.6649<br>26 | 0.0197<br>34 | 0.2293<br>89 | Bo9g0642<br>10.1 |
| 18<br>48 | . | gene | MSTRG.9380 | 0.4455<br>42 | 0.0197<br>8 | 0.2294<br>36 | Bo3g0616<br>50.1 |
| 18<br>47 | . | gene | MSTRG.1053<br>7 | 0.5478<br>55 | 0.0197<br>6 | 0.2294<br>36 | Bo3g1032<br>20.1 |
| 18<br>49 | . | gene | MSTRG.1626<br>3 | 2.6536<br>29 | 0.0197<br>89 | 0.2294<br>36 | Bo5g0056<br>00.1 |
| 18<br>50 | . | gene | MSTRG.2405<br>8 | 1.5321<br>84 | 0.0197<br>91 | 0.2294<br>36 | Bo7g0648<br>90.1 |
| 18<br>51 | . | gene | MSTRG.2911 | 1.5812<br>46 | 0.0198<br>25 | 0.2297<br>09 | Bo1g1397<br>90.1 |
| 18<br>52 | . | gene | MSTRG.1806<br>9 | 1.6379<br>9 | 0.0198<br>48 | 0.2298<br>54 | |

| | | | | | | | |
|---|---|---|---|---|---|---|---|
| 1862 | . | gene | MSTRG.8337 | 0.840251 | 0.020005 | 0.230423 | Bo3g032440.1 |
| 1861 | PEX19-1 | gene | MSTRG.9213 | 1.516379 | 0.020004 | 0.230423 | Bo3g057100.1 |
| 1856 | . | gene | MSTRG.12801 | 0.98449 | 0.019941 | 0.230423 | Bo4g030620.1 |
| 1859 | . | gene | MSTRG.13964 | 0.169409 | 0.019975 | 0.230423 | Bo4g108270.1 |
| 1853 | . | gene | MSTRG.15424 | 1.660744 | 0.019932 | 0.230423 | Bo4g182460.1 |
| 1858 | . | gene | MSTRG.17604 | 0.600683 | 0.019967 | 0.230423 | Bo5g048550.1 |
| 1860 | . | gene | MSTRG.18573 | 1.388848 | 0.020001 | 0.230423 | Bo5g113680.1 |
| 1854 | . | gene | MSTRG.31176 | 1.136766 | 0.019933 | 0.230423 | Bo9g059030.1 |
| 1855 | . | gene | MSTRG.35071 | 8.748384 | 0.019936 | 0.230423 | |
| 1857 | . | gene | MSTRG.13866 | 0.349264 | 0.019958 | 0.230423 | |
| 1864 | . | gene | MSTRG.13668 | 1.262095 | 0.020031 | 0.230469 | Bo4g087080.1 |
| 1863 | . | gene | MSTRG.28342 | 1.765768 | 0.020025 | 0.230469 | Bo8g090240.1 |
| 1865 | . | gene | MSTRG.2061 | 0.664475 | 0.020067 | 0.23077 | Bo1g087680.1 |
| 1868 | . | gene | MSTRG.12375 | 1.590352 | 0.020115 | 0.23089 | Bo4g016140.1 |
| 1867 | CAD4 | gene | MSTRG.18605 | 1.82976 | 0.020108 | 0.23089 | Bo5g114100.1 |
| 1866 | . | gene | MSTRG.28163 | 0.691816 | 0.020106 | 0.23089 | Bo8g082700.1 |
| 1869 | . | gene | MSTRG.35336 | 44.66831 | 0.020121 | 0.23089 | |
| 1870 | . | gene | MSTRG.29483 | 1.733928 | 0.02014 | 0.230986 | Bo8g112580.1 |
| 1873 | . | gene | MSTRG.3516 | 1.597734 | 0.020194 | 0.231093 | Bo2g009110.1 |
| 1874 | . | gene | MSTRG.17605 | 0.316824 | 0.020199 | 0.231093 | Bo5g048570.1 |
| 1871 | . | gene | MSTRG.24900 | 0.2652 | 0.020161 | 0.231093 | Bo7g093030.1 |
| 1872 | . | gene | MSTRG.25386 | 0.624869 | 0.020186 | 0.231093 | Bo7g104340.1 |
| 1875 | . | gene | MSTRG.35098 | 0.504653 | 0.020203 | 0.231093 | |
| 1876 | . | gene | MSTRG.7326 | 0.682471 | 0.020221 | 0.231172 | Bo3g010820.1 |
| 1877 | . | gene | MSTRG.15579 | 1.620264 | 0.020242 | 0.231295 | Bo4g187000.1 |
| 1878 | . | gene | MSTRG.15266 | 1.891768 | 0.0203 | 0.231827 | Bo4g174830.1 |
| 1879 | . | gene | MSTRG.11697 | 0.875461 | 0.020334 | 0.23209 | Bo3g167410.1 |
| 1880 | . | gene | MSTRG.20701 | 0.572433 | 0.020375 | 0.232349 | Bo6g064220.1 |
| 1881 | ATB5-B | gene | MSTRG.30044 | 1.375352 | 0.020385 | 0.232349 | Bo9g009600.1 |

| | | | | | | | |
|---|---|---|---|---|---|---|---|
| 1882 | . | gene | MSTRG.35603 | 1.588069 | 0.020389 | 0.232349 | |
| 1884 | . | gene | MSTRG.4526 | 0.354499 | 0.020481 | 0.233071 | Bo2g050980.1 |
| 1883 | . | gene | MSTRG.21797 | 1.341328 | 0.020477 | 0.233071 | Bo6g099050.1 |
| 1885 | . | gene | MSTRG.34681 | 0.333015 | 0.020485 | 0.233071 | |
| 1890 | . | gene | MSTRG.1201 | 1.454062 | 0.020569 | 0.233174 | Bo1g037320.1 |
| 1891 | . | gene | MSTRG.7051 | 1.506153 | 0.020575 | 0.233174 | Bo3g005620.1 |
| 1887 | . | gene | MSTRG.9134 | 0.664783 | 0.020528 | 0.233174 | Bo3g054890.1 |
| 1886 | . | gene | MSTRG.16977 | 1.740012 | 0.020519 | 0.233174 | Bo5g021970.1 |
| 1893 | . | gene | MSTRG.32728 | 1.671329 | 0.020581 | 0.233174 | Bo9g152130.1 |
| 1888 | . | gene | MSTRG.33982 | 0.656096 | 0.020534 | 0.233174 | |
| 1889 | . | gene | MSTRG.34267 | 1.666027 | 0.020561 | 0.233174 | |
| 1892 | . | gene | MSTRG.36090 | 0.690791 | 0.020576 | 0.233174 | |
| 1894 | . | gene | MSTRG.16968 | 5.30328 | 0.020596 | 0.23322 | Bo5g021810.1 |
| 1896 | . | gene | MSTRG.462 | 2.266672 | 0.020636 | 0.233423 | Bo1g010270.1 |
| 1895 | . | gene | MSTRG.31583 | 0.131201 | 0.020625 | 0.233423 | Bo9g081920.1 |
| 1900 | . | gene | MSTRG.15945 | 0.366909 | 0.020711 | 0.233782 | Bo4g195870.1 |
| 1899 | . | gene | MSTRG.17580 | 1.256739 | 0.020711 | 0.233782 | Bo5g045270.1 |
| 1898 | . | gene | MSTRG.31472 | 11.92068 | 0.02071 | 0.233782 | Bo9g075990.1 |
| 1897 | . | gene | MSTRG.32012 | 1.926972 | 0.020686 | 0.233782 | Bo9g109760.1 |
| 1901 | . | gene | MSTRG.23097 | 1.216051 | 0.020745 | 0.234045 | Bo7g017510.1 |
| 1902 | . | gene | MSTRG.33268 | 1.520734 | 0.020774 | 0.234251 | Bo9g170090.1 |
| 1904 | . | gene | MSTRG.1417 | 1.2099 | 0.020818 | 0.234434 | Bo1g045370.1 |
| 1908 | . | gene | MSTRG.8600 | 1.559551 | 0.020856 | 0.234434 | Bo3g037260.1 |
| 1903 | . | gene | MSTRG.20624 | 3.037582 | 0.020817 | 0.234434 | Bo6g056220.1 |
| 1907 | . | gene | MSTRG.26912 | 4.018215 | 0.020852 | 0.234434 | Bo8g036770.1 |
| 1905 | . | gene | MSTRG.33370 | 1.83273 | 0.020833 | 0.234434 | |
| 1906 | . | gene | MSTRG.7620 | 0.113395 | 0.020841 | 0.234434 | |
| 1909 | . | gene | MSTRG.31937 | 1.101385 | 0.020876 | 0.23453 | Bo9g104660.1 |
| 1910 | . | gene | MSTRG.30797 | 0.801587 | 0.020889 | 0.234556 | Bo9g032420.1 |

| | | | | | | | |
|---|---|---|---|---|---|---|---|
| 19<br>11 | . | gene | MSTRG.3063<br>5 | 2.0005<br>37 | 0.0209 | 0.2345<br>58 | Bo9g0276<br>70.1 |
| 19<br>12 | . | gene | MSTRG.2489<br>6 | 1.3684<br>19 | 0.0209<br>13 | 0.2345<br>77 | |
| 19<br>13 | . | gene | MSTRG.1127<br>0 | 2.7486<br>24 | 0.0209<br>77 | 0.2351<br>51 | Bo3g1477<br>40.1 |
| 19<br>14 | . | gene | MSTRG.1568<br>8 | 1.5746<br>38 | 0.0209<br>86 | 0.2351<br>51 | Bo4g1887<br>50.1 |
| 19<br>28 | . | gene | MSTRG.3301 | 0.9072<br>31 | 0.0211<br>8 | 0.2353<br>95 | Bo1g1575<br>80.1 |
| 19<br>18 | . | gene | MSTRG.4719 | 1.3935<br>81 | 0.0211<br>18 | 0.2353<br>95 | Bo2g0600<br>40.1 |
| 19<br>22 | . | gene | MSTRG.8927 | 3.8217<br>76 | 0.0211<br>39 | 0.2353<br>95 | Bo3g0455<br>30.1 |
| 19<br>23 | . | gene | MSTRG.1047<br>7 | 0.3456<br>1 | 0.0211<br>47 | 0.2353<br>95 | Bo3g1014<br>70.1 |
| 19<br>20 | . | gene | MSTRG.2455<br>3 | 1.7807<br>13 | 0.0211<br>26 | 0.2353<br>95 | Bo7g0820<br>90.1 |
| 19<br>15 | . | gene | MSTRG.2564<br>2 | 1.9867<br>31 | 0.0210<br>66 | 0.2353<br>95 | Bo7g1088<br>90.1 |
| 19<br>16 | . | gene | MSTRG.2589<br>2 | 1.3063<br>94 | 0.0210<br>97 | 0.2353<br>95 | Bo7g1138<br>90.1 |
| 19<br>29 | . | gene | MSTRG.2641<br>3 | 0.6538<br>11 | 0.0211<br>82 | 0.2353<br>95 | Bo8g0059<br>30.1 |
| 19<br>17 | . | gene | MSTRG.2742<br>4 | 0.5000<br>33 | 0.0211<br>14 | 0.2353<br>95 | Bo8g0625<br>30.1 |
| 19<br>25 | . | gene | MSTRG.2902<br>5 | 1.3067<br>38 | 0.0211<br>59 | 0.2353<br>95 | Bo8g1026<br>60.1 |
| 19<br>27 | . | gene | MSTRG.2975<br>0 | 0.7380<br>44 | 0.0211<br>69 | 0.2353<br>95 | Bo9g0023<br>40.1 |
| 19<br>19 | . | gene | MSTRG.1242<br>1 | 0.6083<br>98 | 0.0211<br>26 | 0.2353<br>95 | |
| 19<br>21 | . | gene | MSTRG.3602<br>4 | 0.5276<br>86 | 0.0211<br>36 | 0.2353<br>95 | |
| 19<br>24 | . | gene | MSTRG.2660<br>9 | 1.6568<br>09 | 0.0211<br>55 | 0.2353<br>95 | |
| 19<br>26 | . | gene | MSTRG.3585<br>4 | 1.9696<br>01 | 0.0211<br>66 | 0.2353<br>95 | |
| 19<br>30 | . | gene | MSTRG.3523<br>3 | 0.9148<br>22 | 0.0211<br>83 | 0.2353<br>95 | |
| 19<br>31 | . | gene | MSTRG.3211<br>7 | 1.4473<br>31 | 0.0212<br>07 | 0.2355<br>35 | Bo9g1173<br>30.1 |
| 19<br>33 | . | gene | MSTRG.1095 | 3.8085<br>08 | 0.0212<br>57 | 0.2357<br>15 | Bo1g0306<br>80.1 |
| 19<br>34 | . | gene | MSTRG.1314<br>5 | 1.4170<br>99 | 0.0212<br>61 | 0.2357<br>15 | Bo4g0458<br>60.1 |
| 19<br>35 | . | gene | MSTRG.2723<br>9 | 1.5383<br>38 | 0.0212<br>67 | 0.2357<br>15 | Bo8g0557<br>40.1 |
| 19<br>32 | . | gene | MSTRG.2797<br>2 | 1.1366<br>09 | 0.0212<br>35 | 0.2357<br>15 | Bo8g0794<br>80.1 |
| 19<br>36 | . | gene | MSTRG.3132<br>9 | 1.8808<br>21 | 0.0212<br>79 | 0.2357<br>29 | |
| 19<br>37 | . | gene | MSTRG.5655 | 1.5753<br>29 | 0.0213<br>36 | 0.2362<br>34 | Bo2g1192<br>50.1 |
| 19<br>38 | . | gene | MSTRG.5065 | 0.0353<br>94 | 0.0213<br>53 | 0.2362<br>99 | |
| 19<br>39 | . | gene | MSTRG.2002<br>2 | 0.7385<br>09 | 0.0213<br>68 | 0.2363<br>47 | |

| | | | | | | | |
|---|---|---|---|---|---|---|---|
| 1941 | . | gene | MSTRG.18828 | 0.72421 | 0.021402 | 0.236482 | Bo5g123520.1 |
| 1940 | . | gene | MSTRG.25338 | 0.702156 | 0.021393 | 0.236482 | Bo7g103710.1 |
| 1942 | . | gene | MSTRG.299 | 0.059469 | 0.021416 | 0.236518 | |
| 1943 | KEA6 | gene | MSTRG.33253 | 0.711977 | 0.021448 | 0.23675 | Bo9g169860.1 |
| 1944 | . | gene | MSTRG.10346 | 0.650491 | 0.021486 | 0.237046 | Bo3g097720.1 |
| 1945 | . | gene | MSTRG.2232 | 2.247155 | 0.021529 | 0.2374 | Bo1g099920.1 |
| 1946 | . | gene | MSTRG.26451 | 1.435526 | 0.021544 | 0.237435 | Bo8g008420.1 |
| 1947 | . | gene | MSTRG.11225 | 1.47859 | 0.021558 | 0.237467 | Bo3g144020.1 |
| 1948 | . | gene | MSTRG.2490 | 1.281884 | 0.021625 | 0.237928 | Bo1g112380.1 |
| 1949 | . | gene | MSTRG.4041 | 1.20877 | 0.021628 | 0.237928 | Bo2g024450.1 |
| 1950 | . | gene | MSTRG.10700 | 0.55751 | 0.021633 | 0.237928 | |
| 1952 | . | gene | MSTRG.7675 | 0.692461 | 0.021666 | 0.238044 | Bo3g018650.1 |
| 1951 | . | gene | MSTRG.16241 | 1.794844 | 0.021664 | 0.238044 | Bo5g005290.1 |
| 1953 | . | gene | MSTRG.25407 | 0.794154 | 0.021718 | 0.238485 | Bo7g104690.1 |
| 1954 | . | gene | MSTRG.17630 | 0.893193 | 0.021728 | 0.238485 | |
| 1955 | . | gene | MSTRG.15247 | 3.193188 | 0.021748 | 0.238586 | Bo4g173600.1 |
| 1956 | . | gene | MSTRG.28295 | 0.404214 | 0.021769 | 0.238696 | Bo8g089550.1 |
| 1957 | . | gene | MSTRG.11320 | 0.682841 | 0.02182 | 0.239133 | Bo3g149400.1 |
| 1961 | . | gene | MSTRG.11533 | 2.915951 | 0.021895 | 0.239462 | Bo3g157010.1 |
| 1958 | . | gene | MSTRG.17627 | 0.743738 | 0.02187 | 0.239462 | Bo5g048950.1 |
| 1960 | . | gene | MSTRG.20820 | 1.528578 | 0.021894 | 0.239462 | Bo6g067090.1 |
| 1959 | . | gene | MSTRG.30526 | 1.514902 | 0.021889 | 0.239462 | Bo9g022670.1 |
| 1962 | SPT42 | gene | MSTRG.6653 | 1.517176 | 0.021933 | 0.239756 | Bo2g165830.1 |
| 1963 | . | gene | MSTRG.1735 | 1.565205 | 0.021953 | 0.239845 | Bo1g057630.1 |
| 1965 | . | gene | MSTRG.10574 | 0.656995 | 0.022038 | 0.240423 | Bo3g105780.1 |
| 1968 | . | gene | MSTRG.20318 | 1.828116 | 0.022061 | 0.240423 | Bo6g033210.1 |
| 1967 | . | gene | MSTRG.29415 | 0.273345 | 0.022061 | 0.240423 | Bo8g109630.1 |
| 1964 | . | gene | MSTRG.10787 | 2.80295 | 0.022017 | 0.240423 | |
| 1966 | . | gene | MSTRG.2592 | 1.408247 | 0.022042 | 0.240423 | |

| | | | | | | | |
|---|---|---|---|---|---|---|---|
| 19 72 | . | gene | MSTRG.1671 1 | 1.6516 75 | 0.0221 38 | 0.2407 66 | Bo5g0169 10.1 |
| 19 69 | . | gene | MSTRG.2599 6 | 1.2929 39 | 0.0221 08 | 0.2407 66 | Bo7g1153 90.1 |
| 19 70 | . | gene | MSTRG.2145 1 | 0.1038 32 | 0.0221 28 | 0.2407 66 | |
| 19 71 | . | gene | MSTRG.1241 2 | 1.8136 55 | 0.0221 34 | 0.2407 66 | |
| 19 73 | . | gene | MSTRG.3142 4 | 0.7968 43 | 0.0221 57 | 0.2408 53 | Bo9g0730 70.1 |
| 19 74 | . | gene | MSTRG.431 | 0.8641 02 | 0.0221 78 | 0.2409 56 | Bo1g0087 60.1 |
| 19 76 | . | gene | MSTRG.9645 | 0.4137 72 | 0.0222 12 | 0.2410 87 | Bo3g0664 80.1 |
| 19 75 | . | gene | MSTRG.3181 7 | 0.1381 35 | 0.0222 02 | 0.2410 87 | Bo9g0958 40.1 |
| 19 77 | . | gene | MSTRG.2994 7 | 1.5054 06 | 0.0222 57 | 0.2414 46 | Bo9g0078 80.1 |
| 19 79 | . | gene | MSTRG.9677 | 1.6099 37 | 0.0222 94 | 0.2415 26 | Bo3g0669 50.1 |
| 19 78 | . | gene | MSTRG.1249 5 | 0.7048 46 | 0.0222 81 | 0.2415 26 | Bo4g0217 50.1 |
| 19 82 | . | gene | MSTRG.2598 8 | 0.7999 85 | 0.0223 2 | 0.2415 26 | Bo7g1152 30.1 |
| 19 81 | . | gene | MSTRG.2741 9 | 0.2837 06 | 0.0223 15 | 0.2415 26 | Bo8g0614 30.1 |
| 19 80 | . | gene | MSTRG.3316 0 | 0.7101 39 | 0.0223 02 | 0.2415 26 | Bo9g1665 10.1 |
| 19 84 | . | gene | MSTRG.9106 | 1.6245 59 | 0.0223 82 | 0.2417 36 | Bo3g0545 70.1 |
| 19 88 | . | gene | MSTRG.1013 9 | 2.5458 25 | 0.0224 12 | 0.2417 36 | Bo3g0870 00.1 |
| 19 86 | . | gene | MSTRG.1850 1 | 1.8626 82 | 0.0224 01 | 0.2417 36 | Bo5g1086 30.1 |
| 19 83 | . | gene | MSTRG.7354 | 0.6000 06 | 0.0223 68 | 0.2417 36 | |
| 19 85 | . | gene | MSTRG.1815 5 | 5.2592 04 | 0.0223 93 | 0.2417 36 | |
| 19 87 | . | gene | MSTRG.3595 7 | 4.0793 18 | 0.0224 02 | 0.2417 36 | |
| 19 89 | . | gene | MSTRG.1888 7 | 2.6446 78 | 0.0224 19 | 0.2417 36 | |
| 19 90 | . | gene | MSTRG.1015 9 | 1.2661 95 | 0.0224 45 | 0.2418 03 | Bo3g0872 60.1 |
| 19 91 | . | gene | MSTRG.1061 8 | 0.1438 26 | 0.0224 47 | 0.2418 03 | Bo3g1075 10.1 |
| 19 93 | . | gene | MSTRG.2842 1 | 1.5160 78 | 0.0224 87 | 0.2419 85 | Bo8g0912 40.1 |
| 19 92 | . | gene | MSTRG.2969 6 | 0.7526 26 | 0.0224 77 | 0.2419 85 | Bo8g1179 10.1 |
| 19 95 | . | gene | MSTRG.243 | 0.3897 55 | 0.0225 17 | 0.2419 94 | Bo1g0062 40.1 |
| 19 97 | . | gene | MSTRG.4210 | 0.2734 96 | 0.0225 33 | 0.2419 94 | Bo2g0291 00.1 |
| 19 96 | . | gene | MSTRG.2358 0 | 0.9413 26 | 0.0225 28 | 0.2419 94 | Bo7g0455 60.1 |
| 19 94 | . | gene | MSTRG.2876 9 | 1.4133 29 | 0.0225 04 | 0.2419 94 | Bo8g0988 30.1 |

| | | | | | | | |
|---|---|---|---|---|---|---|---|
| 1998 | ATG10 | gene | MSTRG.19459 | 1.294177 | 0.022545 | 0.242005 | Bo5g141450.1 |
| 1999 | . | gene | MSTRG.11056 | 2.606626 | 0.022562 | 0.242064 | Bo3g138360.1 |
| 2001 | . | gene | MSTRG.1218 | 0.665818 | 0.022612 | 0.242246 | Bo1g037580.1 |
| 2002 | . | gene | MSTRG.8926 | 1.308419 | 0.022613 | 0.242246 | Bo3g045510.1 |
| 2000 | . | gene | MSTRG.4505 | 0.975306 | 0.022592 | 0.242246 | |
| 2003 | . | gene | MSTRG.5039 | 1.23973 | 0.022655 | 0.242568 | Bo2g082390.1 |
| 2004 | . | gene | MSTRG.32331 | 1.29942 | 0.022665 | 0.242568 | |
| 2005 | . | gene | MSTRG.22924 | 0.268034 | 0.022692 | 0.242666 | |
| 2006 | . | gene | MSTRG.2226 | 0.10009 | 0.022697 | 0.242666 | |
| 2010 | . | gene | MSTRG.20339 | 0.687963 | 0.022764 | 0.242892 | Bo6g034790.1 |
| 2007 | . | gene | MSTRG.21913 | 0.817169 | 0.022738 | 0.242892 | |
| 2008 | . | gene | MSTRG.4156 | 1.854703 | 0.022747 | 0.242892 | |
| 2009 | . | gene | MSTRG.35649 | 0.453404 | 0.022762 | 0.242892 | |
| 2011 | . | gene | MSTRG.35040 | 0.170124 | 0.022788 | 0.24294 | |
| 2012 | . | gene | MSTRG.35608 | 2.522719 | 0.022791 | 0.24294 | |
| 2013 | . | gene | MSTRG.19964 | 0.388024 | 0.022827 | 0.243205 | |
| 2015 | . | gene | MSTRG.32023 | 0.71186 | 0.022855 | 0.243262 | Bo9g109880.1 |
| 2014 | . | gene | MSTRG.21067 | 0.374295 | 0.022844 | 0.243262 | |
| 2016 | . | gene | MSTRG.14535 | 0.627275 | 0.02288 | 0.243405 | |
| 2018 | . | gene | MSTRG.2304 | 0.305829 | 0.022954 | 0.243748 | Bo1g102380.1 |
| 2017 | . | gene | MSTRG.7260 | 0.662314 | 0.022924 | 0.243748 | Bo3g010180.1 |
| 2022 | . | gene | MSTRG.26461 | 0.135611 | 0.02299 | 0.243748 | Bo8g009510.1 |
| 2020 | . | gene | MSTRG.28225 | 1.647088 | 0.022958 | 0.243748 | Bo8g083550.1 |
| 2019 | . | gene | MSTRG.34917 | 0.638296 | 0.022955 | 0.243748 | |
| 2021 | . | gene | MSTRG.15408 | 0.609141 | 0.022985 | 0.243748 | |
| 2023 | . | gene | MSTRG.33967 | 1.498689 | 0.022992 | 0.243748 | |
| 2024 | . | gene | MSTRG.7458 | 1.949066 | 0.023032 | 0.243936 | Bo3g013820.1 |
| 2028 | . | gene | MSTRG.13014 | 1.949011 | 0.023099 | 0.243936 | Bo4g040060.1 |
| 2031 | . | gene | MSTRG.14066 | 0.166139 | 0.023122 | 0.243936 | Bo4g115360.1 |

| | | | | | | | |
|---|---|---|---|---|---|---|---|
| 20<br>29 | . | gene | MSTRG.1599<br>1 | 2.0914<br>28 | 0.0231<br>07 | 0.2439<br>36 | Bo4g1984<br>90.1 |
| 20<br>36 | . | gene | MSTRG.2048<br>9 | 1.0381<br>17 | 0.0231<br>57 | 0.2439<br>36 | Bo6g0468<br>10.1 |
| 20<br>25 | . | gene | MSTRG.2307<br>8 | 1.3313<br>25 | 0.0230<br>37 | 0.2439<br>36 | Bo7g0161<br>90.1 |
| 20<br>33 | . | gene | MSTRG.2473<br>8 | 2.1932<br>48 | 0.0231<br>42 | 0.2439<br>36 | Bo7g0886<br>20.1 |
| 20<br>30 | . | gene | MSTRG.2567<br>2 | 3.0993<br>91 | 0.0231<br>21 | 0.2439<br>36 | Bo7g1093<br>40.1 |
| 20<br>26 | . | gene | MSTRG.2744<br>4 | 0.6632<br>92 | 0.0230<br>62 | 0.2439<br>36 | Bo8g0648<br>30.1 |
| 20<br>32 | . | gene | MSTRG.3152<br>3 | 1.5506<br>72 | 0.0231<br>24 | 0.2439<br>36 | Bo9g0777<br>90.1 |
| 20<br>27 | . | gene | MSTRG.1329<br>9 | 1.5043<br>47 | 0.0230<br>67 | 0.2439<br>36 | |
| 20<br>34 | . | gene | MSTRG.1089<br>0 | 0.3075<br>08 | 0.0231<br>42 | 0.2439<br>36 | |
| 20<br>35 | . | gene | gene:Bo5g00<br>8160 | 2.4781<br>14 | 0.0231<br>56 | 0.2439<br>36 | |
| 20<br>38 | . | gene | MSTRG.8504 | 0.5774<br>14 | 0.0232<br>04 | 0.2441<br>06 | Bo3g0359<br>40.1 |
| 20<br>37 | . | gene | MSTRG.1222<br>8 | 1.4995<br>91 | 0.0231<br>98 | 0.2441<br>06 | Bo4g0057<br>20.1 |
| 20<br>39 | . | gene | MSTRG.2439<br>5 | 5.1209<br>43 | 0.0232<br>08 | 0.2441<br>06 | |
| 20<br>40 | . | gene | MSTRG.1793<br>3 | 1.0631<br>95 | 0.0232<br>32 | 0.2441<br>27 | Bo5g0670<br>90.1 |
| 20<br>42 | . | gene | MSTRG.2256<br>5 | 0.7594<br>35 | 0.0232<br>44 | 0.2441<br>27 | Bo6g1210<br>60.1 |
| 20<br>41 | HOS15 | gene | MSTRG.3042<br>1 | 1.5265<br>3 | 0.0232<br>43 | 0.2441<br>27 | Bo9g0202<br>90.1 |
| 20<br>43 | . | gene | MSTRG.3691 | 4.0282<br>96 | 0.0232<br>64 | 0.2441<br>82 | Bo2g0117<br>30.1 |
| 20<br>44 | . | gene | MSTRG.1564<br>8 | 0.5552<br>95 | 0.0232<br>72 | 0.2441<br>82 | Bo4g1880<br>90.1 |
| 20<br>45 | . | gene | MSTRG.2906<br>7 | 2.6213<br>98 | 0.0232<br>83 | 0.2441<br>82 | Bo8g1043<br>20.1 |
| 20<br>47 | . | gene | MSTRG.6673 | 10.814<br>36 | 0.0233<br>34 | 0.2443<br>68 | Bo2g1661<br>30.1 |
| 20<br>48 | . | gene | MSTRG.1960<br>6 | 1.1910<br>61 | 0.0233<br>35 | 0.2443<br>68 | Bo5g1455<br>40.1 |
| 20<br>46 | . | gene | MSTRG.2531<br>0 | 1.5415<br>16 | 0.0233<br>17 | 0.2443<br>68 | Bo7g1012<br>10.1 |
| 20<br>50 | . | gene | MSTRG.1752<br>7 | 0.7167<br>46 | 0.0233<br>78 | 0.2445<br>44 | Bo5g0432<br>00.1 |
| 20<br>49 | . | gene | MSTRG.2916<br>7 | 0.1888<br>86 | 0.0233<br>76 | 0.2445<br>44 | Bo8g1058<br>10.1 |
| 20<br>51 | . | gene | MSTRG.86 | 0.9762<br>05 | 0.0233<br>86 | 0.2445<br>44 | |
| 20<br>55 | . | gene | MSTRG.6476 | 0.1962<br>08 | 0.0234<br>78 | 0.2448<br>08 | Bo2g1590<br>80.1 |
| 20<br>52 | . | gene | MSTRG.2031<br>6 | 1.2978<br>27 | 0.0234<br>31 | 0.2448<br>08 | Bo6g0331<br>50.1 |
| 20<br>54 | . | gene | MSTRG.2389<br>3 | 1.2770<br>46 | 0.0234<br>69 | 0.2448<br>08 | Bo7g0598<br>70.1 |
| 20<br>56 | . | gene | MSTRG.2644<br>8 | 1.5813<br>53 | 0.0234<br>85 | 0.2448<br>08 | Bo8g0073<br>70.1 |

| | | | | | | | |
|---|---|---|---|---|---|---|---|
| 20 57 | . | gene | MSTRG.2813 7 | 1.7650 91 | 0.0234 86 | 0.2448 08 | Bo8g0823 00.1 |
| 20 58 | TSB1 | gene | MSTRG.3214 6 | 1.8904 9 | 0.0234 91 | 0.2448 08 | Bo9g1188 50.1 |
| 20 53 | . | gene | MSTRG.3008 4 | 0.7882 14 | 0.0234 39 | 0.2448 08 | |
| 20 60 | . | gene | MSTRG.1991 2 | 2.0138 39 | 0.0235 23 | 0.2449 05 | Bo6g0105 40.1 |
| 20 59 | . | gene | MSTRG.1743 2 | 0.6705 73 | 0.0235 17 | 0.2449 05 | |
| 20 73 | . | gene | MSTRG.2326 | 1.1934 72 | 0.0237 07 | 0.2452 68 | Bo1g1037 30.1 |
| 20 62 | . | gene | MSTRG.1023 7 | 1.6446 03 | 0.0236 18 | 0.2452 68 | Bo3g0917 70.1 |
| 20 69 | . | gene | MSTRG.1752 4 | 0.6113 53 | 0.0236 73 | 0.2452 68 | Bo5g0431 60.1 |
| 20 61 | ATMSH5 | gene | MSTRG.1849 9 | 0.8744 48 | 0.0236 13 | 0.2452 68 | Bo5g1086 10.1 |
| 20 63 | . | gene | MSTRG.2519 8 | 1.2808 39 | 0.0236 41 | 0.2452 68 | Bo7g0982 70.1 |
| 20 71 | . | gene | MSTRG.2916 3 | 0.1161 58 | 0.0236 98 | 0.2452 68 | Bo8g1057 60.1 |
| 20 65 | . | gene | MSTRG.3099 4 | 1.1657 24 | 0.0236 57 | 0.2452 68 | Bo9g0453 60.1 |
| 20 64 | . | gene | MSTRG.3354 7 | 1.5660 99 | 0.0236 51 | 0.2452 68 | Bo9g1759 60.1 |
| 20 66 | . | gene | MSTRG.1733 3 | 0.1163 89 | 0.0236 59 | 0.2452 68 | |
| 20 67 | . | gene | MSTRG.1760 1 | 0.1624 55 | 0.0236 71 | 0.2452 68 | |
| 20 68 | . | gene | MSTRG.1844 | 2.0039 12 | 0.0236 71 | 0.2452 68 | |
| 20 70 | . | gene | MSTRG.2655 8 | 0.2971 14 | 0.0236 81 | 0.2452 68 | |
| 20 72 | . | gene | MSTRG.3164 1 | 0.8019 84 | 0.0237 01 | 0.2452 68 | |
| 20 76 | . | gene | MSTRG.4128 | 1.1693 62 | 0.0237 53 | 0.2452 79 | Bo2g0267 30.1 |
| 20 75 | . | gene | MSTRG.6213 | 0.9379 46 | 0.0237 52 | 0.2452 79 | Bo2g1479 00.1 |
| 20 79 | . | gene | MSTRG.1027 6 | 2.0530 02 | 0.0237 87 | 0.2452 79 | Bo3g0936 40.1 |
| 20 80 | . | gene | MSTRG.1754 1 | 0.12 | 0.0237 88 | 0.2452 79 | Bo5g0434 30.1 |
| 20 78 | . | gene | MSTRG.2843 0 | 0.7179 6 | 0.0237 7 | 0.2452 79 | Bo8g0914 20.1 |
| 20 77 | . | gene | MSTRG.3153 6 | 3.8043 91 | 0.0237 58 | 0.2452 79 | Bo9g0780 50.1 |
| 20 74 | . | gene | MSTRG.2341 2 | 1.7234 29 | 0.0237 25 | 0.2452 79 | |
| 20 83 | TLP18.3 | gene | MSTRG.2038 | 0.7663 39 | 0.0238 68 | 0.2455 51 | Bo1g0861 90.1 |
| 20 81 | . | gene | MSTRG.3246 | 1.6944 81 | 0.0238 45 | 0.2455 51 | Bo1g1549 00.1 |
| 20 82 | . | gene | MSTRG.7197 | 0.2692 12 | 0.0238 59 | 0.2455 51 | Bo3g0095 40.1 |
| 20 84 | . | gene | MSTRG.1030 1 | 0.2764 22 | 0.0238 71 | 0.2455 51 | Bo3g0940 80.1 |

| | | | | | | | |
|---|---|---|---|---|---|---|---|
| 2085 | . | gene | MSTRG.18091 | 3.01946 | 0.023872 | 0.245551 | Bo5g078760.1 |
| 2087 | . | gene | MSTRG.3387 | 1.609644 | 0.023895 | 0.245559 | Bo2g003980.1 |
| 2086 | . | gene | MSTRG.24339 | 1.236716 | 0.023893 | 0.245559 | Bo7g077520.1 |
| 2088 | . | gene | MSTRG.3593 | 1.542199 | 0.023931 | 0.245726 | Bo2g010140.1 |
| 2089 | . | gene | MSTRG.36140 | 1.284879 | 0.023934 | 0.245726 | |
| 2098 | . | gene | MSTRG.5077 | 1.334881 | 0.024077 | 0.245844 | Bo2g085240.1 |
| 2090 | . | gene | MSTRG.7061 | 1.436467 | 0.023967 | 0.245844 | Bo3g006710.1 |
| 2100 | . | gene | MSTRG.9254 | 1.330456 | 0.024097 | 0.245844 | Bo3g058620.1 |
| 2096 | . | gene | MSTRG.10969 | 2.020176 | 0.024056 | 0.245844 | Bo3g134200.1 |
| 2095 | . | gene | MSTRG.17388 | 1.312198 | 0.024054 | 0.245844 | Bo5g039080.1 |
| 2094 | . | gene | MSTRG.20139 | 0.678023 | 0.024052 | 0.245844 | Bo6g026460.1 |
| 2103 | . | gene | MSTRG.20220 | 1.235125 | 0.024106 | 0.245844 | Bo6g030700.1 |
| 2097 | . | gene | MSTRG.20938 | 0.417965 | 0.024072 | 0.245844 | Bo6g069740.1 |
| 2091 | . | gene | MSTRG.23346 | 0.528203 | 0.023971 | 0.245844 | Bo7g033090.1 |
| 2101 | . | gene | MSTRG.25227 | 0.113975 | 0.024098 | 0.245844 | Bo7g098720.1 |
| 2092 | . | gene | MSTRG.32414 | 0.75314 | 0.023991 | 0.245844 | Bo9g134790.1 |
| 2102 | . | gene | MSTRG.33717 | 5.020203 | 0.024102 | 0.245844 | Bo9g181420.1 |
| 2093 | . | gene | MSTRG.20480 | 0.690222 | 0.024042 | 0.245844 | |
| 2099 | . | gene | MSTRG.24279 | 0.541764 | 0.024086 | 0.245844 | |
| 2104 | . | gene | MSTRG.26373 | 1.494219 | 0.024126 | 0.24587 | Bo8g004280.1 |
| 2105 | . | gene | MSTRG.15325 | 2.086774 | 0.024132 | 0.24587 | |
| 2109 | . | gene | MSTRG.16181 | 2.017393 | 0.024179 | 0.24588 | Bo5g004450.1 |
| 2106 | . | gene | MSTRG.17226 | 0.772231 | 0.024156 | 0.24588 | Bo5g030680.1 |
| 2107 | . | gene | MSTRG.26185 | 1.604694 | 0.02417 | 0.24588 | Bo7g118360.1 |
| 2108 | . | gene | MSTRG.36080 | 1.845719 | 0.024172 | 0.24588 | |
| 2114 | . | gene | MSTRG.11994 | 0.842859 | 0.024261 | 0.246044 | Bo3g181040.1 |
| 2113 | . | gene | MSTRG.17614 | 0.096598 | 0.024248 | 0.246044 | Bo5g048730.1 |
| 2115 | . | gene | MSTRG.21476 | 1.228883 | 0.024264 | 0.246044 | Bo6g086310.1 |
| 2110 | . | gene | MSTRG.23649 | 0.236371 | 0.024217 | 0.246044 | Bo7g049740.1 |

| | | | | | | | |
|---|---|---|---|---|---|---|---|
| 21 12 | . | gene | MSTRG.3062 3 | 6.3127 32 | 0.0242 37 | 0.2460 44 | Bo9g0275 60.1 |
| 21 11 | . | gene | MSTRG.8961 | 1.5352 84 | 0.0242 24 | 0.2460 44 | |
| 21 16 | . | gene | MSTRG.4021 | 0.9287 2 | 0.0243 04 | 0.2463 39 | Bo2g0239 60.1 |
| 21 17 | . | gene | MSTRG.2956 2 | 1.8929 54 | 0.0243 48 | 0.2466 61 | Bo8g1147 00.1 |
| 21 18 | . | gene | MSTRG.1268 2 | 3.8329 86 | 0.0243 71 | 0.2466 96 | Bo4g0263 70.1 |
| 21 19 | . | gene | MSTRG.3549 4 | 5.5281 55 | 0.0243 74 | 0.2466 96 | |
| 21 20 | . | gene | MSTRG.2260 7 | 0.2386 87 | 0.0243 89 | 0.2467 29 | Bo6g1216 50.1 |
| 21 21 | . | gene | MSTRG.2954 3 | 0.0069 44 | 0.0244 1 | 0.2468 08 | Bo8g1144 20.1 |
| 21 22 | . | gene | MSTRG.3453 5 | 0.6717 22 | 0.0244 2 | 0.2468 08 | |
| 21 23 | . | gene | MSTRG.1690 0 | 2.1509 87 | 0.0244 43 | 0.2469 26 | Bo5g0209 30.1 |
| 21 26 | . | gene | MSTRG.9702 | 0.6936 4 | 0.0244 99 | 0.2470 15 | Bo3g0684 70.1 |
| 21 28 | . | gene | MSTRG.1082 0 | 1.3779 18 | 0.0245 09 | 0.2470 15 | Bo3g1236 30.1 |
| 21 27 | . | gene | MSTRG.1185 8 | 1.4439 02 | 0.0245 | 0.2470 15 | Bo3g1750 20.1 |
| 21 24 | . | gene | MSTRG.1870 3 | 1.7800 88 | 0.0244 69 | 0.2470 15 | Bo5g1186 80.1 |
| 21 25 | . | gene | MSTRG.3429 0 | 4.0602 95 | 0.0244 9 | 0.2470 15 | |
| 21 29 | . | gene | MSTRG.3107 7 | 0.5996 01 | 0.0246 96 | 0.2487 84 | Bo9g0544 80.1 |
| 21 30 | . | gene | MSTRG.1496 | 1.6018 71 | 0.0247 11 | 0.2488 17 | Bo1g0494 70.1 |
| 21 31 | . | gene | MSTRG.7119 | 0.2220 51 | 0.0247 48 | 0.2490 7 | Bo3g0075 70.1 |
| 21 33 | . | gene | MSTRG.2043 4 | 1.4376 75 | 0.0248 14 | 0.2494 34 | Bo6g0439 60.1 |
| 21 32 | . | gene | MSTRG.2077 0 | 16.924 75 | 0.0248 1 | 0.2494 34 | Bo6g0663 10.1 |
| 21 34 | . | gene | MSTRG.2141 6 | 1.3136 32 | 0.0248 19 | 0.2494 34 | Bo6g0833 10.1 |
| 21 40 | . | gene | MSTRG.533 | 1.5679 07 | 0.0249 19 | 0.2496 75 | Bo1g0112 80.1 |
| 21 35 | . | gene | MSTRG.1583 | 0.6144 06 | 0.0248 89 | 0.2496 75 | Bo1g0530 70.1 |
| 21 41 | . | gene | MSTRG.7318 | 1.8152 82 | 0.0249 34 | 0.2496 75 | Bo3g0107 30.1 |
| 21 39 | . | gene | MSTRG.7926 | 1.1535 95 | 0.0249 05 | 0.2496 75 | Bo3g0235 10.1 |
| 21 42 | . | gene | MSTRG.1003 8 | 1.6371 43 | 0.0249 41 | 0.2496 75 | Bo3g0835 70.1 |
| 21 36 | HIPL2 | gene | MSTRG.1060 5 | 1.4876 37 | 0.0248 98 | 0.2496 75 | Bo3g1073 20.1 |
| 21 43 | . | gene | MSTRG.1973 3 | 1.5941 03 | 0.0249 48 | 0.2496 75 | Bo5g1514 00.1 |
| 21 37 | . | gene | MSTRG.2779 4 | 1.6208 08 | 0.0248 98 | 0.2496 75 | |

| | | | | | | | |
|---|---|---|---|---|---|---|---|
| 21 38 | . | gene | MSTRG.3443 1 | 0.8882 12 | 0.0249 04 | 0.2496 75 | |
| 21 44 | . | gene | MSTRG.1848 1 | 0.5502 92 | 0.0249 86 | 0.2499 4 | |
| 21 45 | . | gene | MSTRG.2806 6 | 2.1497 34 | 0.0249 98 | 0.2499 44 | Bo8g0812 00.1 |
| 21 48 | . | gene | MSTRG.2485 6 | 1.8118 16 | 0.0250 66 | 0.2501 89 | Bo7g0912 50.1 |
| 21 46 | . | gene | MSTRG.2668 8 | 0.1281 46 | 0.0250 41 | 0.2501 89 | Bo8g0255 30.1 |
| 21 47 | . | gene | MSTRG.2469 9 | 2.0424 19 | 0.0250 48 | 0.2501 89 | |
| 21 49 | . | gene | MSTRG.2715 5 | 0.5359 48 | 0.0250 69 | 0.2501 89 | |
| 21 50 | . | gene | MSTRG.2838 0 | 2.6932 29 | 0.0251 13 | 0.2505 09 | Bo8g0906 70.1 |
| 21 51 | . | gene | MSTRG.2410 5 | 1.2746 37 | 0.0251 43 | 0.2506 98 | Bo7g0656 60.1 |
| 21 52 | ATNADP-ME4 | gene | MSTRG.2146 5 | 1.2281 96 | 0.0251 67 | 0.2507 08 | Bo6g0860 90.1 |
| 21 53 | . | gene | MSTRG.3077 3 | 0.7476 73 | 0.0251 68 | 0.2507 08 | Bo9g0310 20.1 |
| 21 54 | . | gene | MSTRG.147 | 1.5524 39 | 0.0252 44 | 0.2513 5 | Bo1g0052 10.1 |
| 21 55 | . | gene | MSTRG.3597 2 | 0.2920 64 | 0.0252 79 | 0.2515 82 | |
| 21 56 | . | gene | MSTRG.1856 3 | 0.4653 22 | 0.0253 03 | 0.2517 03 | Bo5g1124 80.1 |
| 21 57 | . | gene | MSTRG.1983 4 | 0.8339 14 | 0.0253 25 | 0.2518 02 | |
| 21 58 | . | gene | MSTRG.5739 | 0.4191 79 | 0.0253 5 | 0.2519 39 | Bo2g1257 10.1 |
| 21 59 | . | gene | MSTRG.1466 | 1.7457 91 | 0.0253 72 | 0.2520 39 | Bo1g0480 80.1 |
| 21 63 | . | gene | MSTRG.6893 | 1.3091 04 | 0.0254 54 | 0.2522 28 | Bo3g0031 60.1 |
| 21 65 | . | gene | MSTRG.1312 7 | 1.1121 77 | 0.0254 62 | 0.2522 28 | Bo4g0455 70.1 |
| 21 62 | . | gene | MSTRG.1526 5 | 1.5367 59 | 0.0254 33 | 0.2522 28 | Bo4g1738 20.1 |
| 21 61 | . | gene | MSTRG.2132 5 | 1.4336 03 | 0.0254 24 | 0.2522 28 | Bo6g0809 60.1 |
| 21 60 | . | gene | MSTRG.969 | 0.2224 88 | 0.0254 06 | 0.2522 28 | |
| 21 64 | . | gene | MSTRG.4934 | 0.7762 08 | 0.0254 6 | 0.2522 28 | |
| 21 66 | . | gene | MSTRG.2300 2 | 1.6012 13 | 0.0254 87 | 0.2522 52 | Bo7g0116 70.1 |
| 21 67 | . | gene | MSTRG.3078 7 | 0.8028 47 | 0.0254 87 | 0.2522 52 | Bo9g0322 30.1 |
| 21 70 | . | gene | MSTRG.6332 | 0.3623 6 | 0.0255 49 | 0.2523 73 | Bo2g1518 50.1 |
| 21 69 | . | gene | MSTRG.7467 | 0.5953 51 | 0.0255 49 | 0.2523 73 | Bo3g0139 20.1 |
| 21 71 | . | gene | MSTRG.1129 4 | 1.3060 67 | 0.0255 59 | 0.2523 73 | Bo3g1490 80.1 |
| 21 72 | . | gene | MSTRG.1195 6 | 1.2036 51 | 0.0255 73 | 0.2523 73 | Bo3g1784 10.1 |

| | | | | | | | |
|---|---|---|---|---|---|---|---|
| 2175 | . | gene | MSTRG.14681 | 0.680776 | 0.025594 | 0.252373 | Bo4g141950.1 |
| 2174 | . | gene | MSTRG.18320 | 1.354703 | 0.025586 | 0.252373 | Bo5g095730.1 |
| 2168 | . | gene | MSTRG.10869 | 0.389273 | 0.025538 | 0.252373 | |
| 2173 | . | gene | MSTRG.3449 | 1.787566 | 0.02558 | 0.252373 | |
| 2181 | . | gene | MSTRG.1153 | 1.531969 | 0.025677 | 0.252388 | Bo1g034460.1 |
| 2177 | PIP5K2 | gene | MSTRG.5193 | 0.447379 | 0.025634 | 0.252388 | Bo2g092920.1 |
| 2176 | . | gene | MSTRG.6824 | 67.20341 | 0.025634 | 0.252388 | Bo3g002090.1 |
| 2182 | . | gene | MSTRG.6900 | 4.855954 | 0.025689 | 0.252388 | Bo3g003280.1 |
| 2184 | . | gene | MSTRG.7386 | 2.654484 | 0.025713 | 0.252388 | Bo3g012760.1 |
| 2185 | . | gene | MSTRG.10981 | 1.803843 | 0.025713 | 0.252388 | Bo3g134390.1 |
| 2180 | . | gene | MSTRG.26623 | 1.3861 | 0.025658 | 0.252388 | Bo8g020460.1 |
| 2178 | . | gene | MSTRG.36097 | 0.413792 | 0.02565 | 0.252388 | |
| 2179 | . | gene | MSTRG.18876 | 0.536837 | 0.025653 | 0.252388 | |
| 2183 | . | gene | MSTRG.15767 | 0.297606 | 0.025693 | 0.252388 | |
| 2190 | . | gene | MSTRG.1639 | 0.133893 | 0.025775 | 0.252396 | Bo1g055030.1 |
| 2187 | . | gene | MSTRG.8086 | 1.113143 | 0.025744 | 0.252396 | Bo3g026960.1 |
| 2186 | . | gene | MSTRG.17651 | 0.17107 | 0.02574 | 0.252396 | Bo5g049270.1 |
| 2188 | . | gene | MSTRG.35651 | 0.430371 | 0.025769 | 0.252396 | |
| 2189 | . | gene | MSTRG.34480 | 0.095085 | 0.025774 | 0.252396 | |
| 2191 | . | gene | MSTRG.31769 | 3.623904 | 0.025784 | 0.252396 | |
| 2192 | . | gene | MSTRG.2489 | 0.668459 | 0.025843 | 0.252665 | Bo1g112370.1 |
| 2193 | . | gene | MSTRG.11811 | 0.746262 | 0.025847 | 0.252665 | Bo3g172310.1 |
| 2194 | . | gene | MSTRG.34243 | 0.117571 | 0.025847 | 0.252665 | |
| 2195 | . | gene | MSTRG.20909 | 2.555954 | 0.025861 | 0.252685 | |
| 2196 | . | gene | MSTRG.556 | 1.346632 | 0.02589 | 0.252839 | Bo1g011660.1 |
| 2202 | . | gene | MSTRG.1723 | 0.670995 | 0.025963 | 0.252839 | Bo1g057480.1 |
| 2198 | . | gene | MSTRG.9591 | 1.344672 | 0.025919 | 0.252839 | Bo3g065700.1 |
| 2201 | . | gene | MSTRG.26095 | 0.95636 | 0.025961 | 0.252839 | Bo7g116910.1 |
| 2197 | . | gene | MSTRG.34421 | 1.332808 | 0.025904 | 0.252839 | |

| | | | | | | | |
|---|---|---|---|---|---|---|---|
| 2199 | . | gene | MSTRG.18799 | 0.997791 | 0.025954 | 0.252839 | |
| 2200 | . | gene | MSTRG.22377 | 1.596936 | 0.02596 | 0.252839 | |
| 2203 | . | gene | MSTRG.34158 | 3.960851 | 0.025971 | 0.252839 | |
| 2204 | . | gene | MSTRG.3974 | 0.707265 | 0.02602 | 0.253199 | |
| 2211 | . | gene | MSTRG.1445 | 1.334345 | 0.026127 | 0.253398 | Bo1g047750.1 |
| 2210 | . | gene | MSTRG.7171 | 2.559802 | 0.026125 | 0.253398 | Bo3g009240.1 |
| 2212 | ARR16 | gene | MSTRG.8454 | 1.797251 | 0.026135 | 0.253398 | Bo3g035110.1 |
| 2208 | . | gene | MSTRG.9508 | 0.699013 | 0.026098 | 0.253398 | Bo3g064550.1 |
| 2209 | . | gene | MSTRG.13540 | 1.371949 | 0.026112 | 0.253398 | Bo4g078360.1 |
| 2206 | . | gene | MSTRG.20020 | 1.873381 | 0.026089 | 0.253398 | Bo6g018570.1 |
| 2207 | . | gene | MSTRG.32772 | 1.328169 | 0.026095 | 0.253398 | Bo9g153850.1 |
| 2205 | . | gene | MSTRG.34714 | 0.500666 | 0.026057 | 0.253398 | |
| 2213 | . | gene | MSTRG.16382 | 0.644759 | 0.026196 | 0.253876 | |
| 2215 | . | gene | MSTRG.1200 | 2.594381 | 0.026239 | 0.254065 | Bo1g037310.1 |
| 2214 | . | gene | MSTRG.16923 | 2.018305 | 0.026234 | 0.254065 | Bo5g021320.1 |
| 2221 | . | gene | MSTRG.1117 | 6.614233 | 0.026329 | 0.254115 | Bo1g032020.1 |
| 2220 | . | gene | MSTRG.7190 | 0.907356 | 0.026326 | 0.254115 | Bo3g009460.1 |
| 2228 | . | gene | MSTRG.14810 | 1.406131 | 0.026404 | 0.254115 | Bo4g148130.1 |
| 2224 | ATMES1 | gene | MSTRG.14904 | 0.773574 | 0.026367 | 0.254115 | Bo4g151710.1 |
| 2218 | . | gene | MSTRG.27678 | 1.114832 | 0.026311 | 0.254115 | Bo8g068580.1 |
| 2222 | . | gene | MSTRG.28500 | 0.50165 | 0.026334 | 0.254115 | Bo8g092430.1 |
| 2227 | . | gene | MSTRG.30984 | 0.897168 | 0.026402 | 0.254115 | Bo9g044230.1 |
| 2216 | . | gene | MSTRG.6805 | 1.341807 | 0.026281 | 0.254115 | |
| 2217 | . | gene | MSTRG.26603 | 0.100738 | 0.026289 | 0.254115 | |
| 2219 | . | gene | MSTRG.15283 | 1.183071 | 0.026326 | 0.254115 | |
| 2223 | . | gene | MSTRG.1963 | 0.364407 | 0.026341 | 0.254115 | |
| 2225 | . | gene | MSTRG.10891 | 0.384634 | 0.026384 | 0.254115 | |
| 2226 | . | gene | MSTRG.11072 | 0.673323 | 0.026387 | 0.254115 | |
| 2229 | . | gene | MSTRG.24701 | 0.556639 | 0.02641 | 0.254115 | |

| | | | | | | | |
|---|---|---|---|---|---|---|---|
| 22<br>31 | LAC11 | gene | MSTRG.3385 | 3.0473<br>83 | 0.0264<br>68 | 0.2543<br>34 | Bo2g0039<br>50.1 |
| 22<br>38 | . | gene | MSTRG.9563 | 0.6864<br>48 | 0.0265<br>42 | 0.2543<br>34 | Bo3g0652<br>70.1 |
| 22<br>35 | . | gene | MSTRG.1659<br>5 | 1.7281<br>73 | 0.0265<br>29 | 0.2543<br>34 | Bo5g0121<br>10.1 |
| 22<br>33 | . | gene | MSTRG.1665<br>8 | 0.7869<br>68 | 0.0264<br>99 | 0.2543<br>34 | Bo5g0150<br>50.1 |
| 22<br>30 | . | gene | MSTRG.1862<br>1 | 0.2879<br>04 | 0.0264<br>49 | 0.2543<br>34 | Bo5g1152<br>40.1 |
| 22<br>32 | . | gene | MSTRG.2127<br>6 | 1.3941<br>93 | 0.0264<br>82 | 0.2543<br>34 | Bo6g0800<br>30.1 |
| 22<br>34 | . | gene | MSTRG.2291<br>0 | 2.3478<br>88 | 0.0265<br>08 | 0.2543<br>34 | Bo7g0060<br>60.1 |
| 22<br>37 | . | gene | MSTRG.2444<br>6 | 1.9718<br>37 | 0.0265<br>37 | 0.2543<br>34 | Bo7g0801<br>40.1 |
| 22<br>36 | . | gene | MSTRG.2780<br>8 | 0.8137<br>83 | 0.0265<br>34 | 0.2543<br>34 | Bo8g0727<br>10.1 |
| 22<br>39 | . | gene | MSTRG.2364<br>7 | 0.1378<br>51 | 0.0265<br>52 | 0.2543<br>34 | |
| 22<br>40 | . | gene | MSTRG.8406 | 0.6444<br>66 | 0.0265<br>9 | 0.2545<br>89 | Bo3g0344<br>30.1 |
| 22<br>41 | . | gene | MSTRG.3122<br>4 | 1.3347<br>21 | 0.0266<br>34 | 0.2548<br>95 | Bo9g0598<br>50.1 |
| 22<br>43 | . | gene | MSTRG.1370<br>8 | 0.3304<br>97 | 0.0266<br>8 | 0.2551<br>1 | Bo4g0930<br>00.1 |
| 22<br>42 | . | gene | MSTRG.1889<br>8 | 0.3940<br>9 | 0.0266<br>8 | 0.2551<br>1 | Bo5g1266<br>40.1 |
| 22<br>46 | . | gene | MSTRG.320 | 1.3317<br>09 | 0.0267<br>36 | 0.2551<br>94 | Bo1g0072<br>60.1 |
| 22<br>49 | . | gene | MSTRG.1127<br>2 | 2.4099<br>7 | 0.0267<br>6 | 0.2551<br>94 | Bo3g1477<br>60.1 |
| 22<br>47 | . | gene | MSTRG.2561<br>9 | 0.6233<br>75 | 0.0267<br>44 | 0.2551<br>94 | Bo7g1085<br>10.1 |
| 22<br>45 | . | gene | MSTRG.2939<br>4 | 0.5245<br>62 | 0.0267<br>27 | 0.2551<br>94 | Bo8g1093<br>30.1 |
| 22<br>44 | . | gene | MSTRG.3461<br>6 | 1.6306<br>86 | 0.0267<br>25 | 0.2551<br>94 | |
| 22<br>48 | . | gene | MSTRG.1764<br>5 | 0.0558<br>82 | 0.0267<br>56 | 0.2551<br>94 | |
| 22<br>50 | . | gene | MSTRG.1761 | 0.8620<br>64 | 0.0267<br>87 | 0.2553<br>32 | Bo1g0590<br>50.1 |
| 22<br>53 | . | gene | MSTRG.8096 | 2.7330<br>75 | 0.0268<br>56 | 0.2554<br>36 | Bo3g0271<br>50.1 |
| 22<br>52 | . | gene | MSTRG.1299<br>2 | 0.3348<br>13 | 0.0268<br>5 | 0.2554<br>36 | Bo4g0395<br>30.1 |
| 22<br>55 | . | gene | MSTRG.1438<br>8 | 2.6024<br>33 | 0.0268<br>73 | 0.2554<br>36 | Bo4g1310<br>50.1 |
| 22<br>59 | . | gene | MSTRG.1552<br>4 | 0.0596<br>42 | 0.0269<br>05 | 0.2554<br>36 | Bo4g1861<br>70.1 |
| 22<br>56 | . | gene | MSTRG.1788<br>2 | 1.3427<br>47 | 0.0268<br>77 | 0.2554<br>36 | Bo5g0632<br>40.1 |
| 22<br>51 | . | gene | MSTRG.1837<br>2 | 1.3008<br>13 | 0.0268<br>1 | 0.2554<br>36 | Bo5g0984<br>00.1 |
| 22<br>57 | . | gene | MSTRG.1878<br>5 | 6.4124<br>39 | 0.0268<br>87 | 0.2554<br>36 | Bo5g1226<br>90.1 |
| 22<br>54 | . | gene | MSTRG.2068<br>8 | 1.3023<br>89 | 0.0268<br>63 | 0.2554<br>36 | Bo6g0610<br>60.1 |

| | | | | | | | |
|---|---|---|---|---|---|---|---|
| 2258 | . | gene | MSTRG.34373 | 0.815142 | 0.026898 | 0.255436 | |
| 2262 | . | gene | MSTRG.7525 | 1.417839 | 0.026995 | 0.255953 | Bo3g015700.1 |
| 2260 | . | gene | MSTRG.20098 | 1.210732 | 0.02698 | 0.255953 | Bo6g024780.1 |
| 2261 | ATTRX2 | gene | MSTRG.20703 | 1.569676 | 0.026988 | 0.255953 | Bo6g064280.1 |
| 2266 | . | gene | MSTRG.17557 | 2.886287 | 0.027058 | 0.256052 | Bo5g043850.1 |
| 2263 | . | gene | MSTRG.21931 | 0.650475 | 0.027032 | 0.256052 | Bo6g104060.1 |
| 2267 | . | gene | MSTRG.31146 | 1.615308 | 0.027065 | 0.256052 | Bo9g058480.1 |
| 2265 | . | gene | MSTRG.32750 | 1.594174 | 0.027057 | 0.256052 | Bo9g152500.1 |
| 2264 | . | gene | MSTRG.23419 | 0.759867 | 0.027051 | 0.256052 | |
| 2268 | eIFiso4G1 | gene | MSTRG.4207 | 1.364689 | 0.02708 | 0.25608 | Bo2g029000.1 |
| 2269 | . | gene | MSTRG.25260 | 0.852209 | 0.027137 | 0.256507 | |
| 2272 | . | gene | MSTRG.16224 | 0.364424 | 0.027196 | 0.256609 | Bo5g005070.1 |
| 2273 | . | gene | MSTRG.17886 | 1.393014 | 0.027196 | 0.256609 | Bo5g064300.1 |
| 2271 | . | gene | MSTRG.25496 | 0.088962 | 0.027186 | 0.256609 | Bo7g106210.1 |
| 2270 | . | gene | MSTRG.36005 | 0.189533 | 0.027168 | 0.256609 | |
| 2274 | . | gene | MSTRG.23782 | 0.140243 | 0.02724 | 0.25691 | Bo7g054770.1 |
| 2276 | . | gene | MSTRG.1451 | 1.393555 | 0.027292 | 0.257038 | Bo1g047820.1 |
| 2278 | . | gene | MSTRG.7231 | 1.358063 | 0.027308 | 0.257038 | Bo3g009920.1 |
| 2277 | . | gene | MSTRG.11198 | 0.569886 | 0.027298 | 0.257038 | Bo3g143660.1 |
| 2279 | . | gene | MSTRG.16647 | 1.608228 | 0.027314 | 0.257038 | Bo5g014810.1 |
| 2275 | . | gene | MSTRG.18513 | 3.309617 | 0.027285 | 0.257038 | Bo5g109790.1 |
| 2280 | . | gene | MSTRG.28782 | 1.917823 | 0.027325 | 0.257038 | Bo8g099020.1 |
| 2281 | . | gene | MSTRG.11311 | 0.505017 | 0.027354 | 0.257199 | Bo3g149310.1 |
| 2282 | . | gene | MSTRG.23251 | 1.669966 | 0.027388 | 0.257398 | |
| 2283 | . | gene | MSTRG.22417 | 0.711353 | 0.027419 | 0.257495 | Bo6g119190.1 |
| 2284 | . | gene | MSTRG.33097 | 1.201859 | 0.027431 | 0.257495 | Bo9g165620.1 |
| 2285 | . | gene | MSTRG.34607 | 0.669258 | 0.027434 | 0.257495 | |
| 2286 | MMZ3 | gene | MSTRG.14275 | 1.76927 | 0.027457 | 0.257595 | Bo4g125310.1 |
| 2287 | . | gene | MSTRG.27714 | 0.196607 | 0.027611 | 0.258931 | Bo8g070310.1 |

| | | | | | | | |
|---|---|---|---|---|---|---|---|
| 2288 | . | gene | MSTRG.33281 | 0.93996 | 0.027659 | 0.259266 | Bo9g171330.1 |
| 2289 | . | gene | MSTRG.19560 | 0.803759 | 0.027681 | 0.259364 | Bo5g144820.1 |
| 2290 | . | gene | MSTRG.21375 | 2.229538 | 0.027704 | 0.259377 | Bo6g081670.1 |
| 2291 | . | gene | MSTRG.26671 | 1.319262 | 0.027707 | 0.259377 | Bo8g023240.1 |
| 2292 | . | gene | MSTRG.5259 | 0.368817 | 0.027778 | 0.259923 | |
| 2293 | . | gene | MSTRG.27369 | 3.083734 | 0.027795 | 0.259972 | Bo8g059670.1 |
| 2294 | . | gene | MSTRG.20385 | 2.102405 | 0.02785 | 0.260245 | Bo6g039620.1 |
| 2295 | . | gene | MSTRG.16226 | 0.974967 | 0.027853 | 0.260245 | |
| 2296 | . | gene | MSTRG.16636 | 0.57829 | 0.02786 | 0.260245 | |
| 2297 | . | gene | MSTRG.36043 | 2.373705 | 0.027899 | 0.260495 | |
| 2298 | . | gene | MSTRG.18849 | 1.336542 | 0.027966 | 0.260999 | Bo5g124850.1 |
| 2299 | . | gene | MSTRG.22726 | 1.165066 | 0.028022 | 0.26141 | Bo6g124370.1 |
| 2300 | . | gene | MSTRG.9734 | 1.743758 | 0.028041 | 0.261472 | Bo3g068930.1 |
| 2301 | . | gene | MSTRG.7465 | 0.650736 | 0.028083 | 0.261754 | Bo3g013890.1 |
| 2302 | . | gene | MSTRG.11276 | 1.553594 | 0.028097 | 0.261769 | |
| 2304 | . | gene | MSTRG.16389 | 1.728871 | 0.028146 | 0.261953 | Bo5g008450.1 |
| 2303 | . | gene | MSTRG.19677 | 1.4308 | 0.028141 | 0.261953 | Bo5g148450.1 |
| 2305 | . | gene | MSTRG.32923 | 1.204606 | 0.028153 | 0.261953 | Bo9g160140.1 |
| 2307 | . | gene | MSTRG.9580 | 1.465765 | 0.028198 | 0.262004 | Bo3g065500.1 |
| 2310 | . | gene | MSTRG.10761 | 0.966342 | 0.02822 | 0.262004 | Bo3g116120.1 |
| 2306 | . | gene | MSTRG.24989 | 1.385588 | 0.028186 | 0.262004 | Bo7g095360.1 |
| 2309 | . | gene | MSTRG.28110 | 2.237928 | 0.02822 | 0.262004 | Bo8g081860.1 |
| 2308 | . | gene | MSTRG.31089 | 3.037236 | 0.028216 | 0.262004 | Bo9g055650.1 |
| 2311 | . | gene | MSTRG.4408 | 0.822907 | 0.028257 | 0.262051 | Bo2g045890.1 |
| 2312 | . | gene | MSTRG.33886 | 0.164619 | 0.02826 | 0.262051 | |
| 2313 | . | gene | MSTRG.34604 | 0.626956 | 0.028261 | 0.262051 | |
| 2314 | . | gene | MSTRG.28162 | 1.215409 | 0.028277 | 0.262079 | |
| 2315 | . | gene | MSTRG.19440 | 1.434046 | 0.028325 | 0.262311 | Bo5g141250.1 |
| 2316 | . | gene | MSTRG.15083 | 1.164924 | 0.028326 | 0.262311 | |

| | | | | | | | |
|---|---|---|---|---|---|---|---|
| 2323 | . | gene | MSTRG.8486 | 0.645239 | 0.028427 | 0.262453 | Bo3g035770.1 |
| 2321 | . | gene | MSTRG.11814 | 1.628444 | 0.028418 | 0.262453 | Bo3g172380.1 |
| 2319 | . | gene | MSTRG.17111 | 1.982972 | 0.02839 | 0.262453 | Bo5g027160.1 |
| 2320 | . | gene | MSTRG.20683 | 0.656008 | 0.028397 | 0.262453 | Bo6g061010.1 |
| 2318 | . | gene | MSTRG.29162 | 1.456812 | 0.028383 | 0.262453 | Bo8g105750.1 |
| 2317 | . | gene | MSTRG.34918 | 0.731043 | 0.028366 | 0.262453 | |
| 2322 | . | gene | MSTRG.26833 | 4.787341 | 0.028422 | 0.262453 | |
| 2325 | . | gene | MSTRG.24000 | 1.424982 | 0.02847 | 0.262624 | Bo7g063900.1 |
| 2324 | . | gene | MSTRG.27011 | 20.08131 | 0.028463 | 0.262624 | |
| 2330 | CLC-E | gene | MSTRG.255 | 0.940083 | 0.02854 | 0.262702 | Bo1g006450.1 |
| 2328 | . | gene | MSTRG.15567 | 1.847673 | 0.028521 | 0.262702 | Bo4g186850.1 |
| 2329 | . | gene | MSTRG.18912 | 2.498795 | 0.028536 | 0.262702 | Bo5g126830.1 |
| 2327 | BZR1 | gene | MSTRG.21493 | 1.320115 | 0.02851 | 0.262702 | Bo6g086560.1 |
| 2326 | . | gene | MSTRG.25289 | 1.683309 | 0.028493 | 0.262702 | Bo7g100850.1 |
| 2331 | . | gene | MSTRG.17611 | 0.089432 | 0.028567 | 0.262834 | |
| 2332 | . | gene | MSTRG.31946 | 1.381547 | 0.028588 | 0.262922 | Bo9g104800.1 |
| 2335 | . | gene | MSTRG.1916 | 0.785214 | 0.028636 | 0.262952 | Bo1g076870.1 |
| 2334 | . | gene | MSTRG.6913 | 1.475031 | 0.02863 | 0.262952 | Bo3g003500.1 |
| 2336 | . | gene | MSTRG.20887 | 0.605404 | 0.028641 | 0.262952 | Bo6g068060.1 |
| 2333 | . | gene | MSTRG.26183 | 1.324826 | 0.028616 | 0.262952 | Bo7g118340.1 |
| 2337 | . | gene | MSTRG.7394 | 1.465207 | 0.028671 | 0.263101 | Bo3g012860.1 |
| 2338 | . | gene | MSTRG.17158 | 1.157341 | 0.028681 | 0.263101 | Bo5g027800.1 |
| 2341 | . | gene | MSTRG.3006 | 1.549785 | 0.028746 | 0.263353 | Bo1g144240.1 |
| 2339 | . | gene | MSTRG.32921 | 1.60804 | 0.028726 | 0.263353 | Bo9g160110.1 |
| 2340 | . | gene | MSTRG.4541 | 0.776712 | 0.028743 | 0.263353 | |
| 2342 | . | gene | MSTRG.30882 | 4.368726 | 0.028758 | 0.263353 | Bo9g037250.1 |
| 2346 | . | gene | MSTRG.9499 | 1.714706 | 0.02884 | 0.263517 | Bo3g064440.1 |
| 2349 | . | gene | MSTRG.18873 | 0.507521 | 0.028873 | 0.263517 | Bo5g126270.1 |
| 2344 | . | gene | MSTRG.24142 | 1.327912 | 0.028807 | 0.263517 | Bo7g067210.1 |

| | | | | | | | |
|---|---|---|---|---|---|---|---|
| 2343 | . | gene | MSTRG.31855 | 0.773243 | 0.0288 | 0.263517 | Bo9g097500.1 |
| 2347 | . | gene | MSTRG.31883 | 1.385451 | 0.028859 | 0.263517 | Bo9g100050.1 |
| 2350 | . | gene | MSTRG.32099 | 0.35787 | 0.028874 | 0.263517 | Bo9g115100.1 |
| 2345 | . | gene | MSTRG.27398 | 0.654899 | 0.028835 | 0.263517 | |
| 2348 | . | gene | MSTRG.34048 | 2.520938 | 0.02886 | 0.263517 | |
| 2351 | . | gene | MSTRG.11788 | 0.616026 | 0.028925 | 0.26387 | Bo3g170990.1 |
| 2352 | . | gene | MSTRG.29330 | 0.775121 | 0.028952 | 0.264003 | Bo8g108620.1 |
| 2353 | . | gene | MSTRG.35142 | 0.894107 | 0.029012 | 0.264441 | |
| 2354 | . | gene | MSTRG.7476 | 18.82444 | 0.029025 | 0.264443 | Bo3g014030.1 |
| 2355 | . | gene | MSTRG.3080 | 0.764175 | 0.029067 | 0.26465 | Bo1g147290.1 |
| 2356 | . | gene | MSTRG.30114 | 0.569757 | 0.029072 | 0.26465 | Bo9g010650.1 |
| 2357 | . | gene | MSTRG.30683 | 0.431717 | 0.029099 | 0.264784 | Bo9g028540.1 |
| 2358 | . | gene | MSTRG.14151 | 0.936166 | 0.029118 | 0.264837 | Bo4g119660.1 |
| 2359 | . | gene | MSTRG.28367 | 4.237996 | 0.029191 | 0.26529 | Bo8g090530.1 |
| 2360 | . | gene | MSTRG.12518 | 0.167588 | 0.029192 | 0.26529 | |
| 2361 | . | gene | MSTRG.35017 | 1.142296 | 0.029236 | 0.26558 | |
| 2362 | . | gene | MSTRG.34438 | 0.586849 | 0.029252 | 0.265613 | |
| 2363 | . | gene | MSTRG.7898 | 3.414261 | 0.02928 | 0.265747 | |
| 2364 | . | gene | MSTRG.1955 | 1.759634 | 0.02931 | 0.265907 | Bo1g078690.1 |
| 2365 | . | gene | MSTRG.3879 | 0.752931 | 0.029337 | 0.266047 | Bo2g017800.1 |
| 2366 | . | gene | MSTRG.34343 | 2.274181 | 0.029382 | 0.266337 | |
| 2369 | . | gene | MSTRG.10061 | 0.462678 | 0.02943 | 0.266437 | Bo3g084000.1 |
| 2367 | . | gene | MSTRG.13496 | 1.558758 | 0.029414 | 0.266437 | |
| 2368 | . | gene | MSTRG.13772 | 1.434667 | 0.029429 | 0.266437 | |
| 2370 | . | gene | MSTRG.35877 | 0.172175 | 0.029467 | 0.266657 | |
| 2371 | . | gene | MSTRG.34805 | 0.785344 | 0.029534 | 0.267155 | |
| 2372 | HPY2 | gene | MSTRG.18952 | 1.275802 | 0.029551 | 0.267197 | Bo5g127420.1 |
| 2373 | . | gene | MSTRG.4189 | 28.16827 | 0.029622 | 0.267597 | Bo2g028810.1 |
| 2374 | . | gene | MSTRG.8183 | 0.684025 | 0.029624 | 0.267597 | Bo3g028200.1 |

| | | | | | | | |
|---|---|---|---|---|---|---|---|
| 2375 | . | gene | MSTRG.21868 | 0.264307 | 0.029633 | 0.267597 | Bo6g101240.1 |
| 2376 | . | gene | MSTRG.2277 | 0.089706 | 0.029646 | 0.267598 | |
| 2377 | . | gene | MSTRG.11262 | 1.130104 | 0.029677 | 0.267764 | |
| 2378 | . | gene | MSTRG.5808 | 0.873058 | 0.029719 | 0.267889 | Bo2g129860.1 |
| 2379 | . | gene | MSTRG.9091 | 2.636749 | 0.029722 | 0.267889 | Bo3g054400.1 |
| 2380 | . | gene | MSTRG.35088 | 0.284548 | 0.029728 | 0.267889 | |
| 2382 | . | gene | MSTRG.2407 | 1.233827 | 0.029776 | 0.267999 | Bo1g107070.1 |
| 2381 | . | gene | MSTRG.18954 | 0.739851 | 0.029762 | 0.267999 | Bo5g127450.1 |
| 2383 | . | gene | MSTRG.24768 | 1.552171 | 0.029787 | 0.267999 | Bo7g089020.1 |
| 2384 | . | gene | MSTRG.10822 | 2.070023 | 0.029797 | 0.267999 | |
| 2385 | . | gene | MSTRG.34556 | 0.576444 | 0.029803 | 0.267999 | |
| 2386 | . | gene | MSTRG.35470 | 0.071851 | 0.029877 | 0.268558 | |
| 2387 | . | gene | MSTRG.5530 | 0.499853 | 0.029903 | 0.26868 | |
| 2388 | ILL6 | gene | MSTRG.19841 | 1.612188 | 0.02992 | 0.268716 | Bo6g003330.1 |
| 2413 | . | gene | MSTRG.3920 | 1.303933 | 0.030252 | 0.268887 | Bo2g018510.1 |
| 2398 | . | gene | MSTRG.6644 | 1.426124 | 0.030104 | 0.268887 | Bo2g165730.1 |
| 2409 | . | gene | MSTRG.10420 | 1.293946 | 0.030213 | 0.268887 | Bo3g100620.1 |
| 2399 | . | gene | MSTRG.10455 | 0.362901 | 0.03011 | 0.268887 | Bo3g101180.1 |
| 2405 | . | gene | MSTRG.11093 | 0.934565 | 0.030167 | 0.268887 | Bo3g139880.1 |
| 2390 | . | gene | MSTRG.11330 | 1.462713 | 0.029989 | 0.268887 | Bo3g149520.1 |
| 2389 | . | gene | MSTRG.11840 | 1.219895 | 0.029973 | 0.268887 | Bo3g173740.1 |
| 2391 | . | gene | MSTRG.12253 | 2.351741 | 0.030017 | 0.268887 | Bo4g007090.1 |
| 2397 | . | gene | MSTRG.14192 | 1.449694 | 0.030098 | 0.268887 | Bo4g120230.1 |
| 2411 | . | gene | MSTRG.15337 | 0.215685 | 0.030237 | 0.268887 | Bo4g178120.1 |
| 2393 | . | gene | MSTRG.15531 | 0.620671 | 0.030054 | 0.268887 | Bo4g186260.1 |
| 2401 | . | gene | MSTRG.17340 | 1.251866 | 0.030155 | 0.268887 | Bo5g037410.1 |
| 2408 | . | gene | MSTRG.17666 | 1.735456 | 0.03021 | 0.268887 | Bo5g049530.1 |
| 2403 | . | gene | MSTRG.23219 | 1.5308 | 0.030164 | 0.268887 | Bo7g026760.1 |
| 2404 | . | gene | MSTRG.23393 | 0.402207 | 0.030166 | 0.268887 | Bo7g038690.1 |

| | | | | | | | |
|---|---|---|---|---|---|---|---|
| 2410 | . | gene | MSTRG.24431 | 1.554776 | 0.030221 | 0.268887 | Bo7g078900.1 |
| 2406 | . | gene | MSTRG.28095 | 2.016935 | 0.030186 | 0.268887 | Bo8g081580.1 |
| 2392 | . | gene | MSTRG.31073 | 0.749895 | 0.030036 | 0.268887 | Bo9g053410.1 |
| 2394 | . | gene | MSTRG.33429 | 1.569819 | 0.030056 | 0.268887 | Bo9g174270.1 |
| 2395 | . | gene | MSTRG.18874 | 0.513714 | 0.030089 | 0.268887 | |
| 2396 | . | gene | MSTRG.15687 | 0.603647 | 0.030092 | 0.268887 | |
| 2400 | . | gene | MSTRG.18212 | 1.680038 | 0.030151 | 0.268887 | |
| 2402 | . | gene | MSTRG.35248 | 1.862729 | 0.030159 | 0.268887 | |
| 2407 | . | gene | MSTRG.26591 | 0.643005 | 0.030203 | 0.268887 | |
| 2412 | . | gene | MSTRG.35104 | 0.556691 | 0.030251 | 0.268887 | |
| 2417 | . | gene | MSTRG.7370 | 0.730436 | 0.030303 | 0.268893 | Bo3g012420.1 |
| 2415 | . | gene | MSTRG.15909 | 0.849701 | 0.030297 | 0.268893 | Bo4g195350.1 |
| 2414 | . | gene | MSTRG.20897 | 0.314629 | 0.030285 | 0.268893 | |
| 2416 | . | gene | MSTRG.36088 | 1.671936 | 0.030303 | 0.268893 | |
| 2419 | . | gene | MSTRG.3665 | 1.454749 | 0.030354 | 0.269124 | Bo2g011310.1 |
| 2418 | . | gene | MSTRG.28559 | 2.878572 | 0.030352 | 0.269124 | Bo8g094330.1 |
| 2420 | . | gene | MSTRG.32707 | 1.581068 | 0.030384 | 0.269272 | Bo9g151850.1 |
| 2421 | . | gene | MSTRG.8014 | 0.20319 | 0.030421 | 0.269398 | Bo3g024820.1 |
| 2423 | . | gene | MSTRG.16891 | 1.471543 | 0.030445 | 0.269398 | Bo5g020800.1 |
| 2422 | . | gene | MSTRG.20357 | 0.922498 | 0.030429 | 0.269398 | Bo6g037100.1 |
| 2424 | . | gene | MSTRG.31172 | 12.33398 | 0.030448 | 0.269398 | Bo9g058960.1 |
| 2426 | . | gene | MSTRG.7544 | 3.34973 | 0.030489 | 0.269534 | Bo3g016940.1 |
| 2425 | . | gene | MSTRG.29227 | 0.449732 | 0.030486 | 0.269534 | Bo8g106710.1 |
| 2427 | . | gene | MSTRG.32867 | 1.591825 | 0.030509 | 0.269605 | Bo9g157370.1 |
| 2428 | . | gene | MSTRG.3429 | 0.161655 | 0.030632 | 0.270377 | Bo2g006620.1 |
| 2430 | . | gene | MSTRG.10754 | 0.652428 | 0.030634 | 0.270377 | Bo3g114950.1 |
| 2429 | . | gene | MSTRG.35447 | 0.36873 | 0.030633 | 0.270377 | |
| 2432 | . | gene | MSTRG.1350 | 0.762386 | 0.03071 | 0.270793 | Bo1g040550.1 |
| 2433 | . | gene | MSTRG.16216 | 1.558128 | 0.030719 | 0.270793 | Bo5g004960.1 |

| | | | | | | | |
|---|---|---|---|---|---|---|---|
| 24<br>31 | . | gene | MSTRG.1660<br>6 | 1.5139<br>7 | 0.0307<br>1 | 0.2707<br>93 | Bo5g0132<br>60.1 |
| 24<br>34 | . | gene | MSTRG.3273<br>0 | 1.2384<br>14 | 0.0307<br>79 | 0.2712<br>05 | Bo9g1521<br>70.1 |
| 24<br>35 | RAB1C | gene | MSTRG.764 | 1.8382<br>17 | 0.0308<br>16 | 0.2714<br>14 | Bo1g0176<br>20.1 |
| 24<br>39 | PRORP3 | gene | MSTRG.1030 | 1.4314<br>62 | 0.0308<br>66 | 0.2714<br>14 | Bo1g0257<br>20.1 |
| 24<br>36 | . | gene | MSTRG.1779<br>1 | 1.5257<br>56 | 0.0308<br>39 | 0.2714<br>14 | Bo5g0586<br>90.1 |
| 24<br>37 | . | gene | MSTRG.2981<br>3 | 1.8275<br>93 | 0.0308<br>55 | 0.2714<br>14 | Bo9g0041<br>00.1 |
| 24<br>38 | . | gene | MSTRG.3217<br>9 | 0.7849<br>19 | 0.0308<br>61 | 0.2714<br>14 | Bo9g1204<br>30.1 |
| 24<br>40 | . | gene | MSTRG.3338<br>2 | 1.5799<br>41 | 0.0308<br>93 | 0.2715<br>44 | Bo9g1735<br>80.1 |
| 24<br>41 | . | gene | MSTRG.2811<br>7 | 0.7538<br>47 | 0.0309<br>11 | 0.2715<br>93 | Bo8g0819<br>90.1 |
| 24<br>42 | MEE14 | gene | MSTRG.1018<br>9 | 0.4445<br>2 | 0.0309<br>44 | 0.2717<br>64 | Bo3g0899<br>80.1 |
| 24<br>47 | . | gene | MSTRG.7422 | 0.6488<br>28 | 0.0310<br>25 | 0.2719<br>21 | Bo3g0133<br>40.1 |
| 24<br>46 | . | gene | MSTRG.7950 | 2.2608<br>23 | 0.0310<br>17 | 0.2719<br>21 | Bo3g0238<br>30.1 |
| 24<br>44 | . | gene | MSTRG.1228<br>3 | 1.1525<br>08 | 0.0310<br>01 | 0.2719<br>21 | Bo4g0114<br>70.1 |
| 24<br>43 | . | gene | MSTRG.2551<br>5 | 1.4373<br>54 | 0.0310<br>01 | 0.2719<br>21 | Bo7g1065<br>20.1 |
| 24<br>45 | . | gene | MSTRG.3491<br>3 | 1.0891<br>05 | 0.0310<br>03 | 0.2719<br>21 | |
| 24<br>49 | . | gene | MSTRG.8142 | 1.3325<br>17 | 0.0310<br>83 | 0.2721<br>13 | Bo3g0278<br>20.1 |
| 24<br>48 | . | gene | MSTRG.2507<br>0 | 1.6398<br>75 | 0.0310<br>72 | 0.2721<br>13 | Bo7g0965<br>70.1 |
| 24<br>50 | . | gene | MSTRG.2629<br>3 | 2.0354<br>98 | 0.0310<br>85 | 0.2721<br>13 | |
| 24<br>51 | . | gene | MSTRG.6436 | 2.3029<br>86 | 0.0311<br>62 | 0.2725<br>95 | Bo2g1562<br>70.1 |
| 24<br>52 | FRS5 | gene | MSTRG.8397 | 0.6445<br>99 | 0.0311<br>65 | 0.2725<br>95 | Bo3g0342<br>90.1 |
| 24<br>57 | . | gene | MSTRG.2170 | 0.2741<br>34 | 0.0312<br>29 | 0.2725<br>96 | Bo1g0947<br>10.1 |
| 24<br>54 | . | gene | MSTRG.1034<br>7 | 0.2954<br>37 | 0.0311<br>95 | 0.2725<br>96 | Bo3g0977<br>30.1 |
| 24<br>56 | . | gene | MSTRG.2257<br>0 | 1.3176<br>03 | 0.0312<br>25 | 0.2725<br>96 | Bo6g1211<br>40.1 |
| 24<br>55 | . | gene | MSTRG.2536<br>6 | 1.2605<br>64 | 0.0312<br>07 | 0.2725<br>96 | Bo7g1041<br>00.1 |
| 24<br>53 | . | gene | MSTRG.3333<br>8 | 0.1194<br>43 | 0.0311<br>81 | 0.2725<br>96 | Bo9g1720<br>50.1 |
| 24<br>58 | . | gene | MSTRG.1327<br>8 | 1.4695<br>33 | 0.0312<br>68 | 0.2726<br>93 | Bo4g0539<br>80.1 |
| 24<br>59 | . | gene | MSTRG.1601<br>9 | 0.6763<br>55 | 0.0312<br>75 | 0.2726<br>93 | |
| 24<br>60 | . | gene | MSTRG.2769<br>8 | 0.4252<br>53 | 0.0312<br>78 | 0.2726<br>93 | |
| 24<br>62 | . | gene | MSTRG.1116<br>0 | 1.4784<br>45 | 0.0313<br>27 | 0.2728<br>38 | Bo3g1429<br>40.1 |

| | | | | | | | |
|---|---|---|---|---|---|---|---|
| 24<br>61 | . | gene | MSTRG.1505<br>9 | 1.3252<br>25 | 0.0313<br>14 | 0.2728<br>38 | Bo4g1612<br>20.1 |
| 24<br>64 | . | gene | MSTRG.1942<br>5 | 1.4967<br>74 | 0.0313<br>46 | 0.2728<br>38 | Bo5g1400<br>80.1 |
| 24<br>63 | . | gene | MSTRG.2000<br>5 | 1.2088<br>54 | 0.0313<br>36 | 0.2728<br>38 | |
| 24<br>65 | . | gene | MSTRG.2189<br>8 | 0.5864<br>79 | 0.0313<br>59 | 0.2728<br>45 | Bo6g1037<br>50.1 |
| 24<br>66 | . | gene | MSTRG.1250 | 0.2699<br>62 | 0.0313<br>77 | 0.2728<br>85 | Bo1g0390<br>30.1 |
| 24<br>67 | . | gene | MSTRG.3263<br>5 | 0.8208<br>65 | 0.0314<br>02 | 0.2729<br>95 | |
| 24<br>68 | AGD4 | gene | MSTRG.1661<br>1 | 1.5975<br>03 | 0.0314<br>76 | 0.2734<br>56 | Bo5g0133<br>20.1 |
| 24<br>69 | . | gene | MSTRG.2188<br>9 | 1.5962<br>87 | 0.0314<br>81 | 0.2734<br>56 | Bo6g1036<br>60.1 |
| 24<br>70 | . | gene | MSTRG.1975<br>5 | 1.3868<br>4 | 0.0315<br>1 | 0.2734<br>95 | Bo5g1526<br>30.1 |
| 24<br>71 | . | gene | MSTRG.2083<br>8 | 1.1959<br>3 | 0.0315<br>1 | 0.2734<br>95 | Bo6g0674<br>70.1 |
| 24<br>72 | . | gene | MSTRG.2777<br>2 | 0.7247<br>63 | 0.0315<br>38 | 0.2736<br>23 | Bo8g0710<br>80.1 |
| 24<br>73 | . | gene | MSTRG.3409<br>0 | 1.4541<br>95 | 0.0315<br>55 | 0.2736<br>59 | |
| 24<br>85 | . | gene | MSTRG.3663 | 1.2610<br>51 | 0.0317<br>13 | 0.2737<br>01 | Bo2g0112<br>90.1 |
| 24<br>77 | . | gene | MSTRG.6300 | 1.5620<br>36 | 0.0316<br>39 | 0.2737<br>01 | Bo2g1502<br>00.1 |
| 24<br>74 | . | gene | MSTRG.7335 | 1.6910<br>59 | 0.0315<br>75 | 0.2737<br>01 | Bo3g0109<br>30.1 |
| 24<br>81 | . | gene | MSTRG.9219 | 4.3485<br>87 | 0.0316<br>81 | 0.2737<br>01 | Bo3g0571<br>70.1 |
| 24<br>76 | . | gene | MSTRG.2580<br>1 | 5.9879<br>07 | 0.0316<br>1 | 0.2737<br>01 | Bo7g1113<br>40.1 |
| 24<br>82 | . | gene | MSTRG.2866<br>9 | 0.8441<br>38 | 0.0316<br>92 | 0.2737<br>01 | Bo8g0974<br>10.1 |
| 24<br>75 | . | gene | MSTRG.1627<br>6 | 0.9002<br>45 | 0.0316<br>09 | 0.2737<br>01 | |
| 24<br>78 | . | gene | MSTRG.2169 | 1.3977<br>12 | 0.0316<br>51 | 0.2737<br>01 | |
| 24<br>79 | . | gene | MSTRG.2098<br>1 | 0.8138<br>05 | 0.0316<br>54 | 0.2737<br>01 | |
| 24<br>80 | . | gene | MSTRG.1824<br>1 | 1.5912<br>12 | 0.0316<br>54 | 0.2737<br>01 | |
| 24<br>83 | . | gene | MSTRG.849 | 0.3173<br>95 | 0.0316<br>96 | 0.2737<br>01 | |
| 24<br>84 | . | gene | MSTRG.1678<br>6 | 0.9925<br>97 | 0.0317<br>03 | 0.2737<br>01 | |
| 24<br>86 | ATNADP-ME3 | gene | MSTRG.2501<br>2 | 1.5457<br>56 | 0.0317<br>4 | 0.2738<br>23 | Bo7g0956<br>60.1 |
| 24<br>87 | . | gene | MSTRG.3239<br>5 | 0.8473<br>66 | 0.0318<br>52 | 0.2746<br>77 | Bo9g1336<br>00.1 |
| 24<br>89 | . | gene | MSTRG.2176 | 0.6967<br>18 | 0.0319<br>19 | 0.2749<br>78 | Bo1g0979<br>90.1 |
| 24<br>99 | . | gene | MSTRG.4572 | 0.8582<br>17 | 0.0320<br>73 | 0.2749<br>78 | Bo2g0528<br>20.1 |
| 24<br>91 | . | gene | MSTRG.6096 | 0.8899<br>81 | 0.0319<br>7 | 0.2749<br>78 | Bo2g1391<br>20.1 |

| | | | | | | | |
|---|---|---|---|---|---|---|---|
| 2496 | . | gene | MSTRG.6904 | 2.651735 | 0.032055 | 0.274978 | Bo3g003360.1 |
| 2498 | . | gene | MSTRG.12267 | 1.638142 | 0.032072 | 0.274978 | Bo4g009240.1 |
| 2503 | . | gene | MSTRG.12977 | 0.182224 | 0.032123 | 0.274978 | Bo4g039330.1 |
| 2502 | . | gene | MSTRG.13473 | 0.579133 | 0.032114 | 0.274978 | Bo4g074160.1 |
| 2510 | . | gene | MSTRG.14516 | 0.051013 | 0.032181 | 0.274978 | Bo4g137050.1 |
| 2492 | . | gene | MSTRG.14881 | 1.653328 | 0.031976 | 0.274978 | Bo4g151320.1 |
| 2488 | ATCRK3 | gene | MSTRG.15976 | 1.421439 | 0.0319 | 0.274978 | Bo4g197280.1 |
| 2507 | . | gene | MSTRG.17857 | 0.519575 | 0.032161 | 0.274978 | Bo5g062790.1 |
| 2495 | . | gene | MSTRG.18851 | 1.558624 | 0.032013 | 0.274978 | Bo5g124870.1 |
| 2497 | ALDH22A1 | gene | MSTRG.19524 | 1.302572 | 0.032065 | 0.274978 | Bo5g143310.1 |
| 2505 | MKP2 | gene | MSTRG.19556 | 1.416306 | 0.032143 | 0.274978 | Bo5g144760.1 |
| 2490 | . | gene | MSTRG.20455 | 0.591406 | 0.031927 | 0.274978 | Bo6g044290.1 |
| 2509 | . | gene | MSTRG.23355 | 1.350017 | 0.032179 | 0.274978 | Bo7g033200.1 |
| 2493 | . | gene | MSTRG.28455 | 1.944413 | 0.031978 | 0.274978 | Bo8g091730.1 |
| 2494 | . | gene | MSTRG.28468 | 0.916459 | 0.032011 | 0.274978 | Bo8g091980.1 |
| 2504 | . | gene | MSTRG.32066 | 1.674695 | 0.032129 | 0.274978 | Bo9g111430.1 |
| 2506 | . | gene | MSTRG.32245 | 0.430181 | 0.032153 | 0.274978 | Bo9g123350.1 |
| 2500 | . | gene | MSTRG.31921 | 0.651974 | 0.032081 | 0.274978 | |
| 2501 | . | gene | MSTRG.18096 | 0.649185 | 0.032096 | 0.274978 | |
| 2508 | . | gene | MSTRG.4667 | 0.733538 | 0.032163 | 0.274978 | |
| 2512 | . | gene | MSTRG.13520 | 0.936295 | 0.032212 | 0.275023 | Bo4g076880.1 |
| 2511 | . | gene | MSTRG.34219 | 0.805929 | 0.032212 | 0.275023 | |
| 2513 | . | gene | MSTRG.23460 | 1.281675 | 0.032248 | 0.275221 | Bo7g040640.1 |
| 2514 | . | gene | MSTRG.23298 | 0.655391 | 0.032276 | 0.27535 | Bo7g031280.1 |
| 2515 | . | gene | MSTRG.28374 | 1.601682 | 0.03229 | 0.275358 | Bo8g090610.1 |
| 2516 | . | gene | MSTRG.24030 | 0.36028 | 0.032305 | 0.275377 | Bo7g064470.1 |
| 2522 | . | gene | MSTRG.3043 | 0.725382 | 0.032417 | 0.275583 | Bo1g144750.1 |
| 2523 | . | gene | MSTRG.8479 | 1.22006 | 0.032419 | 0.275583 | Bo3g035710.1 |
| 2518 | LEW3 | gene | MSTRG.12614 | 0.84139 | 0.032385 | 0.275583 | Bo4g025420.1 |

| | | | | | | | |
|---|---|---|---|---|---|---|---|
| 2519 | . | gene | MSTRG.16150 | 0.18136 | 0.032386 | 0.275583 | Bo5g003920.1 |
| 2520 | . | gene | MSTRG.22134 | 0.341825 | 0.032395 | 0.275583 | Bo6g111030.1 |
| 2517 | . | gene | MSTRG.33191 | 1.827036 | 0.032346 | 0.275583 | Bo9g167860.1 |
| 2521 | . | gene | MSTRG.5861 | 0.760448 | 0.032416 | 0.275583 | |
| 2524 | . | gene | MSTRG.18577 | 1.627109 | 0.032451 | 0.275743 | |
| 2525 | . | gene | MSTRG.6806 | 1.656727 | 0.032494 | 0.275956 | Bo3g001820.1 |
| 2526 | . | gene | MSTRG.20598 | 0.400022 | 0.032502 | 0.275956 | |
| 2527 | . | gene | MSTRG.157 | 1.675588 | 0.032527 | 0.276058 | |
| 2528 | . | gene | MSTRG.486 | 0.492343 | 0.032564 | 0.276219 | Bo1g010580.1 |
| 2530 | . | gene | MSTRG.19290 | 1.663175 | 0.032584 | 0.276219 | Bo5g137320.1 |
| 2529 | . | gene | MSTRG.25498 | 0.488926 | 0.032575 | 0.276219 | Bo7g106220.1 |
| 2531 | . | gene | MSTRG.20127 | 2.179403 | 0.032622 | 0.276426 | |
| 2532 | ROL5 | gene | MSTRG.12484 | 1.143886 | 0.032665 | 0.276686 | Bo4g021510.1 |
| 2533 | . | gene | MSTRG.15717 | 0.469739 | 0.032688 | 0.276772 | Bo4g190130.1 |
| 2536 | . | gene | MSTRG.15329 | 1.381669 | 0.032758 | 0.277027 | Bo4g177920.1 |
| 2535 | . | gene | MSTRG.22969 | 1.528585 | 0.032751 | 0.277027 | Bo7g010170.1 |
| 2534 | . | gene | MSTRG.25313 | 1.503872 | 0.032746 | 0.277027 | Bo7g101270.1 |
| 2537 | . | gene | MSTRG.35741 | 0.482301 | 0.03277 | 0.277027 | |
| 2538 | . | gene | MSTRG.7655 | 0.402724 | 0.032835 | 0.277421 | Bo3g018320.1 |
| 2539 | . | gene | MSTRG.16883 | 1.242535 | 0.032842 | 0.277421 | Bo5g020720.1 |
| 2540 | . | gene | MSTRG.15836 | 0.678398 | 0.032864 | 0.277467 | Bo4g192940.1 |
| 2541 | . | gene | MSTRG.8221 | 0.755695 | 0.032874 | 0.277467 | |
| 2542 | . | gene | MSTRG.22537 | 1.145014 | 0.032893 | 0.277518 | Bo6g120760.1 |
| 2543 | . | gene | MSTRG.33510 | 0.361747 | 0.032914 | 0.277589 | Bo9g175300.1 |
| 2544 | . | gene | MSTRG.10615 | 0.95978 | 0.032986 | 0.278085 | Bo3g107490.1 |
| 2546 | . | gene | MSTRG.8106 | 1.582284 | 0.033035 | 0.278283 | Bo3g027260.1 |
| 2545 | . | gene | MSTRG.19690 | 7.352834 | 0.033033 | 0.278283 | Bo5g148720.1 |
| 2547 | . | gene | MSTRG.1870 | 1.631229 | 0.03305 | 0.278296 | Bo1g071110.1 |
| 2548 | . | gene | MSTRG.31863 | 1.648203 | 0.03308 | 0.278444 | Bo9g097610.1 |

| | | | | | | | |
|---|---|---|---|---|---|---|---|
| 25 49 | . | gene | MSTRG.1799 3 | 0.9221 03 | 0.0331 49 | 0.2789 11 | Bo5g0724 60.1 |
| 25 52 | . | gene | MSTRG.1254 0 | 1.8783 54 | 0.0332 03 | 0.2789 93 | Bo4g0244 00.1 |
| 25 53 | . | gene | MSTRG.2877 0 | 1.4697 26 | 0.0332 11 | 0.2789 93 | Bo8g0988 40.1 |
| 25 50 | . | gene | MSTRG.3119 5 | 2.4153 91 | 0.0331 81 | 0.2789 93 | Bo9g0593 80.1 |
| 25 51 | . | gene | MSTRG.1791 9 | 0.3441 39 | 0.0331 92 | 0.2789 93 | |
| 25 54 | . | gene | MSTRG.2496 2 | 0.8117 21 | 0.0332 44 | 0.2791 62 | Bo7g0948 90.1 |
| 25 55 | ATNRT1:2 | gene | MSTRG.2210 3 | 2.8350 05 | 0.0333 16 | 0.2796 61 | Bo6g1104 60.1 |
| 25 59 | . | gene | MSTRG.646 | 0.9125 37 | 0.0334 11 | 0.2799 12 | Bo1g0149 80.1 |
| 25 58 | . | gene | MSTRG.2256 3 | 0.4165 4 | 0.0334 09 | 0.2799 12 | Bo6g1210 40.1 |
| 25 60 | . | gene | MSTRG.2921 3 | 1.4165 32 | 0.0334 15 | 0.2799 12 | Bo8g1064 90.1 |
| 25 56 | . | gene | MSTRG.3078 9 | 0.8591 91 | 0.0333 88 | 0.2799 12 | Bo9g0322 70.1 |
| 25 57 | . | gene | MSTRG.2015 8 | 0.6797 17 | 0.0334 04 | 0.2799 12 | |
| 25 61 | . | gene | MSTRG.3528 0 | 0.5107 8 | 0.0334 25 | 0.2799 12 | |
| 25 62 | . | gene | MSTRG.2969 7 | 1.8304 23 | 0.0334 4 | 0.2799 33 | Bo8g1179 20.1 |
| 25 63 | . | gene | MSTRG.2033 0 | 0.7264 23 | 0.0334 55 | 0.2799 51 | Bo6g0344 70.1 |
| 25 65 | ARA7 | gene | MSTRG.908 | 1.8101 97 | 0.0334 87 | 0.2799 95 | Bo1g0217 30.1 |
| 25 64 | . | gene | MSTRG.3234 8 | 0.7974 14 | 0.0334 81 | 0.2799 95 | Bo9g1309 20.1 |
| 25 66 | ATMPK15 | gene | MSTRG.2155 1 | 0.9212 12 | 0.0335 04 | 0.2800 32 | Bo6g0876 00.1 |
| 25 70 | . | gene | MSTRG.1826 | 0.7050 66 | 0.0336 | 0.2801 3 | Bo1g0672 70.1 |
| 25 78 | . | gene | MSTRG.4077 | 0.6018 27 | 0.0337 | 0.2801 3 | Bo2g0249 40.1 |
| 25 86 | . | gene | MSTRG.4404 | 1.9074 58 | 0.0337 77 | 0.2801 3 | Bo2g0457 80.1 |
| 25 67 | . | gene | MSTRG.4956 | 0.2798 19 | 0.0335 49 | 0.2801 3 | Bo2g0769 30.1 |
| 25 68 | . | gene | MSTRG.5079 | 0.6859 98 | 0.0335 68 | 0.2801 3 | Bo2g0852 60.1 |
| 25 82 | . | gene | MSTRG.1138 5 | 0.6013 99 | 0.0337 45 | 0.2801 3 | Bo3g1514 10.1 |
| 25 84 | . | gene | MSTRG.1549 9 | 0.6023 22 | 0.0337 58 | 0.2801 3 | Bo4g1846 70.1 |
| 25 85 | . | gene | MSTRG.1557 0 | 2.9088 51 | 0.0337 76 | 0.2801 3 | Bo4g1868 90.1 |
| 25 81 | . | gene | MSTRG.1627 3 | 2.5693 17 | 0.0337 3 | 0.2801 3 | Bo5g0057 50.1 |
| 25 76 | CID11 | gene | MSTRG.1765 9 | 1.8294 17 | 0.0336 79 | 0.2801 3 | Bo5g0493 80.1 |
| 25 71 | . | gene | MSTRG.2014 0 | 1.4016 32 | 0.0336 12 | 0.2801 3 | Bo6g0264 70.1 |

| | | | | | | | |
|---|---|---|---|---|---|---|---|
| 25 79 | . | gene | MSTRG.2059 4 | 0.1847 59 | 0.0337 04 | 0.2801 3 | Bo6g0536 60.1 |
| 25 73 | . | gene | MSTRG.2314 1 | 0.8834 24 | 0.0336 37 | 0.2801 3 | Bo7g0213 10.1 |
| 25 74 | . | gene | MSTRG.2424 0 | 1.3589 62 | 0.0336 57 | 0.2801 3 | Bo7g0739 80.1 |
| 25 83 | . | gene | MSTRG.2775 7 | 3.2581 34 | 0.0337 53 | 0.2801 3 | Bo8g0709 00.1 |
| 25 80 | . | gene | MSTRG.3009 5 | 0.5524 | 0.0337 24 | 0.2801 3 | Bo9g0103 20.1 |
| 25 75 | . | gene | MSTRG.3207 9 | 1.7264 34 | 0.0336 58 | 0.2801 3 | Bo9g1126 70.1 |
| 25 69 | . | gene | MSTRG.4154 | 0.0907 31 | 0.0335 93 | 0.2801 3 | |
| 25 72 | . | gene | MSTRG.2054 4 | 0.5868 4 | 0.0336 28 | 0.2801 3 | |
| 25 77 | . | gene | MSTRG.3486 7 | 0.1741 28 | 0.0336 91 | 0.2801 3 | |
| 25 87 | . | gene | MSTRG.1136 7 | 0.3492 04 | 0.0338 61 | 0.2806 23 | Bo3g1500 40.1 |
| 25 88 | . | gene | MSTRG.3068 9 | 1.8473 36 | 0.0338 63 | 0.2806 23 | Bo9g0286 80.1 |
| 25 89 | . | gene | MSTRG.1528 6 | 0.4264 95 | 0.0339 | 0.2808 24 | Bo4g1762 00.1 |
| 25 90 | . | gene | MSTRG.1224 3 | 0.5621 34 | 0.0339 8 | 0.2813 74 | Bo4g0069 50.1 |
| 25 91 | . | gene | MSTRG.3369 | 0.6768 18 | 0.0340 13 | 0.2814 4 | Bo2g0027 50.1 |
| 25 92 | . | gene | MSTRG.1770 3 | 1.4131 12 | 0.0340 3 | 0.2814 4 | Bo5g0521 00.1 |
| 25 93 | . | gene | MSTRG.1774 1 | 0.7043 24 | 0.0340 44 | 0.2814 4 | Bo5g0557 50.1 |
| 25 95 | . | gene | MSTRG.2834 8 | 0.7374 96 | 0.0340 53 | 0.2814 4 | Bo8g0903 00.1 |
| 25 94 | . | gene | MSTRG.2857 7 | 1.2172 62 | 0.0340 52 | 0.2814 4 | |
| 25 96 | . | gene | MSTRG.2586 | 1.8502 63 | 0.0340 7 | 0.2814 74 | Bo1g1191 00.1 |
| 25 98 | . | gene | MSTRG.2866 7 | 1.3853 79 | 0.0341 04 | 0.2815 32 | Bo8g0973 60.1 |
| 25 97 | . | gene | MSTRG.2034 1 | 0.7528 | 0.0340 92 | 0.2815 32 | |
| 25 99 | TUB8 | gene | MSTRG.2512 5 | 1.3605 71 | 0.0341 36 | 0.2816 91 | Bo7g0974 20.1 |
| 26 00 | . | gene | MSTRG.1479 4 | 1.5606 | 0.0341 5 | 0.2816 95 | Bo4g1459 00.1 |
| 26 01 | . | gene | MSTRG.1599 7 | 1.4258 74 | 0.0342 04 | 0.2819 67 | Bo4g1986 00.1 |
| 26 02 | . | gene | MSTRG.1769 5 | 0.3960 21 | 0.0342 09 | 0.2819 67 | Bo5g0519 80.1 |
| 26 04 | . | gene | MSTRG.2980 | 2.1303 03 | 0.0342 54 | 0.2821 2 | Bo1g1437 90.1 |
| 26 05 | . | gene | MSTRG.4564 | 1.3970 87 | 0.0342 67 | 0.2821 2 | Bo2g0526 60.1 |
| 26 03 | . | gene | MSTRG.1831 8 | 0.4057 13 | 0.0342 51 | 0.2821 2 | Bo5g0957 20.1 |
| 26 07 | . | gene | MSTRG.1605 9 | 0.7257 19 | 0.0343 39 | 0.2824 72 | Bo5g0025 50.1 |

| | | | | | | | |
|---|---|---|---|---|---|---|---|
| 2606 | . | gene | MSTRG.36159 | 0.860619 | 0.034334 | 0.282472 | |
| 2608 | . | gene | MSTRG.7928 | 0.891893 | 0.034349 | 0.282472 | |
| 2609 | . | gene | MSTRG.16551 | 3.734289 | 0.034364 | 0.282482 | Bo5g010420.1 |
| 2610 | . | gene | MSTRG.8159 | 1.083263 | 0.034386 | 0.282557 | Bo3g027980.1 |
| 2611 | . | gene | MSTRG.7328 | 2.239015 | 0.034423 | 0.282756 | Bo3g010850.1 |
| 2613 | . | gene | MSTRG.9808 | 0.559691 | 0.034476 | 0.282975 | Bo3g071980.1 |
| 2612 | . | gene | MSTRG.22931 | 2.15993 | 0.034473 | 0.282975 | Bo7g008350.1 |
| 2614 | . | gene | MSTRG.5244 | 0.363763 | 0.034524 | 0.283184 | Bo2g094850.1 |
| 2616 | . | gene | MSTRG.17218 | 1.702765 | 0.034541 | 0.283184 | Bo5g030580.1 |
| 2615 | . | gene | MSTRG.17512 | 0.706424 | 0.034531 | 0.283184 | |
| 2631 | . | gene | MSTRG.471 | 1.529929 | 0.034785 | 0.283447 | Bo1g010380.1 |
| 2623 | . | gene | MSTRG.2208 | 2.120825 | 0.034701 | 0.283447 | Bo1g098520.1 |
| 2622 | . | gene | MSTRG.2727 | 1.890046 | 0.034677 | 0.283447 | Bo1g128550.1 |
| 2630 | . | gene | MSTRG.2841 | 0.297673 | 0.034771 | 0.283447 | Bo1g138630.1 |
| 2621 | . | gene | MSTRG.2882 | 0.356266 | 0.034656 | 0.283447 | Bo1g139350.1 |
| 2617 | . | gene | MSTRG.2957 | 0.113198 | 0.034605 | 0.283447 | Bo1g141390.1 |
| 2625 | . | gene | MSTRG.6614 | 0.720517 | 0.034713 | 0.283447 | Bo2g164190.1 |
| 2619 | . | gene | MSTRG.11130 | 1.115781 | 0.034643 | 0.283447 | Bo3g141540.1 |
| 2629 | . | gene | MSTRG.21244 | 0.313952 | 0.03477 | 0.283447 | Bo6g079420.1 |
| 2620 | . | gene | MSTRG.24418 | 0.27375 | 0.034648 | 0.283447 | Bo7g078740.1 |
| 2632 | PAF2 | gene | MSTRG.26612 | 0.837983 | 0.034785 | 0.283447 | Bo8g019190.1 |
| 2627 | . | gene | MSTRG.30686 | 1.388276 | 0.03473 | 0.283447 | Bo9g028610.1 |
| 2618 | . | gene | MSTRG.32239 | 0.625044 | 0.034637 | 0.283447 | Bo9g122190.1 |
| 2624 | . | gene | MSTRG.8801 | 0.122563 | 0.034711 | 0.283447 | |
| 2626 | . | gene | MSTRG.9924 | 0.430179 | 0.034717 | 0.283447 | |
| 2628 | . | gene | MSTRG.1615 | 1.328763 | 0.034737 | 0.283447 | |
| 2634 | . | gene | MSTRG.3052 | 0.545932 | 0.034901 | 0.284003 | Bo1g144900.1 |
| 2635 | . | gene | MSTRG.7007 | 1.331464 | 0.034912 | 0.284003 | Bo3g005050.1 |
| 2636 | . | gene | MSTRG.20003 | 0.869056 | 0.034961 | 0.284003 | Bo6g018340.1 |

| | | | | | | | |
|---|---|---|---|---|---|---|---|
| 26 41 | . | gene | MSTRG.2143 2 | 0.6494 47 | 0.0349 72 | 0.2840 03 | Bo6g0835 80.1 |
| 26 39 | . | gene | MSTRG.2182 8 | 1.4493 58 | 0.0349 7 | 0.2840 03 | Bo6g0995 80.1 |
| 26 33 | . | gene | MSTRG.2391 8 | 2.3998 37 | 0.0348 96 | 0.2840 03 | Bo7g0613 90.1 |
| 26 37 | . | gene | MSTRG.2563 4 | 1.3690 62 | 0.0349 66 | 0.2840 03 | Bo7g1087 90.1 |
| 26 38 | . | gene | MSTRG.3506 5 | 2.1126 8 | 0.0349 66 | 0.2840 03 | |
| 26 40 | . | gene | MSTRG.7870 | 0.4656 81 | 0.0349 71 | 0.2840 03 | |
| 26 42 | . | gene | MSTRG.8000 | 0.6332 16 | 0.0350 46 | 0.2841 01 | Bo3g0246 10.1 |
| 26 46 | . | gene | MSTRG.1667 9 | 0.7024 09 | 0.0350 57 | 0.2841 01 | Bo5g0163 50.1 |
| 26 47 | CARB | gene | MSTRG.1781 3 | 1.5450 78 | 0.0350 64 | 0.2841 01 | Bo5g0611 00.1 |
| 26 44 | PAD2 | gene | MSTRG.3048 8 | 2.0852 21 | 0.0350 54 | 0.2841 01 | Bo9g0221 40.1 |
| 26 43 | . | gene | gene:Bo6g03 3160 | 0.7086 38 | 0.0350 48 | 0.2841 01 | |
| 26 45 | . | gene | MSTRG.2166 5 | 0.5582 58 | 0.0350 56 | 0.2841 01 | |
| 26 48 | . | gene | MSTRG.2678 8 | 0.6549 68 | 0.0350 92 | 0.2842 19 | |
| 26 49 | . | gene | MSTRG.1530 5 | 0.3967 97 | 0.0351 2 | 0.2842 84 | Bo4g1775 00.1 |
| 26 50 | . | gene | MSTRG.3075 1 | 1.3238 32 | 0.0351 38 | 0.2842 84 | Bo9g0306 00.1 |
| 26 51 | . | gene | MSTRG.3604 4 | 0.4733 04 | 0.0351 4 | 0.2842 84 | |
| 26 52 | . | gene | MSTRG.9107 | 1.6307 29 | 0.0351 53 | 0.2842 86 | Bo3g0545 80.1 |
| 26 53 | . | gene | MSTRG.3130 | 0.7188 43 | 0.0351 85 | 0.2844 4 | Bo1g1500 60.1 |
| 26 54 | . | gene | MSTRG.6999 | 0.7766 59 | 0.0352 43 | 0.2847 14 | Bo3g0049 40.1 |
| 26 56 | . | gene | MSTRG.9554 | 0.8120 92 | 0.0352 59 | 0.2847 14 | Bo3g0651 20.1 |
| 26 55 | . | gene | MSTRG.1516 | 1.5776 49 | 0.0352 51 | 0.2847 14 | |
| 26 57 | . | gene | MSTRG.1927 8 | 1.1324 33 | 0.0353 1 | 0.2850 16 | Bo5g1371 10.1 |
| 26 58 | . | gene | MSTRG.6589 | 1.2582 43 | 0.0353 43 | 0.2851 78 | Bo2g1617 90.1 |
| 26 59 | . | gene | MSTRG.3176 | 1.4773 46 | 0.0353 64 | 0.2852 39 | Bo1g1518 80.1 |
| 26 60 | . | gene | MSTRG.1357 6 | 1.1928 32 | 0.0353 91 | 0.2853 54 | Bo4g0787 60.1 |
| 26 61 | . | gene | MSTRG.1475 4 | 0.4323 26 | 0.0354 4 | 0.2855 87 | Bo4g1452 60.1 |
| 26 62 | . | gene | MSTRG.1577 0 | 0.1244 14 | 0.0354 47 | 0.2855 87 | |
| 26 63 | . | gene | MSTRG.5851 | 0.3844 33 | 0.0355 14 | 0.2860 18 | |
| 26 64 | . | gene | MSTRG.1304 1 | 0.4078 88 | 0.0356 19 | 0.2867 59 | Bo4g0412 30.1 |

| | | | | | | | |
|---|---|---|---|---|---|---|---|
| 26<br>65 | . | gene | MSTRG.3374<br>3 | 1.4918<br>93 | 0.0356<br>35 | 0.2867<br>75 | Bo9g1817<br>90.1 |
| 26<br>66 | . | gene | MSTRG.2115<br>5 | 1.3062<br>33 | 0.0356<br>77 | 0.2870<br>11 | |
| 26<br>67 | . | gene | MSTRG.1972 | 0.5927<br>77 | 0.0357<br>39 | 0.2873<br>93 | Bo1g0789<br>50.1 |
| 26<br>68 | . | gene | MSTRG.1997<br>6 | 0.8054<br>96 | 0.0357<br>52 | 0.2873<br>93 | Bo6g0157<br>50.1 |
| 26<br>71 | . | gene | MSTRG.488 | 0.5624<br>7 | 0.0358<br>31 | 0.2876<br>55 | Bo1g0106<br>00.1 |
| 26<br>69 | . | gene | MSTRG.1132<br>5 | 1.5258<br>04 | 0.0357<br>98 | 0.2876<br>55 | Bo3g1494<br>60.1 |
| 26<br>70 | . | gene | MSTRG.2340<br>3 | 0.1572<br>68 | 0.0358<br>25 | 0.2876<br>55 | Bo7g0388<br>50.1 |
| 26<br>72 | . | gene | MSTRG.1850<br>4 | 0.7383<br>93 | 0.0358<br>38 | 0.2876<br>55 | |
| 26<br>99 | . | gene | MSTRG.2984 | 1.6498<br>89 | 0.0362<br>53 | 0.2878<br>44 | Bo1g1438<br>50.1 |
| 27<br>12 | . | gene | MSTRG.3229 | 1.1926<br>61 | 0.0364<br>18 | 0.2878<br>44 | Bo1g1536<br>00.1 |
| 27<br>01 | . | gene | MSTRG.4032 | 0.6788<br>76 | 0.0362<br>61 | 0.2878<br>44 | Bo2g0242<br>20.1 |
| 26<br>94 | ATRAB8C | gene | MSTRG.6847 | 1.3603<br>29 | 0.0361<br>93 | 0.2878<br>44 | Bo3g0023<br>80.1 |
| 27<br>07 | . | gene | MSTRG.7563 | 1.8434<br>77 | 0.0363<br>54 | 0.2878<br>44 | Bo3g0172<br>30.1 |
| 26<br>97 | . | gene | MSTRG.7771 | 1.5271<br>13 | 0.0362<br>35 | 0.2878<br>44 | Bo3g0210<br>00.1 |
| 26<br>92 | . | gene | MSTRG.8967 | 1.2177<br>5 | 0.0361<br>59 | 0.2878<br>44 | Bo3g0484<br>30.1 |
| 26<br>88 | . | gene | MSTRG.9603 | 1.2852<br>12 | 0.0361<br>08 | 0.2878<br>44 | Bo3g0658<br>30.1 |
| 27<br>06 | . | gene | MSTRG.1017<br>1 | 2.8474<br>28 | 0.0363<br>25 | 0.2878<br>44 | Bo3g0884<br>00.1 |
| 26<br>98 | . | gene | MSTRG.1057<br>5 | 1.8723<br>08 | 0.0362<br>36 | 0.2878<br>44 | Bo3g1058<br>00.1 |
| 27<br>03 | . | gene | MSTRG.1123<br>5 | 0.7592<br>35 | 0.0362<br>89 | 0.2878<br>44 | Bo3g1452<br>30.1 |
| 26<br>90 | LOL2 | gene | MSTRG.1159<br>3 | 1.2368<br>47 | 0.0361<br>3 | 0.2878<br>44 | Bo3g1624<br>70.1 |
| 26<br>83 | . | gene | MSTRG.1201<br>8 | 1.4432<br>37 | 0.0360<br>55 | 0.2878<br>44 | Bo3g1813<br>90.1 |
| 26<br>79 | . | gene | MSTRG.1536<br>7 | 1.3847<br>42 | 0.0360<br>13 | 0.2878<br>44 | Bo4g1816<br>10.1 |
| 27<br>04 | . | gene | MSTRG.1579<br>4 | 1.5990<br>43 | 0.0362<br>96 | 0.2878<br>44 | Bo4g1912<br>60.1 |
| 27<br>13 | . | gene | MSTRG.1661<br>0 | 0.7599<br>66 | 0.0364<br>23 | 0.2878<br>44 | Bo5g0133<br>10.1 |
| 27<br>05 | . | gene | MSTRG.1912<br>3 | 1.1164<br>59 | 0.0363<br>16 | 0.2878<br>44 | Bo5g1328<br>30.1 |
| 26<br>91 | MTO1 | gene | MSTRG.1981<br>4 | 1.3963<br>44 | 0.0361<br>32 | 0.2878<br>44 | Bo5g1564<br>30.1 |
| 27<br>10 | . | gene | MSTRG.2245<br>8 | 1.3053<br>96 | 0.0364<br>11 | 0.2878<br>44 | Bo6g1197<br>70.1 |
| 26<br>78 | . | gene | MSTRG.2391<br>5 | 2.6407<br>19 | 0.0359<br>98 | 0.2878<br>44 | Bo7g0613<br>40.1 |
| 26<br>93 | . | gene | MSTRG.2392<br>0 | 0.3372<br>86 | 0.0361<br>73 | 0.2878<br>44 | Bo7g0614<br>40.1 |

| | | | | | | | |
|---|---|---|---|---|---|---|---|
| 2686 | . | gene | MSTRG.25722 | 2.199235 | 0.036089 | 0.287844 | Bo7g110080.1 |
| 2677 | . | gene | MSTRG.25926 | 0.844977 | 0.035968 | 0.287844 | Bo7g114460.1 |
| 2700 | . | gene | MSTRG.27574 | 0.792207 | 0.03626 | 0.287844 | Bo8g066850.1 |
| 2711 | . | gene | MSTRG.28077 | 0.859652 | 0.036415 | 0.287844 | Bo8g081330.1 |
| 2684 | . | gene | MSTRG.28631 | 0.709181 | 0.036072 | 0.287844 | Bo8g095800.1 |
| 2687 | . | gene | MSTRG.28959 | 1.268787 | 0.036089 | 0.287844 | Bo8g101760.1 |
| 2708 | . | gene | MSTRG.29867 | 1.36699 | 0.036382 | 0.287844 | Bo9g005780.1 |
| 2682 | . | gene | MSTRG.30230 | 0.441211 | 0.036053 | 0.287844 | Bo9g014400.1 |
| 2685 | . | gene | MSTRG.32024 | 0.244798 | 0.036086 | 0.287844 | Bo9g110890.1 |
| 2675 | . | gene | MSTRG.33428 | 1.598442 | 0.03591 | 0.287844 | Bo9g174260.1 |
| 2681 | . | gene | MSTRG.33589 | 1.344554 | 0.036042 | 0.287844 | Bo9g176520.1 |
| 2673 | . | gene | MSTRG.26484 | 2.135178 | 0.035887 | 0.287844 | |
| 2674 | . | gene | MSTRG.34926 | 2.256115 | 0.035909 | 0.287844 | |
| 2676 | . | gene | MSTRG.19342 | 0.085185 | 0.035947 | 0.287844 | |
| 2680 | . | gene | MSTRG.1990 | 1.040418 | 0.036025 | 0.287844 | |
| 2689 | . | gene | MSTRG.11105 | 0.049891 | 0.036116 | 0.287844 | |
| 2695 | . | gene | MSTRG.23424 | 0.64242 | 0.036205 | 0.287844 | |
| 2696 | . | gene | MSTRG.33807 | 0.271844 | 0.036213 | 0.287844 | |
| 2702 | . | gene | MSTRG.9995 | 0.417482 | 0.036281 | 0.287844 | |
| 2709 | . | gene | MSTRG.26854 | 1.445581 | 0.036394 | 0.287844 | |
| 2714 | . | gene | MSTRG.29552 | 0.755472 | 0.036425 | 0.287844 | |
| 2715 | . | gene | MSTRG.4749 | 0.050379 | 0.036457 | 0.287899 | Bo2g061420.1 |
| 2717 | . | gene | MSTRG.23107 | 0.504415 | 0.036475 | 0.287899 | Bo7g018700.1 |
| 2716 | DCL4 | gene | MSTRG.32715 | 0.712039 | 0.036466 | 0.287899 | Bo9g151990.1 |
| 2718 | . | gene | MSTRG.4948 | 1.372121 | 0.036486 | 0.287899 | |
| 2720 | . | gene | MSTRG.3007 | 1.389197 | 0.036523 | 0.287952 | Bo1g144250.1 |
| 2719 | . | gene | MSTRG.8517 | 0.929154 | 0.036517 | 0.287952 | Bo3g036080.1 |
| 2721 | . | gene | MSTRG.21101 | 1.142082 | 0.036533 | 0.287952 | Bo6g076350.1 |
| 2724 | . | gene | MSTRG.21763 | 0.258841 | 0.036578 | 0.287989 | Bo6g097400.1 |

| | | | | | | | |
|---|---|---|---|---|---|---|---|
| 2723 | . | gene | MSTRG.28966 | 0.51147 | 0.036576 | 0.287989 | Bo8g101850.1 |
| 2722 | . | gene | MSTRG.14299 | 0.87841 | 0.036554 | 0.287989 | |
| 2726 | . | gene | MSTRG.13259 | 1.403107 | 0.036628 | 0.288173 | Bo4g053680.1 |
| 2725 | . | gene | MSTRG.23602 | 0.580615 | 0.036622 | 0.288173 | Bo7g046950.1 |
| 2727 | . | gene | MSTRG.21042 | 0.375581 | 0.036659 | 0.288315 | Bo6g074580.1 |
| 2728 | . | gene | MSTRG.27592 | 0.840562 | 0.036695 | 0.288489 | Bo8g067120.1 |
| 2729 | . | gene | MSTRG.7990 | 0.60036 | 0.036751 | 0.288825 | Bo3g024410.1 |
| 2730 | . | gene | MSTRG.20562 | 0.722916 | 0.036817 | 0.289234 | Bo6g051120.1 |
| 2731 | . | gene | MSTRG.32941 | 1.732883 | 0.036838 | 0.289293 | Bo9g161430.1 |
| 2732 | . | gene | MSTRG.9519 | 0.820895 | 0.036868 | 0.289421 | Bo3g064720.1 |
| 2733 | . | gene | MSTRG.11770 | 1.436313 | 0.036958 | 0.289952 | Bo3g170710.1 |
| 2739 | AAP3 | gene | MSTRG.22582 | 0.243951 | 0.03703 | 0.289952 | Bo6g121280.1 |
| 2735 | . | gene | MSTRG.22743 | 0.537554 | 0.036983 | 0.289952 | Bo6g125600.1 |
| 2736 | . | gene | MSTRG.22798 | 1.396152 | 0.037002 | 0.289952 | Bo7g003570.1 |
| 2734 | . | gene | MSTRG.18386 | 0.477998 | 0.036968 | 0.289952 | |
| 2737 | . | gene | MSTRG.28300 | 0.722409 | 0.037007 | 0.289952 | |
| 2738 | . | gene | MSTRG.28023 | 3.665451 | 0.037026 | 0.289952 | |
| 2740 | . | gene | MSTRG.19614 | 0.573926 | 0.037056 | 0.289957 | Bo5g146610.1 |
| 2741 | . | gene | MSTRG.26663 | 2.830693 | 0.037058 | 0.289957 | Bo8g023090.1 |
| 2742 | . | gene | MSTRG.3476 | 0.206024 | 0.03714 | 0.290466 | Bo2g007510.1 |
| 2743 | . | gene | MSTRG.16098 | 0.693184 | 0.03715 | 0.290466 | Bo5g003200.1 |
| 2744 | . | gene | MSTRG.15838 | 0.56117 | 0.037184 | 0.290613 | Bo4g192980.1 |
| 2745 | . | gene | MSTRG.19252 | 0.630385 | 0.037195 | 0.290613 | Bo5g136780.1 |
| 2746 | . | gene | MSTRG.17910 | 0.42929 | 0.037229 | 0.290769 | Bo5g065760.1 |
| 2747 | . | gene | MSTRG.10041 | 2.298269 | 0.037254 | 0.29086 | |
| 2749 | . | gene | MSTRG.5957 | 1.156829 | 0.037338 | 0.291281 | Bo2g134480.1 |
| 2751 | . | gene | MSTRG.7974 | 0.757672 | 0.037363 | 0.291281 | Bo3g024180.1 |
| 2748 | . | gene | MSTRG.8032 | 0.340703 | 0.037328 | 0.291281 | Bo3g025130.1 |
| 2750 | . | gene | MSTRG.27653 | 6.326698 | 0.037356 | 0.291281 | Bo8g068200.1 |

| | | | | | | | |
|---|---|---|---|---|---|---|---|
| 2752 | . | gene | MSTRG.6631 | 0.391037 | 0.03742 | 0.291584 | Bo2g165450.1 |
| 2753 | . | gene | MSTRG.8681 | 1.322161 | 0.037429 | 0.291584 | Bo3g039560.1 |
| 2757 | . | gene | MSTRG.6598 | 0.044085 | 0.037484 | 0.291592 | Bo2g162920.1 |
| 2754 | . | gene | MSTRG.26250 | 1.571865 | 0.037444 | 0.291592 | Bo7g119290.1 |
| 2755 | . | gene | MSTRG.4551 | 0.112231 | 0.037478 | 0.291592 | |
| 2756 | . | gene | MSTRG.35371 | 0.30313 | 0.03748 | 0.291592 | |
| 2759 | . | gene | MSTRG.11604 | 2.241913 | 0.037522 | 0.291679 | Bo3g162720.1 |
| 2758 | . | gene | MSTRG.10289 | 1.834958 | 0.037518 | 0.291679 | |
| 2761 | . | gene | MSTRG.24850 | 0.404339 | 0.03755 | 0.291683 | Bo7g091160.1 |
| 2760 | . | gene | MSTRG.32016 | 2.344259 | 0.037545 | 0.291683 | Bo9g109790.1 |
| 2765 | . | gene | MSTRG.5254 | 0.281226 | 0.037701 | 0.292336 | Bo2g095130.1 |
| 2768 | . | gene | MSTRG.9332 | 0.26788 | 0.037743 | 0.292336 | Bo3g060750.1 |
| 2764 | . | gene | MSTRG.20097 | 0.805733 | 0.037694 | 0.292336 | Bo6g024750.1 |
| 2767 | . | gene | MSTRG.23192 | 0.961737 | 0.037722 | 0.292336 | Bo7g024240.1 |
| 2762 | . | gene | MSTRG.23952 | 1.534738 | 0.037663 | 0.292336 | Bo7g061880.1 |
| 2766 | . | gene | MSTRG.32712 | 1.252021 | 0.03772 | 0.292336 | Bo9g151940.1 |
| 2763 | . | gene | MSTRG.25110 | 1.167108 | 0.037683 | 0.292336 | |
| 2769 | . | gene | MSTRG.32920 | 6.774622 | 0.037747 | 0.292336 | |
| 2770 | . | gene | MSTRG.34487 | 2.212154 | 0.037757 | 0.292336 | |
| 2771 | . | gene | MSTRG.1392 | 1.36598 | 0.037796 | 0.292533 | Bo1g043070.1 |
| 2772 | . | gene | MSTRG.34886 | 1.469571 | 0.037834 | 0.292669 | |
| 2773 | . | gene | MSTRG.15946 | 0.770657 | 0.037841 | 0.292669 | |
| 2774 | . | gene | MSTRG.20717 | 2.303877 | 0.03793 | 0.293256 | Bo6g064440.1 |
| 2775 | . | gene | MSTRG.9284 | 1.794879 | 0.03795 | 0.293303 | Bo3g059040.1 |
| 2776 | . | gene | MSTRG.33518 | 0.81181 | 0.037983 | 0.293448 | Bo9g175480.1 |
| 2777 | . | gene | MSTRG.2115 | 0.673129 | 0.038037 | 0.293766 | Bo1g091590.1 |
| 2778 | . | gene | MSTRG.24064 | 1.440003 | 0.038075 | 0.29395 | Bo7g065000.1 |
| 2779 | . | gene | MSTRG.22590 | 1.740933 | 0.038097 | 0.294014 | |
| 2781 | . | gene | MSTRG.21984 | 0.753495 | 0.038137 | 0.294029 | Bo6g106510.1 |

| | | | | | | | |
|---|---|---|---|---|---|---|---|
| 2782 | . | gene | MSTRG.27327 | 2.05342 | 0.03814 | 0.294029 | Bo8g059080.1 |
| 2780 | . | gene | MSTRG.11157 | 0.73493 | 0.038115 | 0.294029 | |
| 2783 | . | gene | MSTRG.35570 | 0.411795 | 0.038154 | 0.294034 | |
| 2784 | . | gene | MSTRG.32246 | 1.239871 | 0.038186 | 0.294172 | Bo9g123420.1 |
| 2785 | . | gene | MSTRG.17135 | 0.088772 | 0.038203 | 0.294197 | Bo5g027550.1 |
| 2787 | . | gene | MSTRG.14829 | 2.156392 | 0.038257 | 0.294338 | Bo4g149430.1 |
| 2786 | . | gene | MSTRG.29895 | 0.258752 | 0.038255 | 0.294338 | Bo9g006040.1 |
| 2788 | . | gene | MSTRG.30220 | 1.822559 | 0.038262 | 0.294338 | |
| 2790 | . | gene | MSTRG.2956 | 1.187982 | 0.038338 | 0.294707 | Bo1g141370.1 |
| 2789 | . | gene | MSTRG.23579 | 1.855306 | 0.038334 | 0.294707 | Bo7g045520.1 |
| 2791 | . | gene | MSTRG.32354 | 0.750415 | 0.038383 | 0.294947 | Bo9g132980.1 |
| 2794 | . | gene | MSTRG.3680 | 0.485257 | 0.038478 | 0.295095 | Bo2g011520.1 |
| 2792 | . | gene | MSTRG.9019 | 0.604755 | 0.038437 | 0.295095 | Bo3g052120.1 |
| 2796 | ATFBP7 | gene | MSTRG.17306 | 1.263004 | 0.038491 | 0.295095 | Bo5g034790.1 |
| 2795 | . | gene | MSTRG.21364 | 0.694445 | 0.038488 | 0.295095 | Bo6g081480.1 |
| 2793 | AT3BETAHSD/D2 | gene | MSTRG.28768 | 1.344097 | 0.038477 | 0.295095 | Bo8g098820.1 |
| 2798 | . | gene | MSTRG.29047 | 0.061321 | 0.038498 | 0.295095 | Bo8g104090.1 |
| 2797 | . | gene | MSTRG.31545 | 1.369651 | 0.038493 | 0.295095 | Bo9g078240.1 |
| 2799 | . | gene | MSTRG.22559 | 2.020309 | 0.03854 | 0.295307 | Bo6g121000.1 |
| 2802 | FZR3 | gene | MSTRG.7183 | 0.587858 | 0.038626 | 0.295649 | Bo3g009380.1 |
| 2800 | . | gene | MSTRG.20524 | 0.63498 | 0.038612 | 0.295649 | Bo6g049540.1 |
| 2801 | . | gene | MSTRG.26512 | 1.113389 | 0.038616 | 0.295649 | Bo8g011590.1 |
| 2803 | . | gene | MSTRG.34877 | 0.635923 | 0.03866 | 0.295808 | |
| 2804 | . | gene | MSTRG.19789 | 0.59724 | 0.038716 | 0.296125 | Bo5g155090.1 |
| 2806 | . | gene | MSTRG.788 | 0.331016 | 0.03878 | 0.296206 | Bo1g019090.1 |
| 2805 | . | gene | MSTRG.6864 | 1.293604 | 0.038749 | 0.296206 | Bo3g002720.1 |
| 2809 | . | gene | MSTRG.17656 | 2.865724 | 0.038795 | 0.296206 | Bo5g049310.1 |
| 2808 | . | gene | MSTRG.29304 | 0.299528 | 0.038787 | 0.296206 | Bo8g108140.1 |
| 2807 | . | gene | MSTRG.32925 | 4.006 | 0.038786 | 0.296206 | Bo9g160200.1 |

| | | | | | | | |
|---|---|---|---|---|---|---|---|
| 28 10 | . | gene | MSTRG.3182 9 | 1.0103 77 | 0.0388 27 | 0.2963 44 | |
| 28 11 | . | gene | MSTRG.7313 | 1.5688 75 | 0.0388 61 | 0.2964 98 | Bo3g0107 80.1 |
| 28 12 | . | gene | MSTRG.9512 | 1.5734 03 | 0.0388 78 | 0.2965 18 | Bo3g0646 10.1 |
| 28 14 | . | gene | MSTRG.1987 0 | 1.5079 5 | 0.0389 38 | 0.2966 87 | Bo6g0058 10.1 |
| 28 13 | . | gene | gene:Bo2g07 5370 | 0.9145 64 | 0.0389 26 | 0.2966 87 | |
| 28 15 | . | gene | MSTRG.3489 8 | 0.1212 66 | 0.0389 41 | 0.2966 87 | |
| 28 16 | . | gene | MSTRG.2528 3 | 1.2637 56 | 0.0389 81 | 0.2968 28 | Bo7g1007 60.1 |
| 28 17 | . | gene | MSTRG.2393 6 | 0.7783 07 | 0.0389 91 | 0.2968 28 | |
| 28 18 | . | gene | MSTRG.3186 8 | 0.7449 16 | 0.0390 01 | 0.2968 28 | |
| 28 19 | . | gene | MSTRG.1701 | 0.6179 42 | 0.0390 2 | 0.2968 65 | Bo1g0571 90.1 |
| 28 21 | . | gene | MSTRG.1779 | 1.4745 34 | 0.0390 56 | 0.2968 88 | Bo1g0594 20.1 |
| 28 22 | . | gene | MSTRG.3620 | 1.3392 24 | 0.0390 65 | 0.2968 88 | Bo2g0106 00.1 |
| 28 20 | . | gene | MSTRG.7444 | 0.5243 72 | 0.0390 51 | 0.2968 88 | Bo3g0136 20.1 |
| 28 23 | . | gene | MSTRG.3093 7 | 0.2163 6 | 0.0391 19 | 0.2971 94 | |
| 28 24 | . | gene | MSTRG.3477 | 1.7908 62 | 0.0391 35 | 0.2972 16 | Bo2g0075 20.1 |
| 28 27 | . | gene | MSTRG.1430 | 0.6953 84 | 0.0391 82 | 0.2972 39 | Bo1g0465 30.1 |
| 28 28 | . | gene | MSTRG.1158 4 | 0.7167 88 | 0.0392 08 | 0.2972 39 | Bo3g1622 00.1 |
| 28 29 | . | gene | MSTRG.1229 9 | 1.6341 63 | 0.0392 11 | 0.2972 39 | Bo4g0127 20.1 |
| 28 30 | . | gene | MSTRG.2148 7 | 0.6000 11 | 0.0392 22 | 0.2972 39 | Bo6g0865 00.1 |
| 28 25 | . | gene | MSTRG.3502 1 | 8.5972 77 | 0.0391 53 | 0.2972 39 | |
| 28 26 | . | gene | MSTRG.3446 2 | 0.1351 1 | 0.0391 73 | 0.2972 39 | |
| 28 31 | . | gene | MSTRG.2694 0 | 0.1012 02 | 0.0392 84 | 0.2976 06 | |
| 28 32 | . | gene | MSTRG.6591 | 0.5394 31 | 0.0393 14 | 0.2976 41 | Bo2g1618 10.1 |
| 28 33 | . | gene | MSTRG.2108 8 | 1.2561 76 | 0.0393 16 | 0.2976 41 | Bo6g0761 40.1 |
| 28 34 | . | gene | MSTRG.2821 7 | 1.2151 8 | 0.0393 33 | 0.2976 6 | Bo8g0834 70.1 |
| 28 35 | . | gene | MSTRG.1543 3 | 0.2989 4 | 0.0393 59 | 0.2977 55 | Bo4g1825 70.1 |
| 28 36 | . | gene | MSTRG.9474 | 0.7053 89 | 0.0394 14 | 0.2980 62 | |
| 28 37 | . | gene | MSTRG.2862 4 | 1.5603 15 | 0.0394 75 | 0.2983 14 | Bo8g0956 90.1 |
| 28 38 | . | gene | MSTRG.7404 | 0.0800 46 | 0.0394 75 | 0.2983 14 | |

| | | | | | | | |
|---|---|---|---|---|---|---|---|
| 2839 | . | gene | MSTRG.27228 | 0.508602 | 0.039522 | 0.298566 | Bo8g054550.1 |
| 2840 | . | gene | MSTRG.30382 | 1.28724 | 0.039553 | 0.298693 | Bo9g018730.1 |
| 2841 | . | gene | MSTRG.9142 | 1.363215 | 0.039639 | 0.299241 | Bo3g054980.1 |
| 2843 | . | gene | MSTRG.11563 | 1.336096 | 0.03967 | 0.299265 | Bo3g158690.1 |
| 2842 | . | gene | MSTRG.30639 | 2.079626 | 0.039664 | 0.299265 | Bo9g027730.1 |
| 2844 | . | gene | MSTRG.9483 | 0.103006 | 0.039688 | 0.299292 | |
| 2845 | . | gene | MSTRG.11181 | 2.197918 | 0.03974 | 0.299579 | Bo3g143260.1 |
| 2846 | . | gene | MSTRG.9594 | 0.439831 | 0.039772 | 0.299615 | Bo3g065730.1 |
| 2847 | . | gene | MSTRG.23321 | 1.877557 | 0.039773 | 0.299615 | Bo7g031650.1 |
| 2848 | . | gene | MSTRG.830 | 1.312472 | 0.039801 | 0.299725 | Bo1g019700.1 |
| 2849 | . | gene | MSTRG.5075 | 1.274069 | 0.039829 | 0.299797 | Bo2g085220.1 |
| 2850 | . | gene | MSTRG.23914 | 1.107115 | 0.039839 | 0.299797 | Bo7g061310.1 |
| 2851 | . | gene | MSTRG.29986 | 1.504157 | 0.039892 | 0.300091 | Bo9g008490.1 |
| 2852 | . | gene | MSTRG.22927 | 1.876461 | 0.039959 | 0.300453 | |
| 2853 | . | gene | MSTRG.7626 | 6.839804 | 0.039968 | 0.300453 | |
| 2854 | . | gene | MSTRG.30879 | 0.926017 | 0.039984 | 0.300467 | |
| 2855 | . | gene | MSTRG.7475 | 0.312776 | 0.040009 | 0.300474 | Bo3g014020.1 |
| 2856 | . | gene | MSTRG.17674 | 1.863074 | 0.040013 | 0.300474 | Bo5g050640.1 |
| 2857 | . | gene | MSTRG.24951 | 0.572023 | 0.040032 | 0.300514 | |
| 2858 | . | gene | MSTRG.35885 | 3.869914 | 0.040068 | 0.300644 | |
| 2859 | . | gene | MSTRG.34827 | 0.410562 | 0.040077 | 0.300644 | |
| 2860 | . | gene | MSTRG.11361 | 1.765656 | 0.040108 | 0.300696 | Bo3g149950.1 |
| 2861 | . | gene | MSTRG.18000 | 1.669584 | 0.040112 | 0.300696 | Bo5g072540.1 |
| 2868 | HSP60-3A | gene | MSTRG.9597 | 1.590829 | 0.040218 | 0.300705 | Bo3g065760.1 |
| 2862 | . | gene | MSTRG.16630 | 1.488654 | 0.040162 | 0.300705 | Bo5g013660.1 |
| 2865 | . | gene | MSTRG.22689 | 1.597941 | 0.040205 | 0.300705 | Bo6g123950.1 |
| 2866 | . | gene | MSTRG.27699 | 0.36932 | 0.040212 | 0.300705 | Bo8g069990.1 |
| 2864 | . | gene | MSTRG.28579 | 0.279869 | 0.040204 | 0.300705 | Bo8g094720.1 |
| 2869 | . | gene | MSTRG.31033 | 0.530976 | 0.040226 | 0.300705 | Bo9g048870.1 |

| | | | | | | | |
|---|---|---|---|---|---|---|---|
| 28<br>63 | . | gene | MSTRG.3224<br>0 | 1.5475<br>26 | 0.0401<br>99 | 0.3007<br>05 | Bo9g1222<br>20.1 |
| 28<br>67 | . | gene | MSTRG.1442<br>0 | 3.0437<br>96 | 0.0402<br>14 | 0.3007<br>05 | |
| 28<br>73 | . | gene | MSTRG.1396<br>2 | 3.2077<br>24 | 0.0402<br>9 | 0.3007<br>26 | Bo4g1082<br>40.1 |
| 28<br>70 | . | gene | MSTRG.1852<br>0 | 1.5214<br>56 | 0.0402<br>61 | 0.3007<br>26 | Bo5g1099<br>20.1 |
| 28<br>71 | . | gene | MSTRG.2990<br>1 | 1.6411<br>23 | 0.0402<br>68 | 0.3007<br>26 | Bo9g0061<br>40.1 |
| 28<br>72 | . | gene | MSTRG.2677<br>4 | 1.6162<br>44 | 0.0402<br>83 | 0.3007<br>26 | |
| 28<br>74 | . | gene | MSTRG.1485<br>8 | 0.1472<br>13 | 0.0402<br>99 | 0.3007<br>26 | |
| 28<br>75 | . | gene | MSTRG.2899<br>9 | 2.0412<br>07 | 0.0403<br>33 | 0.3008<br>79 | Bo8g1022<br>70.1 |
| 28<br>77 | . | gene | MSTRG.1336 | 1.5353<br>74 | 0.0403<br>95 | 0.3011<br>13 | Bo1g0403<br>30.1 |
| 28<br>81 | . | gene | MSTRG.3204 | 1.6151<br>19 | 0.0404<br>72 | 0.3011<br>13 | Bo1g1521<br>80.1 |
| 28<br>83 | . | gene | MSTRG.9759 | 0.6378<br>02 | 0.0404<br>77 | 0.3011<br>13 | Bo3g0713<br>40.1 |
| 28<br>80 | . | gene | MSTRG.2516<br>1 | 1.2924<br>38 | 0.0404<br>64 | 0.3011<br>13 | Bo7g0978<br>20.1 |
| 28<br>76 | . | gene | MSTRG.1589<br>0 | 1.7167<br>81 | 0.0403<br>92 | 0.3011<br>13 | |
| 28<br>78 | . | gene | MSTRG.1029<br>1 | 0.9516<br>49 | 0.0404<br>26 | 0.3011<br>13 | |
| 28<br>79 | . | gene | MSTRG.3458<br>7 | 5.7977<br>11 | 0.0404<br>63 | 0.3011<br>13 | |
| 28<br>82 | . | gene | MSTRG.3190<br>3 | 0.1195<br>59 | 0.0404<br>75 | 0.3011<br>13 | |
| 28<br>84 | . | gene | MSTRG.381 | 0.5754<br>31 | 0.0405<br>5 | 0.3015<br>53 | Bo1g0080<br>80.1 |
| 28<br>85 | . | gene | MSTRG.2917<br>2 | 1.7477<br>91 | 0.0405<br>9 | 0.3017<br>43 | Bo8g1059<br>10.1 |
| 28<br>88 | . | gene | MSTRG.1192 | 0.2604<br>4 | 0.0406<br>56 | 0.3018<br>56 | Bo1g0372<br>00.1 |
| 28<br>86 | . | gene | MSTRG.1927<br>3 | 1.2067<br>06 | 0.0406<br>21 | 0.3018<br>56 | Bo5g1370<br>30.1 |
| 28<br>89 | . | gene | MSTRG.2415<br>9 | 0.3467<br>42 | 0.0406<br>61 | 0.3018<br>56 | Bo7g0675<br>30.1 |
| 28<br>87 | . | gene | MSTRG.2458<br>6 | 1.5692<br>64 | 0.0406<br>38 | 0.3018<br>56 | Bo7g0825<br>40.1 |
| 28<br>90 | . | gene | MSTRG.1827<br>7 | 0.5840<br>91 | 0.0406<br>81 | 0.3018<br>96 | Bo5g0930<br>80.1 |
| 28<br>93 | . | gene | MSTRG.7069 | 0.2659<br>37 | 0.0407<br>34 | 0.3018<br>99 | Bo3g0067<br>70.1 |
| 28<br>92 | . | gene | MSTRG.1283<br>2 | 1.2819<br>43 | 0.0407<br>32 | 0.3018<br>99 | Bo4g0312<br>10.1 |
| 28<br>91 | . | gene | MSTRG.1940<br>5 | 1.2977<br>69 | 0.0407 | 0.3018<br>99 | Bo5g1398<br>70.1 |
| 28<br>94 | . | gene | MSTRG.2060<br>6 | 0.5931<br>39 | 0.0407<br>37 | 0.3018<br>99 | Bo6g0548<br>30.1 |
| 28<br>95 | . | gene | MSTRG.3464<br>7 | 0.6014<br>1 | 0.0407<br>54 | 0.3019<br>2 | |
| 28<br>96 | . | gene | MSTRG.3477<br>0 | 1.4548<br>38 | 0.0407<br>73 | 0.3019<br>56 | |

| | | | | | | | |
|---|---|---|---|---|---|---|---|
| 2897 | . | gene | MSTRG.6538 | 1.340768 | 0.040794 | 0.302002 | Bo2g160900.1 |
| 2898 | . | gene | MSTRG.22214 | 0.422308 | 0.040868 | 0.302447 | Bo6g113140.1 |
| 2900 | . | gene | MSTRG.16913 | 4.88386 | 0.040916 | 0.302597 | Bo5g021160.1 |
| 2899 | . | gene | MSTRG.5089 | 0.774215 | 0.040907 | 0.302597 | |
| 2901 | . | gene | MSTRG.11867 | 2.03683 | 0.04094 | 0.302671 | Bo3g175190.1 |
| 2902 | . | gene | MSTRG.4059 | 0.866722 | 0.040982 | 0.302874 | |
| 2906 | . | gene | MSTRG.22663 | 0.708597 | 0.041104 | 0.303182 | Bo6g122510.1 |
| 2905 | . | gene | MSTRG.29143 | 0.808757 | 0.041097 | 0.303182 | Bo8g105570.1 |
| 2908 | . | gene | MSTRG.29677 | 0.777039 | 0.041108 | 0.303182 | Bo8g117650.1 |
| 2903 | . | gene | MSTRG.31134 | 1.329135 | 0.041042 | 0.303182 | Bo9g058310.1 |
| 2907 | . | gene | MSTRG.32966 | 0.888025 | 0.041105 | 0.303182 | Bo9g161790.1 |
| 2904 | . | gene | MSTRG.35386 | 0.226943 | 0.04106 | 0.303182 | |
| 2909 | . | gene | MSTRG.2172 | 0.968229 | 0.041134 | 0.303264 | Bo1g095760.1 |
| 2910 | . | gene | MSTRG.19703 | 0.287011 | 0.041241 | 0.30395 | Bo5g149880.1 |
| 2911 | . | gene | MSTRG.2765 | 0.594527 | 0.041263 | 0.30401 | Bo1g133320.1 |
| 2912 | . | gene | MSTRG.22264 | 0.443589 | 0.041279 | 0.30402 | Bo6g115880.1 |
| 2913 | . | gene | MSTRG.19366 | 1.253914 | 0.041366 | 0.304561 | Bo5g139390.1 |
| 2917 | . | gene | MSTRG.2021 | 0.082954 | 0.041438 | 0.304639 | Bo1g084840.1 |
| 2916 | . | gene | MSTRG.2929 | 1.182309 | 0.041431 | 0.304639 | Bo1g141020.1 |
| 2918 | . | gene | MSTRG.18802 | 1.388731 | 0.041448 | 0.304639 | Bo5g123050.1 |
| 2915 | . | gene | MSTRG.27658 | 0.28977 | 0.041419 | 0.304639 | Bo8g068270.1 |
| 2914 | . | gene | MSTRG.28728 | 1.380236 | 0.041393 | 0.304639 | Bo8g098220.1 |
| 2919 | . | gene | MSTRG.4510 | 1.297624 | 0.041509 | 0.304846 | Bo2g050670.1 |
| 2921 | . | gene | MSTRG.6313 | 1.442871 | 0.041527 | 0.304846 | Bo2g150420.1 |
| 2923 | . | gene | MSTRG.19952 | 0.818384 | 0.041547 | 0.304846 | Bo6g013250.1 |
| 2920 | . | gene | MSTRG.22183 | 2.820143 | 0.041509 | 0.304846 | Bo6g112710.1 |
| 2922 | . | gene | MSTRG.34764 | 0.711397 | 0.041536 | 0.304846 | |
| 2925 | . | gene | MSTRG.1945 | 0.896579 | 0.041583 | 0.304855 | Bo1g078490.1 |
| 2924 | . | gene | MSTRG.32125 | 1.175971 | 0.041574 | 0.304855 | Bo9g117490.1 |

| | | | | | | | |
|---|---|---|---|---|---|---|---|
| 29<br>26 | . | gene | MSTRG.3396<br>4 | 0.1220<br>56 | 0.0415<br>91 | 0.3048<br>55 | |
| 29<br>34 | . | gene | MSTRG.6638 | 0.015 | 0.0417<br>38 | 0.3048<br>99 | Bo2g1655<br>90.1 |
| 29<br>35 | . | gene | MSTRG.8878 | 0.7250<br>58 | 0.0417<br>43 | 0.3048<br>99 | Bo3g0445<br>80.1 |
| 29<br>37 | . | gene | MSTRG.9532 | 0.0745<br>23 | 0.0417<br>66 | 0.3048<br>99 | Bo3g0648<br>70.1 |
| 29<br>27 | GPT | gene | MSTRG.1232<br>9 | 2.0400<br>04 | 0.0416<br>17 | 0.3048<br>99 | Bo4g0133<br>00.1 |
| 29<br>29 | . | gene | MSTRG.1516<br>7 | 0.4220<br>4 | 0.0416<br>85 | 0.3048<br>99 | Bo4g1692<br>00.1 |
| 29<br>30 | CYP88A3 | gene | MSTRG.1624<br>0 | 0.6078<br>54 | 0.0416<br>92 | 0.3048<br>99 | Bo5g0052<br>80.1 |
| 29<br>33 | . | gene | MSTRG.1815<br>7 | 0.8626<br>23 | 0.0417<br>17 | 0.3048<br>99 | Bo5g0840<br>00.1 |
| 29<br>32 | . | gene | MSTRG.1851<br>9 | 0.7942<br>22 | 0.0417<br>08 | 0.3048<br>99 | Bo5g1098<br>60.1 |
| 29<br>38 | . | gene | MSTRG.3011<br>3 | 2.6119<br>54 | 0.0417<br>72 | 0.3048<br>99 | Bo9g0106<br>10.1 |
| 29<br>28 | . | gene | MSTRG.3338<br>8 | 1.2455<br>8 | 0.0416<br>78 | 0.3048<br>99 | Bo9g1736<br>90.1 |
| 29<br>31 | . | gene | MSTRG.3187<br>7 | 0.5980<br>6 | 0.0416<br>99 | 0.3048<br>99 | |
| 29<br>36 | . | gene | MSTRG.1692<br>6 | 0.5642<br>92 | 0.0417<br>59 | 0.3048<br>99 | |
| 29<br>39 | . | gene | MSTRG.3530<br>0 | 0.2660<br>03 | 0.0417<br>82 | 0.3048<br>99 | |
| 29<br>43 | . | gene | MSTRG.5032 | 0.6461<br>83 | 0.0419<br>01 | 0.3052<br>54 | Bo2g0823<br>00.1 |
| 29<br>42 | . | gene | MSTRG.1134<br>3 | 0.6768 | 0.0418<br>88 | 0.3052<br>54 | Bo3g1496<br>70.1 |
| 29<br>45 | . | gene | MSTRG.1139<br>6 | 1.6966<br>46 | 0.0419<br>41 | 0.3052<br>54 | Bo3g1526<br>80.1 |
| 29<br>41 | . | gene | MSTRG.1253<br>5 | 1.669 | 0.0418<br>76 | 0.3052<br>54 | Bo4g0242<br>80.1 |
| 29<br>46 | . | gene | MSTRG.1316<br>9 | 1.3524<br>07 | 0.0419<br>45 | 0.3052<br>54 | Bo4g0461<br>90.1 |
| 29<br>44 | . | gene | MSTRG.2332<br>9 | 0.6400<br>8 | 0.0419<br>32 | 0.3052<br>54 | Bo7g0318<br>10.1 |
| 29<br>40 | . | gene | MSTRG.2630<br>9 | 1.7156<br>28 | 0.0418<br>56 | 0.3052<br>54 | Bo8g0012<br>80.1 |
| 29<br>48 | . | gene | MSTRG.2756<br>9 | 0.7196<br>46 | 0.0419<br>59 | 0.3052<br>54 | Bo8g0667<br>90.1 |
| 29<br>47 | . | gene | MSTRG.3349<br>6 | 0.7005<br>68 | 0.0419<br>58 | 0.3052<br>54 | Bo9g1750<br>80.1 |
| 29<br>55 | . | gene | MSTRG.2664 | 1.0662<br>39 | 0.0420<br>63 | 0.3052<br>86 | Bo1g1234<br>70.1 |
| 29<br>54 | . | gene | MSTRG.5313 | 1.4160<br>16 | 0.0420<br>52 | 0.3052<br>86 | Bo2g0961<br>40.1 |
| 29<br>50 | . | gene | MSTRG.1185<br>5 | 1.1604<br>47 | 0.0420<br>18 | 0.3052<br>86 | Bo3g1739<br>50.1 |
| 29<br>49 | . | gene | MSTRG.1566<br>6 | 0.7107<br>06 | 0.0419<br>9 | 0.3052<br>86 | Bo4g1884<br>60.1 |
| 29<br>52 | . | gene | MSTRG.1945<br>8 | 1.2302<br>7 | 0.0420<br>25 | 0.3052<br>86 | Bo5g1414<br>40.1 |
| 29<br>51 | . | gene | MSTRG.2670<br>2 | 1.4393<br>6 | 0.0420<br>19 | 0.3052<br>86 | Bo8g0268<br>40.1 |

| | | | | | | | |
|---|---|---|---|---|---|---|---|
| 2953 | . | gene | MSTRG.25632 | 0.453776 | 0.042047 | 0.305286 | |
| 2956 | . | gene | MSTRG.33437 | 1.617352 | 0.042078 | 0.305294 | |
| 2958 | . | gene | MSTRG.10340 | 0.563298 | 0.042119 | 0.305382 | Bo3g097580.1 |
| 2957 | . | gene | MSTRG.36160 | 0.64029 | 0.042112 | 0.305382 | |
| 2959 | . | gene | MSTRG.7195 | 0.179205 | 0.042151 | 0.305512 | Bo3g009510.1 |
| 2960 | . | gene | MSTRG.29535 | 1.529226 | 0.042217 | 0.30589 | Bo8g114260.1 |
| 2961 | . | gene | MSTRG.10505 | 2.152753 | 0.042232 | 0.305896 | Bo3g102820.1 |
| 2963 | . | gene | MSTRG.995 | 0.506628 | 0.042299 | 0.305917 | Bo1g024180.1 |
| 2968 | . | gene | MSTRG.2246 | 0.474416 | 0.042344 | 0.305917 | Bo1g101110.1 |
| 2967 | . | gene | MSTRG.5437 | 0.986237 | 0.042335 | 0.305917 | Bo2g102160.1 |
| 2969 | . | gene | MSTRG.15189 | 1.465384 | 0.042349 | 0.305917 | Bo4g170660.1 |
| 2962 | . | gene | MSTRG.18303 | 1.393932 | 0.042295 | 0.305917 | Bo5g094500.1 |
| 2964 | . | gene | MSTRG.20814 | 0.648526 | 0.0423 | 0.305917 | Bo6g067010.1 |
| 2965 | . | gene | MSTRG.35365 | 1.608645 | 0.042305 | 0.305917 | |
| 2966 | . | gene | MSTRG.6647 | 1.649888 | 0.042316 | 0.305917 | |
| 2971 | . | gene | MSTRG.29470 | 0.404605 | 0.042425 | 0.306237 | Bo8g112380.1 |
| 2970 | . | gene | MSTRG.33629 | 1.445155 | 0.042413 | 0.306237 | |
| 2972 | . | gene | MSTRG.9983 | 0.585081 | 0.042437 | 0.306237 | |
| 2978 | . | gene | MSTRG.6001 | 0.214315 | 0.042528 | 0.306279 | Bo2g136400.1 |
| 2975 | . | gene | MSTRG.6737 | 0.650369 | 0.042498 | 0.306279 | Bo2g167990.1 |
| 2974 | . | gene | MSTRG.12731 | 1.804377 | 0.042482 | 0.306279 | Bo4g027340.1 |
| 2973 | . | gene | MSTRG.21807 | 0.709822 | 0.042471 | 0.306279 | Bo6g099190.1 |
| 2977 | . | gene | MSTRG.25421 | 2.424828 | 0.042521 | 0.306279 | Bo7g104970.1 |
| 2976 | . | gene | MSTRG.33293 | 0.301063 | 0.042513 | 0.306279 | Bo9g171490.1 |
| 2979 | . | gene | MSTRG.6922 | 2.858065 | 0.042562 | 0.306362 | Bo3g003690.1 |
| 2981 | . | gene | MSTRG.9476 | 1.334664 | 0.042592 | 0.306362 | Bo3g064150.1 |
| 2982 | . | gene | MSTRG.19374 | 1.346095 | 0.042597 | 0.306362 | Bo5g139530.1 |
| 2980 | . | gene | MSTRG.20831 | 0.030635 | 0.042589 | 0.306362 | |
| 2983 | . | gene | MSTRG.8561 | 1.166785 | 0.042613 | 0.30638 | Bo3g036550.1 |

| | | | | | | | |
|---|---|---|---|---|---|---|---|
| 29 85 | . | gene | MSTRG.1756 2 | 0.1075 32 | 0.0426 72 | 0.3065 8 | Bo5g0449 40.1 |
| 29 84 | . | gene | MSTRG.2322 0 | 1.3914 9 | 0.0426 57 | 0.3065 8 | Bo7g0267 70.1 |
| 29 86 | . | gene | MSTRG.2883 9 | 0.6323 44 | 0.0426 96 | 0.3065 8 | Bo8g0999 00.1 |
| 29 87 | . | gene | MSTRG.3171 6 | 0.8931 17 | 0.0426 99 | 0.3065 8 | Bo9g0914 10.1 |
| 29 88 | . | gene | MSTRG.3118 7 | 0.4609 84 | 0.0427 24 | 0.3066 61 | |
| 29 90 | . | gene | MSTRG.1436 9 | 1.5938 53 | 0.0427 99 | 0.3069 95 | Bo4g1307 90.1 |
| 29 89 | . | gene | MSTRG.2529 0 | 0.7238 97 | 0.0427 92 | 0.3069 95 | Bo7g1008 60.1 |
| 29 91 | . | gene | MSTRG.2962 3 | 0.8503 09 | 0.0428 51 | 0.3072 62 | Bo8g1156 90.1 |
| 29 92 | BHLH101 | gene | MSTRG.6858 | 0.0698 65 | 0.0428 74 | 0.3073 26 | Bo3g0026 30.1 |
| 29 95 | . | gene | MSTRG.1017 6 | 0.0155 79 | 0.0429 57 | 0.3076 03 | Bo3g0884 80.1 |
| 29 93 | . | gene | MSTRG.1938 3 | 1.5144 95 | 0.0429 28 | 0.3076 03 | Bo5g1396 30.1 |
| 29 97 | . | gene | MSTRG.2428 4 | 1.5870 5 | 0.0429 87 | 0.3076 03 | Bo7g0756 90.1 |
| 29 94 | . | gene | MSTRG.2557 1 | 0.4901 42 | 0.0429 47 | 0.3076 03 | Bo7g1075 80.1 |
| 29 98 | . | gene | MSTRG.3101 3 | 0.1438 71 | 0.0429 99 | 0.3076 03 | Bo9g0466 10.1 |
| 29 96 | . | gene | MSTRG.3212 1 | 0.5108 44 | 0.0429 76 | 0.3076 03 | Bo9g1174 30.1 |
| 29 99 | BGAL17 | gene | MSTRG.2157 8 | 0.6614 63 | 0.0430 19 | 0.3076 14 | Bo6g0881 30.1 |
| 30 00 | . | gene | MSTRG.3159 5 | 2.0743 49 | 0.0430 39 | 0.3076 14 | Bo9g0831 20.1 |
| 30 01 | . | gene | MSTRG.3378 5 | 0.7572 91 | 0.0430 43 | 0.3076 14 | Bo9g1833 30.1 |
| 30 02 | . | gene | MSTRG.2545 0 | 1.5426 7 | 0.0430 69 | 0.3076 93 | Bo7g1053 60.1 |
| 30 03 | . | gene | MSTRG.3143 2 | 1.0904 23 | 0.0431 27 | 0.3080 07 | Bo9g0732 40.1 |
| 30 04 | . | gene | MSTRG.3062 7 | 1.3872 82 | 0.0431 74 | 0.3082 41 | Bo9g0275 90.1 |
| 30 05 | . | gene | MSTRG.1245 4 | 1.3948 38 | 0.0431 91 | 0.3082 57 | Bo4g0210 40.1 |
| 30 06 | . | gene | MSTRG.7738 | 0.7320 41 | 0.0432 16 | 0.3083 32 | Bo3g0195 50.1 |
| 30 07 | . | gene | MSTRG.2506 4 | 1.4828 89 | 0.0432 41 | 0.3084 07 | Bo7g0964 30.1 |
| 30 08 | . | gene | MSTRG.4861 | 2.2572 78 | 0.0432 72 | 0.3085 27 | Bo2g0684 30.1 |
| 30 09 | . | gene | MSTRG.2391 0 | 1.5385 91 | 0.0432 92 | 0.3085 68 | Bo7g0612 40.1 |
| 30 11 | . | gene | MSTRG.1142 3 | 1.4187 43 | 0.0433 79 | 0.3088 92 | Bo3g1533 00.1 |
| 30 12 | NRPA2 | gene | MSTRG.2353 2 | 2.1936 52 | 0.0434 01 | 0.3088 92 | Bo7g0448 90.1 |
| 30 13 | . | gene | MSTRG.2938 1 | 1.8120 46 | 0.0434 09 | 0.3088 92 | Bo8g1092 30.1 |

| | | | | | | | |
|---|---|---|---|---|---|---|---|
| 3014 | UGT71C4 | gene | MSTRG.29531 | 0.072396 | 0.043409 | 0.308892 | Bo8g114230.1 |
| 3010 | . | gene | MSTRG.34493 | 1.159619 | 0.043352 | 0.308892 | |
| 3015 | . | gene | MSTRG.29448 | 2.01987 | 0.04344 | 0.309005 | Bo8g111030.1 |
| 3016 | . | gene | MSTRG.1594 | 0.42466 | 0.043466 | 0.309089 | Bo1g053340.1 |
| 3017 | . | gene | MSTRG.19358 | 1.240027 | 0.043545 | 0.309552 | Bo5g139280.1 |
| 3020 | . | gene | MSTRG.10312 | 0.275023 | 0.043602 | 0.309646 | Bo3g094280.1 |
| 3019 | . | gene | MSTRG.14463 | 1.8779 | 0.0436 | 0.309646 | Bo4g136190.1 |
| 3018 | . | gene | MSTRG.30533 | 1.16727 | 0.043599 | 0.309646 | Bo9g023760.1 |
| 3022 | . | gene | MSTRG.3706 | 1.612095 | 0.043637 | 0.30969 | Bo2g012050.1 |
| 3021 | . | gene | MSTRG.30617 | 1.097175 | 0.043635 | 0.30969 | |
| 3023 | . | gene | MSTRG.31702 | 0.426279 | 0.043674 | 0.309847 | Bo9g091160.1 |
| 3024 | . | gene | MSTRG.23553 | 0.732451 | 0.043721 | 0.310078 | Bo7g045250.1 |
| 3025 | . | gene | MSTRG.23316 | 0.64666 | 0.043736 | 0.310084 | Bo7g031530.1 |
| 3026 | . | gene | MSTRG.25548 | 1.68742 | 0.043779 | 0.310285 | Bo7g107230.1 |
| 3029 | . | gene | MSTRG.10238 | 1.292466 | 0.043833 | 0.31036 | Bo3g091790.1 |
| 3028 | . | gene | MSTRG.11178 | 1.333219 | 0.043831 | 0.31036 | Bo3g143230.1 |
| 3027 | . | gene | MSTRG.24479 | 1.443086 | 0.043822 | 0.31036 | Bo7g080590.1 |
| 3038 | AtAGAL2 | gene | MSTRG.3512 | 1.929547 | 0.044008 | 0.310462 | Bo2g009050.1 |
| 3037 | . | gene | MSTRG.9246 | 0.741476 | 0.044 | 0.310462 | Bo3g058530.1 |
| 3033 | HEN4 | gene | MSTRG.10469 | 1.307672 | 0.043923 | 0.310462 | Bo3g101340.1 |
| 3034 | . | gene | MSTRG.10788 | 1.735488 | 0.043926 | 0.310462 | Bo3g120920.1 |
| 3030 | . | gene | MSTRG.14791 | 0.679549 | 0.043863 | 0.310462 | Bo4g145870.1 |
| 3035 | . | gene | MSTRG.17720 | 0.783563 | 0.043941 | 0.310462 | Bo5g052340.1 |
| 3032 | ATGSTF11 | gene | MSTRG.19715 | 1.878999 | 0.043899 | 0.310462 | Bo5g150180.1 |
| 3036 | . | gene | MSTRG.24182 | 1.492436 | 0.043978 | 0.310462 | Bo7g069890.1 |
| 3040 | . | gene | MSTRG.28452 | 1.499477 | 0.044022 | 0.310462 | Bo8g091690.1 |
| 3031 | . | gene | MSTRG.5136 | 0.992607 | 0.043881 | 0.310462 | |
| 3039 | . | gene | MSTRG.34200 | 1.82413 | 0.044009 | 0.310462 | |
| 3041 | . | gene | MSTRG.7495 | 0.934786 | 0.04403 | 0.310462 | |

| | | | | | | | |
|---|---|---|---|---|---|---|---|
| 3042 | . | gene | MSTRG.29493 | 1.383098 | 0.044035 | 0.310462 | |
| 3050 | . | gene | MSTRG.1288 | 0.18297 | 0.044201 | 0.310555 | Bo1g039500.1 |
| 3056 | . | gene | MSTRG.1646 | 0.258842 | 0.04427 | 0.310555 | Bo1g055140.1 |
| 3054 | . | gene | MSTRG.7250 | 0.623564 | 0.044255 | 0.310555 | Bo3g010080.1 |
| 3045 | . | gene | MSTRG.9626 | 0.792214 | 0.044139 | 0.310555 | Bo3g066140.1 |
| 3052 | . | gene | MSTRG.11131 | 1.288833 | 0.044225 | 0.310555 | Bo3g141560.1 |
| 3049 | . | gene | MSTRG.13835 | 0.809246 | 0.044168 | 0.310555 | Bo4g101230.1 |
| 3046 | . | gene | MSTRG.19519 | 0.322805 | 0.044147 | 0.310555 | Bo5g143220.1 |
| 3057 | . | gene | MSTRG.21070 | 0.801935 | 0.044273 | 0.310555 | Bo6g074910.1 |
| 3053 | . | gene | MSTRG.26458 | 1.506651 | 0.044241 | 0.310555 | Bo8g009480.1 |
| 3047 | . | gene | MSTRG.27421 | 1.591101 | 0.044159 | 0.310555 | Bo8g061440.1 |
| 3055 | . | gene | MSTRG.29384 | 1.36595 | 0.044266 | 0.310555 | Bo8g109260.1 |
| 3043 | . | gene | MSTRG.14445 | 0.054831 | 0.044081 | 0.310555 | |
| 3044 | . | gene | MSTRG.35419 | 2.831346 | 0.044104 | 0.310555 | |
| 3048 | . | gene | MSTRG.35319 | 0.686692 | 0.044159 | 0.310555 | |
| 3051 | . | gene | MSTRG.35873 | 0.49063 | 0.044202 | 0.310555 | |
| 3058 | . | gene | MSTRG.34555 | 1.198095 | 0.04428 | 0.310555 | |
| 3060 | . | gene | MSTRG.19595 | 1.508627 | 0.044321 | 0.31061 | Bo5g145420.1 |
| 3059 | . | gene | MSTRG.21843 | 0.290051 | 0.044316 | 0.31061 | Bo6g099860.1 |
| 3061 | . | gene | MSTRG.26519 | 0.712211 | 0.044333 | 0.31061 | Bo8g011680.1 |
| 3062 | . | gene | MSTRG.35809 | 0.910008 | 0.044346 | 0.31061 | |
| 3063 | . | gene | MSTRG.18442 | 0.419699 | 0.044385 | 0.31078 | Bo5g103510.1 |
| 3064 | SNRK2-5 | gene | MSTRG.10535 | 0.494944 | 0.044475 | 0.311245 | Bo3g103200.1 |
| 3066 | . | gene | MSTRG.29257 | 1.346865 | 0.044495 | 0.311245 | Bo8g107190.1 |
| 3065 | . | gene | MSTRG.34860 | 1.784428 | 0.044493 | 0.311245 | |
| 3067 | . | gene | MSTRG.7531 | 2.028314 | 0.044522 | 0.311334 | Bo3g015760.1 |
| 3068 | . | gene | MSTRG.386 | 1.1925 | 0.044552 | 0.311442 | Bo1g008160.1 |
| 3069 | . | gene | MSTRG.3551 | 1.223675 | 0.044633 | 0.311905 | Bo2g009690.1 |
| 3071 | . | gene | MSTRG.10570 | 0.485203 | 0.044695 | 0.312064 | Bo3g104710.1 |

| | | | | | | | |
|---|---|---|---|---|---|---|---|
| 3070 | . | gene | MSTRG.10854 | 1.447012 | 0.044687 | 0.312064 | Bo3g127230.1 |
| 3072 | . | gene | MSTRG.30503 | 2.494794 | 0.044699 | 0.312064 | Bo9g022310.1 |
| 3073 | . | gene | MSTRG.23313 | 1.240179 | 0.044722 | 0.312066 | Bo7g031500.1 |
| 3074 | . | gene | MSTRG.13906 | 0.315738 | 0.044728 | 0.312066 | |
| 3075 | . | gene | MSTRG.14790 | 1.239632 | 0.044769 | 0.312195 | Bo4g145860.1 |
| 3076 | . | gene | MSTRG.28686 | 1.167859 | 0.044776 | 0.312195 | Bo8g097680.1 |
| 3079 | . | gene | MSTRG.7439 | 2.393656 | 0.044837 | 0.312205 | Bo3g013590.1 |
| 3081 | . | gene | MSTRG.9574 | 0.540318 | 0.044859 | 0.312205 | Bo3g065370.1 |
| 3078 | . | gene | MSTRG.18280 | 0.656648 | 0.044821 | 0.312205 | Bo5g093110.1 |
| 3080 | . | gene | MSTRG.18346 | 0.304804 | 0.044844 | 0.312205 | Bo5g098070.1 |
| 3077 | . | gene | MSTRG.24597 | 0.359184 | 0.0448 | 0.312205 | Bo7g082680.1 |
| 3082 | . | gene | MSTRG.33493 | 0.776942 | 0.044865 | 0.312205 | Bo9g175040.1 |
| 3083 | . | gene | MSTRG.25232 | 0.436177 | 0.044911 | 0.312331 | Bo7g098800.1 |
| 3084 | . | gene | MSTRG.398 | 4.04786 | 0.044912 | 0.312331 | |
| 3085 | ATGCN3 | gene | MSTRG.30788 | 1.5729 | 0.044933 | 0.312378 | Bo9g032240.1 |
| 3086 | . | gene | MSTRG.28485 | 1.323736 | 0.04495 | 0.312389 | Bo8g092250.1 |
| 3088 | . | gene | MSTRG.19082 | 0.274785 | 0.044998 | 0.312521 | Bo5g132140.1 |
| 3087 | . | gene | MSTRG.22278 | 0.749047 | 0.044992 | 0.312521 | Bo6g116070.1 |
| 3093 | . | gene | MSTRG.249 | 0.852262 | 0.045108 | 0.312647 | Bo1g006370.1 |
| 3092 | . | gene | MSTRG.6909 | 1.591011 | 0.045105 | 0.312647 | Bo3g003450.1 |
| 3089 | . | gene | MSTRG.7895 | 0.76687 | 0.045071 | 0.312647 | Bo3g023040.1 |
| 3094 | . | gene | MSTRG.27543 | 1.702027 | 0.045115 | 0.312647 | Bo8g066470.1 |
| 3091 | ATPDIL1-3 | gene | MSTRG.28274 | 1.949823 | 0.045085 | 0.312647 | Bo8g087250.1 |
| 3090 | . | gene | MSTRG.765 | 0.729742 | 0.045077 | 0.312647 | |
| 3095 | . | gene | MSTRG.34567 | 0.961703 | 0.045118 | 0.312647 | |
| 3096 | . | gene | MSTRG.31986 | 0.786987 | 0.045158 | 0.312822 | |
| 3097 | . | gene | MSTRG.715 | 0.12178 | 0.045195 | 0.31298 | Bo1g016930.1 |
| 3098 | XTR2 | gene | MSTRG.29218 | 0.660457 | 0.045227 | 0.313098 | Bo8g106600.1 |
| 3099 | . | gene | MSTRG.27113 | 2.169832 | 0.045274 | 0.313325 | Bo8g050440.1 |

| | | | | | | | |
|---|---|---|---|---|---|---|---|
| 3100 | . | gene | MSTRG.1061 | 0.674173 | 0.04536 | 0.313694 | Bo1g027190.1 |
| 3102 | . | gene | MSTRG.4201 | 1.219582 | 0.045371 | 0.313694 | Bo2g028950.1 |
| 3101 | . | gene | MSTRG.27438 | 1.336194 | 0.045366 | 0.313694 | Bo8g063780.1 |
| 3109 | . | gene | MSTRG.10083 | 1.524125 | 0.045485 | 0.313715 | Bo3g084300.1 |
| 3107 | . | gene | MSTRG.14277 | 0.213812 | 0.045469 | 0.313715 | Bo4g125340.1 |
| 3110 | . | gene | MSTRG.14888 | 0.828905 | 0.045491 | 0.313715 | Bo4g151450.1 |
| 3105 | . | gene | MSTRG.16650 | 1.309877 | 0.045429 | 0.313715 | Bo5g014900.1 |
| 3103 | . | gene | MSTRG.18231 | 0.567469 | 0.045395 | 0.313715 | Bo5g088280.1 |
| 3108 | . | gene | MSTRG.25854 | 1.263513 | 0.045478 | 0.313715 | Bo7g113200.1 |
| 3104 | . | gene | MSTRG.26331 | 0.630194 | 0.045407 | 0.313715 | Bo8g002670.1 |
| 3106 | . | gene | MSTRG.29226 | 1.39604 | 0.045436 | 0.313715 | Bo8g106690.1 |
| 3111 | . | gene | MSTRG.12920 | 1.105734 | 0.045512 | 0.313757 | Bo4g038600.1 |
| 3113 | . | gene | MSTRG.14967 | 3.681561 | 0.045545 | 0.313783 | Bo4g154670.1 |
| 3112 | . | gene | MSTRG.35594 | 0.549375 | 0.045542 | 0.313783 | |
| 3114 | . | gene | MSTRG.15058 | 0.778946 | 0.045601 | 0.314069 | Bo4g161210.1 |
| 3118 | . | gene | MSTRG.7151 | 0.571358 | 0.045665 | 0.314102 | Bo3g008990.1 |
| 3115 | . | gene | MSTRG.11516 | 0.605299 | 0.045632 | 0.314102 | Bo3g155740.1 |
| 3116 | . | gene | MSTRG.29166 | 0.026288 | 0.045638 | 0.314102 | Bo8g105800.1 |
| 3117 | . | gene | MSTRG.29551 | 0.552049 | 0.045651 | 0.314102 | Bo8g114520.1 |
| 3119 | . | gene | MSTRG.15170 | 1.152703 | 0.045689 | 0.314166 | Bo4g169260.1 |
| 3120 | . | gene | MSTRG.8253 | 0.781099 | 0.045706 | 0.314187 | Bo3g030110.1 |
| 3121 | . | gene | MSTRG.4779 | 2.06279 | 0.045743 | 0.314231 | Bo2g062980.1 |
| 3122 | . | gene | MSTRG.26232 | 1.270916 | 0.045746 | 0.314231 | |
| 3123 | . | gene | MSTRG.3726 | 0.313417 | 0.045757 | 0.314231 | |
| 3125 | . | gene | MSTRG.19998 | 0.733688 | 0.045877 | 0.314856 | Bo6g018240.1 |
| 3124 | . | gene | MSTRG.34109 | 1.349457 | 0.045872 | 0.314856 | |
| 3126 | . | gene | MSTRG.21128 | 1.170915 | 0.045942 | 0.315201 | Bo6g076740.1 |
| 3130 | . | gene | MSTRG.346 | 4.739856 | 0.046051 | 0.315543 | Bo1g007610.1 |
| 3128 | . | gene | MSTRG.5215 | 0.168852 | 0.046038 | 0.315543 | Bo2g093380.1 |

| | | | | | | | |
|---|---|---|---|---|---|---|---|
| 3127 | . | gene | MSTRG.7750 | 1.306757 | 0.046025 | 0.315543 | Bo3g019730.1 |
| 3129 | . | gene | MSTRG.13316 | 1.562162 | 0.046048 | 0.315543 | Bo4g059680.1 |
| 3131 | . | gene | MSTRG.11228 | 1.119065 | 0.046089 | 0.315702 | Bo3g144040.1 |
| 3134 | . | gene | MSTRG.7863 | 1.226323 | 0.046151 | 0.315782 | Bo3g022520.1 |
| 3132 | . | gene | MSTRG.27287 | 1.852388 | 0.046142 | 0.315782 | Bo8g058540.1 |
| 3135 | . | gene | MSTRG.27617 | 0.729888 | 0.046159 | 0.315782 | Bo8g067570.1 |
| 3133 | . | gene | MSTRG.30310 | 0.312782 | 0.046145 | 0.315782 | Bo9g017710.1 |
| 3136 | . | gene | MSTRG.7922 | 2.113832 | 0.046191 | 0.315898 | Bo3g023410.1 |
| 3137 | . | gene | MSTRG.36067 | 9.087586 | 0.046213 | 0.31595 | |
| 3138 | . | gene | MSTRG.1558 | 4.408285 | 0.046249 | 0.316091 | Bo1g051790.1 |
| 3139 | . | gene | MSTRG.23746 | 1.390674 | 0.04628 | 0.316208 | Bo7g053000.1 |
| 3141 | . | gene | MSTRG.23780 | 1.742632 | 0.046326 | 0.316316 | Bo7g054660.1 |
| 3140 | . | gene | MSTRG.24945 | 1.627486 | 0.046323 | 0.316316 | Bo7g093670.1 |
| 3142 | . | gene | MSTRG.8911 | 1.260095 | 0.046381 | 0.316529 | Bo3g045160.1 |
| 3143 | . | gene | MSTRG.11252 | 0.768695 | 0.046387 | 0.316529 | Bo3g146450.1 |
| 3144 | . | gene | MSTRG.9022 | 1.864557 | 0.046412 | 0.31653 | Bo3g052140.1 |
| 3145 | . | gene | MSTRG.29295 | 1.647382 | 0.046416 | 0.31653 | Bo8g107940.1 |
| 3146 | . | gene | MSTRG.22678 | 1.490678 | 0.046451 | 0.316667 | Bo6g123740.1 |
| 3147 | . | gene | MSTRG.25576 | 0.828701 | 0.046503 | 0.316922 | Bo7g107710.1 |
| 3148 | . | gene | MSTRG.19957 | 6.25743 | 0.046556 | 0.316968 | Bo6g014360.1 |
| 3154 | . | gene | MSTRG.22538 | 1.941652 | 0.046613 | 0.316968 | Bo6g120770.1 |
| 3151 | . | gene | MSTRG.23815 | 2.161073 | 0.046587 | 0.316968 | Bo7g056170.1 |
| 3150 | . | gene | MSTRG.27430 | 0.576425 | 0.046583 | 0.316968 | Bo8g062700.1 |
| 3153 | . | gene | MSTRG.32667 | 0.5486 | 0.0466 | 0.316968 | Bo9g150070.1 |
| 3149 | . | gene | MSTRG.22422 | 0.326368 | 0.04657 | 0.316968 | |
| 3152 | . | gene | MSTRG.17519 | 1.475794 | 0.046589 | 0.316968 | |
| 3156 | . | gene | MSTRG.9992 | 0.228897 | 0.046663 | 0.317105 | Bo3g082080.1 |
| 3155 | . | gene | MSTRG.2468 | 1.549858 | 0.046654 | 0.317105 | |
| 3158 | . | gene | MSTRG.10055 | 1.549772 | 0.046694 | 0.317111 | Bo3g083790.1 |

| | | | | | | | |
|---|---|---|---|---|---|---|---|
| 3157 | . | gene | MSTRG.29209 | 1.656249 | 0.046686 | 0.317111 | Bo8g106450.1 |
| 3171 | . | gene | MSTRG.5448 | 0.652022 | 0.046968 | 0.317589 | Bo2g103280.1 |
| 3160 | . | gene | MSTRG.6890 | 1.759365 | 0.046804 | 0.317589 | Bo3g003120.1 |
| 3167 | . | gene | MSTRG.11744 | 1.401705 | 0.046911 | 0.317589 | Bo3g169210.1 |
| 3159 | . | gene | MSTRG.12138 | 0.965838 | 0.046802 | 0.317589 | Bo3g185290.1 |
| 3169 | . | gene | MSTRG.14818 | 1.62983 | 0.046936 | 0.317589 | Bo4g148230.1 |
| 3166 | . | gene | MSTRG.18741 | 1.973205 | 0.046905 | 0.317589 | Bo5g121240.1 |
| 3170 | . | gene | MSTRG.20272 | 1.457926 | 0.046953 | 0.317589 | Bo6g031540.1 |
| 3165 | . | gene | MSTRG.23158 | 1.358112 | 0.0469 | 0.317589 | Bo7g021560.1 |
| 3168 | SSI1 | gene | MSTRG.25101 | 1.199842 | 0.04692 | 0.317589 | Bo7g097020.1 |
| 3172 | . | gene | MSTRG.26505 | 0.417379 | 0.046977 | 0.317589 | Bo8g011280.1 |
| 3173 | . | gene | MSTRG.26649 | 0.083231 | 0.046986 | 0.317589 | Bo8g021800.1 |
| 3164 | . | gene | MSTRG.29490 | 0.793886 | 0.046899 | 0.317589 | Bo8g112720.1 |
| 3161 | . | gene | MSTRG.32814 | 0.361466 | 0.046834 | 0.317589 | Bo9g154500.1 |
| 3162 | . | gene | MSTRG.31867 | 0.671752 | 0.046892 | 0.317589 | |
| 3163 | . | gene | MSTRG.6995 | 0.566153 | 0.046896 | 0.317589 | |
| 3174 | ATGSKB6 | gene | MSTRG.9755 | 1.867316 | 0.047054 | 0.317946 | Bo3g070270.1 |
| 3175 | . | gene | MSTRG.14052 | 1.330327 | 0.047069 | 0.317946 | Bo4g113000.1 |
| 3176 | . | gene | MSTRG.2108 | 1.271422 | 0.047116 | 0.31817 | Bo1g090460.1 |
| 3177 | . | gene | MSTRG.30860 | 2.372563 | 0.047155 | 0.318331 | Bo9g036810.1 |
| 3179 | . | gene | MSTRG.5919 | 1.509226 | 0.047237 | 0.318608 | Bo2g133780.1 |
| 3180 | . | gene | MSTRG.20124 | 1.148497 | 0.047241 | 0.318608 | Bo6g026190.1 |
| 3178 | . | gene | MSTRG.26930 | 0.486749 | 0.047235 | 0.318608 | |
| 3181 | . | gene | MSTRG.24280 | 0.532065 | 0.047261 | 0.318645 | Bo7g075600.1 |
| 3182 | . | gene | MSTRG.11232 | 2.140244 | 0.047281 | 0.318677 | Bo3g145200.1 |
| 3183 | . | gene | MSTRG.3138 | 1.797484 | 0.047303 | 0.318725 | Bo1g150180.1 |
| 3184 | . | gene | MSTRG.35190 | 0.450004 | 0.047319 | 0.318738 | |
| 3185 | . | gene | MSTRG.5910 | 1.226915 | 0.047343 | 0.318799 | Bo2g133690.1 |
| 3186 | . | gene | MSTRG.21474 | 1.132653 | 0.047386 | 0.318982 | |

| | | | | | | | |
|---|---|---|---|---|---|---|---|
| 3187 | . | gene | MSTRG.3631 | 0.546165 | 0.047412 | 0.318983 | Bo2g010810.1 |
| 3189 | . | gene | MSTRG.3646 | 1.475774 | 0.047438 | 0.318983 | Bo2g011040.1 |
| 3190 | . | gene | MSTRG.32494 | 0.811975 | 0.047445 | 0.318983 | Bo9g137060.1 |
| 3188 | . | gene | MSTRG.33178 | 1.202531 | 0.047438 | 0.318983 | Bo9g166720.1 |
| 3194 | . | gene | MSTRG.4385 | 1.354682 | 0.047514 | 0.319041 | Bo2g043400.1 |
| 3192 | . | gene | MSTRG.10458 | 0.490959 | 0.0475 | 0.319041 | Bo3g101220.1 |
| 3195 | CUE1 | gene | MSTRG.26819 | 1.965866 | 0.047528 | 0.319041 | Bo8g030830.1 |
| 3193 | . | gene | MSTRG.29853 | 1.958443 | 0.047505 | 0.319041 | Bo9g005580.1 |
| 3191 | . | gene | MSTRG.32735 | 1.646621 | 0.047474 | 0.319041 | Bo9g152230.1 |
| 3196 | . | gene | MSTRG.23022 | 0.52964 | 0.047544 | 0.319045 | Bo7g014030.1 |
| 3197 | . | gene | MSTRG.19891 | 0.873871 | 0.047577 | 0.31914 | Bo6g008230.1 |
| 3198 | . | gene | MSTRG.32959 | 1.405178 | 0.047587 | 0.31914 | Bo9g161700.1 |
| 3202 | . | gene | MSTRG.3783 | 1.184199 | 0.0477 | 0.319349 | Bo2g013390.1 |
| 3204 | . | gene | MSTRG.6718 | 1.378112 | 0.047723 | 0.319349 | Bo2g167680.1 |
| 3200 | . | gene | MSTRG.7981 | 1.30143 | 0.04765 | 0.319349 | Bo3g024290.1 |
| 3203 | . | gene | MSTRG.25142 | 1.340569 | 0.047709 | 0.319349 | Bo7g097590.1 |
| 3205 | . | gene | MSTRG.28739 | 1.439807 | 0.047723 | 0.319349 | Bo8g098390.1 |
| 3201 | . | gene | MSTRG.31768 | 4.005815 | 0.04769 | 0.319349 | Bo9g094080.1 |
| 3199 | . | gene | MSTRG.21585 | 0.995122 | 0.047647 | 0.319349 | |
| 3206 | . | gene | MSTRG.11536 | 1.254103 | 0.04775 | 0.319428 | Bo3g157110.1 |
| 3207 | . | gene | MSTRG.1185 | 0.309398 | 0.047776 | 0.319505 | Bo1g037120.1 |
| 3208 | . | gene | MSTRG.5622 | 0.717865 | 0.047829 | 0.319759 | Bo2g116600.1 |
| 3209 | . | gene | MSTRG.18509 | 0.095409 | 0.047845 | 0.319766 | Bo5g109730.1 |
| 3211 | . | gene | MSTRG.20780 | 1.158282 | 0.047877 | 0.319782 | Bo6g066520.1 |
| 3210 | . | gene | MSTRG.20991 | 0.649641 | 0.047866 | 0.319782 | |
| 3212 | . | gene | MSTRG.16829 | 6.454717 | 0.047927 | 0.320015 | Bo5g019860.1 |
| 3213 | . | gene | MSTRG.15595 | 1.87474 | 0.047963 | 0.320158 | Bo4g187330.1 |
| 3214 | . | gene | MSTRG.33245 | 2.051289 | 0.048011 | 0.320378 | Bo9g169730.1 |
| 3215 | . | gene | MSTRG.32480 | 0.461469 | 0.048125 | 0.321037 | Bo9g136920.1 |

| | | | | | | | |
|---|---|---|---|---|---|---|---|
| 32<br>17 | . | gene | MSTRG.2144<br>2 | 3.4636<br>62 | 0.0482<br>28 | 0.3215<br>24 | Bo6g0837<br>40.1 |
| 32<br>16 | . | gene | MSTRG.2517<br>2 | 0.5446<br>58 | 0.0482<br>24 | 0.3215<br>24 | Bo7g0979<br>60.1 |
| 32<br>19 | . | gene | MSTRG.1137<br>6 | 0.2228<br>62 | 0.0483<br>04 | 0.3218<br>33 | Bo3g1511<br>90.1 |
| 32<br>18 | . | gene | MSTRG.2894<br>2 | 1.5787<br>06 | 0.0482<br>94 | 0.3218<br>33 | Bo8g1015<br>30.1 |
| 32<br>20 | . | gene | MSTRG.3259<br>8 | 0.0407<br>16 | 0.0483<br>66 | 0.3221<br>41 | Bo9g1481<br>00.1 |
| 32<br>21 | . | gene | MSTRG.3035<br>2 | 1.7260<br>4 | 0.0484<br>21 | 0.3224<br>09 | |
| 32<br>22 | . | gene | MSTRG.1944<br>1 | 1.1689<br>47 | 0.0484<br>51 | 0.3225<br>13 | Bo5g1412<br>70.1 |
| 32<br>23 | . | gene | MSTRG.8286 | 1.0290<br>03 | 0.0484<br>78 | 0.3225<br>87 | Bo3g0316<br>50.1 |
| 32<br>24 | . | gene | MSTRG.1855<br>7 | 0.7031<br>31 | 0.0484<br>99 | 0.3226<br>32 | Bo5g1123<br>70.1 |
| 32<br>25 | . | gene | MSTRG.1218<br>9 | 1.8883<br>35 | 0.0485<br>33 | 0.3227<br>55 | Bo4g0011<br>90.1 |
| 32<br>31 | APK2 (Adenylyl-sulfate<br>kinase) | gene | MSTRG.69 | 3.3131<br>72 | 0.0486<br>76 | 0.3229<br>32 | Bo1g0029<br>60.1 |
| 32<br>29 | . | gene | MSTRG.521 | 0.6710<br>57 | 0.0486<br>48 | 0.3229<br>32 | Bo1g0110<br>40.1 |
| 32<br>37 | . | gene | MSTRG.2147 | 0.5140<br>97 | 0.0487<br>4 | 0.3229<br>32 | Bo1g0943<br>40.1 |
| 32<br>28 | . | gene | MSTRG.2641 | 0.3623<br>32 | 0.0486<br>46 | 0.3229<br>32 | Bo1g1230<br>50.1 |
| 32<br>32 | . | gene | MSTRG.4409 | 1.3538<br>74 | 0.0486<br>92 | 0.3229<br>32 | Bo2g0459<br>00.1 |
| 32<br>27 | PHT3%3B1 | gene | MSTRG.7200 | 1.4758<br>2 | 0.0486<br>13 | 0.3229<br>32 | Bo3g0095<br>70.1 |
| 32<br>26 | . | gene | MSTRG.1010<br>7 | 2.4924<br>65 | 0.0485<br>78 | 0.3229<br>32 | Bo3g0855<br>80.1 |
| 32<br>34 | . | gene | MSTRG.1383<br>3 | 1.6852<br>3 | 0.0487<br>22 | 0.3229<br>32 | Bo4g1011<br>90.1 |
| 32<br>33 | . | gene | MSTRG.1782<br>0 | 1.2851<br>31 | 0.0487<br>08 | 0.3229<br>32 | Bo5g0611<br>60.1 |
| 32<br>30 | . | gene | MSTRG.2408<br>3 | 0.5013<br>14 | 0.0486<br>49 | 0.3229<br>32 | Bo7g0652<br>90.1 |
| 32<br>36 | . | gene | MSTRG.3146<br>3 | 0.7847<br>37 | 0.0487<br>33 | 0.3229<br>32 | Bo9g0759<br>00.1 |
| 32<br>35 | . | gene | MSTRG.3404<br>4 | 0.5345<br>8 | 0.0487<br>26 | 0.3229<br>32 | |
| 32<br>39 | . | gene | MSTRG.2441 | 0.7750<br>15 | 0.0487<br>83 | 0.3230<br>18 | Bo1g1086<br>30.1 |
| 32<br>38 | . | gene | MSTRG.2072<br>8 | 1.6677<br>69 | 0.0487<br>69 | 0.3230<br>18 | Bo6g0647<br>30.1 |
| 32<br>40 | . | gene | MSTRG.1567<br>9 | 0.4206<br>03 | 0.0488<br>35 | 0.3232<br>6 | Bo4g1886<br>50.1 |
| 32<br>43 | . | gene | MSTRG.751 | 1.4727 | 0.0489<br>36 | 0.3233<br>6 | Bo1g0173<br>70.1 |
| 32<br>41 | . | gene | MSTRG.6152 | 1.9369<br>26 | 0.0488<br>94 | 0.3233<br>6 | Bo2g1449<br>10.1 |
| 32<br>42 | . | gene | MSTRG.8960 | 1.3859<br>76 | 0.0489<br>18 | 0.3233<br>6 | Bo3g0471<br>90.1 |
| 32<br>46 | . | gene | MSTRG.1345<br>5 | 0.6039<br>84 | 0.0489<br>53 | 0.3233<br>6 | Bo4g0718<br>80.1 |

| | | | | | | | |
|---|---|---|---|---|---|---|---|
| 32<br>47 | . | gene | MSTRG.3234<br>4 | 0.6179<br>51 | 0.0489<br>56 | 0.3233<br>6 | Bo9g1308<br>90.1 |
| 32<br>44 | . | gene | MSTRG.1369<br>8 | 0.9779<br>22 | 0.0489<br>37 | 0.3233<br>6 | |
| 32<br>45 | . | gene | MSTRG.3594<br>7 | 2.6127<br>05 | 0.0489<br>46 | 0.3233<br>6 | |
| 32<br>48 | . | gene | MSTRG.1126 | 3.4858<br>94 | 0.0490<br>3 | 0.3236<br>64 | Bo1g0331<br>10.1 |
| 32<br>49 | . | gene | MSTRG.7721 | 1.3945<br>25 | 0.0490<br>32 | 0.3236<br>64 | Bo3g0192<br>90.1 |
| 32<br>50 | . | gene | MSTRG.8284 | 19.499<br>94 | 0.0490<br>59 | 0.3237<br>22 | Bo3g0316<br>20.1 |
| 32<br>51 | . | gene | MSTRG.9930 | 1.6451<br>48 | 0.0490<br>71 | 0.3237<br>22 | Bo3g0801<br>30.1 |
| 32<br>52 | . | gene | MSTRG.3397<br>2 | 1.4774<br>05 | 0.0491<br>14 | 0.3239<br>06 | |
| 32<br>54 | . | gene | MSTRG.1916<br>3 | 1.9256<br>32 | 0.0491<br>44 | 0.3239<br>09 | Bo5g1334<br>50.1 |
| 32<br>53 | . | gene | MSTRG.3592<br>9 | 1.5151<br>03 | 0.0491<br>32 | 0.3239<br>09 | |
| 32<br>55 | . | gene | MSTRG.2869<br>1 | 1.6586<br>27 | 0.0491<br>75 | 0.3240<br>08 | Bo8g0977<br>50.1 |
| 32<br>56 | . | gene | MSTRG.1772<br>1 | 0.8468<br>18 | 0.0492<br>12 | 0.3241<br>58 | Bo5g0523<br>50.1 |
| 32<br>57 | . | gene | MSTRG.3056<br>1 | 1.2548<br>8 | 0.0492<br>31 | 0.3241<br>82 | Bo9g0243<br>80.1 |
| 32<br>59 | . | gene | MSTRG.1130<br>5 | 0.1850<br>38 | 0.0492<br>65 | 0.3242<br>06 | Bo3g1492<br>10.1 |
| 32<br>58 | . | gene | MSTRG.2424<br>3 | 0.8077<br>76 | 0.0492<br>6 | 0.3242<br>06 | Bo7g0740<br>30.1 |
| 32<br>60 | . | gene | MSTRG.1639<br>2 | 1.2144 | 0.0492<br>81 | 0.3242<br>09 | Bo5g0085<br>20.1 |
| 32<br>61 | . | gene | MSTRG.1265<br>8 | 1.2906<br>04 | 0.0493<br>38 | 0.3243<br>03 | Bo4g0259<br>30.1 |
| 32<br>65 | . | gene | MSTRG.1727<br>4 | 4.4925<br>49 | 0.0493<br>9 | 0.3243<br>03 | Bo5g0343<br>50.1 |
| 32<br>64 | . | gene | MSTRG.2324<br>9 | 1.4481<br>91 | 0.0493<br>75 | 0.3243<br>03 | Bo7g0303<br>00.1 |
| 32<br>63 | . | gene | MSTRG.2401<br>9 | 1.1398<br>23 | 0.0493<br>51 | 0.3243<br>03 | Bo7g0642<br>90.1 |
| 32<br>62 | . | gene | MSTRG.3412<br>6 | 1.6431<br>39 | 0.0493<br>39 | 0.3243<br>03 | |
| 32<br>66 | . | gene | MSTRG.3560<br>2 | 0.6357<br>04 | 0.0493<br>96 | 0.3243<br>03 | |
| 32<br>67 | . | gene | MSTRG.3435<br>1 | 0.0666<br>68 | 0.0494<br>01 | 0.3243<br>03 | |
| 32<br>68 | . | gene | MSTRG.1600<br>3 | 1.4100<br>04 | 0.0494<br>3 | 0.3243<br>25 | Bo4g1986<br>80.1 |
| 32<br>69 | . | gene | MSTRG.2224<br>0 | 0.2175<br>18 | 0.0494<br>34 | 0.3243<br>25 | |
| 32<br>70 | . | gene | MSTRG.2078<br>7 | 1.5704<br>97 | 0.0494<br>58 | 0.3243<br>8 | Bo6g0666<br>20.1 |
| 32<br>71 | . | gene | MSTRG.2584<br>8 | 1.7025<br>22 | 0.0495<br>61 | 0.3249<br>55 | Bo7g1131<br>40.1 |
| 32<br>72 | . | gene | MSTRG.8683 | 1.8273<br>06 | 0.0496<br>01 | 0.3251<br>22 | Bo3g0396<br>20.1 |
| 32<br>73 | . | gene | MSTRG.3101<br>6 | 0.8302<br>9 | 0.0496<br>35 | 0.3252<br>41 | Bo9g0466<br>40.1 |

| | | | | | | | |
|---|---|---|---|---|---|---|---|
| 32 74 | . | gene | MSTRG.8100 | 4.4162 96 | 0.0496 99 | 0.3255 6 | Bo3g0272 00.1 |
| 32 75 | . | gene | MSTRG.2042 0 | 0.6550 74 | 0.0497 57 | 0.3258 42 | Bo6g0426 00.1 |
| 32 76 | . | gene | MSTRG.2889 5 | 0.5952 92 | 0.0498 17 | 0.3260 58 | Bo8g1008 70.1 |
| 32 77 | . | gene | MSTRG.3315 2 | 1.4872 76 | 0.0498 2 | 0.3260 58 | Bo9g1663 70.1 |
| 32 78 | . | gene | MSTRG.1907 5 | 1.2461 44 | 0.0498 56 | 0.3261 95 | Bo5g1320 00.1 |
| 32 79 | . | gene | MSTRG.3480 3 | 1.1884 68 | 0.0498 84 | 0.3262 8 | |
| 32 81 | . | gene | MSTRG.3267 8 | 0.2203 43 | 0.0499 18 | 0.3262 98 | Bo9g1502 10.1 |
| 32 80 | . | gene | MSTRG.3192 3 | 0.7037 98 | 0.0499 07 | 0.3262 98 | |
| 32 86 | . | gene | MSTRG.2403 | 0.2818 11 | 0.0499 99 | 0.3263 33 | Bo1g1070 00.1 |
| 32 84 | . | gene | MSTRG.2786 | 0.0066 1 | 0.0499 9 | 0.3263 33 | Bo1g1346 30.1 |
| 32 82 | . | gene | MSTRG.9272 | 1.2433 59 | 0.0499 5 | 0.3263 33 | Bo3g0588 90.1 |
| 32 85 | . | gene | MSTRG.1100 0 | 0.7882 72 | 0.0499 98 | 0.3263 33 | Bo3g1346 00.1 |
| 32 83 | . | gene | MSTRG.2254 9 | 0.1711 03 | 0.0499 83 | 0.3263 33 | Bo6g1209 00.1 |

#### Supplementary Table 6

| | GO.ID | Term | Annotated | Significant | Expected | classicFisher |
|---|---|---|---|---|---|---|
| 1 | GO:1901135 | carbohydrate derivative metabolic proces... | 2128 | 247 | 118.69 | 2.80E-29 |
| 2 | GO:0019693 | ribose phosphate metabolic process | 994 | 150 | 55.44 | 3.20E-29 |
| 3 | GO:0046496 | nicotinamide nucleotide metabolic proces... | 681 | 118 | 37.98 | 1.70E-28 |
| 4 | GO:0019362 | pyridine nucleotide metabolic process | 686 | 118 | 38.26 | 3.40E-28 |
| 5 | GO:0009117 | nucleotide metabolic process | 1255 | 171 | 70 | 6.60E-28 |
| 6 | GO:0072524 | pyridine-containing compound metabolic p... | 693 | 118 | 38.65 | 8.60E-28 |
| 7 | GO:0006753 | nucleoside phosphate metabolic process | 1262 | 171 | 70.39 | 1.30E-27 |
| 8 | GO:0055086 | nucleobase-containing small molecule met... | 1306 | 172 | 72.84 | 2.40E-26 |
| 9 | GO:0006733 | oxidoreduction coenzyme metabolic proces... | 777 | 122 | 43.34 | 2.00E-25 |
| 10 | GO:0006732 | coenzyme metabolic process | 985 | 141 | 54.94 | 3.50E-25 |
| 11 | GO:0044265 | cellular macromolecule catabolic process | 1077 | 147 | 60.07 | 4.00E-24 |
| 12 | GO:0051186 | cofactor metabolic process | 1348 | 169 | 75.19 | 1.60E-23 |
| 13 | GO:1901575 | organic substance catabolic process | 2548 | 262 | 142.12 | 4.20E-23 |
| 14 | GO:0044710 | single-organism metabolic process | 7693 | 604 | 429.09 | 1.10E-22 |
| 15 | GO:0009057 | macromolecule catabolic process | 1180 | 152 | 65.82 | 2.30E-22 |
| 16 | GO:0044281 | small molecule metabolic process | 4652 | 407 | 259.47 | 2.60E-22 |
| 17 | GO:0044257 | cellular protein catabolic process | 837 | 121 | 46.69 | 4.40E-22 |
| 18 | GO:0044237 | cellular metabolic process | 13616 | 949 | 759.45 | 4.70E-22 |
| 19 | GO:0008152 | metabolic process | 15047 | 1028 | 839.27 | 5.00E-22 |
| 20 | GO:0046686 | response to cadmium ion | 964 | 132 | 53.77 | 7.40E-22 |
| 21 | GO:0051603 | proteolysis involved in cellular protein... | 799 | 117 | 44.57 | 7.50E-22 |
| 22 | GO:0030163 | protein catabolic process | 850 | 121 | 47.41 | 1.60E-21 |
| 23 | GO:0009056 | catabolic process | 3014 | 289 | 168.11 | 6.10E-21 |
| 24 | GO:1901564 | organonitrogen compound metabolic proces... | 3795 | 343 | 211.67 | 9.50E-21 |
| 25 | GO:0044723 | single-organism carbohydrate metabolic p... | 1953 | 210 | 108.93 | 1.00E-20 |
| 26 | GO:0071704 | organic substance metabolic process | 14197 | 973 | 791.86 | 2.60E-20 |
| 27 | GO:0010038 | response to metal ion | 1167 | 145 | 65.09 | 6.60E-20 |

| | | | | | | |
|---|---|---|---|---|---|---|
| 28 | GO:0006006 | glucose metabolic process | 549 | 89 | 30.62 | 9.20E-20 |
| 29 | GO:0005996 | monosaccharide metabolic process | 674 | 101 | 37.59 | 1.00E-19 |
| 30 | GO:0009259 | ribonucleotide metabolic process | 747 | 107 | 41.67 | 2.40E-19 |
| 31 | GO:0009132 | nucleoside diphosphate metabolic process | 423 | 75 | 23.59 | 4.20E-19 |
| 32 | GO:0019637 | organophosphate metabolic process | 1869 | 198 | 104.25 | 7.80E-19 |
| 33 | GO:0006096 | glycolytic process | 418 | 74 | 23.31 | 7.90E-19 |
| 34 | GO:0006165 | nucleoside diphosphate phosphorylation | 418 | 74 | 23.31 | 7.90E-19 |
| 35 | GO:0006757 | ATP generation from ADP | 418 | 74 | 23.31 | 7.90E-19 |
| 36 | GO:0009135 | purine nucleoside diphosphate metabolic ... | 420 | 74 | 23.43 | 1.00E-18 |
| 37 | GO:0009179 | purine ribonucleoside diphosphate metabo... | 420 | 74 | 23.43 | 1.00E-18 |
| 38 | GO:0009185 | ribonucleoside diphosphate metabolic pro... | 420 | 74 | 23.43 | 1.00E-18 |
| 39 | GO:0046031 | ADP metabolic process | 420 | 74 | 23.43 | 1.00E-18 |
| 40 | GO:0019318 | hexose metabolic process | 591 | 91 | 32.96 | 1.10E-18 |
| 41 | GO:0046939 | nucleotide phosphorylation | 421 | 74 | 23.48 | 1.20E-18 |
| 42 | GO:0009126 | purine nucleoside monophosphate metaboli... | 481 | 80 | 26.83 | 1.50E-18 |
| 43 | GO:0009167 | purine ribonucleoside monophosphate meta... | 481 | 80 | 26.83 | 1.50E-18 |
| 44 | GO:0019941 | modification-dependent protein catabolic... | 640 | 95 | 35.7 | 2.40E-18 |
| 45 | GO:0043632 | modification-dependent macromolecule cat... | 640 | 95 | 35.7 | 2.40E-18 |
| 46 | GO:0009987 | cellular process | 17535 | 1144 | 978.04 | 2.60E-18 |
| 47 | GO:0006511 | ubiquitin-dependent protein catabolic pr... | 630 | 94 | 35.14 | 2.60E-18 |
| 48 | GO:0044724 | single-organism carbohydrate catabolic p... | 623 | 93 | 34.75 | 3.80E-18 |
| 49 | GO:0009123 | nucleoside monophosphate metabolic proce... | 489 | 80 | 27.27 | 4.00E-18 |
| 50 | GO:0009161 | ribonucleoside monophosphate metabolic p... | 489 | 80 | 27.27 | 4.00E-18 |
| 51 | GO:0009150 | purine ribonucleotide metabolic process | 500 | 81 | 27.89 | 4.50E-18 |
| 52 | GO:0009199 | ribonucleoside triphosphate metabolic pr... | 470 | 78 | 26.22 | 4.50E-18 |
| 53 | GO:0009205 | purine ribonucleoside triphosphate metab... | 470 | 78 | 26.22 | 4.50E-18 |
| 54 | GO:0046034 | ATP metabolic process | 470 | 78 | 26.22 | 4.50E-18 |
| 55 | GO:0009144 | purine nucleoside triphosphate metabolic... | 472 | 78 | 26.33 | 5.80E-18 |
| 56 | GO:0009141 | nucleoside triphosphate metabolic proces... | 475 | 78 | 26.49 | 8.40E-18 |

| | | | | | | |
|---|---|---|---|---|---|---|
| 57 | GO:0044238 | primary metabolic process | 12871 | 886 | 717.9 | 8.50E-18 |
| 58 | GO:0006094 | gluconeogenesis | 352 | 65 | 19.63 | 1.20E-17 |
| 59 | GO:0043248 | proteasome assembly | 381 | 68 | 21.25 | 1.30E-17 |
| 60 | GO:0051788 | response to misfolded protein | 381 | 68 | 21.25 | 1.30E-17 |
| 61 | GO:0005975 | carbohydrate metabolic process | 2247 | 222 | 125.33 | 1.70E-17 |
| 62 | GO:0044248 | cellular catabolic process | 2418 | 234 | 134.87 | 2.30E-17 |
| 63 | GO:0046364 | monosaccharide biosynthetic process | 376 | 67 | 20.97 | 2.50E-17 |
| 64 | GO:0044711 | single-organism biosynthetic process | 4962 | 407 | 276.76 | 2.80E-17 |
| 65 | GO:0019319 | hexose biosynthetic process | 358 | 65 | 19.97 | 2.90E-17 |
| 66 | GO:0006508 | proteolysis | 1506 | 165 | 84 | 4.00E-17 |
| 67 | GO:0006163 | purine nucleotide metabolic process | 561 | 85 | 31.29 | 4.40E-17 |
| 68 | GO:0044763 | single-organism cellular process | 10539 | 747 | 587.83 | 7.10E-17 |
| 69 | GO:0016052 | carbohydrate catabolic process | 708 | 98 | 39.49 | 8.40E-17 |
| 70 | GO:0006807 | nitrogen compound metabolic process | 8509 | 626 | 474.6 | 9.00E-17 |
| 71 | GO:0080129 | proteasome core complex assembly | 286 | 56 | 15.95 | 1.40E-16 |
| 72 | GO:0006950 | response to stress | 6641 | 510 | 370.41 | 1.80E-16 |
| 73 | GO:0072521 | purine-containing compound metabolic pro... | 592 | 86 | 33.02 | 3.80E-16 |
| 74 | GO:0010498 | proteasomal protein catabolic process | 407 | 68 | 22.7 | 4.50E-16 |
| 75 | GO:1901360 | organic cyclic compound metabolic proces... | 8191 | 601 | 456.87 | 1.20E-15 |
| 76 | GO:0043933 | macromolecular complex subunit organizat... | 1106 | 129 | 61.69 | 1.30E-15 |
| 77 | GO:0006793 | phosphorus metabolic process | 3315 | 290 | 184.9 | 1.60E-15 |
| 78 | GO:0071822 | protein complex subunit organization | 1062 | 125 | 59.23 | 1.80E-15 |
| 79 | GO:0046483 | heterocycle metabolic process | 7367 | 549 | 410.91 | 2.60E-15 |
| 80 | GO:0009628 | response to abiotic stimulus | 4939 | 396 | 275.48 | 3.50E-15 |
| 81 | GO:0016051 | carbohydrate biosynthetic process | 1417 | 152 | 79.04 | 4.70E-15 |
| 82 | GO:0006091 | generation of precursor metabolites and ... | 957 | 115 | 53.38 | 6.10E-15 |
| 83 | GO:0006796 | phosphate-containing compound metabolic ... | 3170 | 277 | 176.81 | 9.40E-15 |
| 84 | GO:0044699 | single-organism process | 15394 | 1007 | 858.63 | 2.70E-14 |
| 85 | GO:0065003 | macromolecular complex assembly | 1079 | 123 | 60.18 | 3.20E-14 |

| | | | | | | |
|---|---|---|---|---|---|---|
| 86 | GO:0035966 | response to topologically incorrect prot... | 730 | 94 | 40.72 | 3.30E-14 |
| 87 | GO:0006790 | sulfur compound metabolic process | 1232 | 135 | 68.72 | 3.60E-14 |
| 88 | GO:0006090 | pyruvate metabolic process | 767 | 97 | 42.78 | 3.90E-14 |
| 89 | GO:0006725 | cellular aromatic compound metabolic pro... | 7883 | 574 | 439.69 | 4.00E-14 |
| 90 | GO:0006139 | nucleobase-containing compound metabolic... | 6691 | 501 | 373.2 | 4.30E-14 |
| 91 | GO:0070271 | protein complex biogenesis | 1047 | 120 | 58.4 | 4.60E-14 |
| 92 | GO:0006461 | protein complex assembly | 1035 | 119 | 57.73 | 4.70E-14 |
| 93 | GO:0044267 | cellular protein metabolic process | 4625 | 370 | 257.97 | 5.70E-14 |
| 94 | GO:0043436 | oxoacid metabolic process | 3444 | 291 | 192.09 | 1.10E-13 |
| 95 | GO:0006082 | organic acid metabolic process | 3445 | 291 | 192.15 | 1.20E-13 |
| 96 | GO:0006996 | organelle organization | 3328 | 283 | 185.62 | 1.20E-13 |
| 97 | GO:0016043 | cellular component organization | 5109 | 399 | 284.96 | 1.50E-13 |
| 98 | GO:1901137 | carbohydrate derivative biosynthetic pro... | 1167 | 128 | 65.09 | 1.60E-13 |
| 99 | GO:0034641 | cellular nitrogen compound metabolic pro... | 7743 | 561 | 431.88 | 2.60E-13 |
| 100 | GO:0009058 | biosynthetic process | 10020 | 695 | 558.88 | 4.50E-13 |
| 101 | GO:0034622 | cellular macromolecular complex assembly | 1046 | 117 | 58.34 | 5.10E-13 |
| 102 | GO:1901576 | organic substance biosynthetic process | 9774 | 680 | 545.16 | 5.60E-13 |
| 103 | GO:0019538 | protein metabolic process | 5124 | 397 | 285.8 | 5.70E-13 |
| 104 | GO:0043623 | cellular protein complex assembly | 1002 | 113 | 55.89 | 7.60E-13 |
| 105 | GO:0044260 | cellular macromolecule metabolic process | 9350 | 652 | 521.51 | 1.80E-12 |
| 106 | GO:0043161 | proteasome-mediated ubiquitin-dependent ... | 250 | 45 | 13.94 | 3.00E-12 |
| 107 | GO:0006970 | response to osmotic stress | 1616 | 156 | 90.14 | 1.10E-11 |
| 108 | GO:0006098 | pentose-phosphate shunt | 290 | 48 | 16.18 | 1.30E-11 |
| 109 | GO:0051156 | glucose 6-phosphate metabolic process | 290 | 48 | 16.18 | 1.30E-11 |
| 110 | GO:0009314 | response to radiation | 2140 | 193 | 119.36 | 1.40E-11 |
| 111 | GO:0043170 | macromolecule metabolic process | 10029 | 686 | 559.38 | 1.50E-11 |
| 112 | GO:0006739 | NADP metabolic process | 302 | 49 | 16.84 | 1.70E-11 |
| 113 | GO:0019752 | carboxylic acid metabolic process | 3262 | 269 | 181.94 | 1.70E-11 |
| 114 | GO:0071840 | cellular component organization or bioge... | 5748 | 428 | 320.6 | 1.90E-11 |

| | | | | | | |
|---|---|---|---|---|---|---|
| 115 | GO:0043094 | cellular metabolic compound salvage | 347 | 53 | 19.35 | 2.70E-11 |
| 116 | GO:0009651 | response to salt stress | 1526 | 148 | 85.12 | 2.70E-11 |
| 117 | GO:0044272 | sulfur compound biosynthetic process | 845 | 96 | 47.13 | 2.80E-11 |
| 118 | GO:0009853 | photorespiration | 322 | 50 | 17.96 | 5.30E-11 |
| 119 | GO:0050896 | response to stimulus | 10816 | 727 | 603.28 | 7.50E-11 |
| 120 | GO:0022607 | cellular component assembly | 1470 | 141 | 81.99 | 1.80E-10 |
| 121 | GO:0009416 | response to light stimulus | 2039 | 181 | 113.73 | 2.30E-10 |
| 122 | GO:0044283 | small molecule biosynthetic process | 2301 | 199 | 128.34 | 2.50E-10 |
| 123 | GO:0044712 | single-organism catabolic process | 1861 | 168 | 103.8 | 3.20E-10 |
| 124 | GO:0016144 | S-glycoside biosynthetic process | 300 | 45 | 16.73 | 1.50E-09 |
| 125 | GO:0019758 | glycosinolate biosynthetic process | 300 | 45 | 16.73 | 1.50E-09 |
| 126 | GO:0019761 | glucosinolate biosynthetic process | 300 | 45 | 16.73 | 1.50E-09 |
| 127 | GO:1901659 | glycosyl compound biosynthetic process | 311 | 46 | 17.35 | 1.60E-09 |
| 128 | GO:0032787 | monocarboxylic acid metabolic process | 2388 | 201 | 133.19 | 1.90E-09 |
| 129 | GO:1901657 | glycosyl compound metabolic process | 412 | 55 | 22.98 | 2.00E-09 |
| 130 | GO:0016143 | S-glycoside metabolic process | 368 | 51 | 20.53 | 2.10E-09 |
| 131 | GO:0019757 | glycosinolate metabolic process | 368 | 51 | 20.53 | 2.10E-09 |
| 132 | GO:0019760 | glucosinolate metabolic process | 368 | 51 | 20.53 | 2.10E-09 |
| 133 | GO:0010035 | response to inorganic substance | 2535 | 210 | 141.39 | 2.90E-09 |
| 134 | GO:0016310 | phosphorylation | 1688 | 151 | 94.15 | 5.00E-09 |
| 135 | GO:0051641 | cellular localization | 2264 | 189 | 126.28 | 1.10E-08 |
| 136 | GO:0044085 | cellular component biogenesis | 2363 | 195 | 131.8 | 1.50E-08 |
| 137 | GO:0006081 | cellular aldehyde metabolic process | 693 | 76 | 38.65 | 1.60E-08 |
| 138 | GO:0051649 | establishment of localization in cell | 2099 | 176 | 117.08 | 2.80E-08 |
| 139 | GO:0009250 | glucan biosynthetic process | 433 | 53 | 24.15 | 7.30E-08 |
| 140 | GO:1901566 | organonitrogen compound biosynthetic pro... | 2847 | 222 | 158.8 | 1.40E-07 |
| 141 | GO:0006396 | RNA processing | 1247 | 114 | 69.55 | 1.40E-07 |
| 142 | GO:0055114 | oxidation-reduction process | 684 | 72 | 38.15 | 1.90E-07 |
| 143 | GO:0019748 | secondary metabolic process | 1119 | 104 | 62.41 | 2.30E-07 |

| | | | | | | |
|---|---|---|---|---|---|---|
| 144 | GO:0008104 | protein localization | 1973 | 163 | 110.05 | 2.50E-07 |
| 145 | GO:0042221 | response to chemical | 6773 | 463 | 377.78 | 2.90E-07 |
| 146 | GO:0070727 | cellular macromolecule localization | 1876 | 156 | 104.64 | 3.10E-07 |
| 147 | GO:0046907 | intracellular transport | 1967 | 162 | 109.71 | 3.30E-07 |
| 148 | GO:0033365 | protein localization to organelle | 884 | 86 | 49.31 | 3.90E-07 |
| 149 | GO:0072594 | establishment of protein localization to... | 872 | 85 | 48.64 | 4.20E-07 |
| 150 | GO:0006073 | cellular glucan metabolic process | 612 | 65 | 34.14 | 5.30E-07 |
| 151 | GO:0044042 | glucan metabolic process | 612 | 65 | 34.14 | 5.30E-07 |
| 152 | GO:0016071 | mRNA metabolic process | 536 | 59 | 29.9 | 5.50E-07 |
| 153 | GO:0034613 | cellular protein localization | 1820 | 151 | 101.51 | 5.50E-07 |
| 154 | GO:0006464 | cellular protein modification process | 2990 | 227 | 166.77 | 7.50E-07 |
| 155 | GO:0036211 | protein modification process | 2990 | 227 | 166.77 | 7.50E-07 |
| 156 | GO:0015031 | protein transport | 1875 | 154 | 104.58 | 7.70E-07 |
| 157 | GO:0045184 | establishment of protein localization | 1875 | 154 | 104.58 | 7.70E-07 |
| 158 | GO:0048193 | Golgi vesicle transport | 555 | 60 | 30.96 | 8.10E-07 |
| 159 | GO:0044249 | cellular biosynthetic process | 9100 | 596 | 507.57 | 9.60E-07 |
| 160 | GO:0006886 | intracellular protein transport | 1793 | 148 | 100.01 | 9.60E-07 |
| 161 | GO:0043412 | macromolecule modification | 3608 | 265 | 201.24 | 1.20E-06 |
| 162 | GO:0019682 | glyceraldehyde-3-phosphate metabolic pro... | 536 | 58 | 29.9 | 1.20E-06 |
| 163 | GO:0008380 | RNA splicing | 417 | 48 | 23.26 | 1.80E-06 |
| 164 | GO:0044550 | secondary metabolite biosynthetic proces... | 546 | 58 | 30.45 | 2.10E-06 |
| 165 | GO:0000096 | sulfur amino acid metabolic process | 598 | 62 | 33.35 | 2.20E-06 |
| 166 | GO:0070647 | protein modification by small protein co... | 523 | 56 | 29.17 | 2.50E-06 |
| 167 | GO:0007005 | mitochondrion organization | 215 | 30 | 11.99 | 3.50E-06 |
| 168 | GO:0033036 | macromolecule localization | 2407 | 185 | 134.25 | 4.30E-06 |
| 169 | GO:0051179 | localization | 5647 | 386 | 314.97 | 4.50E-06 |
| 170 | GO:0009266 | response to temperature stimulus | 1715 | 139 | 95.66 | 5.40E-06 |
| 171 | GO:0006833 | water transport | 300 | 37 | 16.73 | 5.50E-06 |
| 172 | GO:0042044 | fluid transport | 300 | 37 | 16.73 | 5.50E-06 |

| | | | | | | |
|---|---|---|---|---|---|---|
| 173 | GO:0000097 | sulfur amino acid biosynthetic process | 525 | 55 | 29.28 | 5.90E-06 |
| 174 | GO:0034660 | ncRNA metabolic process | 591 | 60 | 32.96 | 6.00E-06 |
| 175 | GO:0000103 | sulfate assimilation | 20 | 8 | 1.12 | 6.30E-06 |
| 176 | GO:1901605 | alpha-amino acid metabolic process | 936 | 85 | 52.21 | 7.10E-06 |
| 177 | GO:0019252 | starch biosynthetic process | 280 | 35 | 15.62 | 7.20E-06 |
| 178 | GO:0009735 | response to cytokinin | 491 | 52 | 27.39 | 7.70E-06 |
| 179 | GO:0006520 | cellular amino acid metabolic process | 1185 | 102 | 66.1 | 9.00E-06 |
| 180 | GO:0051604 | protein maturation | 119 | 20 | 6.64 | 9.30E-06 |
| 181 | GO:1901607 | alpha-amino acid biosynthetic process | 672 | 65 | 37.48 | 1.20E-05 |
| 182 | GO:0006626 | protein targeting to mitochondrion | 184 | 26 | 10.26 | 1.20E-05 |
| 183 | GO:0070585 | protein localization to mitochondrion | 184 | 26 | 10.26 | 1.20E-05 |
| 184 | GO:0072655 | establishment of protein localization to... | 184 | 26 | 10.26 | 1.20E-05 |
| 185 | GO:0009100 | glycoprotein metabolic process | 349 | 40 | 19.47 | 1.40E-05 |
| 186 | GO:0032446 | protein modification by small protein co... | 374 | 42 | 20.86 | 1.40E-05 |
| 187 | GO:0006486 | protein glycosylation | 337 | 39 | 18.8 | 1.40E-05 |
| 188 | GO:0009101 | glycoprotein biosynthetic process | 337 | 39 | 18.8 | 1.40E-05 |
| 189 | GO:0043413 | macromolecule glycosylation | 337 | 39 | 18.8 | 1.40E-05 |
| 190 | GO:0070085 | glycosylation | 337 | 39 | 18.8 | 1.40E-05 |
| 191 | GO:0015833 | peptide transport | 2053 | 159 | 114.51 | 1.40E-05 |
| 192 | GO:0015931 | nucleobase-containing compound transport | 242 | 31 | 13.5 | 1.40E-05 |
| 193 | GO:0006605 | protein targeting | 1628 | 131 | 90.8 | 1.40E-05 |
| 194 | GO:0051234 | establishment of localization | 5308 | 361 | 296.06 | 1.60E-05 |
| 195 | GO:0042886 | amide transport | 2064 | 159 | 115.12 | 1.90E-05 |
| 196 | GO:0016192 | vesicle-mediated transport | 865 | 78 | 48.25 | 2.10E-05 |
| 197 | GO:0006555 | methionine metabolic process | 168 | 24 | 9.37 | 2.20E-05 |
| 198 | GO:0009640 | photomorphogenesis | 420 | 45 | 23.43 | 2.30E-05 |
| 199 | GO:0007276 | gamete generation | 310 | 36 | 17.29 | 2.70E-05 |
| 200 | GO:0007292 | female gamete generation | 227 | 29 | 12.66 | 2.80E-05 |
| 201 | GO:0000271 | polysaccharide biosynthetic process | 892 | 79 | 49.75 | 3.50E-05 |

| | | | | | | |
|---|---|---|---|---|---|---|
| 202 | GO:0043085 | positive regulation of catalytic activit... | 174 | 24 | 9.71 | 3.90E-05 |
| 203 | GO:0044282 | small molecule catabolic process | 1083 | 92 | 60.41 | 4.00E-05 |
| 204 | GO:0042592 | homeostatic process | 644 | 61 | 35.92 | 4.10E-05 |
| 205 | GO:0009639 | response to red or far red light | 743 | 68 | 41.44 | 4.50E-05 |
| 206 | GO:0051128 | regulation of cellular component organiz... | 565 | 55 | 31.51 | 4.70E-05 |
| 207 | GO:0010605 | negative regulation of macromolecule met... | 1161 | 97 | 64.76 | 4.70E-05 |
| 208 | GO:0008652 | cellular amino acid biosynthetic process | 788 | 71 | 43.95 | 5.10E-05 |
| 209 | GO:0044093 | positive regulation of molecular functio... | 178 | 24 | 9.93 | 5.70E-05 |
| 210 | GO:0009560 | embryo sac egg cell differentiation | 224 | 28 | 12.49 | 5.70E-05 |
| 211 | GO:0009165 | nucleotide biosynthetic process | 503 | 50 | 28.06 | 6.00E-05 |
| 212 | GO:1901293 | nucleoside phosphate biosynthetic proces... | 504 | 50 | 28.11 | 6.30E-05 |
| 213 | GO:0065008 | regulation of biological quality | 1975 | 150 | 110.16 | 6.60E-05 |
| 214 | GO:0033692 | cellular polysaccharide biosynthetic pro... | 726 | 66 | 40.49 | 7.10E-05 |
| 215 | GO:0006810 | transport | 5202 | 349 | 290.15 | 7.20E-05 |
| 216 | GO:0000302 | response to reactive oxygen species | 561 | 54 | 31.29 | 7.30E-05 |
| 217 | GO:0006979 | response to oxidative stress | 842 | 74 | 46.96 | 7.90E-05 |
| 218 | GO:0010033 | response to organic substance | 5208 | 349 | 290.48 | 7.90E-05 |
| 219 | GO:0044264 | cellular polysaccharide metabolic proces... | 900 | 78 | 50.2 | 8.10E-05 |
| 220 | GO:0009086 | methionine biosynthetic process | 160 | 22 | 8.92 | 8.40E-05 |
| 221 | GO:0009408 | response to heat | 497 | 49 | 27.72 | 8.60E-05 |
| 222 | GO:0016567 | protein ubiquitination | 341 | 37 | 19.02 | 9.10E-05 |
| 223 | GO:0006913 | nucleocytoplasmic transport | 354 | 38 | 19.74 | 9.20E-05 |
| 224 | GO:0051169 | nuclear transport | 354 | 38 | 19.74 | 9.20E-05 |
| 225 | GO:0009892 | negative regulation of metabolic process | 1218 | 99 | 67.94 | 0.00011 |
| 226 | GO:0034470 | ncRNA processing | 464 | 46 | 25.88 | 0.00012 |
| 227 | GO:0001558 | regulation of cell growth | 199 | 25 | 11.1 | 0.00013 |
| 228 | GO:0009404 | toxin metabolic process | 386 | 40 | 21.53 | 0.00013 |
| 229 | GO:0005982 | starch metabolic process | 361 | 38 | 20.14 | 0.00014 |
| 230 | GO:0006635 | fatty acid beta-oxidation | 310 | 34 | 17.29 | 0.00014 |

| | | | | | | |
|---|---|---|---|---|---|---|
| 231 | GO:0010439 | regulation of glucosinolate biosynthetic... | 16 | 6 | 0.89 | 0.00015 |
| 232 | GO:0006401 | RNA catabolic process | 213 | 26 | 11.88 | 0.00015 |
| 233 | GO:0009066 | aspartate family amino acid metabolic pr... | 238 | 28 | 13.27 | 0.00016 |
| 234 | GO:0007049 | cell cycle | 1306 | 104 | 72.84 | 0.00016 |
| 235 | GO:0000023 | maltose metabolic process | 226 | 27 | 12.61 | 0.00016 |
| 236 | GO:0005976 | polysaccharide metabolic process | 1082 | 89 | 60.35 | 0.00016 |
| 237 | GO:0019439 | aromatic compound catabolic process | 471 | 46 | 26.27 | 0.00017 |
| 238 | GO:0044270 | cellular nitrogen compound catabolic pro... | 471 | 46 | 26.27 | 0.00017 |
| 239 | GO:0034440 | lipid oxidation | 340 | 36 | 18.96 | 0.00018 |
| 240 | GO:0030243 | cellulose metabolic process | 216 | 26 | 12.05 | 0.00019 |
| 241 | GO:0032844 | regulation of homeostatic process | 64 | 12 | 3.57 | 0.00019 |
| 242 | GO:0046700 | heterocycle catabolic process | 475 | 46 | 26.49 | 0.00021 |
| 243 | GO:0009743 | response to carbohydrate | 740 | 65 | 41.27 | 0.00021 |
| 244 | GO:0034637 | cellular carbohydrate biosynthetic proce... | 783 | 68 | 43.67 | 0.00021 |
| 245 | GO:0006974 | cellular response to DNA damage stimulus | 395 | 40 | 22.03 | 0.00021 |
| 246 | GO:0019395 | fatty acid oxidation | 317 | 34 | 17.68 | 0.00021 |
| 247 | GO:0022604 | regulation of cell morphogenesis | 182 | 23 | 10.15 | 0.00021 |
| 248 | GO:0070646 | protein modification by small protein re... | 230 | 27 | 12.83 | 0.00022 |
| 249 | GO:0009791 | post-embryonic development | 3851 | 263 | 214.8 | 0.00023 |
| 250 | GO:0044262 | cellular carbohydrate metabolic process | 1066 | 87 | 59.46 | 0.00025 |
| 251 | GO:0009260 | ribonucleotide biosynthetic process | 307 | 33 | 17.12 | 0.00025 |
| 252 | GO:0046390 | ribose phosphate biosynthetic process | 307 | 33 | 17.12 | 0.00025 |
| 253 | GO:0009067 | aspartate family amino acid biosynthetic... | 196 | 24 | 10.93 | 0.00026 |
| 254 | GO:0042023 | DNA endoreduplication | 220 | 26 | 12.27 | 0.00026 |
| 255 | GO:0009407 | toxin catabolic process | 359 | 37 | 20.02 | 0.00026 |
| 256 | GO:0090487 | secondary metabolite catabolic process | 359 | 37 | 20.02 | 0.00026 |
| 257 | GO:0010629 | negative regulation of gene expression | 1053 | 86 | 58.73 | 0.00027 |
| 258 | GO:0051510 | regulation of unidimensional cell growth | 173 | 22 | 9.65 | 0.00027 |
| 259 | GO:0065009 | regulation of molecular function | 386 | 39 | 21.53 | 0.00027 |

| | | | | | | |
|---|---|---|---|---|---|---|
| 260 | GO:0046148 | pigment biosynthetic process | 453 | 44 | 25.27 | 0.00027 |
| 261 | GO:0044707 | single-multicellular organism process | 6086 | 396 | 339.46 | 0.00028 |
| 262 | GO:0034655 | nucleobase-containing compound catabolic... | 234 | 27 | 13.05 | 0.00029 |
| 263 | GO:0051273 | beta-glucan metabolic process | 234 | 27 | 13.05 | 0.00029 |
| 264 | GO:0000394 | RNA splicing, via endonucleolytic cleava... | 222 | 26 | 12.38 | 0.00029 |
| 265 | GO:0044786 | cell cycle DNA replication | 223 | 26 | 12.44 | 0.00032 |
| 266 | GO:0042440 | pigment metabolic process | 567 | 52 | 31.63 | 0.00032 |
| 267 | GO:0006457 | protein folding | 431 | 42 | 24.04 | 0.00034 |
| 268 | GO:0048285 | organelle fission | 513 | 48 | 28.61 | 0.00034 |
| 269 | GO:0030258 | lipid modification | 353 | 36 | 19.69 | 0.00038 |
| 270 | GO:0009060 | aerobic respiration | 99 | 15 | 5.52 | 0.00038 |
| 271 | GO:1901361 | organic cyclic compound catabolic proces... | 489 | 46 | 27.27 | 0.0004 |
| 272 | GO:0050790 | regulation of catalytic activity | 303 | 32 | 16.9 | 0.00042 |
| 273 | GO:0006007 | glucose catabolic process | 215 | 25 | 11.99 | 0.00043 |
| 274 | GO:0015780 | nucleotide-sugar transport | 26 | 7 | 1.45 | 0.00043 |
| 275 | GO:1901264 | carbohydrate derivative transport | 60 | 11 | 3.35 | 0.00043 |
| 276 | GO:0010256 | endomembrane system organization | 423 | 41 | 23.59 | 0.00044 |
| 277 | GO:0006997 | nucleus organization | 145 | 19 | 8.09 | 0.00046 |
| 278 | GO:0040008 | regulation of growth | 549 | 50 | 30.62 | 0.00049 |
| 279 | GO:0065007 | biological regulation | 9001 | 563 | 502.05 | 0.00049 |
| 280 | GO:0032204 | regulation of telomere maintenance | 61 | 11 | 3.4 | 0.0005 |
| 281 | GO:0019320 | hexose catabolic process | 218 | 25 | 12.16 | 0.00052 |
| 282 | GO:0046365 | monosaccharide catabolic process | 218 | 25 | 12.16 | 0.00052 |
| 283 | GO:0007275 | multicellular organism development | 5904 | 382 | 329.31 | 0.00057 |
| 284 | GO:0019321 | pentose metabolic process | 72 | 12 | 4.02 | 0.00059 |
| 285 | GO:0006259 | DNA metabolic process | 1159 | 91 | 64.65 | 0.00065 |
| 286 | GO:0048638 | regulation of developmental growth | 500 | 46 | 27.89 | 0.00065 |
| 287 | GO:0022412 | cellular process involved in reproductio... | 272 | 29 | 15.17 | 0.00066 |
| 288 | GO:0022402 | cell cycle process | 1025 | 82 | 57.17 | 0.00068 |

| | | | | | | |
|---|---|---|---|---|---|---|
| 289 | GO:0010228 | vegetative to reproductive phase transit... | 744 | 63 | 41.5 | 0.00069 |
| 290 | GO:0042542 | response to hydrogen peroxide | 338 | 34 | 18.85 | 0.00069 |
| 291 | GO:0030244 | cellulose biosynthetic process | 127 | 17 | 7.08 | 0.00071 |
| 292 | GO:0060249 | anatomical structure homeostasis | 74 | 12 | 4.13 | 0.00077 |
| 293 | GO:0006839 | mitochondrial transport | 249 | 27 | 13.89 | 0.00077 |
| 294 | GO:0006397 | mRNA processing | 187 | 22 | 10.43 | 0.00078 |
| 295 | GO:0006325 | chromatin organization | 851 | 70 | 47.47 | 0.00079 |
| 296 | GO:0000741 | karyogamy | 96 | 14 | 5.35 | 0.00087 |
| 297 | GO:0048284 | organelle fusion | 96 | 14 | 5.35 | 0.00087 |
| 298 | GO:0042742 | defense response to bacterium | 826 | 68 | 46.07 | 0.00091 |
| 299 | GO:0009561 | megagametogenesis | 304 | 31 | 16.96 | 0.00092 |
| 300 | GO:0033554 | cellular response to stress | 2433 | 171 | 135.7 | 0.00093 |
| 301 | GO:0009657 | plastid organization | 623 | 54 | 34.75 | 0.00095 |
| 302 | GO:0051276 | chromosome organization | 1099 | 86 | 61.3 | 0.00099 |
| 303 | GO:0010332 | response to gamma radiation | 98 | 14 | 5.47 | 0.00107 |
| 304 | GO:0006611 | protein export from nucleus | 144 | 18 | 8.03 | 0.00113 |
| 305 | GO:0090407 | organophosphate biosynthetic process | 1211 | 93 | 67.55 | 0.00113 |
| 306 | GO:0050793 | regulation of developmental process | 1569 | 116 | 87.51 | 0.00116 |
| 307 | GO:0051274 | beta-glucan biosynthetic process | 145 | 18 | 8.09 | 0.00122 |
| 308 | GO:0051640 | organelle localization | 244 | 26 | 13.61 | 0.00123 |
| 309 | GO:0016109 | tetraterpenoid biosynthetic process | 169 | 20 | 9.43 | 0.00123 |
| 310 | GO:0016117 | carotenoid biosynthetic process | 169 | 20 | 9.43 | 0.00123 |
| 311 | GO:0009218 | pyrimidine ribonucleotide metabolic proc... | 257 | 27 | 14.33 | 0.00123 |
| 312 | GO:0009220 | pyrimidine ribonucleotide biosynthetic p... | 257 | 27 | 14.33 | 0.00123 |
| 313 | GO:0043247 | telomere maintenance in response to DNA ... | 58 | 10 | 3.24 | 0.00127 |
| 314 | GO:0042180 | cellular ketone metabolic process | 297 | 30 | 16.57 | 0.00129 |
| 315 | GO:0016579 | protein deubiquitination | 89 | 13 | 4.96 | 0.00129 |
| 316 | GO:0051168 | nuclear export | 146 | 18 | 8.14 | 0.00132 |
| 317 | GO:0071322 | cellular response to carbohydrate stimul... | 220 | 24 | 12.27 | 0.00135 |

| | | | | | | |
|---|---|---|---|---|---|---|
| 318 | GO:0009642 | response to light intensity | 476 | 43 | 26.55 | 0.00136 |
| 319 | GO:0032501 | multicellular organismal process | 6471 | 411 | 360.93 | 0.00136 |
| 320 | GO:0006220 | pyrimidine nucleotide metabolic process | 259 | 27 | 14.45 | 0.00138 |
| 321 | GO:0006221 | pyrimidine nucleotide biosynthetic proce... | 259 | 27 | 14.45 | 0.00138 |
| 322 | GO:0042732 | D-xylose metabolic process | 69 | 11 | 3.85 | 0.00145 |
| 323 | GO:0016458 | gene silencing | 608 | 52 | 33.91 | 0.00155 |
| 324 | GO:0009553 | embryo sac development | 396 | 37 | 22.09 | 0.00158 |
| 325 | GO:0006364 | rRNA processing | 412 | 38 | 22.98 | 0.00174 |
| 326 | GO:0022603 | regulation of anatomical structure morph... | 251 | 26 | 14 | 0.00184 |
| 327 | GO:0007030 | Golgi organization | 386 | 36 | 21.53 | 0.00187 |
| 328 | GO:0000956 | nuclear-transcribed mRNA catabolic proce... | 200 | 22 | 11.16 | 0.00188 |
| 329 | GO:0006402 | mRNA catabolic process | 200 | 22 | 11.16 | 0.00188 |
| 330 | GO:0010043 | response to zinc ion | 104 | 14 | 5.8 | 0.00191 |
| 331 | GO:0009644 | response to high light intensity | 359 | 34 | 20.02 | 0.00192 |
| 332 | GO:0016072 | rRNA metabolic process | 415 | 38 | 23.15 | 0.00197 |
| 333 | GO:0016108 | tetraterpenoid metabolic process | 176 | 20 | 9.82 | 0.00201 |
| 334 | GO:0016116 | carotenoid metabolic process | 176 | 20 | 9.82 | 0.00201 |
| 335 | GO:0006261 | DNA-dependent DNA replication | 402 | 37 | 22.42 | 0.00205 |
| 336 | GO:0006631 | fatty acid metabolic process | 719 | 59 | 40.1 | 0.00206 |
| 337 | GO:0033044 | regulation of chromosome organization | 189 | 21 | 10.54 | 0.00208 |
| 338 | GO:0048519 | negative regulation of biological proces... | 2256 | 157 | 125.83 | 0.00222 |
| 339 | GO:0051716 | cellular response to stimulus | 4609 | 299 | 257.07 | 0.00222 |
| 340 | GO:0007010 | cytoskeleton organization | 841 | 67 | 46.91 | 0.00223 |
| 341 | GO:0006084 | acetyl-CoA metabolic process | 141 | 17 | 7.86 | 0.00226 |
| 342 | GO:0031365 | N-terminal protein amino acid modificati... | 255 | 26 | 14.22 | 0.00229 |
| 343 | GO:0000723 | telomere maintenance | 73 | 11 | 4.07 | 0.0023 |
| 344 | GO:0032200 | telomere organization | 73 | 11 | 4.07 | 0.0023 |
| 345 | GO:0071166 | ribonucleoprotein complex localization | 130 | 16 | 7.25 | 0.00243 |
| 346 | GO:0071426 | ribonucleoprotein complex export from nu... | 130 | 16 | 7.25 | 0.00243 |

| | | | | | | |
|---|---|---|---|---|---|---|
| 347 | GO:0009746 | response to hexose | 310 | 30 | 17.29 | 0.00248 |
| 348 | GO:0044767 | single-organism developmental process | 6852 | 430 | 382.18 | 0.00249 |
| 349 | GO:0072528 | pyrimidine-containing compound biosynthe... | 270 | 27 | 15.06 | 0.00251 |
| 350 | GO:0006399 | tRNA metabolic process | 167 | 19 | 9.31 | 0.00253 |
| 351 | GO:0006406 | mRNA export from nucleus | 119 | 15 | 6.64 | 0.00259 |
| 352 | GO:0051028 | mRNA transport | 119 | 15 | 6.64 | 0.00259 |
| 353 | GO:0071427 | mRNA-containing ribonucleoprotein comple... | 119 | 15 | 6.64 | 0.00259 |
| 354 | GO:0006089 | lactate metabolic process | 155 | 18 | 8.65 | 0.00259 |
| 355 | GO:0019243 | methylglyoxal catabolic process to D-lac... | 155 | 18 | 8.65 | 0.00259 |
| 356 | GO:0061727 | methylglyoxal catabolic process to lacta... | 155 | 18 | 8.65 | 0.00259 |
| 357 | GO:0071702 | organic substance transport | 2952 | 199 | 164.65 | 0.0026 |
| 358 | GO:0006637 | acyl-CoA metabolic process | 143 | 17 | 7.98 | 0.00262 |
| 359 | GO:0035383 | thioester metabolic process | 143 | 17 | 7.98 | 0.00262 |
| 360 | GO:0048856 | anatomical structure development | 6862 | 430 | 382.74 | 0.00276 |
| 361 | GO:0034976 | response to endoplasmic reticulum stress | 684 | 56 | 38.15 | 0.00276 |
| 362 | GO:0019953 | sexual reproduction | 480 | 42 | 26.77 | 0.00277 |
| 363 | GO:0009438 | methylglyoxal metabolic process | 156 | 18 | 8.7 | 0.00278 |
| 364 | GO:0042182 | ketone catabolic process | 156 | 18 | 8.7 | 0.00278 |
| 365 | GO:0051596 | methylglyoxal catabolic process | 156 | 18 | 8.7 | 0.00278 |
| 366 | GO:0006403 | RNA localization | 132 | 16 | 7.36 | 0.00284 |
| 367 | GO:0006405 | RNA export from nucleus | 132 | 16 | 7.36 | 0.00284 |
| 368 | GO:0050657 | nucleic acid transport | 132 | 16 | 7.36 | 0.00284 |
| 369 | GO:0050658 | RNA transport | 132 | 16 | 7.36 | 0.00284 |
| 370 | GO:0051236 | establishment of RNA localization | 132 | 16 | 7.36 | 0.00284 |
| 371 | GO:0016053 | organic acid biosynthetic process | 1803 | 128 | 100.57 | 0.00285 |
| 372 | GO:0046394 | carboxylic acid biosynthetic process | 1803 | 128 | 100.57 | 0.00285 |
| 373 | GO:0031327 | negative regulation of cellular biosynth... | 805 | 64 | 44.9 | 0.00289 |
| 374 | GO:0051172 | negative regulation of nitrogen compound... | 805 | 64 | 44.9 | 0.00289 |
| 375 | GO:0046185 | aldehyde catabolic process | 157 | 18 | 8.76 | 0.00298 |

| | | | | | | |
|---|---|---|---|---|---|---|
| 376 | GO:0009062 | fatty acid catabolic process | 383 | 35 | 21.36 | 0.00299 |
| 377 | GO:0000398 | mRNA splicing, via spliceosome | 146 | 17 | 8.14 | 0.00326 |
| 378 | GO:0051052 | regulation of DNA metabolic process | 316 | 30 | 17.63 | 0.00329 |
| 379 | GO:0042762 | regulation of sulfur metabolic process | 36 | 7 | 2.01 | 0.00332 |
| 380 | GO:0001678 | cellular glucose homeostasis | 12 | 4 | 0.67 | 0.00333 |
| 381 | GO:0071333 | cellular response to glucose stimulus | 12 | 4 | 0.67 | 0.00333 |
| 382 | GO:0000375 | RNA splicing, via transesterification re... | 159 | 18 | 8.87 | 0.00341 |
| 383 | GO:0000377 | RNA splicing, via transesterification re... | 159 | 18 | 8.87 | 0.00341 |
| 384 | GO:0034284 | response to monosaccharide | 317 | 30 | 17.68 | 0.00345 |
| 385 | GO:0015980 | energy derivation by oxidation of organi... | 172 | 19 | 9.59 | 0.00352 |
| 386 | GO:0045333 | cellular respiration | 172 | 19 | 9.59 | 0.00352 |
| 387 | GO:0061024 | membrane organization | 318 | 30 | 17.74 | 0.00361 |
| 388 | GO:0019750 | chloroplast localization | 160 | 18 | 8.92 | 0.00365 |
| 389 | GO:0051644 | plastid localization | 160 | 18 | 8.92 | 0.00365 |
| 390 | GO:0009890 | negative regulation of biosynthetic proc... | 814 | 64 | 45.4 | 0.00372 |
| 391 | GO:0010558 | negative regulation of macromolecule bio... | 784 | 62 | 43.73 | 0.00374 |
| 392 | GO:2000113 | negative regulation of cellular macromol... | 784 | 62 | 43.73 | 0.00374 |
| 393 | GO:0048229 | gametophyte development | 875 | 68 | 48.8 | 0.00374 |
| 394 | GO:0032504 | multicellular organism reproduction | 694 | 56 | 38.71 | 0.00375 |
| 395 | GO:0006623 | protein targeting to vacuole | 225 | 23 | 12.55 | 0.00384 |
| 396 | GO:0072665 | protein localization to vacuole | 225 | 23 | 12.55 | 0.00384 |
| 397 | GO:0072666 | establishment of protein localization to... | 225 | 23 | 12.55 | 0.00384 |
| 398 | GO:0005984 | disaccharide metabolic process | 292 | 28 | 16.29 | 0.00384 |
| 399 | GO:0006352 | DNA-templated transcription, initiation | 28 | 6 | 1.56 | 0.00389 |
| 400 | GO:1900376 | regulation of secondary metabolite biosy... | 28 | 6 | 1.56 | 0.00389 |
| 401 | GO:0072527 | pyrimidine-containing compound metabolic... | 279 | 27 | 15.56 | 0.00395 |
| 402 | GO:0000280 | nuclear division | 461 | 40 | 25.71 | 0.00397 |
| 403 | GO:0009617 | response to bacterium | 1124 | 84 | 62.69 | 0.00399 |
| 404 | GO:0016125 | sterol metabolic process | 293 | 28 | 16.34 | 0.00403 |

| | | | | | | |
|---|---|---|---|---|---|---|
| 405 | GO:0044703 | multi-organism reproductive process | 491 | 42 | 27.39 | 0.00414 |
| 406 | GO:0009059 | macromolecule biosynthetic process | 5990 | 377 | 334.1 | 0.00414 |
| 407 | GO:0009909 | regulation of flower development | 653 | 53 | 36.42 | 0.0042 |
| 408 | GO:0090304 | nucleic acid metabolic process | 5718 | 361 | 318.93 | 0.00423 |
| 409 | GO:0009069 | serine family amino acid metabolic proce... | 463 | 40 | 25.82 | 0.00427 |
| 410 | GO:0031324 | negative regulation of cellular metaboli... | 926 | 71 | 51.65 | 0.00428 |
| 411 | GO:0016126 | sterol biosynthetic process | 281 | 27 | 15.67 | 0.00435 |
| 412 | GO:0006487 | protein N-linked glycosylation | 163 | 18 | 9.09 | 0.00445 |
| 413 | GO:0006260 | DNA replication | 479 | 41 | 26.72 | 0.00452 |
| 414 | GO:0032876 | negative regulation of DNA endoreduplica... | 13 | 4 | 0.73 | 0.00459 |
| 415 | GO:0070370 | cellular heat acclimation | 13 | 4 | 0.73 | 0.00459 |
| 416 | GO:0048878 | chemical homeostasis | 540 | 45 | 30.12 | 0.00496 |
| 417 | GO:0044802 | single-organism membrane organization | 313 | 29 | 17.46 | 0.00535 |
| 418 | GO:0006606 | protein import into nucleus | 245 | 24 | 13.67 | 0.00545 |
| 419 | GO:0044744 | protein targeting to nucleus | 245 | 24 | 13.67 | 0.00545 |
| 420 | GO:1902593 | single-organism nuclear import | 245 | 24 | 13.67 | 0.00545 |
| 421 | GO:0072329 | monocarboxylic acid catabolic process | 398 | 35 | 22.2 | 0.00548 |
| 422 | GO:0009668 | plastid membrane organization | 286 | 27 | 15.95 | 0.0055 |
| 423 | GO:0010027 | thylakoid membrane organization | 286 | 27 | 15.95 | 0.0055 |
| 424 | GO:0032268 | regulation of cellular protein metabolic... | 693 | 55 | 38.65 | 0.00565 |
| 425 | GO:0015976 | carbon utilization | 14 | 4 | 0.78 | 0.00615 |
| 426 | GO:0006498 | N-terminal protein lipidation | 248 | 24 | 13.83 | 0.00632 |
| 427 | GO:0006499 | N-terminal protein myristoylation | 248 | 24 | 13.83 | 0.00632 |
| 428 | GO:0009749 | response to glucose | 156 | 17 | 8.7 | 0.00638 |
| 429 | GO:0009756 | carbohydrate mediated signaling | 208 | 21 | 11.6 | 0.00639 |
| 430 | GO:0010182 | sugar mediated signaling pathway | 208 | 21 | 11.6 | 0.00639 |
| 431 | GO:1901362 | organic cyclic compound biosynthetic pro... | 5416 | 341 | 302.09 | 0.00648 |
| 432 | GO:0031047 | gene silencing by RNA | 476 | 40 | 26.55 | 0.00675 |
| 433 | GO:0019344 | cysteine biosynthetic process | 375 | 33 | 20.92 | 0.00677 |

| | | | | | | |
|---|---|---|---|---|---|---|
| 434 | GO:0034645 | cellular macromolecule biosynthetic proc... | 5815 | 364 | 324.34 | 0.0068 |
| 435 | GO:0051246 | regulation of protein metabolic process | 700 | 55 | 39.04 | 0.00687 |
| 436 | GO:0071705 | nitrogen compound transport | 3004 | 198 | 167.55 | 0.00689 |
| 437 | GO:0032502 | developmental process | 7007 | 433 | 390.83 | 0.00695 |
| 438 | GO:0040029 | regulation of gene expression, epigeneti... | 581 | 47 | 32.41 | 0.00712 |
| 439 | GO:0048518 | positive regulation of biological proces... | 1709 | 119 | 95.32 | 0.00717 |
| 440 | GO:0009902 | chloroplast relocation | 158 | 17 | 8.81 | 0.00723 |
| 441 | GO:0051667 | establishment of plastid localization | 158 | 17 | 8.81 | 0.00723 |
| 442 | GO:1902589 | single-organism organelle organization | 778 | 60 | 43.39 | 0.00724 |
| 443 | GO:0018377 | protein myristoylation | 251 | 24 | 14 | 0.0073 |
| 444 | GO:0034504 | protein localization to nucleus | 251 | 24 | 14 | 0.0073 |
| 445 | GO:0006534 | cysteine metabolic process | 377 | 33 | 21.03 | 0.00731 |
| 446 | GO:0022414 | reproductive process | 4067 | 261 | 226.84 | 0.00739 |
| 447 | GO:0044242 | cellular lipid catabolic process | 435 | 37 | 24.26 | 0.00741 |
| 448 | GO:0048831 | regulation of shoot system development | 719 | 56 | 40.1 | 0.00763 |
| 449 | GO:0051170 | nuclear import | 252 | 24 | 14.06 | 0.00766 |
| 450 | GO:0000003 | reproduction | 4072 | 261 | 227.12 | 0.00783 |
| 451 | GO:0008156 | negative regulation of DNA replication | 15 | 4 | 0.84 | 0.00802 |
| 452 | GO:0019217 | regulation of fatty acid metabolic proce... | 15 | 4 | 0.84 | 0.00802 |
| 453 | GO:0042593 | glucose homeostasis | 15 | 4 | 0.84 | 0.00802 |
| 454 | GO:2000104 | negative regulation of DNA-dependent DNA... | 15 | 4 | 0.84 | 0.00802 |
| 455 | GO:1902580 | single-organism cellular localization | 467 | 39 | 26.05 | 0.00817 |
| 456 | GO:1902582 | single-organism intracellular transport | 467 | 39 | 26.05 | 0.00817 |
| 457 | GO:0009070 | serine family amino acid biosynthetic pr... | 381 | 33 | 21.25 | 0.0085 |
| 458 | GO:0050826 | response to freezing | 187 | 19 | 10.43 | 0.00859 |
| 459 | GO:0010212 | response to ionizing radiation | 123 | 14 | 6.86 | 0.00873 |
| 460 | GO:1902578 | single-organism localization | 2211 | 149 | 123.32 | 0.00886 |
| 461 | GO:0048609 | multicellular organismal reproductive pr... | 664 | 52 | 37.04 | 0.00892 |
| 462 | GO:0007034 | vacuolar transport | 242 | 23 | 13.5 | 0.00911 |

| | | | | | | |
|---|---|---|---|---|---|---|
| 463 | GO:0017038 | protein import | 441 | 37 | 24.6 | 0.00913 |
| 464 | GO:0051656 | establishment of organelle localization | 162 | 17 | 9.04 | 0.0092 |
| 465 | GO:0048522 | positive regulation of cellular process | 1290 | 92 | 71.95 | 0.00933 |
| 466 | GO:0051260 | protein homooligomerization | 24 | 5 | 1.34 | 0.00937 |
| 467 | GO:0044765 | single-organism transport | 2200 | 148 | 122.71 | 0.00961 |

#### Supplementary Table 7

RT-qPCR primers used to validate RNAseq data.

| *Gene of Interest* | *Primer Orientation* | *Sequence* |
|---|---|---|
| *Housekeeping Gene UBC* | *Forward* | *GATCCACCCACCTCGTGTAG* |
| *Housekeeping Gene UBC* | *Reverse* | *CTGGAGGGAAGTGAATGGTAAC* |
| *MYB28 (Bo2g161590)* | *Forward* | *TCACGAACATGGAGAAGGTG* |
| *MYB28 (Bo2g161590)* | *Reverse* | *ACCACTTGTTGCCACGAGCA* |
| *MAM3 (Bo7g098000)* | *Forward* | *ACCGCAGAAGCTTGAGATTG* |
| *MAM3 (Bo7g098000)* | *Reverse* | *CTTCATCCACCTCATTTCCC* |
| *CYP79F1 (Bo5g021810)* | *Forward* | *GACGAGAGGGTGGAGTTATG* |
| *CYP79F1 (Bo5g021810)* | *Reverse* | *AATTCAACGCATTGAGCTTT* |
| *APS3 (Bo1g057690)* | *Forward* | *CAATTCGCTTCGTCTCGACG* |
| *APS3 (Bo1g057690)* | *Reverse* | *CTCAATGTCGCTGAGAATAGCG* |

#### Supplementary Figure 3

Alignment of MYB28 C7 of the high glucoraphanin broccoli line HG Inbred, *Brassica villosa* and Ironman using the Clustal Omega Multiple Sequence Aligner (Sievers, Wilm et al. 2011).

```
CLUSTAL O(1.2.4) multiple sequence alignment

Ironman         ATGTCAAGAAAACCGTGTTGTGTCGGAGAAGGGCTGAAGAAAGGGGCATGGACCACCGAA 60
HGInbred        ATGTCAAGAAAACCGTGTTGTGTCGGAGAAGGGCTGAAGAAAGGGGCATGGACCACCGAA 60
BVillosa        ATGTCAAGAAAACCGTGTTGTGTCGGAGAAGGGCTGAAGAAAGGGGCATGGACCACCGAA 60
                ************************************************************

Ironman         GAAGATAAGAAACTCATCTCTTACATCCACGAACATGGAGAAAGAGGCTGGCGCGACATT 120
HGInbred        GAAGATAAGAAACTCATCTCTTACATCCACGAACATGGAGAAGGAGGCTGGCGCGACATT 120
BVillosa        GAAGATAAGAAACTCATCTCTTACATCCACGAACATGGAGAAGGAGGCTGGCGCGACATT 120
                ****************************************** *****************

Ironman         CCCCAAAAAGCTGGTTAAATTTTTTTTTTGTAACACAGCTGGTTAAATATCTATGAAATT 180
HGInbred        CCCCAAAAAGCTGGTTAAATTTTTTTTTTGTAACACAGCTGGTTAAATATCTATGAAATT 180
BVillosa        CCCCAAAAAGCTGGTTAAATTTTTTTTTTGTAACACAGCTGGTTAAATATCTATGAAATT 180
                ************************************************************

Ironman         ATACACATGCTTATTTAGTAATCGTACTTGTATGAAATTCTAATTAATTATGATTATGGT 240
HGInbred        ATACACATGCTTATTTAGTAATCGTACTTGTATGAAATTCTAATTAATTATGATTATGGT 240
BVillosa        ATACACATGCTTATTTAGTAATCGTACTTGTATGAAATTCTAATTAATTATGATTATGGT 240
                ************************************************************

Ironman         ATGTAATTAGGGTTAAAAAGGTGTGGAAAGAGTTGTAGACTGCGTTGGACTTACTACCTA 300
HGInbred        ATGTAATTAGGGTTAAAAAGGTGTGGAAAGAGTTGTAGACTGCGTTGGACTTACTACCTA 300
BVillosa        ATGTAATTAGGGTTAAAAAGGTGTGGAAAGAGTTGTAGACTGCGTTGGACTAACTACCTA 300
                *************************************************** ********

Ironman         AAACCTGATGTCAAAAGAGGCGAGTTTAGTTCAGAGGAGGAACAGATTATTATTATGCTT 360
HGInbred        AAACCTGATGTCAAAAGAGGCGAGTTTAGTTCAGAGGAGGAACAGATTATTATTATGCTT 360
BVillosa        AAACCTGATGTCAAAAGAGGCGAGTTTAGTTCAGAGGAGGAACAGATTATTATTTTGCTT 360
                ****************************************************** *****

Ironman         CATGCTTCTCGTGGCAACAAGTACGTTTATATGCTTAAATGTGTGTGTATGCATGGATCC 420
HGInbred        CATGCTTCTCGTGGCAACAAGTACGTTTATATGCTTAAATGTGTGTGTATGCATGGATCC 420
BVillosa        CATGCTTCTCGTGGCAACAAGTACGTTTATATGCTTAAATGTGTGTGTATGCATGGATCC 420
                ************************************************************

Ironman         TCCAATGAATAATGAAATGCATGAAAAAGTTCCATATATTATTTTAAAGATACGATCATT 480
HGInbred        TCCAATGAATAATGAAATGCATGAAAAAGTTCCATATATTATTTTAAAGATACGATCATT 480
BVillosa        TCCAATTAAAAATGAAATGCATGAAAAAGTTCCATATATTATTTTAAAGATACGATCATT 480
                ****** ** **************************************************

Ironman         ATTAGTTAATATAGTCCCTTCCGCGAAAAGCTCTACAAAGGAATCATATTAATACCGTCT 540
HGInbred        ATTAGTTAATATAGTCCCTTCCGCGAAAAGCTCTACAAAGGAATCATATTAATACCGTCT 540
BVillosa        ATTAGTTAATATAGTCCCTTCCGCGAAAAGCTCTACAAAGGAATCATATTAATACCGTCT 540
                ************************************************************

Ironman         TTGATATCTCAATCTCTCTTCTTGCCTTTAGGTGGTCGGACATAGCGAGACATTTACCTA 600
HGInbred        TTGATATCTCAATCTCTCTTCTTGCCTTTAGGTGGTCGGACATAGCGAGACATTTACCTA 600
BVillosa        TTGATATCTCAATCTCTCTTCTTGCCTTTAGGTGGTCGGACATAGCGAGACATTTACCTA 600
                ************************************************************

Ironman         GAAGAACAGACAATGAGGTCAAGAATTACTGGAACACTCATCTTAAAAAGCGTTTGATCG 660
HGInbred        GAAGAACAGACAATGAGGTCAAGAATTACTGGAACACTCATCTTAAAAAGCGTTTGATCG 660
BVillosa        GAAGAACAGACAATGAGGTCAAGAATTACTGGAACACTCATCTTAAAAAGCGTTTGATCG 660
                ************************************************************

Ironman         AACAGGGTATTGATCCCGTGACTCACAAGCCACTGGCTTCTAATTCCAACCCTACCGTCA 720
HGInbred        AACAGGGTATTGATCCCGTGACTCACAAGCCACTGGCTTCTAATTCCAACCCTACCGTCA 720
BVillosa        AACAGGGTATTGATCCCGTGACTCACAAGCCACTGGCTTCTAATTCCAACCCTACCGTCA 720
                ************************************************************
```

```
Ironman         ACACGCCTCCAGAGAATTTGCATTCCCTTGCTGCGCCTAGTTCCGACAAGCAATACTCCC 780
HGInbred        ACACGCCTCCAGAGAATTTGCATTCCCTTGCTGCGCCTAGTTCCGACAAGCAACACTCCC 780
BVillosa        ACACGCCTCCAGAGAATTTGCATTCCCTTGCTGCGCCTAGTTCCGACAAGGAATACTCCC 780
                ************************************************** ** ******

Ironman         GGTCGAGCTCATTTCCTTCCCTGTCTCGTCTTACCAACAAAGATGGGACACCAGTTCAAG 840
HGInbred        GGTCGAGCTCATTGCCTTCCCTGTCTCGTCTTACCAACAAAGATGGGACACCAGTTCAAG 840
BVillosa        GGTCGAGCTCATTGCCTTCCCTGTCTCGTCTTACCAACAAAGATGGGACACCAGTTCAAG 840
                ************* **********************************************

Ironman         GCGGTTCCTTGAGTCACAAGAAACGTTTTAAGAGGTCGAGTTCTACATCAAGGCTTTTGA 900
HGInbred        GCGGTTCCTTGAGTCACAAGAAACGTTTTAAGAGGTCGAGTTCTACATCAAGGCTTTTGA 900
BVillosa        GCGGTTCCTTGAGTCACAAGAAACGTTTTAAGAGGTCGAGTTCTACATCAAGGCTTTTGA 900
                ************************************************************

Ironman         ACAAAGTTGCGGCTAAGGTCACTTCTGTAAAAGAAATATTGTCGGCTTCCATGGAAGGTA 960
HGInbred        ACAAAGTTGCGGCTAAGGTCACTTCTGTAAAAGAAATATTGTCGGCTTCCATGGAAGGTA 960
BVillosa        ACAAAGTTGCGGCTAAGGTCACTTCTGTAAAAGAAATATTGTCGGCTTCCATGGAAGGTA 960
                ************************************************************

Ironman         GCTTGAGCGCTACTACATTACCATATGCAAGCTATTCTGATGGCTTCTCTGAGCAGATTC 1020
HGInbred        GCTTGAGCGCTACTACATTACCATATGCAAGCTATTCTGATGGCTTCTCTGAGCAGATTC 1020
BVillosa        GCTTGAGCGCTACTACATTACCATATGCAAGCTATTCTGATGGCTTCTCTGAGCAGATTC 1020
                ************************************************************

Ironman         GCAATGGAGAGGGTAGTTCCAACGCGTTCCTGACAAATACTCTCGCCGAGTTCGATCCCT 1080
HGInbred        GCAATGGAGAGGGTAGTTCCAACGCGTTCCTGACAAATACTCTCGCCGAGTTCGATCCCT 1080
BVillosa        GCAATGGAGAGGATAGTTCCAACGCGTTCCTGACAAATACTCTCGCCGAGTTCGATCCCT 1080
                ************ ***********************************************

Ironman         TCTCCCAATCACCGTTGTACAGTGAGCATGAGATCAACGCTACTTCTGATCTCGGTATGG 1140
HGInbred        TCTCCCAATCACCGTTGTACAGTGAGCATGAGATCAACGCTACTTCTGATCTCGGTATGG 1140
BVillosa        TCTCCCAATCACCGTTGTACAGTGAGCATGAGATCAACGCTACTTCTGATCTCGGAATGG 1140
                ******************************************************* ****

Ironman         ATTACGATTTCTCACATTTTCTTGAAAAGCTTGGGAGAGATGACCACAACGAGGAGAACG 1200
HGInbred        ATTACGATTTCTCACATTTTCTTGAAAAGCTTGGGAGAGATGACCACAACGAGGAGAACG 1200
BVillosa        ATTACGATTTCTCACATTTTCTTGAAAAGCTTTGGAGAGATGACCACAACGAGGAGAACG 1200
                ******************************** ***************************

Ironman         ATATGAATGTCGAGTATGGTCATGATGATCTTCTTATGTCCGATGTGTCTCAAGAAGTCT 1260
HGInbred        ATATGAATGTCGAGTATGGTCATGATGATCTTCTTATGTCCGATGTGTCTCAAGAAGTCT 1260
BVillosa        ATATGAATGTCGAGTATGGTCATGATGATCTTCTTATGTCCGATGTGTCTCAAGAAGTCT 1260
                ************************************************************

Ironman         CATCAACAAGCGTTGATGATCGAGACAATATATTTGAAAATTTTGAGGGTTGGTCAAATT 1320
HGInbred        CATCAACAAGCGTTGATGATCGACACAATATATTTGAAAATTTTGAGGGTTGGTCAAATT 1320
BVillosa        CATCAACAAGCGTTGATGATCGAGACAATATATTTGAAAATTTTGAGGGTTGGTCAAATT 1320
                *********************** ************************************

Ironman         ATCTTCTTGACCATGCGGATTTCGTATATGACACGGAGTCTGATACCCTCATATGA     1376
HGInbred        ATCTTCTTGACCATGCGGATTTCGTATATGACACGGAGTCTGATACCCTCATATGA     1376
BVillosa        ATCTTCTTGGCCATGCGGATTTCGTATATGACACGGAGTCTGATACCCTCATATGA     1376
                ********* **********************************************
```

#### Supplementary Figure 4

Alignment of MYB28 C9 of the high glucoraphanin broccoli line HG Inbred, *Brassica villosa* and Ironman using the Clustal Omega Multiple Sequence Aligner (Sievers, Wilm et al. 2011).

```
CLUSTAL O(1.2.4) multiple sequence alignment

BVillosa        ATGTCAAGAAAGCCATGTTGTGTCGGAGAAGGGCTGAAGAAAGGGGCATGGACCACCGAG 60
Ironman         ATGTCAAGAAAGCCATGTTGTGTCGGAGAAGGGCTGAAGAAAGGGGCATGGACCACCGAG 60
HGInbred        ATGTCAAGAAAGCCATGTTGTGTCGGAGAAGGGCTGAAGAAAGGGGCATGGACCACCGAG 60
                ************************************************************

BVillosa        GAAGATAAGAAACTCATCTCTTACATCCATGAACATGGAGAAGGAGGCTGGCGCGACATT 120
Ironman         GAAGATAAGAAACTCATCTCTTACATCCATGAACATGGAGAAGGAGGCTGGCGCGACATT 120
HGInbred        GAAGATAAGAAACTCATCTCTTACATCCATGAACATGGAGAAGGAGGCTGGCGCGACATT 120
                ************************************************************

BVillosa        CCTCAAAAAGCTGGTTAATATCTATTATATATTTTTTGGTAAATTTTTAAAACATATATG 180
Ironman         CCTCAAAAAGCTGGTTAATATCTATTATATATATTTTGGTAAATTTTTAAAACATATATG 180
HGInbred        CCTCAAAAAGCTGGTTAATATCTATTATATATATTTTGGTAAATTTTTAAAACATATATG 180
                ******************************** ***************************

BVillosa        TTTGTTTGGTATTTGATGTATGAAAGTTTTATGTTGAATATGGTGTTTTACTAGGATTGA 240
Ironman         TTTGTTTGGTATTTGATGTATGAAAGTTTTATATTGAATGTGGTGTTTTACTAGGATTGA 240
HGInbred        TTTGTTTGGTATTTGATGTATGAAAGTTTTATATTGAATGTGGTGTTTTACTAGGATTGA 240
                ******************************** ****** ********************

BVillosa        AAAGGTGTGGAAAGAGTTGTAGACTGCGATGGACTAACTACCTAAAACCTGAAATCAAAA 300
Ironman         AAAGGTGTGGAAAGAGTTGTAGACTGCGATGGACTAACTACCTAAAACCTGAAATCAAAA 300
HGInbred        AAAGGTGTGGAAAGAGTTGTAGACTGCGATGGACTAACTACCTAAAACCTGAAATCAAAA 300
                ************************************************************

BVillosa        GAGGCGAGTTTAGTTCAGAGGAGGAACAGATTATCATCATGCTTCATGCTTCTCGTGGAA 360
Ironman         GAGGCGAGTTTAGTTCAGAGGAGGAACAGATTATCATCATGCTTCATGCTTCTCGTGGAA 360
HGInbred        GAGGCGAGTTTAGTTCAGAGGAGGAACAGATTATCATCATGCTTCATGCTTCTCGTGGAA 360
                ************************************************************

BVillosa        ACAAGTAAGTATATATGTGTGTATGGGTCCTCCAATGATTATCATTCTGATTTTGTTTCT 420
Ironman         ACAAGTAAGTATATATGTGTGTATGGGTCCTCCAATGATTATCATTCTGATTTTGTTTCT 420
HGInbred        ACAAGTAAGTATATATGTGTGTATGGGTCCTCCAATGATTATCATTCTGATTTTGTTTCT 420
                ************************************************************

BVillosa        TTTCCATGTATTTGACAATTCTCTGATGCAAAATATGTGTTGGATTAGCAATTTTTGACT 480
Ironman         TTTCCATGTATTTGACAATTCTCTGATGCAAAATATGTGTTGGATTAGCAATTTTTGACT 480
HGInbred        TTTCCATGTATTTGACAATTCTCTGATGCAAAATATGTGTTGGATTAGCAATTTTTGACT 480
                ************************************************************

BVillosa        AAAAATTGCAATACCACACATCATTTTAGTTTCAAAATAAATTTTATTAACTTTGGTCTT 540
Ironman         AAAAATTGCAATAGCACACATCATTTTAGTTTCAAAATAAATTTTATTAACTTTGGTCTT 540
HGInbred        AAAAATTGCAATAGCACACATCATTTTAGTTTCAAAATAAATTTTATTAACTTTGGTCTT 540
                ************* **********************************************

BVillosa        CCATCTTATCTCTTTCACTATTCTGGTCTTTAGGTGGTCAGTCATAGCGAGACATTTACC 600
Ironman         CCATCTTATCTCTTTCACTATTCTGGTCTTTAGGTGGTCGGTCATAGCGAGACATTTACC 600
HGInbred        CCATCTTATCTCTTTCACTATTCTGGTCTTTAGGTGGTCGGTCATAGCGAGACATTTACC 600
                *************************************** ********************

BVillosa        TAGAAGAACAGACAATGAGATCAAGAACTACTGGAACACACATCTCAAGAAACGTTTGAT 660
Ironman         TAGAAGAACAGACAATGAGATCAAGAACTACTGGAACACACATCTCAAGAAACGTTTGAT 660
HGInbred        TAGAAGAACAGACAATGAGATCAAGAACTACTGGAACACACATCTCAAGAAACGTTTGAT 660
                ************************************************************

BVillosa        CGAACAGGGTACTGATCCCGTGACTCACAAGCCACTAGCTTCTAATACAAACCCTACTGT 720
Ironman         CGAACAGGGTACTGATCCCGTGACTCACAAGCCACTAGCTTCTAATACAAACCCTACTGT 720
HGInbred        CGAACAGGGTACTGATCCCGTGACTCACAAGCCACTAGCTTCTAATACAAACCCTACTGT 720
                ************************************************************
```

```
BVillosa        ACCTGAGAATTTGCATTCCCTAGATGCATCTAGTAATTCCGACAAGCAATACTCCCGGTC 780
Ironman         ACCTGAGAATTTGCATTCCCTAGATGCATCTAG---TTCCGACAAGCAATACTCCCGGTC 777
HGInbred        ACCTGAGAATTTGCATTCCCTAGATGCATCTAG---TTCCGACAAGCAATACTCCCGGTC 777
                *********************************   ************************

BVillosa        AACCTCAATGCCTTCCATGTCTTGTACTCCTTCCTCCGGTTTCAACACGGTTTTCGAGAA 840
Ironman         AAGCTCAATGCCTTCCATGTCTTGTACTCCTTCCTCCGGTTTCAACACGGTTTTCGAGAA 837
HGInbred        AAGCTCAATGCCTTCCATGTCTTGTACTCCTTCCTCCGGTTTCAACACGGTTTTCGAGAA 837
                ** *********************************************************

BVillosa        TACCAGCAAAGATGGGACACCAGTTCGTGAGGACGATTCCTTGAGTCGCAAGAAACGTTT 900
Ironman         TACCAGCAAAGATGGGACACCAGTTCGTGAGGACGATTCCTTGAGTCGCAAGAAACGTTT 897
HGInbred        TACCAGCAAAGATGGGACACCAGTTCGTGAGGACGATTCCTTGAGTCGCAAGAAACGTTT 897
                ************************************************************

BVillosa        TAAGAAATCAAGTTCTACATCAAGGCTTTTGAACAAAGTTGCGGCTAAGGCCACTTCCAT 960
Ironman         GAAGAAATCAAGTTCTACATCAAGGCTTTTGAACAAAGTTGCGGCTAAGGCCACTTCCAT 957
HGInbred        GAAGAAATCAAGTTCTACATCAAGGCTTTTGAACAAAGTTGCGGCTAAGGCCACTTCCAT 957
                 ***********************************************************

BVillosa        GAAAGAAGCTTTGTCTGCTTCCATGGAAGGTAGCTTGAATGCTAATACAAGCTTTTCCAA 1020
Ironman         GAAAAAAGCTTTGTCTGCTTCCATGGAAGGTAGCTTGAATGCTAATATAAGCTTTTCCAA 1017
HGInbred        GAAAGAAGCTTTGTCTGCTTCCATGGAAGGTAGCTTGAATGCTAATACAAGCTTTTCCAA 1017
                **** ****************************************** ************

BVillosa        TGGCTACTCTGAGCAGATTCTCAATGAAGATGATAGTTCTAATGCATCCCTCATAAACAC 1080
Ironman         TGGCTACTCTGAGCAGATTCTCAATGAAGATGATAGTTCTAATGCATCCCTCATAAACAC 1077
HGInbred        TGGCTACTCTGAGCAGATTCTCAATGAAGATGATAGTTCTAATGCATCCCTCATAAACAC 1077
                ************************************************************

BVillosa        TCTCGCCGAGTTCGATCCCTTCCTCCAAACAACGTTTTACCCTGAGAATGAGATGAATAC 1140
Ironman         TCTCGCCGAGTTCGATCCCTTCCTCCAAACAACGTTTTACCCTGAGAATGAGATGAATAC 1137
HGInbred        TCTCGCCGAGTTCGATCCCTTCCTCCAAACAACGTTTTACCCTGAGAATGAAATGAATAC 1137
                *************************************************** ********

BVillosa        TACTTCTGATCTCGGTATAGATCAGGACTACTTCTCACATTTTCTCGAAAATTTCGGCAG 1200
Ironman         TACTTCTGATCTCGGTATAGATCAGGACTACTTCTCACATTTTCTCGAAAATTTCGGCAA 1197
HGInbred        TACTTCTGATCTCGATATAGATCAGGACTACTTCTCACATTTTCTCGAAAATTTCGGCAG 1197
                ************** ******************************************** 

BVillosa        AGATGATGACCACAATGAGGAGCACTACATGAATCATAACTATGGTCATGGTCTTCTTAT 1260
Ironman         CC---------ATAATGAGGAGCACTACATGAATCATAACTATGGTCATGGTCTTCTTAT 1248
HGInbred        AGATGATGACCACAATGAGGAGCACTACATGAATCATAACTATGGTCATGATCTTCTTAT 1257
                           * ************************************* *********

BVillosa        GTCCGATGTGTCCCAAGAAGTCTCATCAACTAGCGTTGATGATCAAGACAATACTAATGA 1320
Ironman         GTCCTATGTGTCCCAAGAAGTCTCATCAACTAGCGTTGATGATCAAGACAATACTAATGA 1308
HGInbred        GTCCGATGTGTCCCAAGAAGTCTCATCAACTAGCGTTGATGATCAAGACAATACTAATGA 1317
                **** *******************************************************
```

#### Supplementary Figure 5

Alignment of MYB29 C9 of the high glucoraphanin broccoli line HG Inbred, *Brassica villosa* and the reference *Brassica oleracea* TO1000 CDS sequence taken from Ensembl Plants (Bolser, Staines et al. 2016) using the Clustal Omega Multiple Sequence Aligner (Sievers, Wilm et al. 2011).

```
CDS             CCCGTGGCAACAAGTGGTCGGTGATAGCGAGACATTTGCCAAAAAGGACAGACAACGAGG 310
HGInbred        TTGTCCGCACAAGGTGGTCGGTGATAGCGAGACATTTGCCAAAAAGGACAGACAACGAGG 1542
BVillosa        TTGTCCGCACAAGGTGGTCGGTGATAGCGAGACATTTGCCAAAAAGGACAGACAACGAGG 773
                      ***  * ***********************************************

CDS             TCAAGAACTATTGGAACACACATCTCAAAAAACGTCTGATCGATCAGGGCATCGATCCCT 370
HGInbred        TCAAGAACTATTGGAACACACATCTCAAAAAACGTCTGATCGATCAGGGCATCGATCCCT 1602
BVillosa        TCAAGAACTATTGGAACACACATCTCAAAAAACGTCTGATCGATCAGGGCATCGATCCCT 833
                ************************************************************

CDS             TGACCCACAAGCCACTTGCTTCTAACCCTAACCCGGCCACGCCCAGGACTTCTGAAGGCC 430
HGInbred        TGACCCACAAGCCACTTGCTTCTAACCCTAACCCGGCCACGCCCAGGACTTCTGAAGGCC 1662
BVillosa        TGACCCACAAGCCACTTGCTTCTAACCCTAATCCGGCCACGCCCAGGACTTCTGAAGTCC 893
                ******************************* ************************* **

CDS             AAGATGACTCAAACCCGATTAACCTGGATGAGCAATCACAATCTGGTTCGATGTCTCCAA 490
HGInbred        AAGATGACTCAAACCCGATTAACCTGGATGAGCAATCACAATCTGGTTCGATGTCTCCAA 1722
BVillosa        AAGATGACTCAAACCCGAGTAACCTGGATGAGCAATCACAATCTGGTTCGATGTCTCCAA 953
                ****************** *****************************************

CDS             AGTCTCTTCCTCATTCATCAAGCTCCTGCAATCTACAAACGATAATCAACAGTGATGAGA 550
HGInbred        AGTCTCTTCCTCATTCATCAAGCTCCTGCAATCTACAAACGATAATCAACAGTGATGAGA 1782
BVillosa        AGTCTCTTCCTCATTCATCAAGCTCCTGCAATCTACAAACGATAATCAACAGTGATGAGA 1013
                ************************************************************

CDS             CACCGAGAAACAATGGTTCCTTGAGCTCCAAGAAATGTCGTTTCAAGAAACTGAGCTCTA 610
HGInbred        CACCGAGAAACAATGGTTCCTTGAGCTCCAAGAAATGTCGTTTCAAGAAACTGAGCTCTA 1842
BVillosa        CACCGAGATACAATGGTTCCTTGAGCTCCAAGAAATGTCGTTTCAAGAAACTGAGCTCTA 1073
                ******** ***************************************************

CDS             CATCAAAACTGTTAAACAAAGTTGCAACTAAAGCGTCTTCCATTGGCAATATGCTTTCAG 670
HGInbred        CATCAAAACTGTTAAACAAAGTTGCAACTAAAGCGTCTTCCATTGGCAATATGCTTTCAG 1902
BVillosa        CATCAAAACTGTTAAACAAAGTTGCAACTAAAGCGTCTTCCATTGGCAATATGCTTTCAG 1133
                ************************************************************

CDS             CTTCCATGGAAGGATCCTTAATTCGCTCAACAACACTGTCTCCATGTCTCAATGA----- 725
HGInbred        CTTCCATGGAAGGATCCTTAATTCGCTCAACAACACTGTCTCCATGTCTCAATGATGTCT 1962
BVillosa        CTTCCATGGAAGGATCCTTAATTCGCTCAACAACGCTGTCTCCATGTCTCAATGATGTCT 1193
                ********************************** ********************

CDS             ------------------------TATGGACGAGTTTGATCCATTCTCTCACTCCTCTGA 761
HGInbred        TCTCCGAAACTAGTCAATTTCTCAGATGGACGAGTTTGATCCATTCTCTCACTCCTCTGA 2022
BVillosa        TCTCGGAAACTAGTCAATTT--CAGATGGACGAATTTGATCCATTCTCTCACTCCTCTGA 1251
                         ******** **************************

CDS             ACAACATATAACTGATCATATGAAGGAAGATATCAACATGGACTTTGATCTCAACAATTC 821
HGInbred        ACAACATATAACTGATCATATGAAGGAAGATATCAACATGGACTTTGATCTCAACAATTC 2082
BVillosa        ACAACATATAACTGATCATATGAAGGAAGATATCAACATGGACTTTGATCTCAACAATTC 1311
                ************************************************************

CDS             GGAATTCGATTTCTCGCAGTTTATCGAGCAATTTAGTAACATTGAAGCCGAAGAAGCCAA 881
HGInbred        GGAATTCGATTTCTCGCAGTTTATCGAGCAATTTAGTAACATTGAAGCCGAAGAAGCCAA 2142
BVillosa        GGAATTCGATTTCTCGCAGTTTATCGAGCAATTTAGTAACATTGAAGCCGAAGAAGCCAA 1371
                ************************************************************

CDS             TGCCATTGGAGGATATAACCAAGATCTCCTGTTGTCTGATGTCTCATCAACAAGCGTTGA 941
HGInbred        TGCCATTGGAGGATATAACCAAGATCTCCTGTTGTCTGATGTCTCATCAACAAGCGTTGA 2202
BVillosa        TGCCATTGGAGGATATAACCAAGATCTCCTGTTGTCTGATGTCTCATCAACAAGCGTTGA 1431
                ************************************************************
```

```
CDS             TGAAGACAATGTGATGCAAAACATAACCGGTTGGTCCAACTATCTTGTTGACGATTCCGA 1001
HGInbred        TGAAGACAATGTGATGCAAAACATAACCGGTTGGTCCAACTATCTTGTTGACGATTCCGA 2262
BVillosa        TGAAGACAATATGATGCAAAACATAACCGGTTGGTCCAACTATCTTGTTGACGATTCCGA 1491
                ********** *************************************************

CDS             TTTTCCTTATGACACGAACCAAGATTTCGACGACAAGAACTTCATATGA   1050
HGInbred        TTTTCCTTATGACACGAACCAAGATTTCGACGACAAGAACTTCATATGA   2311
BVillosa        TTTACCTTATGACACGAACCAAGATTTCGACGACAAGAACTTCATATGA   1540
```

#### Supplementary Figure 6

MAM3 C2 Alignment of the high glucoraphanin broccoli line HG Inbred, *Brassica villosa*, Ironman and the reference *Brassica oleracea* TO1000 CDS sequence taken from Ensembl Plants (Bolser, Staines et al. 2016) using the Clustal Omega Multiple Sequence Aligner (Sievers, Wilm et al. 2011).

```
CDS             ATGGCATCGTCACTTCTGACATCTTCCGGAATTATCCCTACCACCAGTTCCAACATGGTT 60
BVillosa        ATGGCATCGTCACTTCTGCCATCTTCCGGAATTATCCCTACCACCGGTTCCAACATGGTT 60
Ironman         ATGGCATCGTCACTTCTGACATCTTCCGGAATTATCCCTACCACCAGTTCCAACATGGTT 60
HGInbred        ATGGCATCGTCACTTCTGACATCTTCCGGAATTATCCCTACCACCGGTTCCAACATGGTT 60
                ****************** ************************** **************

CDS             GTCCGGTCATTCTTACCCTTTGGATCTGTCCGCCTGACCCGTCCGTACAACAAGTCGTCC 120
BVillosa        GTCCGGTCATTCTTACCCTTTGGATCTGTCCGCCTGACCCGTCCGTACAACAAGTCGTCC 120
Ironman         GTCCGGTCATTCTTACCCTTTGGATCTGTCCGCCTGACCCGTCCGTACAACAAGTCGTCC 120
HGInbred        GTCCGGTCATTCTTACCCTTTGGATCTGTCCGCCTGACCCGTCCGTACAACAAGTCGTCC 120
                ************************************************************

CDS             TTGCTCATCTCATGTTGCTCCTCTGTGTCCAAGAAGGCTGAGACTAGTGGTACTGACCTC 180
BVillosa        TTGCTCATCTCATGTTGCTCCTCTGTGTCCAAGAAGGCTGAGACTAGTGGTACTGACCTC 180
Ironman         TTGCTCATCTCATGTTGCTCCTCTGTGTCCAAGAAGGCTGAGACTAGTGGTACTGACCTC 180
HGInbred        TTGCTCATCTCATGTTGCTCCTCTGTGTCCAAGAAGGCTGAGACTAGTGGTACTGACCTC 180
                ************************************************************

CDS             AAAACCGTTGTGGAACGGTGGCCAGAGTATATTCCGAACAAGCTCCCCGACAAGAACTAC 240
BVillosa        AAAACCGTTGTGGAACGGTGGCCAGAGTATATTCCGAACAAGCTCCCCGACAAGAACTAC 240
Ironman         AAAACCGTTGTGGAACGGTGGCCAGAGTATATTCCGAACAAGCTCCCCGACAAGAACTAC 240
HGInbred        AAAACCGTTGTGGAACGGTGGCCAGAGTATATTCCGAACAAGCTCCCCGACAAGAACTAC 240
                ************************************************************

CDS             GTGCGTGTATTTGATACGACGCTCCGTGACGGCGAACAATCTCCCGGTGCAGCCCTTACT 300
BVillosa        GTGCGTGTATTTGATACGACGCTCCGTGACGGCGAACAATCTCCCGGTGCAGCCCTTACT 300
Ironman         GTGCGTGTATTTGATACGACGCTCCGTGACGGCGAACAATCTCCCGGTGCAGCCCTTACT 300
HGInbred        GTGCGTGTATTTGATACGACGCTCCGTGACGGCGAACAATCTCCCGGTGCAGCCCTTACT 300
                ************************************************************

CDS             CCACCGCAGAAGATAGAGATTGCCCGGCAGCTCGCTAAACTCAGAGTAGACATCATGGAA 360
BVillosa        CCACCGCAGAAGATAGAGATTGCCCGGCAGCTCGCTAAACTCAGAGTAGACATCATGGAA 360
Ironman         CCACCGCAGAAGATAGAGATTGCCCGGCAGCTCGCTAAACTCAGAGTAGACATCATGGAA 360
HGInbred        CCACCGCAGAAGATAGAGATTGCCCGGCAGCTCGCTAAACTCAGAGTAGACATCATGGAA 360
                ************************************************************

CDS             GTTGGTTTTCCCGTGTCATCAGAGGAAGAGTTCGAAACCATCAAAACCATCGCCAAAACC 420
BVillosa        GTTGGTTTTCCCGTGTCATCAGAGGAAGAGTTCGAAACCATCAAAACCATCGCCAAAACC 420
Ironman         GTTGGTTTTCCCGTGTCATCAGAGGAAGAGTTCGAAACCATCAAAACCATCGCCAAAACC 420
HGInbred        GTTGGTTTTCCCGTGTCATCAGAGGAAGAGTTCGAAACCATCAAAACCATCGCCAAAACC 420
                ************************************************************

CDS             GTGGGAAACGAGGTTGATGAGGAAACAGGTTATATCCCAGTGATATGCGTCATCGCACGA 480
BVillosa        GTGGGAAACGAGGTTGTTTCTTTATTTCCTTCTTCAAACTATATCGGTTTTTTTTCA-TA 479
Ironman         GTGGGAAACGAGGTTGTTTCTTTATTTCCTTCTTCAAACTATATCGTTTTTTTTCATAAG 480
HGInbred        GTGGGAAACGAGGTTGTTTCTTTATTTCCTTCTTCAAACTATATCGTTTTTTTTCATAAA 480
                **************** *     *     ** *         **     *  *

CDS             AGCAAAGAAAGAGATATAAAGGCGGCTTGGG------AGTCAGTGAAGTACGCAAAGAGG 534
BVillosa        AAAAAACT-----ATATCAAGTTTATTTGATAAAGAGAGTTTATAATGAAATTCAAATTA 534
Ironman         --AAAACT-----ATATCAAGTTTATTTGATAAAGAGAGTTTATAATGAAATTCAAATTA 533
HGInbred        AAAAAACT-----ATATCAAGTTTATTTGATAAAGAGAGTTTATAATGAAATTCAAATTA 535
```

#### Supplementary Figure 7

MAM3 C7 Alignment of the high glucoraphanin broccoli line HG Inbred, *Brassica villosa*, Ironman and the reference *Brassica oleracea* TO1000 CDS sequence taken from Ensembl Plants (Bolser, Staines et al. 2016) using the Clustal Omega Multiple Sequence Aligner (Sievers, Wilm et al. 2011).

```
CDS             ATGGCTTCGTCACTTCTGACATCTCCCACAATGATCCCCACCACCGGTTCCACTGTGGTG 60
BVillosa        ATGGCTTCATCACTTCTGACATCTCCCACAATGATACCCACCACCGGTTCCACTGTGGTT 60
HGInbred        ATGGCTTCGTCACTTCTGACATCTCCCACAATGATCCCCACCACCGGTTCCACTGTGGTG 60
Ironman         ATGGCTTCGTCACTTCTGACATCTCCCACAATGATCCCCACCACCGGTTCCACTGTGGTG 60
                ******** ************************** *********************** 

CDS             GTCCGATCAGGCTTACCTTTTAGATCTTCCTTGCCCTCCCTACGCCTGAATCGTCCGTAC 120
BVillosa        GTCCGATCAGGCTTACCTTT---ATCTTCCATGCCCTCCCTACGCCTGAATCGTCCGTAC 117
HGInbred        GTCCGATCAGGCTTACCTTTTAGATCTTCCTTGCCCTCCCTACGCCTGAATCGTCCGTAC 120
Ironman         GTCCGATCAGGCTTACCTTTTAGATCTTCCTTGCCCTCCCTACGCCTGAATCGTCCGTAC 120
                ********************   ******* *****************************

CDS             AACAAGTCATCCTTGTCTATCTCATGTTGCTCCCCTGTGTCTAAAATGGGGGGTACTAGT 180
BVillosa        AACAAGTCATCCTTGTCTATCTCATGTTGCTCCCCTGTGTCTAAAAAGGGGGGTACTAGT 177
HGInbred        AACAAGTCATCCTTGTCTATCTCATGTTGCTCCCCTGTGTCTAAAATGGGGGGTACTAGT 180
Ironman         AACAAGTCATCCTTGTCTATCTCATGTTGCTCCCCTGTGTCTAAAATGGGGGGTACTAGT 180
                ********************************************** *************

CDS             GCGGGTGACCTCAAACCCGTCTTGGAAAGGTGGCCAGAGTACATACCGCACAAGCTTCCC 240
BVillosa        GCGAGTGACCTCAAGCCCGTCGTGGAAAGGTGGCCAGAGTACATACCGCACAAGCTTCCC 237
HGInbred        GCGGGTGACCTCAAACCCGTCTTGGAAAGGTGGCCAGAGTACATACCGCACAAGCTTCCC 240
Ironman         GCGGGTGACCTCAAACCCGTCTTGGAAAGGTGGCCAGAGTACATACCGCACAAGCTTCCC 240
                *** ********** ****** **************************************

CDS             GACAAGAATTACGTGCGTGTATTCGACACGACGCTCCGTGACGGCGAACAAGCTCCAGGT 300
BVillosa        GACAAGAATTACGTGCGTGTATTCGACACGACGCTCCGTGACGGCGAACAAGCTCCAGGT 297
HGInbred        GACAAGAATTACGTGCGTGTATTCGACACGACGCTCCGTGACGGCGAACAAGCTCCAGGT 300
Ironman         GACAAGAATTACGTGCGTGTATTCGACACGACGCTCCGTGACGGCGAACAAGCTCCAGGT 300
                ************************************************************

CDS             GGAGCTCTTACTCCACCGCAGAAGCTTGAGATTGCTAGACAGCTCGCAAAACTCCGAGTA 360
BVillosa        GGAGCTCTTACTCCACCGCAGAAGCTTGGGATTGCAAGACAGCTCGCAAAACTCCGAGTA 357
HGInbred        GGAGCTCTTACTCCACCGCAGAAGCTTGAGATTGCTAGACAGCTCGCAAAACTCCGAGTA 360
Ironman         GGAGCTCTTACTCCACCGCAGAAGCTTGAGATTGCTAGACAGCTCGCAAAACTCCGAGTA 360
                **************************** ****** ************************

CDS             GACATCATGGAAGTTGGTTTTCCAGTGTCCTCTGAGGAAGAGTTCGAAACAGTCAAAACC 420
BVillosa        GACATCATGGAAGTTGGTTTTCCGGTGTCGTCTGAGGAAGAGTTCGAAACAGTCAAAACC 417
HGInbred        GACATCATGGAAGTTGGTTTTCCGGTGTCCTCTGAGGAAGAGTTCGAAACAGTCAAAACC 420
Ironman         GACATCATGGAAGTTGGTTTTCCGGTGTCCTCTGAGGAAGAGTTCGAAACAGTCAAAACC 420
                *********************** ***** ******************************

CDS             ATCGCCAAGACCGTGGGAAATGAGGTGGATGAAGCAACAGGTTACGTCCCAGTGATATGC 480
BVillosa        ATCGCCAAGACCGTTGGAAACGAGGTAATTGTTTCTAGTTATCAATTTTATGAATACTGT 477
HGInbred        ATCGCCAAGACCGTGGGAAATGAGGTTGTTTCTATATTTCCTTCAATTTATTATATAAGG 480
Ironman         ATCGCCAAGACCGTGGGAAATGAGGTTGTTTCTATATTTCCTTCAATTTATTATATAAGG 480
                ************** ***** *****   *           *    *           * 
```

#### Supplementary Figure 8

CYP79F1 Alignment of the high glucoraphanin broccoli line HG Inbred, *Brassica villosa*, Ironman and the reference *Brassica oleracea* TO1000 CDS sequence taken from Ensembl Plants (Bolser, Staines et al. 2016) using the Clustal Omega Multiple Sequence Aligner (Sievers, Wilm et al. 2011).

```
HGInbred        ATGACAATGATGATGATCCTTACCACATCGTTACCATACCCTTTTCAAATCCTACTAGTC 60
CDS             ATGACAATGATGATGAGCCTTACCACATCGTTACCATACCCTTTTCAAATCCTACTAGTC 60
Ironman         ATGACAATGATGATGAGCCTTACCACATCGTTACCATACCCTTTTCAAATCCTACTAGTC 60
BVillosa        ATGACAATGATGATGAGCCTTACCACATCGTTACCATACCCTTTTCAAATCCTACTAGTC 60
                **************** *******************************************

HGInbred        TTTATCTTATCCATGGCATCAATCACTTTGCTTGGCCGAATACTATCAAGACCCACCAAA 120
CDS             TTTATCTTATCCATGGCGTCAATCACTTTGCTTGGCCGAATACTATCAAGACCCACCAAA 120
Ironman         TTTAGCTTATCCATGGCATCAATCACTTTGCTTGGCCGAATACTATCAAGACCCTCCAAA 120
BVillosa        TTTATCTTATCCATGGCATCAATCACTTTGCTTGGCCGAATACTATCAAGACCCACCAAA 120
                **** ************ ************************************ *****

HGInbred        ACCAAAGACCGGTCTCGTCAGCTTCCTCCCGGACCACCAGGATGGCCCATCCTAGGCAAT 180
CDS             ACCAAAGACCGGTCTCGTCAGCTTCCTCCCGGACCACCAGGATGGCCCATCCTAGGCAAT 180
Ironman         ACCAAAAACCGGTCTCGTCAGCTTCCTCCCGGACCACCAGGATGGCCCATCCTAGGCAAT 180
BVillosa        ACCAAAGACCGGTCTCGTCAGCTTCCTCCCGGACCACCAGGATGGCCCATCCTAGGCAAT 180
                ****** *****************************************************

HGInbred        TTACCCGAACTAATGATGACTCGTCCTAGGCACAAATATGTCGACATTGCCATGAAAGGG 240
CDS             TTACCCGAACTAATGATGACTCGTCCTAGGCACAAATATGTCGGCATTGCCATGGAAGGG 240
Ironman         TTACCCGAACTAATGATGACTCGTCCTAGGCACAAATATGTCGGCATTGCCATGGAAGGG 240
BVillosa        TTACCCGAACTAATGATGACTCGTCCTAGGCACAAATATGTCGGCATTGCCATGGAAGGG 240
                ******************************************* ********** *****

HGInbred        CAAAAACCGGATATCGCATGTTTCAACTTCGCCGGAACACACGCCATCATCATAAACTCC 300
CDS             CAAAAACCGGATATCGCATGTTTCAACTTCGCCGGAACACACGCCATCATCATAAACTCC 300
Ironman         CAAAAACCGGATATCGCATGTTTCAACTTCGCCGGAACACACGCCATCATCATAAACTCC 300
BVillosa        CAAAAACCGGATATCGCATGTTTCAACTTCGCCGGAACACACGCCATCATCATAAACTCC 300
                ************************************************************

HGInbred        GACGAGATCGCTCGAGAAGCGCTTAAAGAGAGAGACGCCGACTTCGCAGACCGGCCTAAT 360
CDS             GACGAGATCGCTCGAGAAGCGCTTAAAGAGAGAGACGCCGACTTCGCAGACCGGCCTAAT 360
Ironman         GACGAGATCGCTCGAGAAGCGCTTAAAGAGAGAGACGCCGACTTCGCAGACCGGCCTAAT 360
BVillosa        GACGAGATCGCTCGAGAAGCGCTTAAAGAGAGAGACGCCGACTTCGCAGACCGGCCTAAT 360
                ************************************************************

HGInbred        CTTTTCAACATGAGAACCATCGGAGGCAATCACAAATCAATGGGGAACTCACCCTACGGT 420
CDS             CTTTTCAACATGAGAACCATCGGAGGCAATCACAAATCAATGGGGAACACACCCTACGGT 420
Ironman         CTTTTCAACATGAGAACCATCGGAGGCAATCACAAATCAATGGGGAACACACCCTACGGT 420
BVillosa        CTTTTCAACATGAGAACCATCGGAGGCAATCACAAATCAATGGGGAACACACCCTGCGGT 420
                ************************************************ ****** ****

HGInbred        GAACAGTTCATGAAGATGAAAAGAGTGATCACAACGGAGATTATGTCCGTTAAAACATTA 480
CDS             GAACAGTTCATGAAGATGAAAAGAGTGATCAGAACGGAGATTATGTCCGTTAAAACATTA 480
Ironman         GAACAGTTCATGAAGATGAAAAGAGTGATCAGAACGGAGATTATGTCCGTTAAAACATTA 480
BVillosa        GAACAGTTCATGAAGATGAAAAGAGTGATCAGAACGGAGATTATGTCCGTTAAAACATTA 480
                ******************************* ****************************

HGInbred        AACATGTTAGTAGCTGCGAGAACTATTGAAGCGGACAACCTCCTTGCTTACCTTCTCTCC 540
CDS             AACATGTTAGAAGCTGCGAGAACTATTGAAGCGGACAACCTCCTTGCTTACCTTCTCTCC 540
Ironman         AACATGTTAGAAGCTGCGAGAACTATTGAAGCGGACAACCTCCTTGCTTACCTTCTCTCC 540
BVillosa        AACATGTTAGAAGCTGCGAGAACTATTGAAGCGGACAACCTCCTTGCTTACCTTCTCTCC 540
                ********** *************************************************

HGInbred        ATGTACAAACGGTCTGAGACTGCTGACGTTAGAGAGTTCTCGAGGGTTTATGGTTACGCT 600
CDS             ATGTACAAACGGTCTGAGACTGCTGACGCTAGAGAGTTCTCGAGGGGTTATGGTTACGCT 600
Ironman         ATGTACAAACGGTCTGAGACTGCTGACGCTAGAGAGTTCTCGAGGGTTTATGGTTACGCT 600
BVillosa        ATGTACAAACGGTCTGAGACTGCTGACGCTAGAGAGTTCTCGAGGGTTTATGGTTACGCT 600
                **************************** ***************** *************
```

```
HGInbred        GTGACCATGAGATTGTTGTTTGGAAGGAGACATGTCACGAAAGAAAATGTTTTCTCCGAC 660
CDS             GTGACCATGAGATTGTTGTTTGGAAGGAGACATGTCACGAAAGAAAATGTTTTCTCCGAC 660
Ironman         GTGACCATGAGATTGTTGTTTGGAAGGAGACATGTCACGAAAGAAAATGTTTTCTCCGAC 660
BVillosa        GTGACCATGAGATTGTTGTTTGGAAGGAGACATGTCACGAAAGAAAATGTTTTCTCCGAC 660
                ************************************************************

HGInbred        GAGGGAAGGTTAGGACAAGCGGAGAAAGATCATCTTGATGCGATTTTCGAAACCTTAAAC 720
CDS             GAGGGAAGGTTAGGACAAGCGGAGAAAGATCATCTTGATGCGATTTTCGAAACCTTAAAC 720
Ironman         GAGGGAAGGTTAGGACAAGCGGAGAAAGATCATCTTGATGCGATTTTCGAAACCTTAAAC 720
BVillosa        GAGGGAAGGTTAGGACAAGCGGAGAAAGATCATCTTGATGCGATTTTCGAAACCTTAAAC 720
                ************************************************************

HGInbred        TGTTTGCCGAGTTTCTCTCCGGCGGATTACTTGGAAAAATGGTTTAGAGGTTGGAACATT 780
CDS             TGTTTGCCGAGTTTCTCTCCGGCGGATTACTTGGAAAAATGGTTTAGAGGTTGGAACATT 780
Ironman         TGTTTGCCGAGTTTCTCTCCGGCGGATTACTTGGAAAAATGGTTTAGAGGTTGGAACATT 780
BVillosa        TGTTTGCCGAGTTTCTCTCCGGCGGATTACTTGGAAAAATGGTTTAGAGGTTGGAACATT 780
                ************************************************************

HGInbred        GATGGTCAAGAGGAGAGGGTGGTAATGTACTGTAATAAAGTTCGCAGTTACAACAATCCG 840
CDS             GATGGTCAAGAGGAGAGGGTGGTAATGTCCTGTAATAAAGTTCGCAGTTACAACAATCCG 840
Ironman         GATGGTCAAGAGGAGAGGGTGGTAATGTCCTGTAATAAAGTTCGCAGTTACAACAATCCG 840
BVillosa        GATGGTCAAGAGGAGAGGGTGGTAATGTCCTGTAATAAAGTTCGCAGTTACAACAATCCG 840
                **************************** *******************************

HGInbred        ATCATTGACGAGAGGGTGGAGTTATGGAGGGAAAAAGGTGGTAAAGCAGCTGTTGAAGAT 900
CDS             ATCATTGACGAGAGGGTGGAGTTATGGAGGGAAAAAGGTGGTAAAGCATCTGTTGAAGAT 900
Ironman         ATCATTGACGAGAGGGTGGAGTTATGGAGGGAAAAAGGTGGTAAAGCATCTGTTGAAGAT 900
BVillosa        ATCATTGACGAGAGGGTGGAGTTATGGAGGGAAAAAGGTGGTAAAGCATCTGTTGAAGAT 900
                ************************************************ ***********

HGInbred        TGGATTGATACTTTCATTACGCTAAAAGATGAAAATGGAAAGTATTATATCACCCCGGAT 960
CDS             TGGATTGATACTTTCATTACGCTAAAAGATGAAAATGGAAAGTATTATATCACCCCGGAT 960
Ironman         TGGATTGATACTTTCATTACGCTAAAAGATGAAAATGGAAAGTATTATATCACCCCGGAT 960
BVillosa        TGGATTGATACTTTCATTACGCTAAAAGATGAAAATGGAAAGTATTATATCACCCCGGAT 960
                ************************************************************

HGInbred        GAGGTCAAAGCTCAATGCGTTGTAAGTAAACACTAGTACTATTTTACTACTATAATTGAT 1020
CDS             GAGGTCAAAGCTCAATGCGTTGAATTTTGTATAGCAGC---------------------- 998
Ironman         GAGGTCAAAGCTCAATGCGTTGTAAGTAAACACTAGTACTATTTTACTACTATAATTGAT 1020
BVillosa        GAGGTCAAAGCTCAATGCGTTGTAAGTAAACACTAGTACTATTTTACTACTATAATTGAT 1020
                ********************** *   *

HGInbred        TATATAACATTAGACAATTACAAGATATCAATGCAAAATATTATGATCATCGAAATAATA 1080
CDS             ------------------------------------------------------------ 998
Ironman         TATATAACATTAGACAATTACAAGATATTAATACAAAATATTATGATCATCGAAATAATA 1080
BVillosa        TATATAACATTAGACAATTACAAGATATTAA------------TGATCATCGAAATAATA 1068

HGInbred        CATTACAGTATAGATATACTACTGTTACTATTGAAAGTGGCAGTAGATAGGGATAGTTTT 1140
CDS             ------------------------------------------------------------ 998
Ironman         CATTACAGTATAGATATACTACTGATACTATTGAAAGTGGCAGTAGATAGGGATAGTTTT 1140
BVillosa        CATTACAGTATAGATATACTACTGATACTATTGAAAGTGGCAGTAGATAGGGATAGTTTT 1128

HGInbred        ACATGATTTTTGTTTCAAAAGAAATATGTCTACTAAAAAATAATATTTAGATTTTTTTTT 1200
CDS             ------------------------------------------------------------ 998
Ironman         ACATGATTTTTGTTTCAAAAGAAATATCTCTACTAAAAATGATATTTAGATTTTTTTTTT 1200
BVillosa        ACATGATTTTTGTTTCAAAAGAAATATCTCTACTAAAAATGATATTTAGATTTTTTTTTT 1188

HGInbred        GTTATATGGAGAATCAAGTTTTATTAATTGTTTTAGGATCCCATTAAATCTATTGTTTGA 1260
CDS             ------------------------------------------------------------ 998
Ironman         GTTATATGGAGAATCAAGTTTTATTAATTGTTTTAGGATCCCATTAAATCTATTGTTTGA 1260
BVillosa        GTTATATGGAGAATCAAGTTTTATTAATTGTTTTAGGATCCCATTAAATCTATTGTTTGA 1248

HGInbred        TGTTTTGTTTCAGGAATTTTGTATAGCAGCGATCGATAATCCGGCAAATAACATGGAATG 1320
CDS             ------------------------------GATCGATAATCCGGCAAATAACATGGAATG 1028
```

```
Ironman         TGTTTTGTTTCAGGAATTTTGTATAGCAGCGATCGATAATCCGGCAAATAACATGGAATG 1320
BVillosa        TGTTTTGTTTCAGGAATTTTGTATAGCAGCGATCGATAATCCGGCAAATAACATGGAATG 1308
                                              ******************************

HGInbred        GACACTTGCGGAAATGTTAAAGAACCCGGAGATTCTCAAAAAAGCTTTGAAGGAGTTAGA 1380
CDS             GACACTTGCGGAAATGTTAAAGAACCCGGAGATTCTCAAAAAAGCTTTGAAGGAGTTAGA 1088
Ironman         GACACTTGCGGAAATGTTAAAGAACCCGGAGATTCTCAAAAAAGCTTTGAAGGAGTTAGA 1380
BVillosa        GACACTTGCGGAAATGTTAAAGAACCCGGAGATTCTCAAAAAAGCTTTGAAGGAGTTAGA 1368
                ************************************************************

HGInbred        CGAAGTGGTGGGAAGAGACAGGCTTGTTCAGGAATCTGACATACCAAATCTAAACTACTT 1440
CDS             CGAAGTGGTGGGAAGAGACAGGCTTGTTCAGGAATCTGACATACCAAATCTAAACTATTT 1148
Ironman         CGAAGTGGTGGGAAGAGACAGGCTTGTTCAGGAATCTGACATACCAAATCTAAACTATTT 1440
BVillosa        CGAAGTGGTGGGAAGAGACAGGCTTGTTCAGGAATCTGACATACCAAATCTAAACTATTT 1428
                ********************************************************* **

HGInbred        GAAAGCTTGTTGCAGAGAAACATTCAGGATTCACCCTAGTGCTCATTATGTCCCTACACA 1500
CDS             AAAAGCTTGTTGCAGAGAAACATTCAGGATTCACCCTAGTGCTCATTATGTCCCTACACA 1208
Ironman         AAAAGCTTGTTGCAGAGAAACATTCAGGATTCACCCTAGTGCTCATTATGTCCCTACACA 1500
BVillosa        AAAAGCTTGTTGCAGAGAAACATTCAGGATTCACCCTAGTGCTCATTATGTCCCTACACA 1488
                 ***********************************************************

HGInbred        TGTGGCTCGTCAAGATACCACCCTCGGGGGTTATTTCATTCCCAAAGGTAAAAAGTTGTA 1560
CDS             TGTGGCTCGTCAAGATACCACCCTCGGGGGTTATTTCATTCCCAAAGGTAGCCACATTCA 1268
Ironman         TGTGGCTCGTCAAGATACCACCCTCGGGGGTTATTTCATTCCCAAAGGTAAAAAGTTGTA 1560
BVillosa        TGTGGCTCGTCAAGATACCACCCTCGGGGGTTATTTCATTCCCAAAGGTAAAAAGTTGTA 1548
                **************************************************    *   *   *
```

#### Supplementary Figure 9

SOT18 Alignment of the high glucoraphanin broccoli line HG Inbred, *Brassica villosa*, Ironman and the reference *Brassica oleracea* TO1000 CDS sequence taken from Ensembl Plants (Bolser, Staines et al. 2016) using the Clustal Omega Multiple Sequence Aligner (Sievers, Wilm et al. 2011).

```
CDS             ATGGCTTCGTCACTTCTGACATCTCCCACAATGATCCCCACCACCGGTTCCACTGTGGTG 60
BVillosa        ATGGCTTCATCACTTCTGACATCTCCCACAATGATACCCACCACCGGTTCCACTGTGGTT 60
HGInbred        ATGGCTTCGTCACTTCTGACATCTCCCACAATGATCCCCACCACCGGTTCCACTGTGGTG 60
Ironman         ATGGCTTCGTCACTTCTGACATCTCCCACAATGATCCCCACCACCGGTTCCACTGTGGTG 60
                ******** ************************** *********************** 

CDS             GTCCGATCAGGCTTACCTTTTAGATCTTCCTTGCCCTCCCTACGCCTGAATCGTCCGTAC 120
BVillosa        GTCCGATCAGGCTTACCTTT---ATCTTCCATGCCCTCCCTACGCCTGAATCGTCCGTAC 117
HGInbred        GTCCGATCAGGCTTACCTTTTAGATCTTCCTTGCCCTCCCTACGCCTGAATCGTCCGTAC 120
Ironman         GTCCGATCAGGCTTACCTTTTAGATCTTCCTTGCCCTCCCTACGCCTGAATCGTCCGTAC 120
                ********************   ******* *****************************

CDS             AACAAGTCATCCTTGTCTATCTCATGTTGCTCCCCTGTGTCTAAAATGGGGGGTACTAGT 180
BVillosa        AACAAGTCATCCTTGTCTATCTCATGTTGCTCCCCTGTGTCTAAAAAGGGGGGTACTAGT 177
HGInbred        AACAAGTCATCCTTGTCTATCTCATGTTGCTCCCCTGTGTCTAAAATGGGGGGTACTAGT 180
Ironman         AACAAGTCATCCTTGTCTATCTCATGTTGCTCCCCTGTGTCTAAAATGGGGGGTACTAGT 180
                ********************************************** *************

CDS             GCGGGTGACCTCAAACCCGTCTTGGAAAGGTGGCCAGAGTACATACCGCACAAGCTTCCC 240
BVillosa        GCGAGTGACCTCAAGCCCGTCGTGGAAAGGTGGCCAGAGTACATACCGCACAAGCTTCCC 237
HGInbred        GCGGGTGACCTCAAACCCGTCTTGGAAAGGTGGCCAGAGTACATACCGCACAAGCTTCCC 240
Ironman         GCGGGTGACCTCAAACCCGTCTTGGAAAGGTGGCCAGAGTACATACCGCACAAGCTTCCC 240
                *** ********** ****** **************************************

CDS             GACAAGAATTACGTGCGTGTATTCGACACGACGCTCCGTGACGGCGAACAAGCTCCAGGT 300
BVillosa        GACAAGAATTACGTGCGTGTATTCGACACGACGCTCCGTGACGGCGAACAAGCTCCAGGT 297
HGInbred        GACAAGAATTACGTGCGTGTATTCGACACGACGCTCCGTGACGGCGAACAAGCTCCAGGT 300
Ironman         GACAAGAATTACGTGCGTGTATTCGACACGACGCTCCGTGACGGCGAACAAGCTCCAGGT 300
                ************************************************************

CDS             GGAGCTCTTACTCCACCGCAGAAGCTTGAGATTGCTAGACAGCTCGCAAAACTCCGAGTA 360
BVillosa        GGAGCTCTTACTCCACCGCAGAAGCTTGGGATTGCAAGACAGCTCGCAAAACTCCGAGTA 357
HGInbred        GGAGCTCTTACTCCACCGCAGAAGCTTGAGATTGCTAGACAGCTCGCAAAACTCCGAGTA 360
Ironman         GGAGCTCTTACTCCACCGCAGAAGCTTGAGATTGCTAGACAGCTCGCAAAACTCCGAGTA 360
                **************************** ****** ************************

CDS             GACATCATGGAAGTTGGTTTTCCAGTGTCCTCTGAGGAAGAGTTCGAAACAGTCAAAACC 420
BVillosa        GACATCATGGAAGTTGGTTTTCCGGTGTCGTCTGAGGAAGAGTTCGAAACAGTCAAAACC 417
HGInbred        GACATCATGGAAGTTGGTTTTCCGGTGTCCTCTGAGGAAGAGTTCGAAACAGTCAAAACC 420
Ironman         GACATCATGGAAGTTGGTTTTCCGGTGTCCTCTGAGGAAGAGTTCGAAACAGTCAAAACC 420
                *********************** ***** ******************************

CDS             ATCGCCAAGACCGTGGGAAATGAGGTGGATGAAGCAACAGGTTACGTCCCAGTGATATGC 480
BVillosa        ATCGCCAAGACCGTTGGAAACGAGGTAATTGTTTCTAGTTATCAATTTTATGAATACTGT 477
HGInbred        ATCGCCAAGACCGTGGGAAATGAGGTTGTTTCTATATTTCCTTCAATTTATTATATAAGG 480
Ironman         ATCGCCAAGACCGTGGGAAATGAGGTTGTTTCTATATTTCCTTCAATTTATTATATAAGG 480
                ************** ***** *****   *           *    *           * 
```

#### Supplementary Figure 10

APS3 Alignment of the high glucoraphanin broccoli line HG Inbred, *Brassica villosa*, Ironman and the reference Brassica oleracea TO1000 CDS sequence taken from Ensembl Plants (Bolser, Staines et al. 2016) using the Clustal Omega Multiple Sequence Aligner (Sievers, Wilm et al. 2011).

```
BVillosa        ATGGCTTCCATGTCCACCGCCTTCCCCAAAACAACTCCTTTCCTCTCTCATCCTCTCACC 60
CDS             ATGGCTTCCATGTCCACCGCCTTCCCCAAAACAACTCCTTTCCTCTCTCATCCTCTCACC 60
Ironman         ATGGCTTCCATGTCCACCGCCTTCCCCAAAACAACTCCTTTCCTCTCTCATCCTCTCACC 60
HGInbred        ATGGCTTCCATGTCCACCGCCTTCCCCAAAACAACTCCTTTCCTCTCTCATCCTCTCACC 60
                ************************************************************

BVillosa        AAATCCGATTCCCTGATCGCAACCACCGTTTCATTTCCCTCCAGCGCGAGATCTCGCAGC 120
CDS             AAATCCGATTCCCTGATCGCAACCACCGTTTCATTTCCCTCCAGCGCGAGATCTCGCAGC 120
Ironman         AAATCCGATTCCCTGATCGCAACCACCGTTTCATTTCCCTCCAGCGCGAGATCTCGCAGC 120
HGInbred        AAATCCGATTCCCTGATCGCAACCACCGTTTCATTTCCCTCCAGCGCGAGATCTCGCAGC 120
                ************************************************************

BVillosa        CTCCGAAGAGCCACCCTCCGCGTACGATCCGCGCTGATCGAGCCAGACGGCGGGAAGCTG 180
CDS             CTCCGAAGAGCCACCCTCCGCGTACGATCCGCGCTGATCGAGCCAGACGGCGGGAAGCTG 180
Ironman         CTCAGAAGAGCCACCCTCCGCGTACGATCCGCGCTGATCGAGCCAGACGGCGGGAAGCTG 180
HGInbred        CTCAGAAGAGCCACCCTCCGCGTACGATCCGCGCTGATCGAGCCAGACGGCGGGAAGCTG 180
                *** ********************************************************

BVillosa        GTGGAGCTTGTGGTCCCGGAGACGCGACGAAGGGAGAAGAAGCACGAGGCGGCGGATTTG 240
CDS             GTGGAGCTTGTGGTCCCGGAGACGCGACGAAGGGAGAAGAAGCACGAGGCGGCGGATTTG 240
Ironman         GTGGAGCTTGTGGTCCCGGAGACGCGACGAAGGGAGAAGAAGCACGAGGCGGCGGATTTG 240
HGInbred        GTGGAGCTTGTGGTCCCGGAGACGCGACGAAGGGAGAAGAAGCACGAGGCGGCGGATTTG 240
                ************************************************************

BVillosa        CCGAGAGTGAAACTGACGGCGATCGATTTGCAGTGGATGCACGTGTTGAGCGAAGGCTGG 300
CDS             CCGAGAGTGAAACTGACGGCGATCGATTTGCAGTGGATGCACGTGTTGAGCGAAGGCTGG 300
Ironman         CCGAGAGTGAAACTGACGGCGATCGATTTGCAGTGGATGCACGTGTTGAGCGAAGGCTGG 300
HGInbred        CCGAGAGTGAAACTGACGGCGATCGATTTGCAGTGGATGCACGTGTTGAGCGAAGGCTGG 300
                ************************************************************

BVillosa        GCAAGTCCGCTTAGAGGGTTTATGCGGGAAGCTGAGTTCCTTCAAACTCTTCATTTCAAT 360
CDS             GCAAGTCCGCTTAGAGGGTTTATGCGGGAAGCTGAGTTCCTTCAAACTCTTCATTTCAAT 360
Ironman         GCAAGTCCGCTTAGAGGGTTTATGCGGGAAGCTGAGTTCCTTCAAACTCTTCATTTCAAT 360
HGInbred        GCAAGTCCGCTTAGAGGGTTTATGCGGGAAGCTGAGTTCCTTCAAACTCTTCATTTCAAT 360
                ************************************************************

BVillosa        TCGCTTCGTCTCGACGACGGATCCGTCGTCAATATGTCTGTACCTATCGTTCTCGCCATC 420
CDS             TCGCTTCGTCTCGACGACGGATCCGTCGTCAATATGTCTGTACCTATCGTTCTCGCCATC 420
Ironman         TCGCTTCGTCTCGACGACGGATCCGTCGTCAATATGTCTGTACCTATCGTTCTCGCCATC 420
HGInbred        TCGCTTCGTCTCGACGACGGATCCGTCGTCAATATGTCTGTACCTATCGTTCTCGCCATC 420
                ************************************************************

BVillosa        GATGATGAACAGAAAGCTCTAATAGGAGAATCTAAACGTGTCGCGCTTGTTGATTCCGAT 480
CDS             GATGATGAACAGAAAGCTCTAATCGGAGAATCTAAACGTGTCGCGCTTGTTGATTCCGAT 480
Ironman         GATGATGAACAGAAAGCTCTAATCGGAGAATCTAAACGTGTCGCGCTTGTTGATTCCGAT 480
HGInbred        GATGATGAACAGAAAGCTCTAATCGGAGAATCTAAACGTGTCGCGCTTGTTGATTCCGAT 480
                *********************** ************************************

BVillosa        GATAATCCAATCGCTATTCTCAGCGAGTAAGTTCATCCTC---CTTGTTAATCTTGAGAT 537
CDS             GATAATCCAATCGCTATTCTCAGCG----------------------------------- 505
Ironman         GATAATCCAATCGCTATTCTCAGCGAGTAAGTTCATCCTCCTCGTTGTTAATCTTGAGAT 540
HGInbred        GATAATCCAATCGCTATTCTCAGCGAGTAAGTTCATCCTCCTCGTTGTTAATCTTGAGAT 540
                *************************

BVillosa        CTCTGCTGTGGTTTCGGGCCTGGTTAATCATGGGCTTTGGCCCAAAACCTTCTCGAATTG 597
CDS             ------------------------------------------------------------ 505
Ironman         CTCTGCTGTGGTTTCGGGCCTGGTGAATCATGGGCTTTGGCCCAAAACCTTCTCGAATTG 600
HGInbred        CTCTGCTGTGGTTTCGGGCCTGGTGAATCATGGGCTTTGGCCCAAAACCTTCTCGAATTG 600
```

```
BVillosa        AAAAAAGAAACTTGTAGCCGGACTAGTCTTTCATACTAATATGAGATTAGTTGGCTTCTT 657
CDS             ------------------------------------------------------------ 505
Ironman         AAAAAAAAACTTGTAG----------CCGGACATACTAATATGAGATTAGTTGGCTTCTT 650
HGInbred        AAAAAAAAACTTGTAG----------CCGGACATACTAATATGAGATTAGTTGGCTTCTT 650

BVillosa        CGGTCAATAAAAAAAAAAACAAGTGAATAGGTTCCTCCAAATTTTAAATATGATTTAATC 717
CDS             ------------------------------------------------------------ 505
Ironman         CGGTCAATAAAA-AAAAAACAAGTGAATAGGTTCCTCCAAATTTTAAATATGATTTAATC 709
HGInbred       CGGTCAATAAAA-AAAAAACAAGTGAATAGGTTCCTCCAAATTTTAAATATGATTTAATC  709

BVillosa        AATCTTAGATTTT-------------------GTAATTCCTCTGTTGCTTGAGACATTTT 758
CDS             ------------------------------------------------------------ 505
Ironman         AATCTTAGATTTTGTAGTTTTTTTCTTCTTCTGTAATTCCTCTGTTGCTTGAGACATTTT 769
HGInbred        AATCTTAGATTTTGTAGTTTTTTTCTTCTTCTGTAATTCCTCTGTTGCTTGAGACATTTT 769

BVillosa        TAGGAATCAATCTTCGCATCACTTAATAGTATATTTATAGTTTGCTTTAACAAATATGTG 818
CDS             ------------------------------------------------------------ 505
Ironman         TAGGAATCAATCTTCGCATCACTTAATAGTATATTTATAGTTTGCTTTAACAAATATGTG 829
HGInbred        TAGGAATCAATCTTCGCATCACTTAATAGTATATTTATAGTTTGCTTTAACAAATATGTG 829

BVillosa        CTTGTTTGTTTTGGTTTCCTCAGCATTGAGATATACAAACATCCCAAAGAGGAAAGAATA 878
CDS             ----------------------ACATTGAGATATACAAACATCCCAAAGAGGAAAGAATA 543
Ironman         CTTGTTTGTTTTGGTTTCCTCAGCATTGAGATATACAAACATCCCAAAGAGGAAAGAATA 889
HGInbred        CTTGTTTGTTTTGGTTTCCTCAGCATTGAGATATACAAACATCCCAAAGAGGAAAGAATA 889
                                       *************************************

BVillosa        GCTAGGACTTGGGGCACGACCGCACCGGGCTTGCCTTTTGCGGAAGAGGCGATAACCAAT 938
CDS             GCTAGAACTTGGGGCACGACCGCACCGGGCTTGCCTTTTGCGGAAGAGGCGATAACCAAT 603
Ironman         GCTAGAACTTGGGGCACGACCGCACCGGGCTTGCCTTTTGCGGAAGAGGCGATAACCAAT 949
HGInbred        GCTAGAACTTGGGGCACGACCGCACCGGGCTTGCCTTTTGCGGAAGAGGCGATAACCAAT 949
                ***** ******************************************************

BVillosa        GCTGGAAACTGGCTCATTGGTGGTGATCTTGAGGTTCTCGAACCTGTCAAGTACAATGAC 998
CDS             GCTGGAAACTGGCTCATTGGTGGTGATCTTGAGGTTCTCGAACCTGTCAAGTACAATGAC 663
Ironman         GCTGGAAACTGGCTCATTGGTGGTGATCTTGAGGTTCTCGAACCTGTCAAGTACAATGAC 1009
HGInbred        GCTGGAAACTGGCTCATTGGTGGTGATCTTGAGGTTCTCGAACCTGTCAAGTACAATGAC 1009
                ************************************************************

BVillosa        GGGCTTGACCGTTTCAGGCTTTCGCCATTTGAACTCCGGAAAGAGCTTGAGAAACGTGGC 1058
CDS             GGGCTTGACCGTTTCAGGCTTTCGCCATTTGAACTCCGGAAAGAGCTTGAGAAACGTGGC 723
Ironman         GGGCTTGACCGTTTCAGGCTTTCGCCATTTGAACTCCGGAAAGAGCTTGAGAAACGTGGC 1069
HGInbred        GGGCTTGACCGTTTCAGGCTTTCGCCATTTGAACTCCGGAAAGAGCTTGAGAAACGTGGC 1069
                ************************************************************

BVillosa        GCGGATGCTGTCTTTGCGTTCCAGCTCAGGAACCCTGTTCACAACGGACACGCCCTTCTC 1118
CDS             GCGGATGCTGTCTTTGCGTTCCAGCTCAGGAACCCTGTTCACAACGGACACGCCCTTCTC 783
Ironman         GCGGATGCTGTCTTTGCGTTCCAGCTCAGGAACCCTGTTCACAACGGACACGCCCTTCTC 1129
HGInbred        GCGGATGCTGTCTTTGCGTTCCAGCTCAGGAACCCTGTTCACAACGGACACGCCCTTCTC 1129
                ************************************************************

BVillosa        ATGACTGACACGCGCCGGAGACTGCTTGAGATGGGTTACAAAAACCCGATCCTTTTGCTT 1178
CDS             ATGACTGACACGCGCCGGAGACTGCTTGAGATGGGTTACAAAAACCCGATCCTTTTGCTT 843
Ironman         ATGACTGACACGCGCCGGAGACTGCTTGAGATGGGTTACAAAAACCCGATCCTTTTGCTT 1189
HGInbred        ATGACTGACACGCGCCGGAGACTGCTTGAGATGGGTTACAAAAACCCGATCCTTTTGCTT 1189
                ************************************************************

BVillosa        CATCCACTTGGAGGGTTCACAAAGGCGGATGATGTTCCTCTAAGCTGGCGAATGAAACAG 1238
CDS             CATCCACTTGGAGGGTTCACAAAGGCGGATGATGTTCCTCTAAGCTGGCGAATGAAACAG 903
Ironman         CATCCACTTGGAGGGTTCACAAAGGCGGATGATGTTCCTCTAAGCTGGCGAATGAAACAG 1249
HGInbred        CATCCCCTTGGAGGGTTCACAAAGGCGGATGATGTTCCTCTAAGCTGGCGAATGAAACAG 1249
                ***** ******************************************************

BVillosa        CACGAGAAGGTGCTAGAGGATGGTGTTCTTGATCCGGAGACGACTGTGGTTTCCATATTC 1298
CDS             CACGAGAAGGTGCTAGAGGATGGTGTTCTTGATCCGGAGACGACTGTGGTTTCCATATTC 963
```

```
Ironman         CACGAGAAGGTGCTAGAGGATGGTGTTCTTGATCCGGAGACGACTGTGGTTTCCATATTC 1309
HGInbred        CACGAGAAGGTGCTAGAGGATGGTGTTCTTGATCCGGAGACGACTGTGGTTTCCATATTC 1309
                ************************************************************

BVillosa        CCATCTCCAATGCTTTACGCTGGTCCAACCGAAGTGCAGTGGCATGCAAAGGCTAGGATC 1358
CDS             CCATCTCCAATGCTTTACGCTGGTCCAACCGAAGTGCAGTGGCATGCAAAGGCTAGGATC 1023
Ironman         CCATCTCCAATGCTTTACGCTGGTCCAACCGAAGTGCAGTGGCATGCAAAGGCTAGGATC 1369
HGInbred        CCATCTCCAATGCTTTACGCTGGTCCAACCGAAGTGCAGTGGCATGCAAAGGCTAGGATC 1369
                ************************************************************

BVillosa        AATGCCGGTGCCAATTTCTACATTGTGGGTAGAGATCCAGCTGGAATGGGTCACCCTGTG 1418
CDS             AATGCCGGTGCCAATTTCTACATTGTGGGTAGAGATCCAGCTGGAATGGGTCACCCTGTG 1083
Ironman         AATGCCGGTGCCAATTTCTACATTGTGGGTAGAGATCCAGCTGGAATGGGTCACCCTGTG 1429
HGInbred        AATGCCGGTGCCAATTTCTACATTGTGGGTAGAGATCCAGCTGGAATGGGTCACCCTGTG 1429
                ************************************************************

BVillosa        GAGAAACGTGATCTGTATGATGCTGATCACGGGAAGAAAGTACTAAGCATGGCTCCTGGA 1478
CDS             GAGAAACGTGATCTGTATGATGCTGATCACGGGAAGAAAGTACTAAGCATGGCTCCTGGA 1143
Ironman         GAGAAACGTGATCTGTATGATGCTGATCACGGGAAGAAAGTACTAAGCATGGCTCCTGGA 1489
HGInbred        GAGAAACGTGATCTGTATGATGCTGATCACGGGAAGAAAGTACTAAGCATGGCTCCTGGA 1489
                ************************************************************

BVillosa        CTCGAACGGCTCAACATTCTTCCTTTCAGGGTATATTTAAATCTGCATACTCAACCTCGT 1538
CDS             CTCGAACGGCTCAACATTCTTCCTTTCAGGGTTGCTGCGT-------------------- 1183
Ironman         CTCGAACGGCTCAACATTCTTCCTTTCAGGGTACATTTAAATCTGCATACTCAACCTCGG 1549
HGInbred        CTCGAACGGCTCAACATTCTTCCTTTCAGGGTACATTTAAATCTGCATACTCAACCTCGG 1549
                ********************************   *

BVillosa        TATCTCCTTCAAAGATTTAAGGCTTTTGAGTTCTGATTTTTCTGGTTTCAATCTATTTAG 1598
CDS             ------------------------------------------------------------ 1183
Ironman         TATCTCCTTCAAAGATTTA-GGCTTTTGAGTTCTGATTTTTCTGGTTTCAATCTGTTTAG 1608
HGInbred        TATCTCCTTCAAAGATTTA-GGCTTTTGAGTTCTGATTTTTCTGGTTTCAATCTGTTTAG 1608

BVillosa        GTTGCTGCGTATGATAAGACACAAGGCAAGATGGCCTTCTTTGATCCATCGAGGGCTCAA 1658
CDS             ----------ATGATAAGACACAAGGCAAGATGGCCTTCTTTGATCCATCGAGGGCTCAA 1233
Ironman         GTTGCTGCGTATGATAAGACACAAGGCAAGATGGCCTTCTTTGATCCATCGAGGGCTCAA 1668
HGInbred        GTTGCTGCGTATGATAAGACACAAGGCAAGATGGCCTTCTTTGATCCATCGAGGGCTCAA 1668
                          **************************************************

BVillosa        GACTTCTTGTTCATCTCTGGCACTAAGGTAACTACAACTTCTCAGACTCATTGAGACATA 1718
CDS             GACTTCTTGTTCATCTCTGGCACTAAG--------------------------------- 1260
Ironman         GACTTCTTGTTCATCTCTGGCACTAAGGTAACTACAACTTCTCAGACTCATTGAGACATA 1728
HGInbred        GACTTCTTGTTCATCTCTGGCACTAAGGTAACTACAACTTCTCAGACTCATTGAGACATA 1728
                ***************************

BVillosa        ATAGAGTACAGACTGTTCTGTTTCTGATTAAAACTTGAATTTGGTTTGGTTCTATAGATG 1778
CDS             ---------------------------------------------------------ATG 1263
Ironman         ATAGAGTACAGAGTGTTCTGTTTCTGGTTAAAGCTTGAATTTGGTTTGGTTCTATAGATG 1788
HGInbred        ATAGAGTACAGAGTGTTCTGTTTCTGGTTAAAGCTTGAATTTGGTTTGGTTCTATAGATG 1788
                                                                         ***

BVillosa        CGAGGACTGGCAAAGAACAAAGAAAGCCCACCAGATGGATTTATGTGCCCCGGAGGCTGG 1838
CDS             CGAGGACTGGCAAAGAACAAAGAAAGCCCACCAGATGGATTTATGTGCCCCGGAGGCTGG 1323
Ironman         CGAGGACTGGCAAAGAACAAAGAAAGCCCACCAGATGGATTTATGTGCCCCGGAGGCTGG 1848
HGInbred        CGAGGACTGGCAAAGAACAAAGAAAGCCCACCAGATGGATTTATGTGCCCCGGAGGCTGG 1848
                ************************************************************

BVillosa        AAAGTGCTTGTTGATTACTACGATAGTTTGAGTCTATCCGGAAATGCCAGACTCCCGGAA 1898
CDS             AAAGTGCTTGTTGATTACTACGATAGTTTGAGTCTATCCGGAAATGCCAGACTCCCGGAA 1383
Ironman         AAAGTGCTTGTTGATTACTACGATAGTTTGAGTCTATCCGGAAATGCCAGACTCCCGGAA 1908
HGInbred        AAAGTGCTTGTTGATTACTACGATAGTTTGAGTCTATCCGGAAATGCCAGACTCCCGGAA 1908
                ************************************************************

BVillosa        AAGGTTCCGGTTTAA   1913
CDS             AAGGTTCCGGTTTAA   1398
Ironman         AAGGTTCCGGTTTAA   1923
HGInbred        AAGGTTCCGGTTTAA   1923
```

#### Supplementary Figure 11

Confirmation of three significantly upregulated genes in MYB28$^{v/v}$ broccoli in comparison to the MYB28$^{b/b}$ broccoli sulphur metabolism using qRT-PCR. Expression normalised to Ubiquitin-C. Unpaired t-test performed between lines, n=3, * = p<0.05, ** = p<0.01, *** = p<0.001.

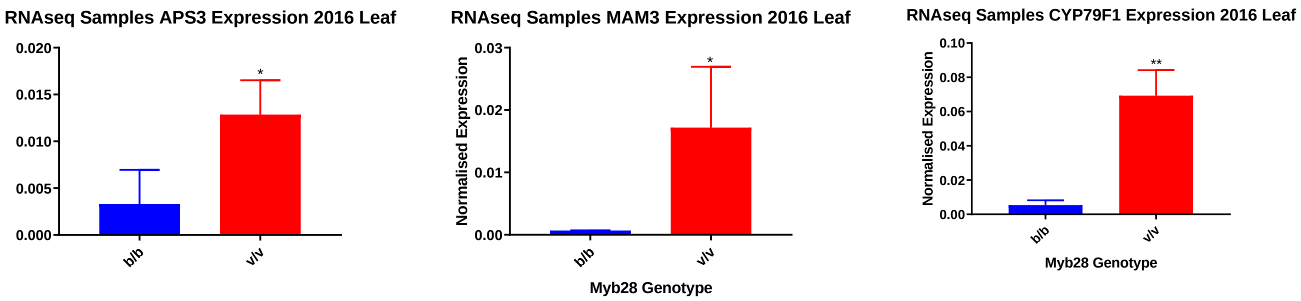

#### Metabolite Analysis

##### Supplementary Figure 12

Plot layout of plants for study year 1 (top) and study year 2 (bottom). Individual plants were given a number and colour code throughout analysis. Plots were generated using a random number generator and were randomised with blocks for column and row. Note: Year 2 also included an additional line '1086' which was also sequenced in the whole genome sequence analysis. Data for this line are not included in the current manuscript.

### Year 1

| | | | | | |
|---|---|---|---|---|---|
| 13 | 29 | 21 | 12 | 4 | 10 |
| 22 | 18 | 1 | 7 | 16 | 30 |
| 9 | 14 | 19 | 5 | 24 | 26 |
| 3 | 27 | 28 | 17 | 6 | 11 |
| 20 | 2 | 8 | 23 | 25 | 15 |

### Year 2

| | | | | | |
|---|---|---|---|---|---|
| 25 | 1 | 15 | 28 | 35 | 17 |
| 36 | 21 | 2 | 26 | 16 | 6 |
| Blank | 5 | 20 | 4 | 29 | 37 |
| 22 | 19 | 30 | 12 | 31 | 10 |
| 13 | 34 | 7 | Blank | 8 | 27 |
| 3 | 23 | 32 | Blank | 14 | 33 |
| 18 | Blank | 24 | 38 | 9 | 11 |

### Key

| |
|---|
| 1-10 = Ironman |
| 11-20 = 1199 |
| 21-30 = HG Inbred |
| 31-38 = 1086 |
| Blank Plants |

#### Supplementary Table 8

Two-way ANOVA with Sidak's multiple comparisons test of glucoraphanin levels in leaves compared to florets of plants grown in the first trial year (2016) GraphPad Prism v.8.2.0

| | | | | | |
|---|---|---|---|---|---|
| Table Analyzed | Glucoraphanin Leaves vs Florets 2016 | | | | |
| Two-way ANOVA | Ordinary | | | | |
| Alpha | 0.05 | | | | |
| Source of Variation | % of total variation | P value | P value summary | Significant? | |
| Interaction | 1.38 | 0.1086 | ns | No | |
| Row Factor | 60.54 | <0.0001 | **** | Yes | |
| Column Factor | 20.34 | <0.0001 | **** | Yes | |
| ANOVA table | SS (Type III) | DF | MS | F (DFn, DFd) | P value |
| Interaction | 149.5 | 2 | 74.73 | F (2, 53) = 2.315 | P=0.1086 |
| Row Factor | 6556 | 2 | 3278 | F (2, 53) = 101.6 | P<0.0001 |
| Column Factor | 2203 | 1 | 2203 | F (1, 53) = 68.25 | P<0.0001 |
| Residual | 1711 | 53 | 32.28 | | |
| Difference between column means | | | | | |
| Predicted (LS) mean of Leaves | 16.65 | | | | |
| Predicted (LS) mean of Florets | 28.88 | | | | |
| Difference between predicted means | -12.23 | | | | |
| SE of difference | 1.48 | | | | |
| 95% CI of difference | -15.20 to -9.261 | | | | |
| Compare each cell mean with the other cell mean in that row | | | | | |
| Number of families | 1 | | | | |
| Number of comparisons per family | 3 | | | | |
| Alpha | 0.05 | | | | |
| Sidak's multiple comparisons test | Predicted (LS) mean diff. | 95.00% CI of diff. | Significant? | Summary | Adjusted P Value |

| | | | | | | | | |
|---|---|---|---|---|---|---|---|---|
| Leaves - Florets | | | | | | | | |
| MYB28$^{b/b}$ | -8.692 | -14.96 to -2.428 | Yes | ** | 0.0036 | | | |
| MYB28$^{b/v}$ | -11.54 | -17.81 to -5.278 | Yes | **** | <0.0001 | | | |
| MYB28$^{v/v}$ | -16.46 | -22.89 to -10.02 | Yes | **** | <0.0001 | | | |
| Test details | Predicted (LS) mean 1 | Predicted (LS) mean 2 | Predicted (LS) mean diff. | SE of diff. | N1 | N2 | t | DF |
| Leaves - Florets | | | | | | | | |
| MYB28$^{b/b}$ | 5.898 | 14.59 | -8.692 | 2.541 | 10 | 10 | 3.421 | 53 |
| MYB28$^{b/v}$ | 16.11 | 27.65 | -11.54 | 2.541 | 10 | 10 | 4.543 | 53 |
| MYB28$^{v/v}$ | 27.95 | 44.41 | -16.46 | 2.61 | 9 | 10 | 6.304 | 53 |

#### Supplementary Table 9

Two-way ANOVA with Sidak's multiple comparisons test of glucoraphanin levels in leaves compared to florets of plants grown in the second trial year (2017) performed using GraphPad Prism v.8.2.0

| | | | | | |
|---|---|---|---|---|---|
| Table Analyzed | Glucoraphanin Leaves vs Florets 2017 | | | | |
| Two-way ANOVA | Ordinary | | | | |
| Alpha | 0.05 | | | | |
| Source of Variation | % of total variation | P value | P value summary | Significant? | |
| Interaction | 8.096 | <0.0001 | **** | Yes | |
| Row Factor | 50.76 | <0.0001 | **** | Yes | |
| Column Factor | 36.75 | <0.0001 | **** | Yes | |
| ANOVA table | SS (Type III) | DF | MS | F (DFn, DFd) | P value |
| Interaction | 174.1 | 2 | 87.05 | F (2, 39) = 14.68 | P<0.0001 |
| Row Factor | 1092 | 2 | 545.8 | F (2, 39) = 92.02 | P<0.0001 |
| Column Factor | 790.3 | 1 | 790.3 | F (1, 39) = 133.2 | P<0.0001 |
| Residual | 231.3 | 39 | 5.931 | | |
| Difference between column means | | | | | |
| Predicted (LS) mean of Leaves | 4.174 | | | | |
| Predicted (LS) mean of Florets | 13.06 | | | | |
| Difference between predicted means | -8.89 | | | | |
| SE of difference | 0.7701 | | | | |
| 95% CI of difference | -10.45 to -7.332 | | | | |
| Data summary | | | | | |
| Number of columns (Column Factor) | 2 | | | | |
| Number of rows (Row Factor) | 3 | | | | |
| Number of values | 45 | | | | |
| Compare each cell mean with the other cell mean in that row | | | | | |
| Number of families | 1 | | | | |

| | | | | | | | | |
|---|---|---|---|---|---|---|---|---|
| Number of comparisons per family | 3 | | | | | | | |
| Alpha | 0.05 | | | | | | | |
| Sidak's multiple comparisons test | Predicted (LS) mean diff. | 95.00% CI of diff. | Significant? | Summary | Adjusted P Value | | | |
| Leaves - Florets | | | | | | | | |
| MYB28$^{b/b}$ | -3.035 | -6.363 to 0.2923 | No | ns | 0.0829 | | | |
| MYB28$^{b/v}$ | -11.18 | -14.51 to -7.851 | Yes | **** | <0.0001 | | | |
| MYB28$^{v/v}$ | -12.46 | -15.78 to -9.128 | Yes | **** | <0.0001 | | | |
| Test details | Predicted (LS) mean 1 | Predicted (LS) mean 2 | Predicted (LS) mean diff. | SE of diff. | N1 | N2 | t | DF |
| Leaves - Florets | | | | | | | | |
| MYB28$^{b/b}$ | 0.8809 | 3.916 | -3.035 | 1.334 | 10 | 5 | 2.276 | 39 |
| MYB28$^{b/v}$ | 2.687 | 13.87 | -11.18 | 1.334 | 10 | 5 | 8.38 | 39 |
| MYB28$^{v/v}$ | 8.953 | 21.41 | -12.46 | 1.334 | 10 | 5 | 9.338 | 39 |

#### Supplementary Table 10

| Leaf MYB28 Expression | | | | | |
|---|---|---|---|---|---|
| **Tukey's multiple comparisons test** | **Predicted (LS) mean diff.** | **95.00% CI of diff.** | **Significant?** | **Summary** | **Adjusted P Value** |
| **Year 1** | | | | | |
| MYB28b/b vs. MYB28v/b | -0.1033 | -0.1751 to -0.03163 | Yes | ** | 0.0033 |
| MYB28b/b vs. MYB28v/v | -0.1172 | -0.1870 to -0.04742 | Yes | *** | 0.0006 |
| MYB28v/b vs. MYB28v/v | -0.01387 | -0.08559 to 0.05784 | No | ns | 0.8842 |
| **Year 2** | | | | | |
| MYB28b/b vs. MYB28v/b | -0.1265 | -0.2369 to -0.01618 | Yes | * | 0.0216 |
| MYB28b/b vs. MYB28v/v | -0.3282 | -0.4386 to -0.2179 | Yes | **** | <0.0001 |
| MYB28v/b vs. MYB28v/v | -0.2017 | -0.3120 to -0.09132 | Yes | *** | 0.0002 |
| **Floret MYB28 Expression** | | | | | |
| **Tukey's multiple comparisons test** | **Predicted (LS) mean diff.** | **95.00% CI of diff.** | **Significant?** | **Summary** | **Adjusted P Value** |
| **Year 1** | | | | | |
| MYB28b/b vs. MYB28v/b | -0.01879 | -0.06298 to 0.02541 | No | ns | 0.5629 |
| MYB28b/b vs. MYB28v/v | -0.1012 | -0.1454 to -0.05705 | Yes | **** | <0.0001 |
| MYB28v/b vs. MYB28v/v | -0.08245 | -0.1266 to -0.03826 | Yes | *** | 0.0001 |
| **Year 2** | | | | | |
| MYB28b/b vs. MYB28v/b | -0.07261 | -0.1220 to -0.02320 | Yes | ** | 0.0024 |
| MYB28b/b vs. MYB28v/v | -0.1862 | -0.2357 to -0.1368 | Yes | **** | <0.0001 |
| MYB28v/b vs. MYB28v/v | -0.1136 | -0.1631 to -0.06423 | Yes | **** | <0.0001 |
| **Leaf Glucoraphanin** | | | | | |
| **Tukey's multiple comparisons test** | **Predicted (LS) mean diff.** | **95.00% CI of diff.** | **Significant?** | **Summary** | **Adjusted P Value** |
| **Year 1** | | | | | |
| MYB28b/b vs. MYB28b/v | -10.21 | -15.76 to -4.654 | Yes | *** | 0.0001 |
| MYB28b/b vs. MYB28v/v | -22.05 | -27.76 to -16.35 | Yes | **** | <0.0001 |

| | | | | | |
|---|---|---|---|---|---|
| MYB28b/v vs. MYB28v/v | -11.84 | -17.55 to -6.138 | Yes | **** | <0.0001 |
| **Year 2** | | | | | |
| MYB28b/b vs. MYB28b/v | -1.806 | -7.360 to 3.747 | No | ns | 0.7143 |
| MYB28b/b vs. MYB28v/v | -8.072 | -13.63 to -2.519 | Yes | ** | 0.0027 |
| MYB28b/v vs. MYB28v/v | -6.266 | -11.82 to -0.7122 | Yes | * | 0.0235 |
| **Floret Glucoraphanin** | | | | | |
| **Tukey's multiple comparisons test** | **Predicted (LS) mean diff.** | **95.00% CI of diff.** | **Significant?** | **Summary** | **Adjusted P Value** |
| **Year 1** | | | | | |
| MYB28b/b vs. MYB28b/v | -13.06 | -17.10 to -9.017 | Yes | **** | <0.0001 |
| MYB28b/b vs. MYB28v/v | -29.82 | -33.85 to -25.78 | Yes | **** | <0.0001 |
| MYB28b/v vs. MYB28v/v | -16.76 | -20.80 to -12.72 | Yes | **** | <0.0001 |
| **Year 2** | | | | | |
| MYB28b/b vs. MYB28b/v | -9.95 | -15.66 to -4.237 | Yes | *** | 0.0004 |
| MYB28b/b vs. MYB28v/v | -17.49 | -23.21 to -11.78 | Yes | **** | <0.0001 |
| MYB28b/v vs. MYB28v/v | -7.543 | -13.26 to -1.830 | Yes | ** | 0.0072 |
| **Leaf Glucoiberin** | | | | | |
| **Tukey's multiple comparisons test** | **Predicted (LS) mean diff.** | **95.00% CI of diff.** | **Significant?** | **Summary** | **Adjusted P Value** |
| **Year 1** | | | | | |
| MYB28b/b vs. MYB28b/v | -1.871 | -2.280 to -1.462 | Yes | **** | <0.0001 |
| MYB28b/b vs. MYB28v/v | 0.6958 | 0.2758 to 1.116 | Yes | *** | 0.0006 |
| MYB28b/v vs. MYB28v/v | 2.567 | 2.147 to 2.987 | Yes | **** | <0.0001 |
| **Year 2** | | | | | |
| MYB28b/b vs. MYB28b/v | -0.632 | -1.041 to -0.2232 | Yes | ** | 0.0013 |
| MYB28b/b vs. MYB28v/v | 0.2654 | -0.1434 to 0.6742 | No | ns | 0.2694 |
| MYB28b/v vs. MYB28v/v | 0.8974 | 0.4886 to 1.306 | Yes | **** | <0.0001 |
| **Floret Glucoiberin** | | | | | |

| Tukey's multiple comparisons test | Predicted (LS) mean diff. | 95.00% CI of diff. | Significant? | Summary | Adjusted P Value |
|---|---|---|---|---|---|
| **Year 1** | | | | | |
| MYB28b/b vs. MYB28b/v | -3.301 | -4.044 to -2.558 | Yes | **** | <0.0001 |
| MYB28b/b vs. MYB28v/v | 2.306 | 1.563 to 3.049 | Yes | **** | <0.0001 |
| MYB28b/v vs. MYB28v/v | 5.607 | 4.864 to 6.350 | Yes | **** | <0.0001 |
| **Year 2** | | | | | |
| MYB28b/b vs. MYB28b/v | -2.369 | -3.420 to -1.318 | Yes | **** | <0.0001 |
| MYB28b/b vs. MYB28v/v | 0.7662 | -0.2848 to 1.817 | No | ns | 0.1909 |
| MYB28b/v vs. MYB28v/v | 3.135 | 2.084 to 4.186 | Yes | **** | <0.0001 |
| **Leaf Indolyl Methyl (IND)** | | | | | |
| **Tukey's multiple comparisons test** | **Predicted (LS) mean diff.** | **95.00% CI of diff.** | **Significant?** | **Summary** | **Adjusted P Value** |
| **Year 1** | | | | | |
| MYB28b/b vs. MYB28b/v | 8.644 | 6.598 to 10.69 | Yes | **** | <0.0001 |
| MYB28b/b vs. MYB28v/v | 15.54 | 13.44 to 17.64 | Yes | **** | <0.0001 |
| MYB28b/v vs. MYB28v/v | 6.894 | 4.791 to 8.996 | Yes | **** | <0.0001 |
| **Year 2** | | | | | |
| MYB28b/b vs. MYB28b/v | 0.3804 | -1.666 to 2.427 | No | ns | 0.8954 |
| MYB28b/b vs. MYB28v/v | 0.3894 | -1.657 to 2.436 | No | ns | 0.8907 |
| MYB28b/v vs. MYB28v/v | 0.00895 | -2.037 to 2.055 | No | ns | >0.9999 |
| **Leaf 1-methoxyindolyl methyl (1-MIND)** | | | | | |
| **Tukey's multiple comparisons test** | **Predicted (LS) mean diff.** | **95.00% CI of diff.** | **Significant?** | **Summary** | **Adjusted P Value** |
| **Year 1** | | | | | |
| MYB28b/b vs. MYB28b/v | 0.1285 | 0.04846 to 0.2085 | Yes | *** | 0.0009 |
| MYB28b/b vs. MYB28v/v | 0.334 | 0.2518 to 0.4162 | Yes | **** | <0.0001 |
| MYB28b/v vs. MYB28v/v | 0.2055 | 0.1233 to 0.2878 | Yes | **** | <0.0001 |
| **Year 2** | | | | | |

| | | | | | |
|---|---|---|---|---|---|
| MYB28b/b vs. MYB28b/v | -0.002046 | -0.08208 to 0.07799 | No | ns | 0.9979 |
| MYB28b/b vs. MYB28v/v | 0.04591 | -0.03413 to 0.1259 | No | ns | 0.3569 |
| MYB28b/v vs. MYB28v/v | 0.04796 | -0.03208 to 0.1280 | No | ns | 0.3257 |
| **Leaf 4-methoxyindolyl methyl (4-MIND)** | | | | | |
| **Tukey's multiple comparisons test** | **Predicted (LS) mean diff.** | **95.00% CI of diff.** | **Significant?** | **Summary** | **Adjusted P Value** |
| **Year 1** | | | | | |
| MYB28b/b vs. MYB28b/v | 0.7913 | -0.7647 to 2.347 | No | ns | 0.4432 |
| MYB28b/b vs. MYB28v/v | 0.7539 | -0.8447 to 2.353 | No | ns | 0.4958 |
| MYB28b/v vs. MYB28v/v | -0.03737 | -1.636 to 1.561 | No | ns | 0.9983 |
| **Year 2** | | | | | |
| MYB28b/b vs. MYB28b/v | 0.1853 | -1.371 to 1.741 | No | ns | 0.9556 |
| MYB28b/b vs. MYB28v/v | 0.1177 | -1.438 to 1.674 | No | ns | 0.9818 |
| MYB28b/v vs. MYB28v/v | -0.06754 | -1.624 to 1.488 | No | ns | 0.994 |
| **Floret Indolyl Methyl (IND)** | | | | | |
| **Tukey's multiple comparisons test** | **Predicted (LS) mean diff.** | **95.00% CI of diff.** | **Significant?** | **Summary** | **Adjusted P Value** |
| **Year 1** | | | | | |
| MYB28b/b vs. MYB28b/v | 0.6335 | -0.3281 to 1.595 | No | ns | 0.2556 |
| MYB28b/b vs. MYB28v/v | 1.022 | 0.06008 to 1.983 | Yes | * | 0.0352 |
| MYB28b/v vs. MYB28v/v | 0.3882 | -0.5734 to 1.350 | No | ns | 0.5915 |
| **Year 2** | | | | | |
| MYB28b/b vs. MYB28b/v | 0.8719 | -0.4880 to 2.232 | No | ns | 0.274 |
| MYB28b/b vs. MYB28v/v | 0.2351 | -1.125 to 1.595 | No | ns | 0.9071 |
| MYB28b/v vs. MYB28v/v | -0.6368 | -1.997 to 0.7230 | No | ns | 0.4952 |
| **Floret 1-methoxyindolyl methyl (1-MIND)** | | | | | |
| **Tukey's multiple comparisons test** | **Predicted (LS) mean diff.** | **95.00% CI of diff.** | **Significant?** | **Summary** | **Adjusted P Value** |
| **Year 1** | | | | | |

| | | | | | |
|---|---|---|---|---|---|
| MYB28b/b vs. MYB28b/v | -0.13 | -0.3425 to 0.08256 | No | ns | 0.3067 |
| MYB28b/b vs. MYB28v/v | 0.2374 | 0.02490 to 0.4500 | Yes | * | 0.0256 |
| MYB28b/v vs. MYB28v/v | 0.3674 | 0.1549 to 0.5800 | Yes | *** | 0.0004 |
| **Year 2** | | | | | |
| MYB28b/b vs. MYB28b/v | -0.01168 | -0.3123 to 0.2889 | No | ns | 0.9951 |
| MYB28b/b vs. MYB28v/v | 0.1305 | -0.1701 to 0.4311 | No | ns | 0.5457 |
| MYB28b/v vs. MYB28v/v | 0.1422 | -0.1584 to 0.4427 | No | ns | 0.4884 |
| **Floret 4-methoxyindolyl methyl (4-MIND)** | | | | | |
| **Tukey's multiple comparisons test** | **Predicted (LS) mean diff.** | **95.00% CI of diff.** | **Significant?** | **Summary** | **Adjusted P Value** |
| **Year 1** | | | | | |
| MYB28b/b vs. MYB28b/v | 0.3124 | -3.161 to 3.786 | No | ns | 0.9739 |
| MYB28b/b vs. MYB28v/v | -3.339 | -6.812 to 0.1341 | No | ns | 0.0617 |
| MYB28b/v vs. MYB28v/v | -3.651 | -7.125 to -0.1782 | Yes | * | 0.0375 |
| **Year 2** | | | | | |
| MYB28b/b vs. MYB28b/v | 5.318 | 0.4063 to 10.23 | Yes | * | 0.0313 |
| MYB28b/b vs. MYB28v/v | 1.995 | -2.917 to 6.907 | No | ns | 0.5877 |
| MYB28b/v vs. MYB28v/v | -3.323 | -8.235 to 1.589 | No | ns | 0.238 |
| **Leaf Sulphate** | | | | | |
| **Tukey's multiple comparisons test** | **Predicted (LS) mean diff.** | **95.00% CI of diff.** | **Significant?** | **Summary** | **Adjusted P Value** |
| **Year 1** | | | | | |
| MYB28b/b vs. MYB28b/v | -30.44 | -85.95 to 25.07 | No | ns | 0.3892 |
| MYB28b/b vs. MYB28v/v | -50.54 | -107.6 to 6.491 | No | ns | 0.0921 |
| MYB28b/v vs. MYB28v/v | -20.1 | -77.13 to 36.93 | No | ns | 0.674 |
| **Year 2** | | | | | |
| MYB28b/b vs. MYB28b/v | 191.1 | 135.6 to 246.6 | Yes | **** | <0.0001 |
| MYB28b/b vs. MYB28v/v | 437.2 | 381.7 to 492.7 | Yes | **** | <0.0001 |

| | | | | | |
|---|---|---|---|---|---|
| MYB28b/v vs. MYB28v/v | 246 | 190.5 to 301.6 | Yes | **** | <0.0001 |
| **Floret Sulphate** | | | | | |
| **Tukey's multiple comparisons test** | **Predicted (LS) mean diff.** | **95.00% CI of diff.** | **Significant?** | **Summary** | **Adjusted P Value** |
| **Year 1** | | | | | |
| MYB28b/b vs. MYB28b/v | -11.61 | -27.60 to 4.391 | No | ns | 0.1938 |
| MYB28b/b vs. MYB28v/v | -2.003 | -18.00 to 13.99 | No | ns | 0.9501 |
| MYB28b/v vs. MYB28v/v | 9.603 | -6.394 to 25.60 | No | ns | 0.3198 |
| **Year 2** | | | | | |
| MYB28b/b vs. MYB28b/v | 1.476 | -21.15 to 24.10 | No | ns | 0.9862 |
| MYB28b/b vs. MYB28v/v | -6.67 | -29.29 to 15.95 | No | ns | 0.7542 |
| MYB28b/v vs. MYB28v/v | -8.146 | -30.77 to 14.48 | No | ns | 0.6577 |
| **Leaf SMCSO** | | | | | |
| **Tukey's multiple comparisons test** | **Predicted (LS) mean diff.** | **95.00% CI of diff.** | **Significant?** | **Summary** | **Adjusted P Value** |
| **Year 1** | | | | | |
| MYB28b/b vs. MYB28b/v | 0.07463 | -7.146 to 7.295 | No | ns | 0.9997 |
| MYB28b/b vs. MYB28v/v | 0.1621 | -7.256 to 7.580 | No | ns | 0.9985 |
| MYB28b/v vs. MYB28v/v | 0.08745 | -7.331 to 7.506 | No | ns | 0.9996 |
| **Year 2** | | | | | |
| MYB28b/b vs. MYB28b/v | 1.048 | -6.173 to 8.268 | No | ns | 0.9348 |
| MYB28b/b vs. MYB28v/v | -9.451 | -16.67 to -2.231 | Yes | ** | 0.0073 |
| MYB28b/v vs. MYB28v/v | -10.5 | -17.72 to -3.278 | Yes | ** | 0.0026 |
| **Floret SMCSO** | | | | | |
| **Tukey's multiple comparisons test** | **Predicted (LS) mean diff.** | **95.00% CI of diff.** | **Significant?** | **Summary** | **Adjusted P Value** |
| **Year 1** | | | | | |
| MYB28b/b vs. MYB28b/v | 19.55 | 1.005 to 38.10 | Yes | * | 0.0369 |
| MYB28b/b vs. MYB28v/v | 20.5 | 1.947 to 39.05 | Yes | * | 0.0275 |

| | | | | | |
|---|---|---|---|---|---|
| MYB28b/v vs. MYB28v/v | 0.9422 | -17.61 to 19.49 | No | ns | 0.9916 |
| **Year 2** | | | | | |
| MYB28b/b vs. MYB28b/v | -8.52 | -34.75 to 17.71 | No | ns | 0.7106 |
| MYB28b/b vs. MYB28v/v | -1.74 | -27.97 to 24.49 | No | ns | 0.9857 |
| MYB28b/v vs. MYB28v/v | 6.779 | -19.45 to 33.01 | No | ns | 0.8048 |
| **Floret Total Sulphur** | | | | | |
| **Tukey's multiple comparisons test** | **Predicted (LS) mean diff.** | **95.00% CI of diff.** | **Significant?** | **Summary** | **Adjusted P Value** |
| **Year 1** | | | | | |
| MYB28b/b vs. MYB28b/v | -33.47 | -109.7 to 42.80 | No | ns | 0.4231 |
| MYB28b/b vs. MYB28v/v | -70.17 | -146.4 to 6.100 | No | ns | 0.0678 |
| MYB28b/v vs. MYB28v/v | -36.7 | -113.0 to 39.57 | No | ns | 0.3654 |
| **Year 2** | | | | | |
| MYB28b/b vs. MYB28b/v | -72.15 | -152.6 to 8.354 | No | ns | 0.0744 |
| MYB28b/b vs. MYB28v/v | -96.16 | -176.7 to -15.66 | Yes | * | 0.0245 |
| MYB28b/v vs. MYB28v/v | -24.01 | -104.5 to 56.49 | No | ns | 0.6513 |
| **Floret Cysteine** | | | | | |
| **Tukey's multiple comparisons test** | **Predicted (LS) mean diff.** | **95.00% CI of diff.** | **Significant?** | **Summary** | **Adjusted P Value** |
| **Year 1** | | | | | |
| MYB28b/b vs. MYB28b/v | 1.211 | -3.579 to 6.000 | No | ns | 0.7824 |
| MYB28b/b vs. MYB28v/v | 0.3026 | -4.486 to 5.092 | No | ns | 0.9845 |
| MYB28b/v vs. MYB28v/v | -0.9079 | -5.697 to 3.881 | No | ns | 0.87 |
| **Year 2** | | | | | |
| MYB28b/b vs. MYB28b/v | 2.063 | -3.341 to 7.468 | No | ns | 0.5798 |
| MYB28b/b vs. MYB28v/v | 3.136 | -2.268 to 8.541 | No | ns | 0.3045 |
| MYB28b/v vs. MYB28v/v | 1.073 | -4.332 to 6.478 | No | ns | 0.8585 |
| **Floret Methionine** | | | | | |

| Tukey's multiple comparisons test | Predicted (LS) mean diff. | 95.00% CI of diff. | Significant? | Summary | Adjusted P Value |
|---|---|---|---|---|---|
| **Year 1** | | | | | |
| MYB28b/b vs. MYB28b/v | 1.363 | -3.426 to 6.152 | No | ns | 0.734 |
| MYB28b/b vs. MYB28v/v | -0.6032 | -5.392 to 4.186 | No | ns | 0.94 |
| MYB28b/v vs. MYB28v/v | -1.966 | -6.755 to 2.823 | No | ns | 0.5349 |
| **Year 2** | | | | | |
| MYB28b/b vs. MYB28b/v | 3.44 | -1.964 to 8.845 | No | ns | 0.2457 |
| MYB28b/b vs. MYB28v/v | 3.641 | -1.763 to 9.046 | No | ns | 0.2119 |
| MYB28b/v vs. MYB28v/v | 0.2011 | -5.204 to 5.606 | No | ns | 0.9946 |
